# Abiotic drivers of protein abundance variation among natural populations

**DOI:** 10.1101/2020.03.27.011676

**Authors:** Joshua Niklas Ebner, Danilo Ritz, Stefanie von Fumetti

## Abstract

Identifying when and where environmental change induces molecular responses in natural populations is an important goal in contemporary ecology. It can aid in identifying molecular signatures of populations experiencing stressful conditions and potentially inform if species are approaching the limits of their tolerance niches. Achieving this goal is hampered by our limited understanding of the influence of environmental variation on the molecular systems of most ecologically relevant species as the pathways underlying fitness-affecting plastic responses have primarily been studied in model organisms under controlled laboratory conditions. In this study, we establish relationships between protein abundance patterns and the abiotic environment by profiling the proteomes of 24 natural populations of the caddisfly *Crunoecia irrorata.* We subsequently relate these profiles to natural variations in the abiotic characteristics of their freshwater spring habitats which shows that protein abundances and networks respond to abiotic variation according to the functional roles these proteins have. We provide evidence that geographic and past and present environmental differences between sites affect protein abundances and identifications, and that baseline reaction norms are ubiquitous and can be used as information rather than noise in comparative field studies. Taking this natural variation into account is a prerequisite if we are to identify the effects environmental change has on natural populations.

## 1. Introduction

By continuously sensing and integrating environmental cues, organisms are able to maintain fitness via physiological and phenotypic plasticity^1–3^. These responses allow individuals to adjust to environmental variation, a feat that directly influences the outcome of evolution and the likelihood of extinction^4–6^. Due to the reversible nature of plasticity and its contribution to fitness, it has the potential to moderate the loss of global biodiversity during environmental change^7–10^. The advantage of species’ plasticity during such change depends on the threshold-limits of their more- or less robust molecular systems that underlie plastic responses^11,12^. In many, if not all species, these networks of interacting macromolecules are likely tuned to the fluctuations in the environment over space and time, one reason why substantial variation in gene expression and protein abundance profiles within and between populations can be observed^13–18^. Attributing this variation to environmental cues experienced by the organism can not only give insights into the molecular mechanisms underlying regulatory plastic responses^19–21^ but also inform about population health and how species at one site compensate for environmental change not encountered at another site. For most species, the cues and underlying molecular pathways involved are largely unknown but increasingly studied by identifying large-scale expression changes, providing fundamental knowledge on adaptive processes in response to environmental challenges such as global warming and pollution^22–25^. It has been repeatedly shown that the correlation between variation in messenger RNA expression and protein abundances is modest^26,27^. Phenomena such as pre, co- and post-transcriptional and translational modification^28,29^, protein turnover^30^ and the inherently stochastic nature of gene expression contribute to this pronounced difference^31,32^. Consequently, the same amount of protein can be formed with up to 30-fold variation of the mRNA level, and with the same mRNA level, the amount of protein formed can vary up to 20-fold^26^. Taken together, the molecular signatures of plasticity triggered by environmental (selective) pressure might frequently only be detectable on the protein level and against a pre-established background of environmentally induced molecular variation. To this end, protein analysis represents an essential complement for studying the physiological consequences of gene expression variability. Furthermore, protein activity, stability and abundance are essential to plastic responses due to their close connection to the organismal phenotype and physiological processes^33–35^. Yet, our knowledge on the concrete environmental cues that induce protein abundance variation among natural populations is limited due to a lack of field-based studies that relate abundance variation to variation in environmental conditions.

In this study, we proposed to tackle two questions. First, what is the standing protein abundance variation between natural populations? Second, are differences in the abiotic environment linked to this variation and if so, do functional profiles of proteins reflect these differences? To this end, 24 populations of the caddisfly *Crunoecia irrorata* (Trichoptera: Lepidostomatidae, Curtis 1834) were used as a study system for two reasons: i) The increasing need to monitor population health of non-model species (i.e. species without sequenced genomes) via systems-wide approaches and ii) the continuous exposure of larval individuals to surrounding aqueous conditions. Population-wide liquid chromatography-tandem mass spectrometry (LC-MS/MS) was used to examine the patterns of protein identification and abundance variation of this species, and we tested the hypothesis that this variation relates systematically to abiotic variables including elevation, geographic distance and past climatic conditions. As cellular conditions change rapidly, often in seconds or minutes^36,37^, functional physiological changes must necessarily occur before or during transcriptional and translational changes. We therefore focus on abiotic point-measurements, i.e. we measured abiotic variables at the same time as organismal sampling to obtain a “physiological snapshot” of these natural populations. Such a characterization of the “normal state” of a given system - the ranges of its variation^38^-could aid in defining baselines of molecular responses that must be exceeded in order to be considered potentially adaptive changes or physiological responses triggered by environmental change.

## 2. Material and Methods

### 2.1. Organism collection and environmental variables

Eight *C. irrorata* larval individuals of similar sizes were collected from 27 (near-)natural freshwater springs, i.e. from 9 springs in three study regions across Germany: Rhoen Biosphere Reserve (R), Harz National Park (H) and Black Forest (BF) (**SI1 Table 1**). Sites were distributed in a randomized design between low (275 m) and high (967 m) elevations. Individuals from one spring were pooled into one sample since pooling has been shown to match mean protein abundances of the individuals making up the pool^39^, thereby reducing the biological variance compared with that between individuals^40^. After identification, larvae were washed with dH_2_O and transferred on-site into 5.0 ml Protein LoBind tubes (Eppendorf, Germany) containing 1 ml of RNAlater® (Invitrogen™). Proteins of various tissues are well preserved in RNAlater®, and biological information is largely kept intact^41–43^. Samples were stored at ~4 °C in a portable fridge (Dometic TropiCool TC 33) until storage at −20 °C was possible. From each spring, additional larvae were collected, conserved in EtOH and their head capsule widths (*n* = 88) were measured using a Leica S9i stereo microscope, serving as a proxy control for possible instar stage distribution differences between sampling regions. Abiotic variables were measured at each spring using a multi-parameter portable meter (MultiLine^®^ Multi 3650 IDS). Inorganic nutrient contents were determined from 30 ml of spring water via Ion Chromatography (IC) using a 940 Professional IC Vario ONE/SeS/PP (Metrohm) and iron (Fe) content via Inductively Coupled Plasma Emission Spectrometry (ICP-OES) using a 5100 ICP-OES (Agilent Technologies) (**SI2, SI1 Figure 1**). In order to assess the influence of past climatic differences between sites on protein abundance profiles, we collected macroclimate data (BioClim1-19) from the WorldClim v.2^44^ database based on spring coordinates using the raster package v.3.0.12^45^ (resolution: 5-minute of a longitude/latitude degree), presenting various indices of environmental data calculated from 30 years of average monthly data (1971-2000).

### 2.2. Protein extraction

Prior to protein extraction, samples were thawed on ice and centrifuged at 5,000 rpm. RNA-later was decanted and larvae transferred from RNAlater^®^ into 1.5 ml protein LoBind tubes containing 1 ml cold 10x phosphate-buffered saline (PBS). After vortexing for 10 s, larvae were transferred into 1.5 ml protein LoBind tubes containing 400 μl cold lysis buffer (1% sodium deoxycholate [SDC], 10 mM TCEP, 100 mM Tris, pH 8.5 [adjusted with NaOH/HCL]). Samples were in-tube homogenized with a sterile glass pestle followed by 10 min incubation at 4 °C and 3 x 1 ultrasonication (Bandelin Sonoplus HD270). Samples were spun at 5.000 rpm for 10 min at −4 °C, and 200 μl supernatant was subsequently transferred to a new tube. To desalt absorbed RNAlater^®^, trichloroacetic acid (TCA)/acetone precipitation was performed according to the protocol by Luis Sanchez^47^, resulting in a pellet that was dissolved in 200 μl lysis buffer. After 10×1 s sonication, samples were incubated for 10 min at 95 °C at 300 rpm in a Thermomixer C (PCR 96 heating block, Eppendorf). At this point, protein concentrations were determined via a Bicinchoninic acid assay (BCA; Thermo Scientific™ Pierce™ BCA Protein Assay Kit) according to manufacturer’s instructions (**SI1 Table 2**). After letting samples cool down at room temperature, they were spun down at 5,000 rpm for 10 s. Four μl of 0.75 M chloroacetamide solution was added to each sample and incubated at 37 °C for 10 min at 500 rpm and again spun down at 5.000 rpm for 10 s. After checking if the pH of each sample was around 8, 1 μg trypsin (Sequencing Grade Modified Trypsin, Promega) was added to 50 μg extracted proteins per sample which then were digested overnight at 37 °C and 300 rpm. Samples were acidified with 50 μl 5% trifluoroacetic acid (TFA) and peptides purified using PreOmics cartridges (Martinsried) according to manufacturer’s instructions. Eluted peptides were transferred to a 96-well plate and concentrated to dryness by applying vacuum for 2 hr. Peptides were subsequently dissolved in 20 μl 0.1% formic acid by 10 x 1 s ultrasonication and shaking at 14.000 rpm at 25 °C for 5 min. After spinning dissolved peptides down at 4.000 rpm for 10 min, peptide concentrations were determined based on absorbance values using a SPECTROstar Nano Absorbance Plate Reader (BMG Labtech) (**SI1 Table 2**). Peptides were diluted to a concentration of 0.5 μg/μl in 0.1% formic acid. IRT peptides (Biognosys AG, Schlieren, Switzerland) were added to the wells to control for LC-MS performance, and samples were stored at −20 °C prior to LC-MS/MS analysis.

### 2.3. LC-MS/MS analysis

Samples were subjected to LC-MS/MS analysis using an Orbitrap Fusion Lumos Tribrid Mass Spectrometer fitted with an EASY-nLC 1200 (both Thermo Fisher Scientific) and a custom-made column heater set to 60 °C. Peptides were resolved using an RP-HPLC column (75 μm x 36 cm) packed in-house with C18 resin (ReproSil-Pur C18-AQ, 1.9 μm resin; Dr. Maisch GmbH) at a flow rate of 0.2 μl/min. The following gradient was used for peptide separation: from 5% B to 12% B over 5 min to 35% B over 65 min to 50% B over 20 min to 95% B over 2 min followed by 18 min at 95% B. Buffer A was 0.1% formic acid in water, and buffer B was 80% acetonitrile, 0.1% formic acid in water. The mass spectrometer was operated in Data-Dependent Acquisition (DDA) mode with a cycle time of 3 s between master scans. Each master scan was acquired in the Orbitrap at a resolution of 120.000 full width at half maximum (at 200 m/z, MS1) and a scan range from 375 to 1,600 m/z followed by MS/MS (MS2) scans of the most intense precursors in the linear ion trap at “Rapid” scan rate with isolation of the quadrupole set to 1.4 m/z. Maximum ion injection time was set to 50 ms (MS1) and 35 ms (MS2) with an AGC target of 1.0E6 and 1.0E4, respectively. Monoisotopic precursor selection (MIPS) was set to peptide, and the intensity threshold was set to 5.0E3. Peptides were fragmented by HCD (higher-energy collisional dissociation) with collision energy set to 35%, and one microscan was acquired for each spectrum. The dynamic exclusion duration was set to 30s.

### 2.4. Protein Identification

Raw spectra (Thermo raw files) were submitted to an Andromeda^46^ search in MaxQuant v1.6.10.43^47^. The “match between runs” option was enabled (match time window: 0.7 min, alignment time window: 20 min). Instrument type was set to Orbitrap, precursor mass tolerance to 15 ppm and fragment ion tolerance to 0.05 Da. Enzyme specificity was set to fully tryptic, with a maximum of two missed cleavages. MS/MS spectra were searched against a previously described database consisting of translated gene-prediction sequences of Trichoptera species^48^, additionally including candidate protein-coding regions of *Rhyacophila fasciata*^49^; BioProject: PRJNA219600). All searches included a contaminants database (as implemented in MaxQuant, 267 sequences). For protein identification, unique and razor peptides were used. The peptide spectrum-match false discovery rate (FDR) and the protein FDR were set to 0.01 (based on the target-decoy approach). Oxidation of methionine (M) and acetylation (Protein N-term) were specified as variable and carbamidomethylation of cysteines (C) as fixed modifications. Enzyme specificity was set to “Trypsin/P”. Minimum peptide length was set to 7. The “evidence.txt” and “summary.txt” output files were used for LC-MS/MS quality control using R package artMS v.1.4.2^50^ (results are given in **SI3)**. To investigate identification differences between the three regions, we analyzed samples in 3 independent MaxQuant runs (settings as above). Overlapping and unique Majority protein IDs (IDs of those proteins that have at least half of the peptides that the leading protein has) were compared via a Venn diagram plotted via package VennDiagram v.1.6.20^51^.

### 2.5. Statistical analysis

Data analysis was conducted in R v.3.6.2^52^. Strongly correlating (Pearson’s R^2^ > 0.7) abiotic and BioClim variables were identified with function cor.test and removed from further analysis, resulting in 11 abiotic and 7 BioClim variables kept for analysis (**SI2, SI1 Figure 2)**. The in MaxQuant implemented label-free quantification (LFQ) option computed 54.2% missing values (**SI1 Figure 3**), making data imputation unreliable. Therefore, protein raw intensities were calculated through summation of peptide intensities^53^ (only proteins with *n* = 2 unique peptides). Raw intensity values were log_2_-transformed, quantile normalization was performed using function normalize.quantiles from package preprocessCore v.1.46.0^54^ and missing values (*n* = 1512 = 5.37%) imputed using function impute.knn from package impute v.1.60.0^55^. These relative abundance values were tested for association with abiotic variables: First, unsupervised non-metric multidimensional scaling (nMDS) ordination was performed to visualize clustering of protein abundances between populations. A Bray-Curtis^56^ distance matrix was calculated based on relative protein abundance values for each population and used as input to the metaMDS function implemented in vegan v.2.5.6^57^ (stress-plot: **SI1 Figure 4**). Weighted correlation network analysis (WGCNA) v.1.68^58^ was used to identify suites of co-regulated proteins in an unsupervised way. These protein networks could identify important functional groups of proteins and candidate proteins underlying plastic responses, similar to gene expression analyses^59,60^. A sample network was constructed to identify outlying samples with a standardized connectivity score of less than −2.5^61^. A signed protein co-abundance network was constructed for all 24 populations independent of sampling region. The network was constructed with a soft threshold power (β) of 18 as this value was found to be appropriate by the function pickSoftThreshold to reach a scale-free topology index (R^2^) of at least 0.90. We used the Dynamic Tree Cut approach to merge highly correlated modules using a height-cut of 0.20^62^. The abundances of these modules can be summarized as the abundance of a single “eigengene”, calculated as the first principal component of the abundances of all proteins in a module across samples^62^. These eigengenes were correlated (Pearson’s correlation) with abiotic variables to identify module-environment (**SI1 Figure 5)**. Analogous to the *in situ* abiotic variables, a WGCNA was independently performed for bioclimatic variables (WGCNA_BC_, **SI1 Figure 6**). Differentially abundant proteins (DAPs) between the three sampling regions were identified using DAPAR v.1.16.11 implemented in ProStaR v.1.16.10^63^. Used functions, diagnostic plots and results of this analysis are given in **SI4**. Norms of reaction for individual proteins were assessed by estimating Bayesian generalized linear models (GLMs) for continuous response variables using the stan_glm function in the rstanarm package v2.17.4^64^. Each protein was assessed independently via scatterplots. To assess the effect of geographic distance on protein abundance divergence between populations, a Mantel test was performed between geographic- and (Euclidean) protein-distance matrices in ade4 v.1.7.13^65^. Geographic distances between springs were calculated based on longitude and latitude of springs using package geosphere v.1.5.10^66^ and the implemented function distm (Vincentry great circle distances). Pairwise spectrum distances (cosine distances) between MS/MS runs (proteome-wide distances independent of peptide-identification and therefore of the homology-based protein database) were calculated using the DISMS2 algorithm^67^.

### 2.6. Functional annotation of protein sequences

The first majority protein ID of each identified protein was parsed from the MaxQuant output and corresponding amino-acid sequences extracted from the database using function ucsc-faSomeRecords as implemented in Bioconda^68^. All sequences were queried against the Swiss-Prot database^69^ (accessed 10 January 2020) using stand-alone blastp v.2.2.28 with default parameters (E-value 0.001). Protein domains were determined using stand-alone InterProScan (IPS) v.5.39-77.0^70^ with default parameters. All proteins were assigned to existing Cluster of Orthologous Groups (COGs)^71^ and Gene Ontology (GO) terms in EggNOG v5.0^72^ via eggNOG-Mapper^73^ (Taxonomic scope: Insecta, Orthologs: all orthologs, GO evidence: non-electronic terms, E-value: 0.001). To test whether any GO terms were overrepresented in WGCNA modules and DAPs, we sorted proteins in selected modules by their Gene Significance (GS) for the significantly correlated abiotic variable and DAPs by their adjusted *p*-values and performed rank-based tests for each GO term assigned to these proteins by applying Kolmogorov-Smirnov tests via package topGO v.2.38.1^74^ (“weight01” algorithm, nodeSize = 10). To further identify and substantiate biological functions and processes associated with modules, a domain-based analysis was conducted. To this end, the IPS Protein Families (Pfam^75^) annotations of the proteins belonging to a certain module were used as entry for domain-centric GO (dcGO) enrichment (FDR *p*-value < 0.01)^76^.

## 3. Results

### 3.1. Global drivers of protein abundance differentiation

Mass spectra matched to 4536 distinct peptide sequences (mean number of amino acids: 16, range: 8-44). These peptide sequences mapped to 1173 proteins that were commonly identified in all 24 *C. irrorata* populations. Protein identifications were largely overlapping (**SI1 Figure 7b**) and pairwise peptide-level correlations between samples were above 0.8 (**SI3**). Protein abundance profiles exhibited variation between populations but showed association with sampling regions, whereby populations from any one region clustered together mostly along axis 2 of the nMDS (**Figure 1a**). Therefore, proteins with significant, positive eigenvector loadings along this axis may play a dominant role in proteome differentiation over geographic range. The top proteins with positive loadings (> 0.005, *n* = 69) were enriched for GO terms *small molecule metabolic process* (GO:0044281, BP), *oxireductase activity* (GO:0016491, MF) and *mitochondrion* (GO:0005739, CC). More similar protein abundance profiles were found between populations occupying adjacent springs when compared to springs further apart (*p*_Mantel_ = 0.006, **SI1 Figure 8**) and pairwise spectrum distances between LC-MS/MS runs were positively correlated with environmental distances between springs (*p*_Mantel_ = 0.009), whereby spectral profiles were more similar between abiotically more similar springs (**Figure 1b**). Environmental differences between springs did not correlate with geographic distance (*p* = 0.33; **SI1 Figure 9**).

**Figure 1.**
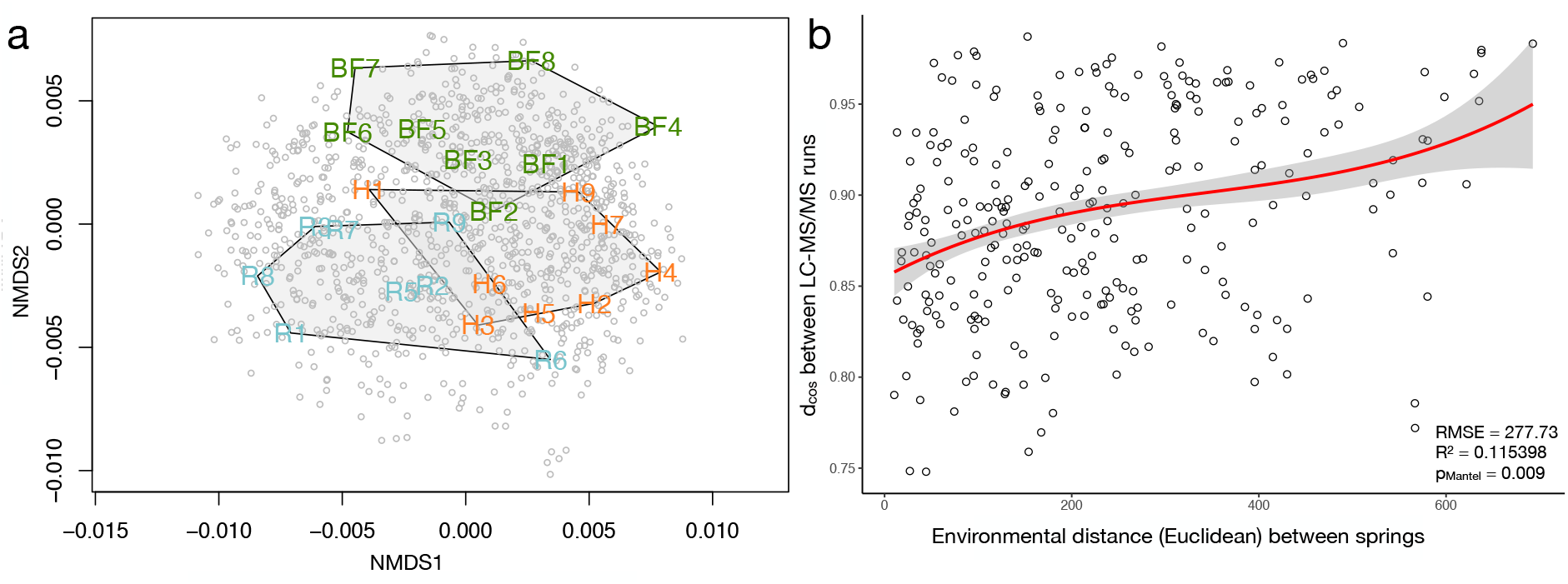
Ordination and spectral distance results. (**a**) Non-Metric Multi-Dimensional Scaling (nMDS) based on abundances of proteins identified in all populations illustrating the similarities and differences in abundance profiles of populations across three sampling regions. Relative proximity of sample-labels represents overall degree of abundance similarity between populations and grey points represent single proteins. (**b**) Correlation between environmental distances between springs and spectral distances between population-wide LC-MS/MS runs.

Ninety-four DAPs were identified between H and BF (FDR = 1.34%), 116 between R and BF (FDR = 1.44%) and 107 between R and H (FDR = 1.12%) (**SI2**). These DAPs were predominantly enriched in GO terms related to cellular-level process [e.g., *(negative) regulation of cellular processes* (GO:0048523; GO:0050794), *regulation of cellular component assembly* (GO:0051128) and *cellular localization* (GO:0051641)]. Protein families consistently showing the highest frequencies (> *n* = 10) across the full set of DAPs were J: *translation, ribosomal structure and biogenesis* and O: *post-translational modification/protein turnover, and chaperones* (**Figure 2a-c**).

**Figure 2.**
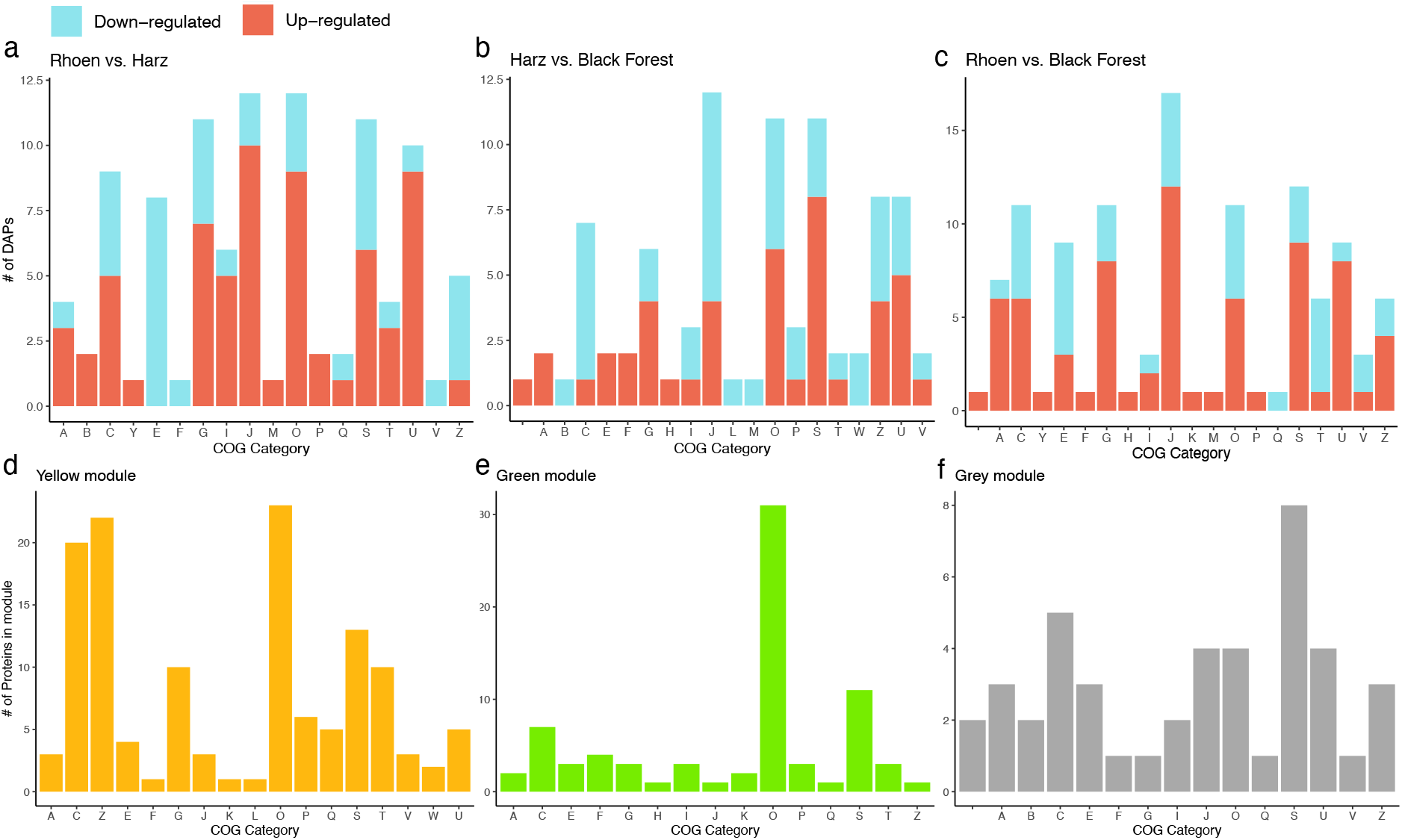
Number of up- and down-regulated DAPs between sampling regions (red and blue, respectively) and module member proteins (yellow, green and grey) regarding their Cluster of Orthologous Group (COG) protein family annotation. (**a**) Rhoen vs. Harz. (**b**) Harz vs. Black Forest. (**c**) Rhoen vs. Black Forest. (**d**) “Yellow” module. (**e**) “Green” module. (**f**) “Grey” module. One-letter abbreviations for the functional categories: A, RNA processing and modification; B, chromatin structure and dynamics; C, energy production and conversion; Y, nuclear structure; E, amino acid transport and metabolism; F, nucleotide transport and metabolism; G, carbohydrate transport and metabolism; H, coenzyme transport and metabolism; I, lipid transport and metabolism; J, translation, ribosomal structure and biogenesis; K, transcription; L, translation, ribosomal structure and biogenesis; M, cell wall/membrane/envelope biogenesis; O, post-translational modification, protein turnover, and chaperones; P, inorganic ion transport and metabolism; W, extracellular structures; Q, secondary metabolites biosynthesis, transport, and catabolism; S, unknown function; T, signal transduction mechanisms; U, intracellular trafficking, secretion, and vesicular transport; V, defense mechanisms; Z, cytoskeleton.

### 3.2. Protein reaction norms

We observed pronounced baseline abundance differences between geographical regions (examples: **Figure 3**): For example, the reaction norm of phosphoglycerate mutase, a pH-sensitive enzyme^77,78^, had a higher baseline abundance in populations from BF springs which had significantly higher mean pH than springs from R and H (**SI1 Figure 10a**). Multiple proteins showed pronounced abundance changes in relation to abiotic conditions. For example, 13 proteins showed increasing and 20 decreasing reaction norms according to spring elevation and pH variation was accompanied by 71 increasing and 5 decreasing norms of reaction. Temperature variation was accompanied by 29 increasing and 33 decreasing reaction norms whereby a high frequency of decreasing proteins belonged to the chaperonin Cpn60/TCP-1 family and a high frequency of increasing proteins to the calreticulin/calnexin family (**SI1 Figure 16**). Both families play a role in the physiological integration of temperature variation, both in the cold^79,80^ and warmth^81^.

**Figure 3.**
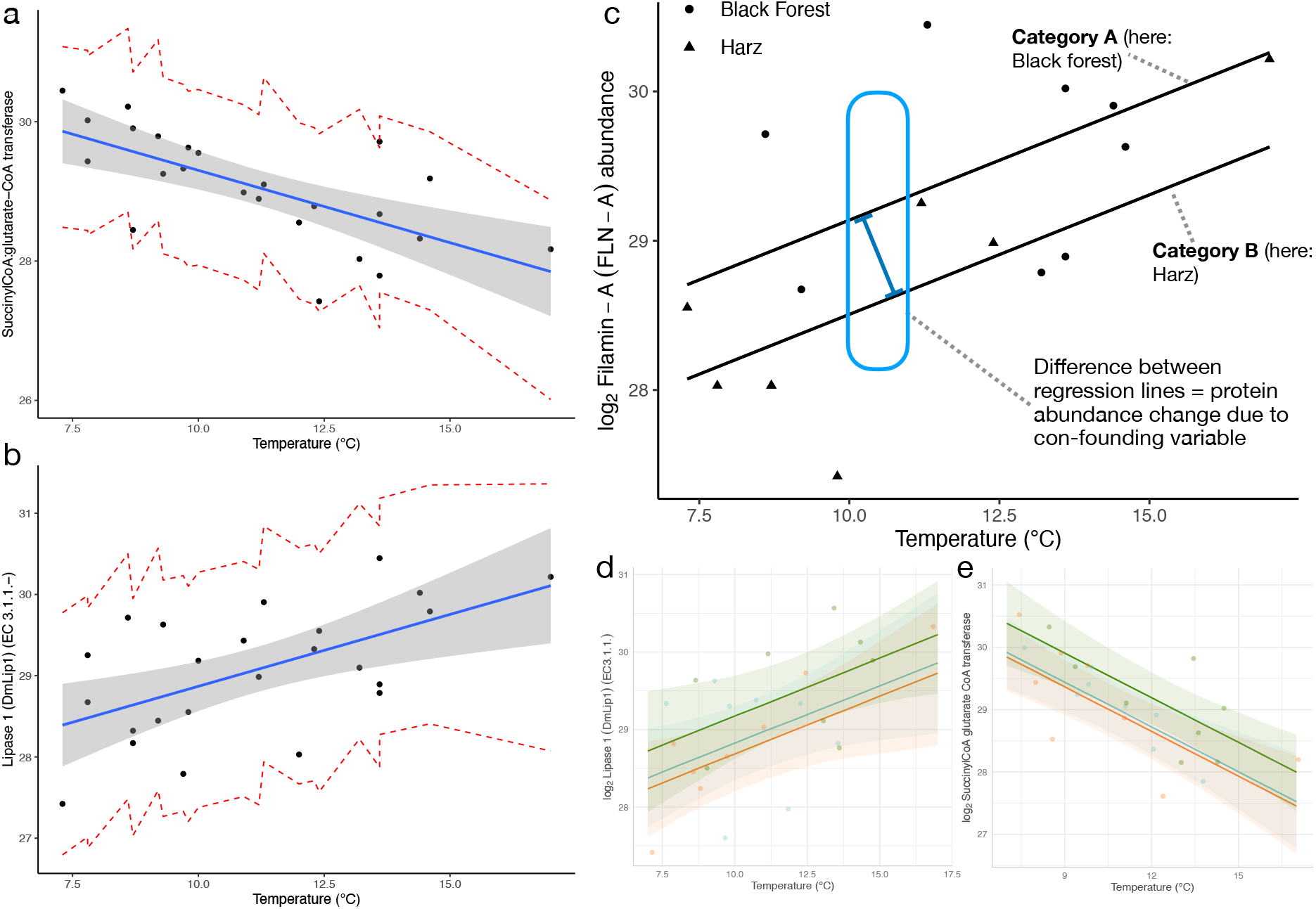
Illustration and concept of protein reaction norms and baseline differences. Protein reaction norms are the relative abundance changes of a single protein across a range of environments. (**a**) Regression model and predicted values of Succinyl-CoA-glutarate-CoA transferase in relation to spring temperature (°C). (**b**) Regression model and predicted values of Lipase 1 in relation to spring temperature (°C). (**c**) Reaction norms of filamin A in response to variation in temperature (°C). Category A: E.g. a site experiencing environmental change such as nutrient influx, temperature change or pollution; Category B: A site experiencing no such environmental change. (**d**) Regression as in (**b)** but categorized by sampling region (color-code as in **Figure 1a**). (**e**) Regression as in (**a)** but categorized by sampling region.

### 3.3. Network analyses

The WGCNA assigned all 1173 proteins to eight co-expression modules (designated randomly by colors). Five module eigengenes were significantly correlated with distinct abiotic variables (**Figure 4a**): Both, “brown” (*n* = 477) and “red” (*n* = 59) modules correlated with magnesium (r_brown_ = −0.46, r_red_ = −0.42) and sulfate (r_brown_ = −0.43, r_red_ = −0.49). “Green” (*n* = 81), “yellow” (*n* = 133) and “grey” (*n* = 73) modules correlated positively with oxygen (r_green_ = 0.4), pH (r_yellow_ = 0.47) and nitrate (r_grey_ = 0.47), respectively. Correlations between module membership and protein significance values were used to further corroborate these relationships, with correlations ranging from 0.5 for the “yellow” module to 0.66 for the “grey” module (**Figure 4f-i**). Gene Ontology enrichment of the highly underrepresented “brown” module detected many significant GO terms across all GO categories (Cellular Compartment (CC), Molecular Function (MF) and BF). Gene Ontology terms within BP and MF included many terms associated with unfolded protein stimulus [e.g., *chaperone binding* (GO:0051082), *chaperone-mediated protein folding* (GO:0061077) and *cellular response to topologically incorrect protein* (GO:0035967)]. The “red” module was associated with the same abiotic variables as the “brown” module, but had no enriched GO terms. The “green” module was enriched with four GO terms in the BP category: *Intestinal stem cell homeostasis* (GO:0036335), *sleep* (GO:0030431), *response to endoplasmic reticulum stress* (GO:0034976) and *RNA interference* (GO:0016246) and one GO term in the CC category: *Intracellular membrane-bounded organelle* (GO:0043231). Lastly, the “yellow” module was enriched with only one GO term in the BP category: *Electron transport chain* (GO:0022900) but with three GO terms in the CC category: *Cytoplasmic part* (GO:0044444), *intracellular membrane bounded organelle* (GO:0043231) and *endomembrane system* (GO:0012505) and two GO terms in the MF category: *Hydrolase activity* (GO:0016787) and *catalytic activity acting on a protein* (GO:0140096).

**Figure 4.**
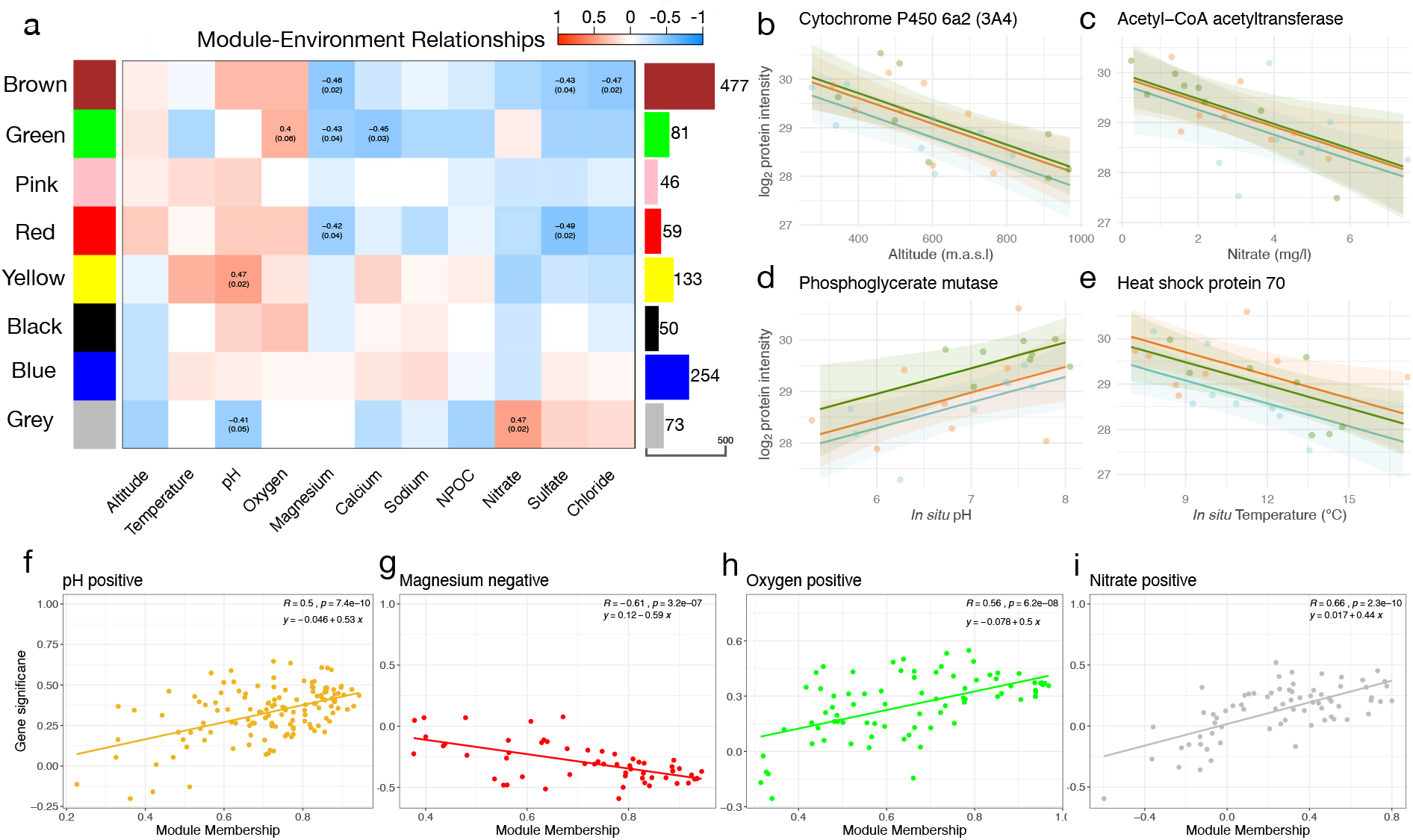
Network analysis of *Crunoecia irrorata* protein abundances in relation to abiotic variables. **(a)** Correlations between module eigengenes (rows) and abiotic variables (columns). The bar graph and numbers on the right indicate number of proteins belonging to each module. The strength of the correlations between abiotic factors and protein co-abundance modules are indicated by color intensity. The numbers in the cells give Pearson’s correlation coefficients between the module “eigengene” and the abiotic factor and the *p*-value of the correlation test (not listed for cells with *p* > 0.05 except “green”-oxygen (*p* = 0.06)). **(b-e)** Scatterplots showing protein reaction norms for four proteins with high gene significance (GS) values in relation to the abiotic variable shown on the x-axis. Reaction norms are colored by sampling region to illustrate baseline differences between sampling regions according to color-scheme in **Fig. 1.(b)** Cytochrome P450 6a2 decreasing in abundance with elevation of springs. **(c)** Acetyl-CoA acetyltransferase abundance decreasing with increasing nitrate concentration of spring water. **(d)** Phosphoglycerate mutase increasing in abundance with increasing *in situ* pH. **(e)** Heat shock protein 70 decreasing in abundance with water temperature. **(f-i)** Scatterplots illustrating the relationship between a protein’s module membership score (x-axis) and the protein’s significance for the abiotic variable (y-axis). Higher correlations between these parameters indicate stronger associations of the **(f)** “yellow”, **(g)** “red”, **(h)** “green” and **(i)** “grey” modules with their associated abiotic variables (pH, magnesium, oxygen and nitrate, respectively).

Domain-centric GO enrichment of the “yellow” module (Pfam_slim_ level = 3) computed significantly enriched GO terms such as *regulation of sodium ion transport* (GO:0002028), *pH reduction* (GO:0045851) and *retina layer formation* (GO:0010842). The “green” module included significant GO terms such as *oxidoreductase activity* (GO:0016491), *oxidation-reduction process* (GO:0055114) and *response to oxidative stress* (GO:0006979). The “grey” module, related negatively to pH and positively to nitrate concentrations was enriched for similar GO terms as the “yellow” module such as *pH reduction* (GO:0045851), *regulation of intracellular pH* (GO:0051453) but also terms related to nitrogen such as *response to xenobiotic stimulus* (GO:0009410) and *cellular nitrogen compound metabolic process* (GO: 0034641). The “yellow” module had higher frequencies of proteins associated with C: *energy production and conversion*, Z: *cytoskeleton* and O: *post-translational modification/protein turnover, and chaperones*. The “green” module included 39.5 % proteins associated with O: *post-translational modification/protein turnover, and chaperones* and the “grey” module showed COG underrepresentation and many proteins with S: *unknown function* (**Figure 3e-f**).

The strongest relationship in the WGCNA_BC_ emerged between the “green_BC_” module (*n* = 214) and bio6 (r = 0.72), representing the minimum temperatures (°C) of the coldest month experienced by *C. irrorata* populations in the corresponding sites over 30 years (**Figure 5**). This module contained a high frequency of proteins related to the cytoskeleton COG family (**Figure 5b**) and was enriched for GO terms related to the regulation of cytoskeleton and tissue integrity, including terms such as regulation of *microtubule-based process* (GO:0032886), *actin filament bundle assembly* (GO:0051017), *actin-mediated cell contraction* (GO:0070252), *homotypic cell-cell adhesion* (GO:0034109), *positive regulation of wound healing* (GO:0090303), *actin filament polymerization* (GO:0030041) and *polarized epithelial cell differentiation* (GO:0030859). In the Cellular Compartment (CC) category, proteins were associated with the *plasma membrane* (GO:0044459, GO:0120025), *outer and inner dense plaque of desmosome* (GO:0090636), *terminal web* (GO:1990357) and the *myosin II complex* (GO:0016460).

**Figure 5.**
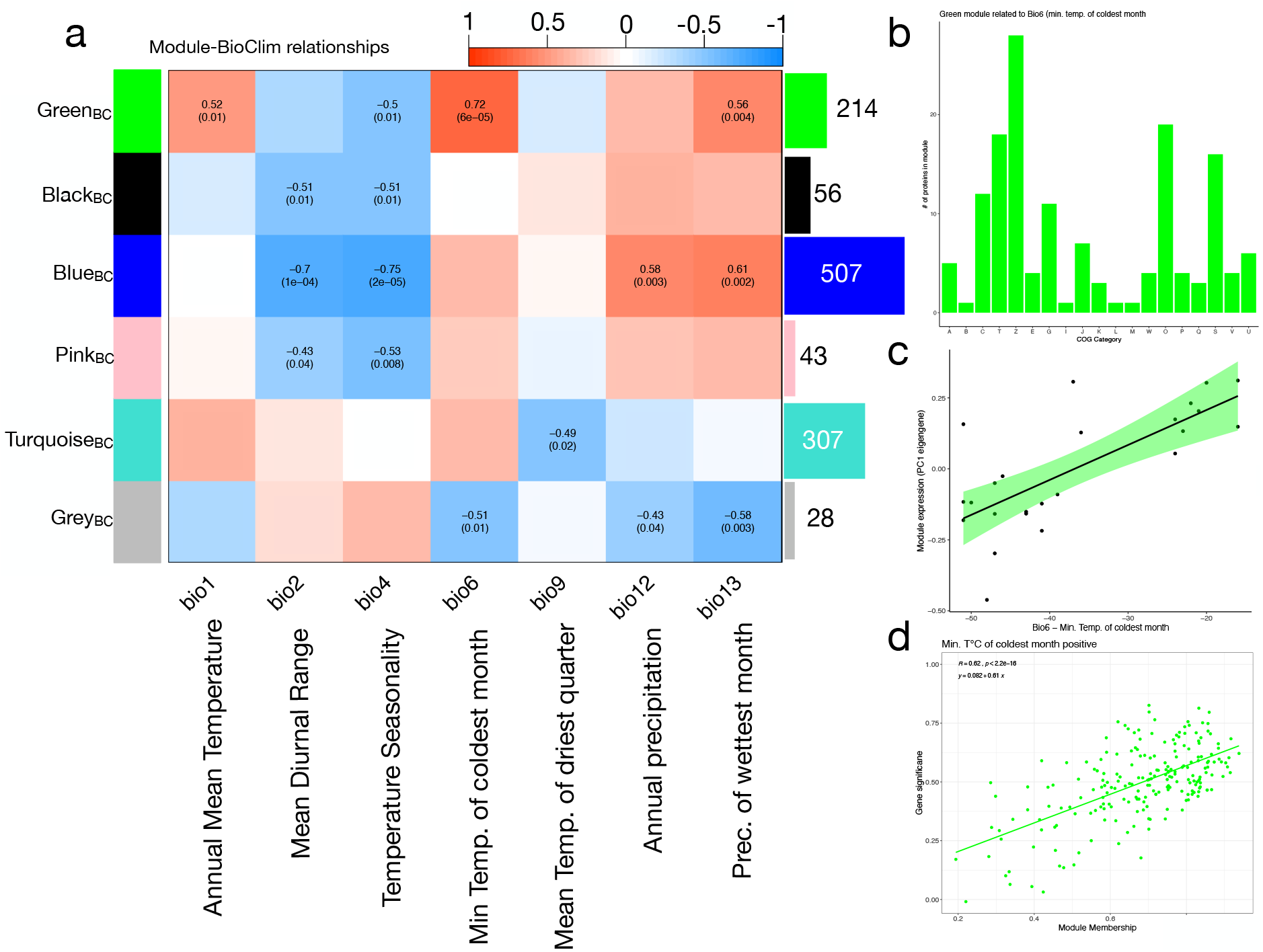
Network analysis of *Crunoecia irrorata* protein abundances in relation to bioclimatic variables (WGCNA_BC_). (**a**) Correlations between module eigengenes (rows) and bioclimatic variables (columns). Colors and numbers convey information identical to **Figure 4a**. (**b**) COG family distribution of proteins belonging to the “green_BC_” module. One-letter abbreviations are identical to **Figure 2**. (**c**) “Green_BC_” module eigengene expression positively associated with BioClim Bio6. (**d**) Scatterplot illustrating the relationship between protein’s “green_BC_” membership score (x-axis) and the protein’s significance for Bio6 (min. temperature of coldest month) (y-axis).

## 4. Discussion

Relating the most variable proteins that drive proteome differentiation among populations to their associated protein families and functions can provide broad insight into which physiological systems respond observably and dynamically to abiotic variation. As a result, any environmental change that is to be deduced from molecular profiles that are associated with any of these physiological systems are potentially confounded. In this study, the differentiation among population proteomes was associated with environmental heterogeneity between springs, geographic distance between populations and past climatic conditions. These findings indicate that the presence or absence of spectra associated with certain peptides/proteins may be an important indicator of environmental differences and that relative protein abundance differences are influenced by the geographic distance between sites. Even though no relationship between protein abundances and environmental distance emerged, protein reaction norms and association of protein modules with abiotic variation show that protein abundances of *C. irrorata* populations are regulated by the species’ integration of variation in abiotic conditions commonly experienced in its habitat. The absence of such a relationship (i.e. the majority of identified proteins did not exhibit a relation or a norm of reaction) was to be expected since cells maintain their own internal environment, e.g. via ion-transport^82^. Proteins functioning in metabolic pathways and energy acquisition separated sampling regions according to the nMDS, indicating that nutrient availability and diet differ between populations and affect protein abundances. Further, protein families showing high frequencies among all DAPs likely contribute substantially to proteome-divergence. The high frequencies of DAPs belonging to the J: *translation, ribosomal structure and biogenesis* and O: *Post-translational modification, protein turnover, and chaperones* families and the observation that the majority of enriched GO terms within DAPs were related to cellular-level processes indicate that the majority of proteins determining such region-wide proteome divergence are to a large degree involved in cellular processes/signaling. This points to differences in cellular environmental sensing^83^, and post-translational modification (PTM), regulatory modifications that activate, change or suppress protein functions^84^. Our findings therefore substantiate the importance of PTMs in integrating environmental cues and regulating physiological processes, suggesting that PTM profile differences between natural populations are substantial and are agents of plasticity. As a detailed discussion of all significant module-abiotic relationships and annotation profiles is beyond the scope of this research article, we focused the discussion on certain protein reaction norms and protein networks that corresponded with differences in abiotic profiles of springs. We first establish functional connections between modules and abiotic variation as this information is of great relevance to studies investigating molecular abundance and activity differentiation between populations^85,86^.

### 4.1. Functional relationships between modules and abiotic variation

We highlight functional relationships between modules and abiotic variables with the examples of the “yellow” module, related to pH, and the “green” module, related to oxygen concentration. Proteins in the “yellow” module show functional profiles related to cell-internal and external pH conditions: Member proteins such as succinate dehydrogenase and peptidyl-prolyl cis-trans isomerase are activated by changes in pH and modulate intracellular pH homeostasis, respectively^87,88^. Enrichment for *electron transport chain* (GO:0022900) of member proteins such as electron transfer flavoprotein (subunits alpha/beta (C- and N-terminal)) indicate a relationship between aqueous pH and the electron transport system (ETS). Fifteen proteins in this module are found in this oligomeric enzyme complex within the inner mitochondrial membrane and of these, 7 show increasing reaction norms with increasing pH (**SI1 Figure 10 & 11**). The ETS of freshwater invertebrates is dependent on external pH, whereby its activity is reduced at lower pH^89^. The relationship between this module and pH variation is further corroborated when analyzing its Pfam and COG family distributions (**SI1 Figure 12b, Figure 2d**). Higher frequencies of proteins with a zinc finger (LIM type) domain, filamin/ABP280 repeat structure and a gelsolin-like domain such as galectin-8 and a high frequency of proteins related to the Z: *cytoskeleton* family (*n* = 22) likely reflect the pH-dependent regulation of the structure and contractility of the actin-based cytoskeleton, as proteins with these domains and within this family are involved in the control of cytoskeletal function^90–93^. These actin-binding proteins and their ability to cross-link actin filaments is pH-dependent^94–96^ and their regulation of the cytoskeleton an integral response of organisms to various environmental contexts including changes in pH^97,98^. Proteins in the “green” module, associated with variation in oxygen concentration, were characterized by high frequencies of heat shock protein 70 family and insect cuticle domains (**SI1 Figure 12a**) and were predominantly associated with the COG family O: *Post-translational modification, protein turnover, and chaperones* (**Figure 2e**). Translated, unfolded proteins require oxygen to form disulfide bonds^99^ which may explain why varying oxygen levels seem to induce regulation of the unfolded protein response. The relationship between the insect cuticle and oxygen concentration is less understood but our data points to key proteins involved in regulating cuticle characteristics such as permeability in response to varying oxygenation of freshwater, potentially to release reactive oxygen species (ROS)^100^ or to facilitate penetration of dissolved oxygen^101^. Multiple enriched GO terms such as *oxidoreductase activity* (GO:0016491), *oxidoreductase activity, acting on CH_2_-OH group of donors* (GO:0016614) and *oxidation-reduction process* (GO:0055114) and GO terms related to externally-induced oxidative stress such as *response to oxidative stress* (GO:0006979) and *response to abiotic stimulus* (GO:0009628) were enriched for this module (**SI2**), providing further evidence for this relationship. The relation between GO enrichment of *intestinal stem cell homeostasis* (GO:0036335) and *regulation of stem cell differentiation* (GO:2000736) in this module and varying oxygen levels of springs suggests that *C. irrorata* larvae actively regulate oxygen levels in the low oxygen/ROS niches in which insect stem cells reside, potentially to avoid the effects of ROS on stem cells such as DNA damage and senescence^102,103^.

Overall, the outlined functional relationships indicate the coordinated regulation of protein abundances in response to abiotic variation, validating the present approach as a way to connect protein dynamics with changes in abiotic conditions. Given this functional relationship between pH variation and the “yellow” module, testable hypotheses about adaptation and phenotypically plastic responses to pH variation can be formulated: For example, the enriched GO term *retina layer formation* in the “yellow” module may reflect a previous observation that retinal gene expression changes in response to a drop in local pH^104^, hinting at the pH-induced regulation of stemmata physiology in the Trichoptera larval ocelli, a potential adaptation to live underwater.

### 4.2. Variation of biomarker proteins

A consequence of the relation between adaptive physiology and environment is that baselines and activity levels of protein biomarkers may shift according to abiotic variation of the studied system. Eco-toxicological studies showed, that enzymatic variability across time and/or space is often not fully explained by stress factors such as pollution^105,106^. Additionally, abundance measures of protein biomarkers are regularly applied in eco-toxicological research, including freshwater invertebrates^107^. Members include e.g. the cytochrome P450 mixed function oxidases and enzymes such as catalase and filamin-A^108–111^. Catalase, filamin-A, ATP-citrate(pro-5)-lyase and the two cytochromes Cyp62a and Cyp4c1 exhibited significant reaction norms in relation to elevation (**SI1 Figure 13**). Both, *Cyp6a2* and *Cyp4c1* are responsible for the metabolism of numerous xenobiotics and endogenous compounds, including organophosphorus insecticides such as dichlorodiphenyltrichloroethane (DDT)^112–114^. Reaction norms of these two cytochromes over an elevation range hints at a potential presence of organophosphorus insecticides in groundwater-aquifers in Central Europe. Moreover, it might indicate that the contemporary application of organophosphorus insecticides in agricultural practice results in groundwater contamination as anthropogenic input of xenobiotics should increase with decreased elevation and corresponding increased settlement density. The described biomarker norms of reaction exemplify that sources and nature of variability in organism-environment relationships need to be understood and made explicit; i.e. used as information rather than dismissed as noise in eco-toxicological analyses of natural populations.

### 4.3. Adjusting for baseline deviations

In comparative field-studies that investigate the systems-wide regulation of proteins in response to environmental change (e.g. comparing populations experiencing drought vs. no drought or a shift in mean temperature vs. no shift), the influence of that change on organismal physiology is commonly assessed through the log_2_ fold-change (FC) values of pairwise comparisons between abundances of identified proteins^115,116^. Since these compared abundances are directly influenced by abiotic variation and therefore by environmental differences between sites, results of these comparisons via e.g. differential expression analysis may over-, or even under-estimate the actual differences between compared groups. Including information on baseline abundance shifts due to confounding variables could be achieved by accounting for the height difference between regression lines (e.g. Filamin-A difference between regions/sites) in the differential analysis performed on the same input expression/abundance matrix. For example, subtracting the height difference (~0.7 log_2_ abundance) between region-specific reaction norms from the output of the differential abundance analysis between H and BF (log_2_ FC = −4.26, *p*_adj_ = 0.0026; **SI2**) results in a FC closer to - 3.56. This value should then represent the protein abundance change between sites that resulted from abiotic or biotic differences other than water temperature.

### 4.4. Temperature tolerance

Despite being a key environmental variable in determining the physiology of ectothermic organisms, *in situ* water temperature was not a regulating abiotic factor of a protein-network. In contrast, proteins regulating the cytoskeleton appear to be related to past extreme climatic conditions. “Green_BC”_ eigengene expression increased with higher minimum extreme temperatures and annual mean temperature (**Figure 5**). Cytoskeletal integrity is perturbed by a variety of cell stress responses and plays a crucial role in maintenance of cell homeostasis^117,118^. This relationship echoes previous findings that show cytoskeletal regulation in response to temperature change via e.g. associated cryoprotective dehydration^102–106^. Population-wide regulation of cytoskeletal proteins in response to previously experienced thermal extremes indicates heritable protein abundance variation between populations attributable to heritable gene expression variation^124,125^, supporting the notion that extreme environmental temperatures exert a strong selective pressure on ectothermic species^126^. Interestingly, the “green_BC_” module also shows decreasing eigengene expression with increasing temperature seasonality, i.e. the standard deviation of the mean monthly temperature. This indicates that cytoskeleton regulation is more significant when organisms experience less-variable temperature regimes (i.e. longer stretches of warmer or colder temperatures).

Integral parts of the cellular stress response include the evolutionary conserved heat-shock- and cold-shock-response (HSR and CSR), characterized by the expression of heat-shock proteins (Hsps). Putatively cold-adapted species have been shown to constitutively express Hsps to facilitate protein folding at low temperatures^127,128^, a phenomenon also observed in *C. irrorata*^48^. Here we observed constitutive and increasing but also *decreasing* Hsp reaction norms with increasing *in situ* temperatures (**SI1 Figure 14 & 16**). Chaperones with decreasing norms of reaction such as Hsp83 and Hsp90 might be integral for alleviating cold-induced protein denaturing and increased protein folding time. These patterns indicate evolutionarily sub-divided roles within the Hsp gene family in putatively cold-adapted species, with certain Hsps functioning during HSRs and others during CSRs. In light of the present data, chaperonines belonging to the Chaperonin Cpn60/TCP-1 protein family such as Hsp68 could be indicative of cold-adaptation of *C. irrorata*, especially since chaperonins govern cell growth at cold temperatures and springs show constant cold temperatures^79,129,130^. Species from such stable and cold environments might be ill-adapted for climate-induced warming of aquatic ecosystems^131,132^. Somewhat counterintuitively, recent cellular evidence indicates that for certain aquatic species, the stable cold environment might be a conditioning factor such that they evolved to be able to protect themselves from temperature-induced cellular damage^128,133,134^.

### 4.5. Limitations and implications for the study of natural populations

This work assumes that if two populations have a different quantity of protein it means that the proportion of individuals which express that protein differs between populations. Inter-individual differences are not only pronounced but directly influence the population-level outcome of environmental change^135,136^. Consequently, even in the absence of a significant population-level response to the environment, some individuals may still respond plastically to changing conditions. Due to this strong inter-individual variation, it may be challenging to apply omics-wide assessments on individuals if the goal is to obtain a “screening” of the physiological status of populations by relating their expression or abundance profiles to external conditions. Overall, utilizing pooled population-proteome profiles can help keeping sample sizes smaller than inter-individual studies and as shown in this study, does not apparently mask abundance patterns in response to environmental variation, including established biomarker proteins. This shows the feasibility of the approach for detecting population-wide physiological responses to environmental change, potentially in response to toxic substances not previously known to be present in the studied ecosystem. The systematic characterization of such relationships in a non-targeted approach, using a non-model organism, has the potential to give unprecedented insights into the molecular functioning of a variety of ecologically relevant species, exemplifying that plastic responses need to be considered when investigating adaptive processes to changing environments. We further recognize that there are many other sources of confounding factors that complicate the interpretation of the observed molecular responses. Our study design did not account for biotic influences on molecular profiles such as predator presence^137^, parasitism status^138^ and microbial differences between sites and individuals or populations^139,140^. Lastly, information on other abiotic factors such as average, prevailing conditions of the studied system are additionally needed to further refine baseline levels of protein abundances and to understand how natural environmental conditions might mask and alter the molecular responses of natural populations.

## 5. Conclusion

To our knowledge, this is the first study to systematically investigate the relationships between protein abundances/identifications and abiotic conditions in natural populations of a freshwater species. The primary findings are that the variation in proteome profiles of pooled population samples are attributable to abiotic but also geographic and past climatic differences between sites. Variables such as pH, oxygen concentration and past thermal minima “explained” part of the variation and need to be considered when comparing sites experiencing differential environmental change. Additionally, protein families related to cellular information processing and PTM regulation contribute to proteome divergence overall and likely play a vital role in physiological adjustment to varying abiotic conditions. More fundamentally, protein norms of reaction are ubiquitous and often reflect the functional roles these proteins have within multicellular organisms. Additionally, signatures of local adaptation (e.g. warming temperatures or differences in oxygenation) can be gleaned through such comparative analyses, whereby up-regulation of cytoskeleton-related proteins in populations experiencing warmer temperatures on a historical timescale are just one example. This considerable variability in protein abundance patterns likely distorts abundance differences in comparative field studies, leading to over- or under-estimation of environmental change experienced by organisms. Integrating this natural variation appears to be an important step in systems-wide biomarker development and for disentangling the differences between populations experiencing differential environmental change.

## Data availability

We provided all raw, transformed and filtered data required to interpret, replicate, and build upon our findings in supporting information files **SI1-SI4** (combining quantitative and annotation information). Mass spectrometry raw data have been deposited to the ProteomeXchange Consortium via the PRIDE partner repository^141^ with the data set identifier PXD017959. R scripts used in this study are available as **SI5**.

## Author’s Contributions

This study is part of the doctoral research of J.N.E. under the supervision of S.F. J.N.E. conceived the idea., S.F. and J.N.E. conceived of study design and conducted field-work. D.R. and J.N.E. generated the mass spectrometry data. J.N.E. analyzed data and wrote the manuscript.

## Acknowledgements

We want to thank Stefan and Christian Zaenker and Haris Mujagic for indispensable help during field-work. We thank the Harz National Park authorities for granting sampling permission and area access. Part of the calculations was performed at sciCORE (http://scicore.unibas.ch/) scientific computing center at the University of Basel. This study was funded by the Swiss National Science Foundation (SNF) under grant number 31003A_176234.

