## Supplemental Information 1 for "Abiotic drivers of protein abundance variation among natural populations"

**Affiliations:**
[1]Geoecology Research Group, Department of Environmental Sciences, University of Basel, Basel, Switzerland
[2]Proteomics Core Facility, Biozentrum, University of Basel, Basel, Switzerland

### Contents:

#### Supplemental tables

**Table 1** | Information on sampled freshwater springs (code (by the Authors, does not reflect official coding), elevation, sampling date, region, latitude and longitude (decimal values))

**Table 2** | Peptide and protein concentration measurements of population-wide samples subjected to LC-MS/MS analysis

#### Supplemental figures

**Figure 1** | Dotplots of mean values of all abiotic measurements

**Figure 2** | Cor.test output of pairwise comparisons of abiotic variables

**Figure 3** | NAs of MaxQuant LFQ

**Figure 4** | nMDS stress plot

**Figure 5** | Statistics and results of the WGCNA analysis

**Figure 6** | Statistics and results of the $WGCNA_{BC}$ analysis

**Figure 7** | Capsule width and protein ID overlap

**Figure 8** | Scatterplot and linear regression of protein abundance distance and geographic distance between springs

**Figure 9 |** Scatterplot and linear regression of environmental distance and geographic distance between springs

**Figure 10 |** Boxplots and significance results of Kruskal-Wallis pairwise comparisons of 4 abiotic variables between sampling regions

**Figure 11** | Reaction norms of 33 proteins in relation to elevation of springs

**Figure 12 |** Pathways related to changes in aqueous pH

**Figure 13 |** Pfam family frequency in modules "green" and "yellow" of the WGCNA

**Figure 14 |** Exemplary reaction norms (polynomial regression) of population-wide protein biomarker abundances (y-axis) in relation to spring altitude and temperature (x-axis)

**Figure 15 |** Heat-shock protein reaction norms

**SI1 Table 1 |** Code = Spring- (population)-codes, Elev. = elevation (m.a.s.l), Sample Date = Sampling of springs (identifying individuals and storing them in RNAlater, measuring of abiotic variables and noting coordinates) occurred between 07:00 and 18:00 on all sampling days), Regions = One of the three regions in which springs were probed (Rhoen = Rhoen Biosphere Reserve, Harz = Harz National Park, BF = "Black Forest"), Lat. & Long. = latitudes and longitudes in decimal values for each sampled spring as recorded with the "GPS Coordinates" App for Android, referenced with a *e*Trex Summit® HC handheld GPS (Garmin).

| Code | Elev. | Sample Date | Regions | Lat. (Decimal) | Long. (Decimal) |
|---|---|---|---|---|---|
| R1 | 370 | 29.07.2019 | Rhoen | 50.613586 | 9.764174 |
| R2 | 570 | 29.07.2019 | Rhoen | 50.5993729 | 9.99908903 |
| R3 | 600 | 29.07.2019 | Rhoen | 50.5889192 | 10.0041492 |
| R4 | 818 | 29.07.2019 | Rhoen | 50.4678952 | 10.0136548 |
| R5 | 815 | 29.07.2019 | Rhoen | 50.4658716 | 10.0054564 |
| R6 | 730 | 29.07.2019 | Rhoen | 50.5035052 | 9.92957296 |
| R7 | 606 | 29.07.2019 | Rhoen | 50.517887 | 9.932069 |
| R8 | 275 | 30.07.2019 | Rhoen | 50.6096721 | 9.64546653 |
| R9 | 340 | 30.07.2019 | Rhoen | 50.5993789 | 9.66936553 |
| H1 | 332 | 31.07.2019 | Harz | 51.637179 | 10.406294 |
| H2 | 390 | 31.07.2019 | Harz | 51.70213 | 10.343042 |
| H3 | 764 | 01.08.2019 | Harz | 51.767917 | 10.610061 |
| H4 | 696 | 01.08.2019 | Harz | 51.767861 | 10.612028 |
| H5 | 482 | 01.08.2019 | Harz | 51.742708 | 10.675715 |
| H6 | 517 | 01.08.2019 | Harz | 51.742801 | 10.675805 |
| H7 | 601 | 02.08.2019 | Harz | 51.726299 | 10.564328 |
| H8 | 597 | 02.08.2019 | Harz | 51.643169 | 10.696474 |
| H9 | 577 | 02.08.2019 | Harz | 51.647887 | 10.741779 |
| SW1 | 912 | 19.08.2019 | BF | 47.652422 | 7.975453 |
| SW2 | 912 | 19.08.2019 | BF | 47.652422 | 7.975456 |
| SW3 | 967 | 19.08.2019 | BF | 47.652778 | 7.973611 |
| SW4 | 346 | 19.08.2019 | BF | 47.597678 | 7.840197 |
| SW5 | 498 | 20.08.2019 | BF | 47.662372 | 7.901392 |
| SW6 | 460 | 20.08.2019 | BF | 47.65695 | 7.907233 |
| SW7 | 511 | 21.08.2019 | BF | 47.655839 | 7.767608 |
| SW8 | 589 | 21.08.2019 | BF | 47.658853 | 7.770014 |
| SW9 | 598 | 21.08.2019 | BF | 47.659061 | 7.7702 |

**SI1 Table 2 |** Sample number, spring code, protein concentrations (µg/µl) peptide concentrations (µg/µl) and amount of protein used for trypsin digestion (µg/µl) of pooled larvae samples (A – X).

| Sample number specification | Spring Code | Protein concentration (µg/µl) | Peptide concentration (µg/µl) | Amount of protein used for digestion with trypsin (µg/µl) |
|---|---|---|---|---|
| A | R1 | 0.42458 | 1.451 | ~50 µg |
| B | R2 | 0.925299 | 0.551 | ~50 µg |
| C | R3 | 0.427037 | 1.973 | ~50 µg |
| D | R5 | 1.72706 | 0.496 | ~50 µg |
| E | R6 | 0.563432 | 1.697 | ~50 µg |
| F | R7 | 0.997794 | 0.553 | ~50 µg |
| G | R8 | 1.70433 | 0.755 | ~50 µg |
| H | R9 | 0.215078 | 1.908 | ~50 µg |
| I | H1 | 0.271602 | 1.381 | ~50 µg |
| J | H2 | 1.085652 | 1.636 | ~50 µg |
| K | H3 | 0.217535 | 0.951 | ~50 µg |
| L | H4 | 0.773546 | 1.155 | ~50 µg |
| M | H5 | 0.540086 | 0.946 | ~50 µg |
| N | H6 | 0.522882 | 1.551 | ~50 µg |
| O | H7 | 0.542543 | 1.671 | ~50 µg |
| P | H9 | 0.258084 | 1.583 | ~50 µg |
| Q | SW1 | 0.213849 | 1.123 | ~50 µg |
| R | SW2 | 0.301707 | 1.553 | ~50 µg |
| S | SW3 | 0.186818 | 1.38 | ~50 µg |
| T | SW4 | 0.11862 | 1.409 | ~50 µg |
| U | SW5 | 0.280818 | 1.283 | ~50 µg |
| V | SW6 | 0.287573 | 1.197 | ~50 µg |
| W | SW7 | 0.414752 | 2.027 | ~50 µg |
| X | SW8 | 0.288806 | 0.51 | ~50 µg |

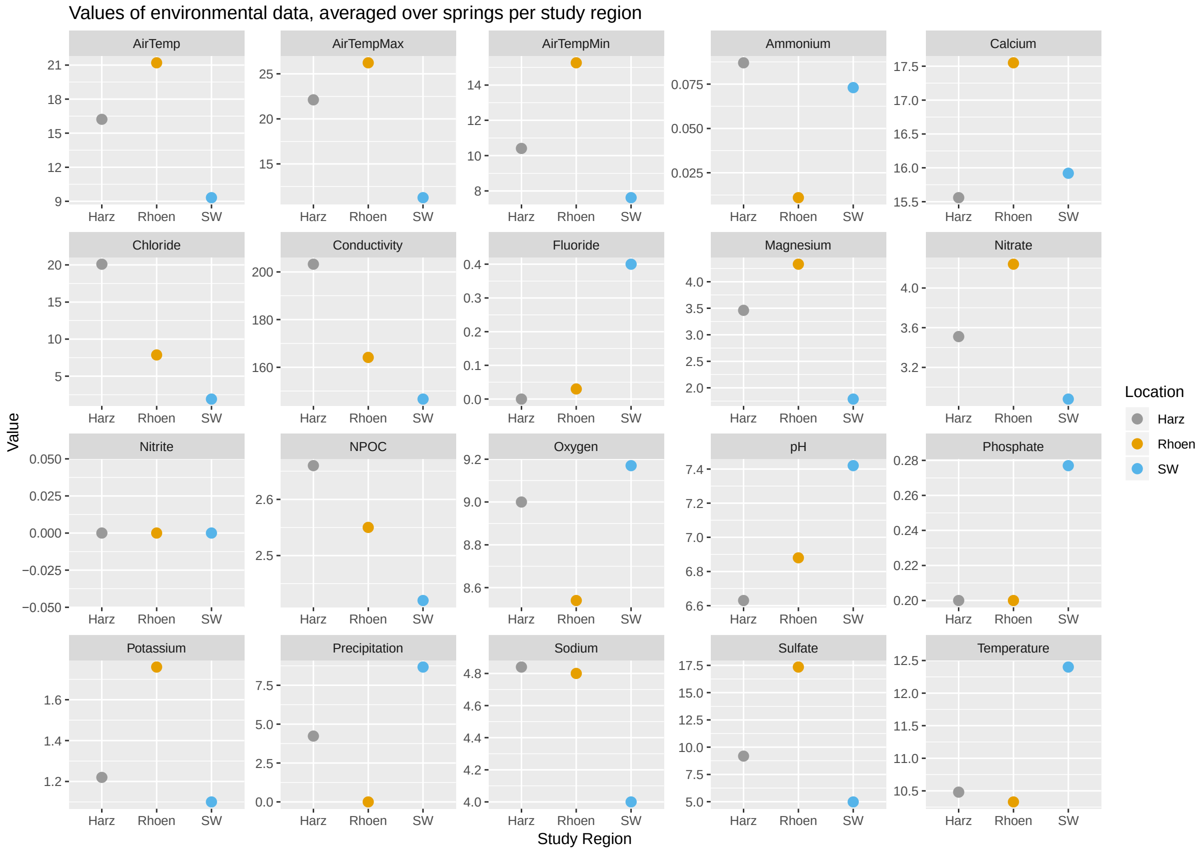


**SI1 Figure 1** | Dotplots of mean values of all abiotic measurements ($n = 9$ per region).

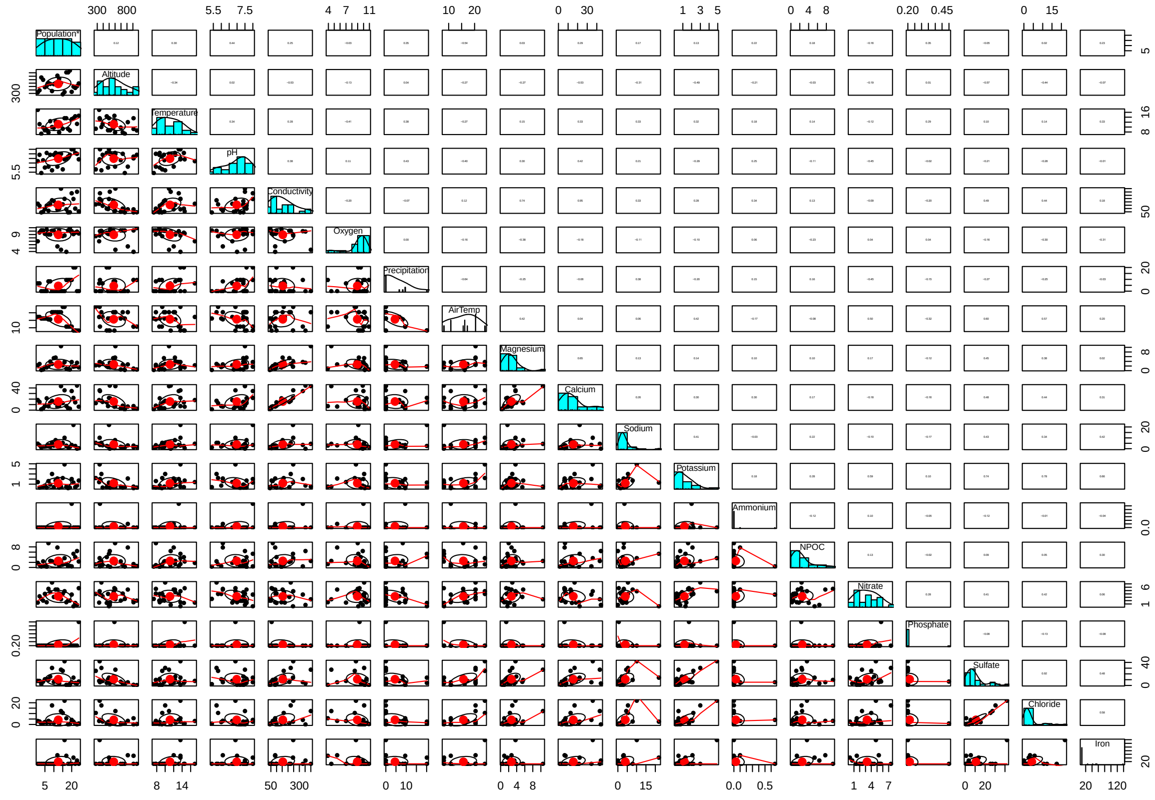


**SI1 Figure 2** | Cor.test output of pairwise comparisons of abiotic variables showing scatterplots and Pearson's correlation values. Of strongly correlating (>0.7) variables, one

was chosen to be included in analyses (for a reduced set of abiotic variables included in statistical analyses, see **SI2**).

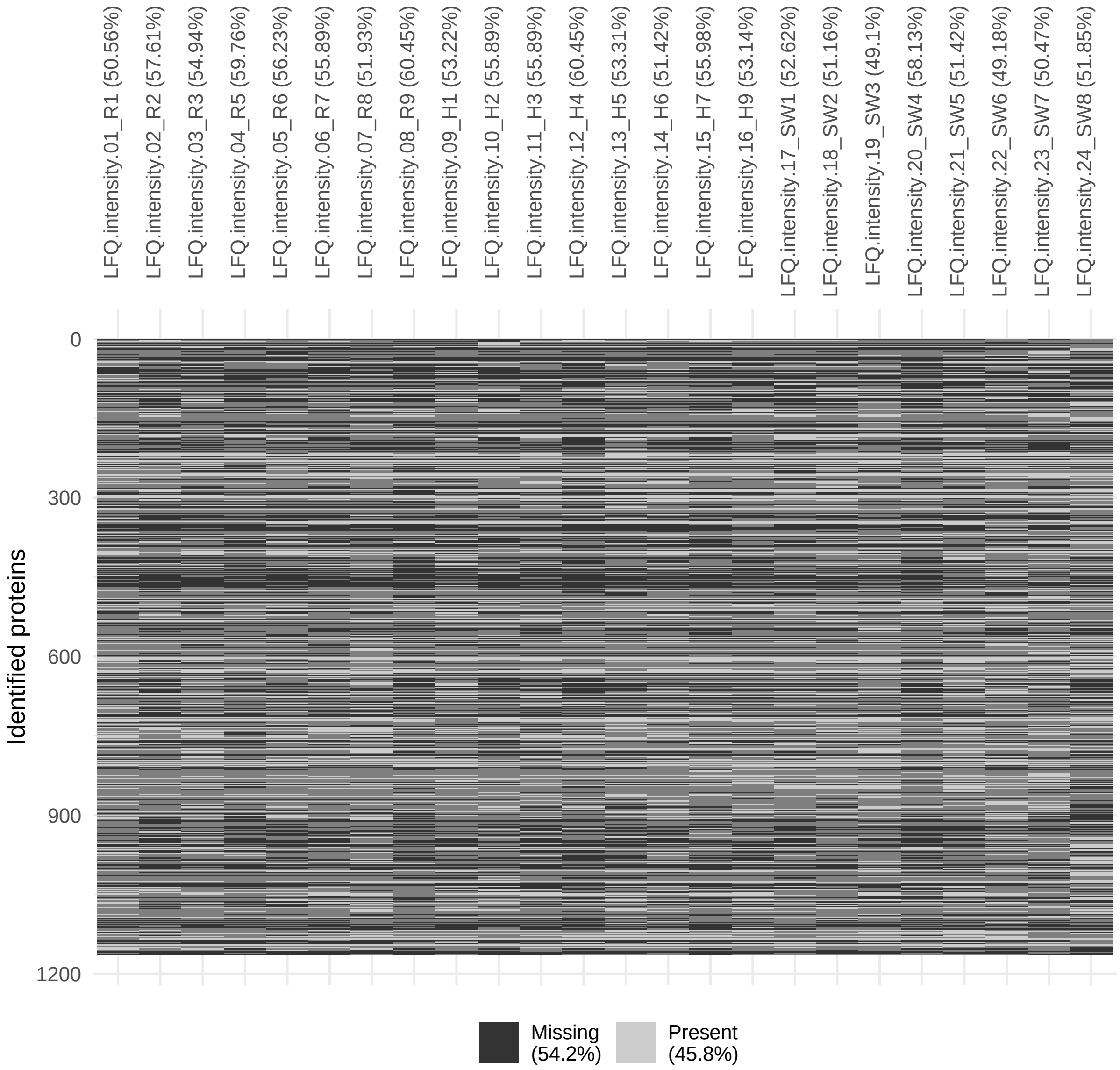


**SI1 Figure 3 |** Number of missing values (NAs) of label-free quantification (LFQ) values in the MaxQuant output. Plot was generated using the vis_miss function of package visdat v.0.5.3 (Tierney 2019).

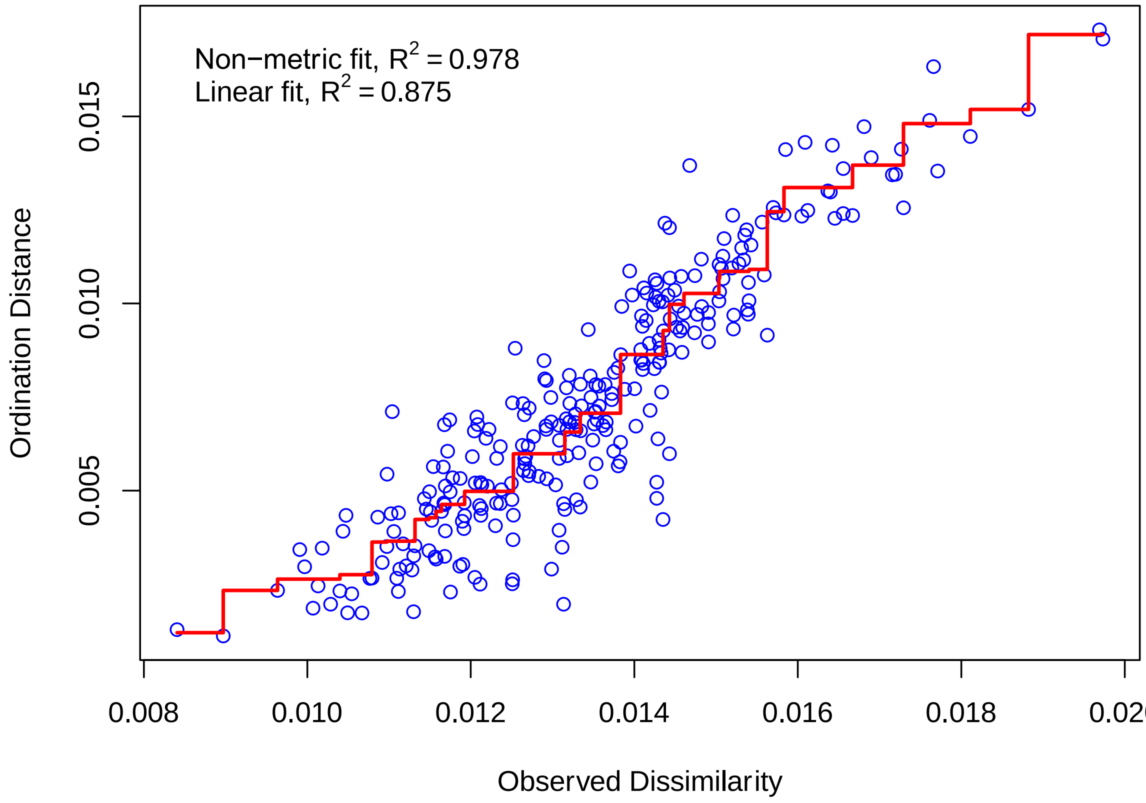


**SI1 Figure 4 |** nMDS stress plot.

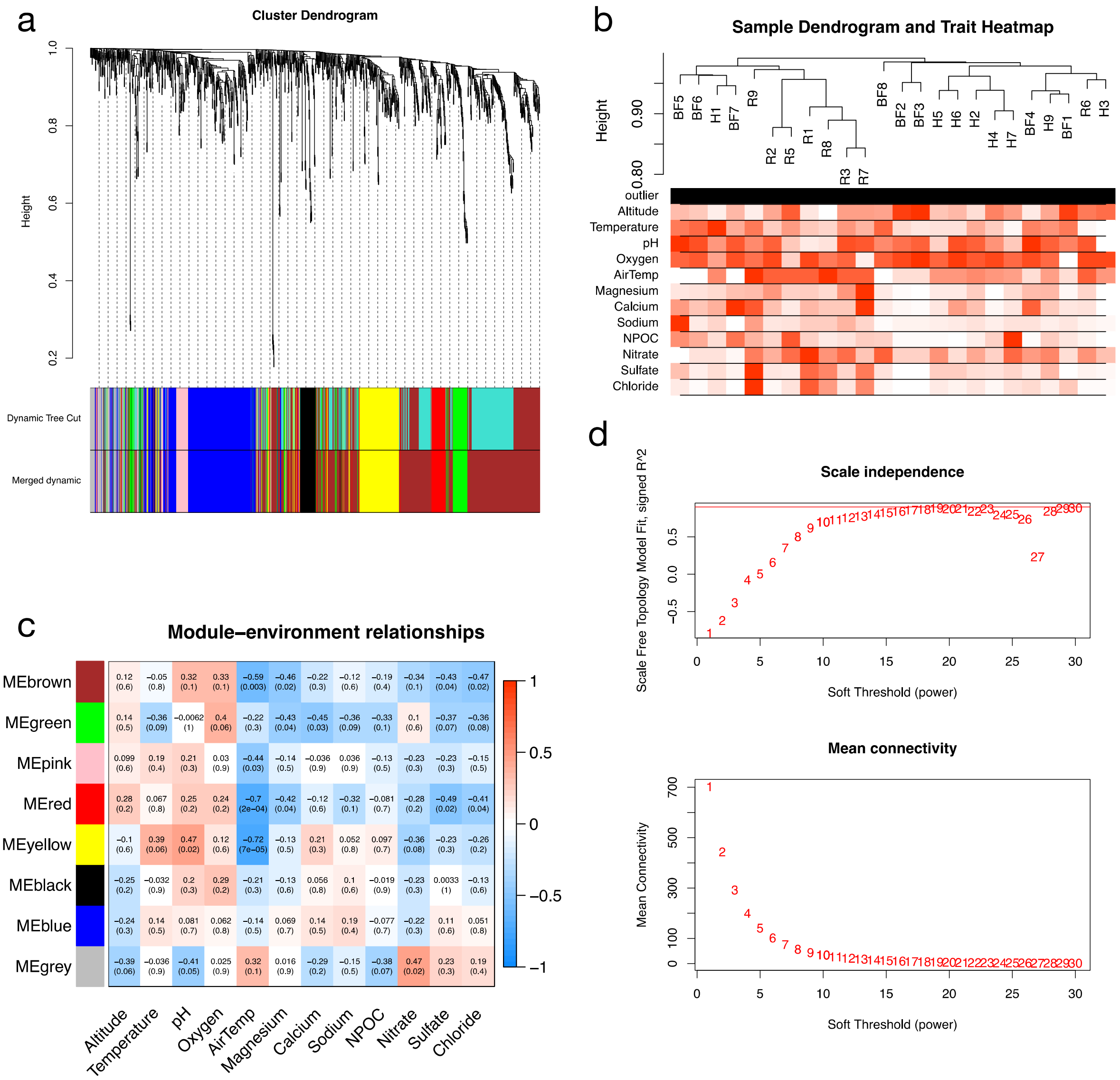


**SI1 Figure 5 | Statistics and results of the WGCNA analysis. (a)** Clustering of proteins with dissimilarity based on topological overlap, together with assigned module colors (Dynamic Tree Cut; modules with more than $n$ = 20 proteins). Merged dynamic below shows concatenated modules of Dynamic Tree Cut modules that showed a correlation of > 0.80, representing the modules used in the further analysis. **(b)** Sample dendogram and abiotic heatmap showing no outliers in the data. **(c)** Module-Environment relationships considering all 24 populations and 12 abiotic variables. **(d)** Analysis of global network topology for various soft-thresholding powers using protein abundance data of all 24 *C. irrorata* populations. Upper panel shows the scale-free fit index (y-axis) as a function of the soft-thresholding power (x-axis). The lower panel displays the mean connectivity (degree, y-axis) as a function of the soft-thresholding power (x-axis). We chose the power 18 for the global analysis, which is the lowest power for which the scale-free topology fit index reaches 0.90 (red cut-off line in left panel.

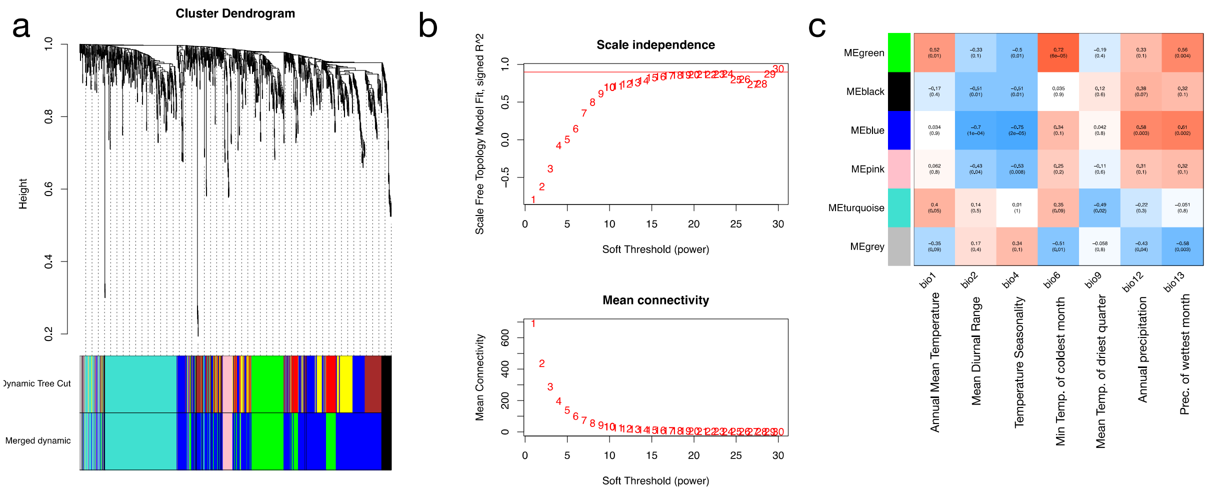


**SI1 Figure 6 | Statistics and results of the WGCNA$_{BC}$ analysis. (a)** Clustering of proteins with dissimilarity based on topological overlap, together with assigned module colors (Dynamic Tree Cut; modules with more than *n* = 20 proteins). Merged dynamic below shows concatenated modules of Dynamic Tree Cut modules that showed a correlation of > 0.80, representing the modules used in the further analysis. **(b)** Analysis of global network topology for various soft-thresholding powers using protein abundance data of all 24 *C. irrorata* populations. Upper panel shows the scale-free fit index (y-axis) as a function of the soft-thresholding power (x-axis). The lower panel displays the mean connectivity (degree, y-axis) as a function of the soft-thresholding power (x-axis). We chose the power 18 for the BioClim analysis, which is the lowest power for which the scale-free topology fit index reaches 0.90 (red cut-off line in left panel; same as for the *in situ* abiotic variable WGCNA). **(c)** Module-BioClim relationships considering all 24 populations and 7 selected BioClim variables.

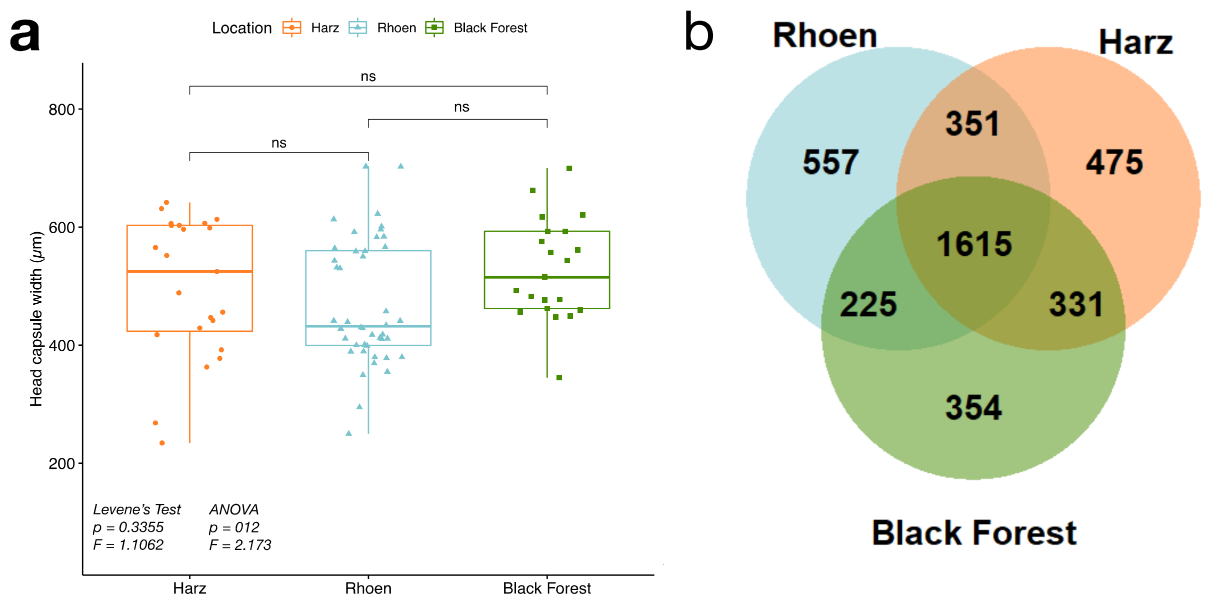

**SI1 Figure 7 |** (**a**) Boxplots of head capsule widths of randomly sampled individuals ($n = 88$) and significance results of pairwise comparison between sampling regions. (**b**) Venn diagram of number of identified majority protein IDs among sampling regions.

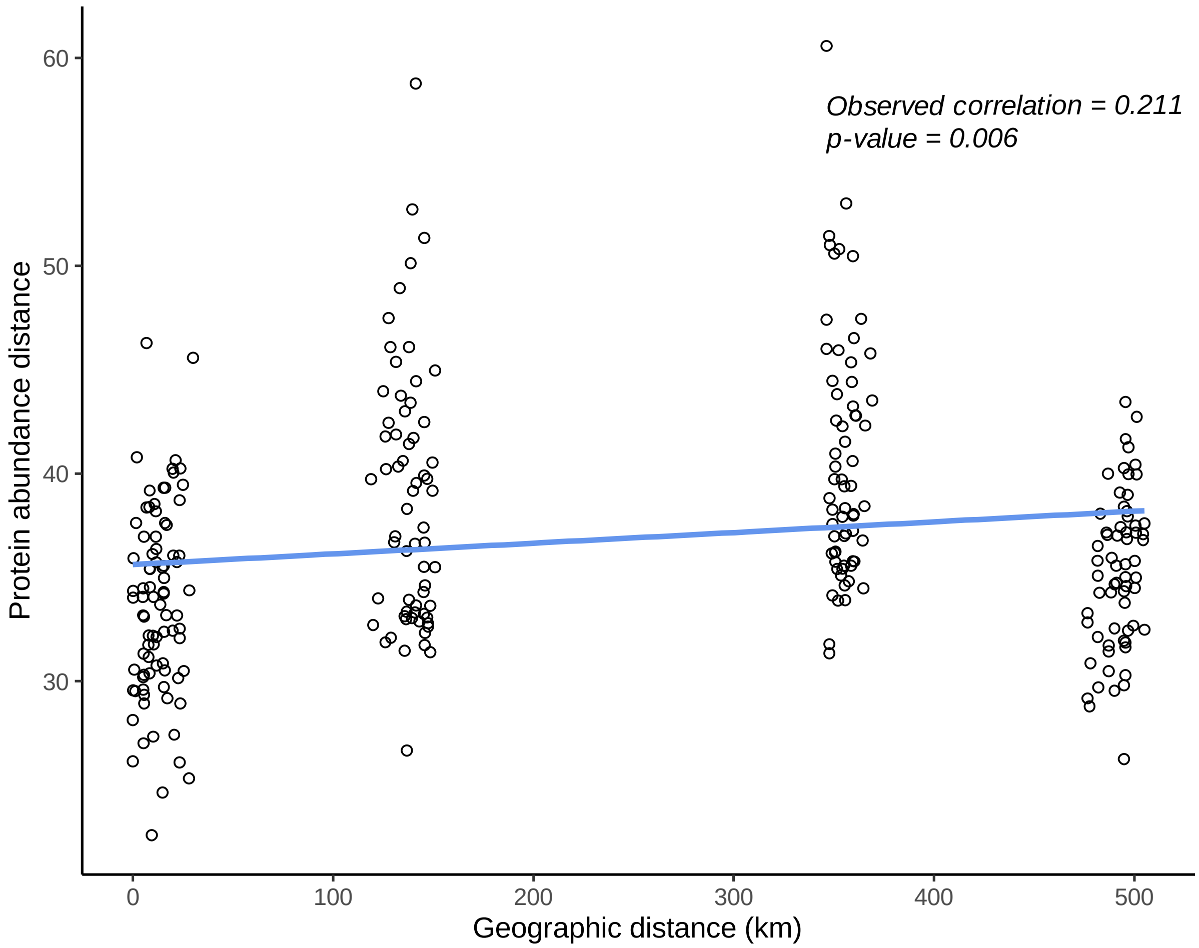


**SI1 Figure 8 |** Scatterplot and linear regression of protein abundance distance (Euclidean) between springs (y-axis) and geographic distance (km) between springs. Mantel test statistics are given in the top right corner (observed correlation and $p$-value).

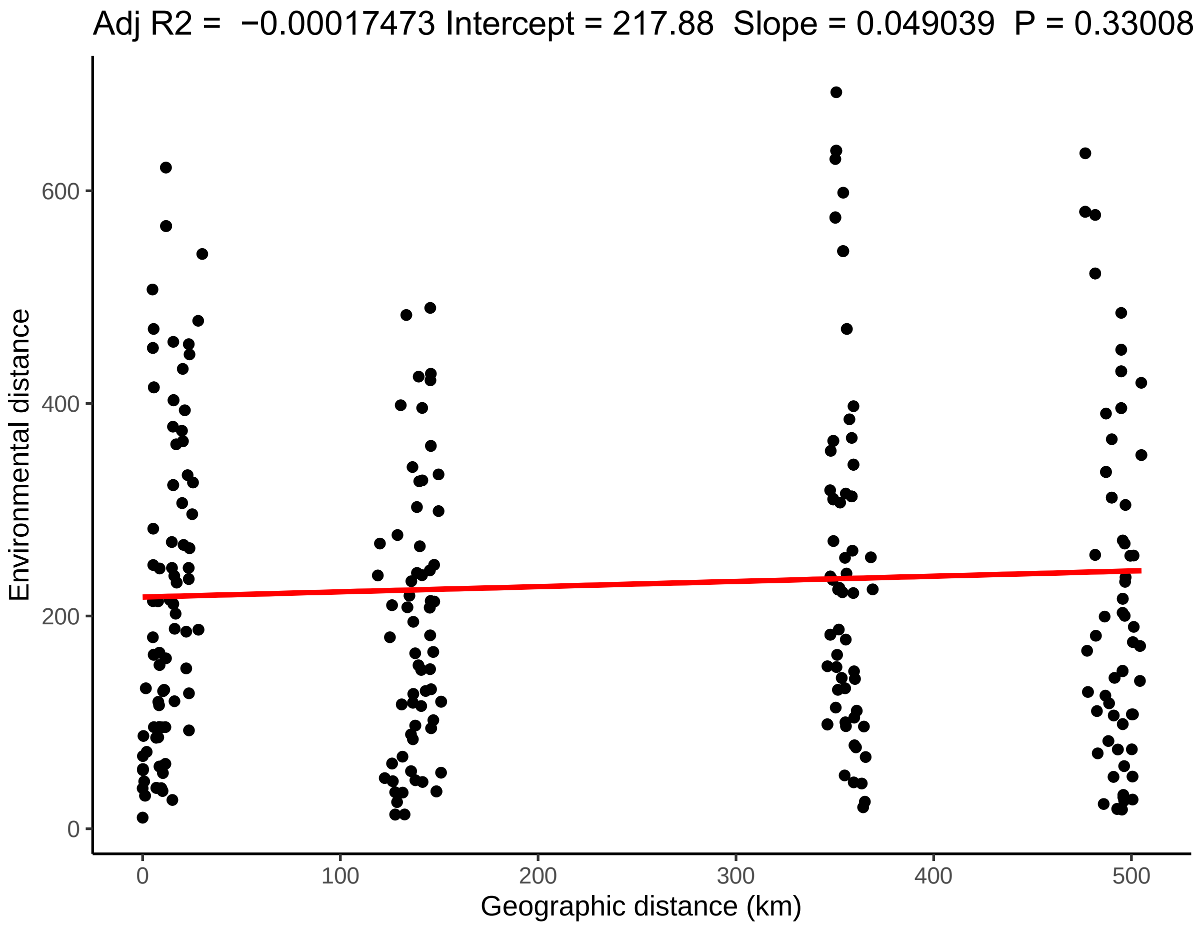


**SI1 Figure 9** | Scatterplot and linear regression of environmental distance (Euclidean) between springs (y-axis; based on 11 abiotic variables) and geographic distance (km) between springs. Adjusted $R_2$, intercept, slope and significance of the linear model are given on top of the scatterplot.

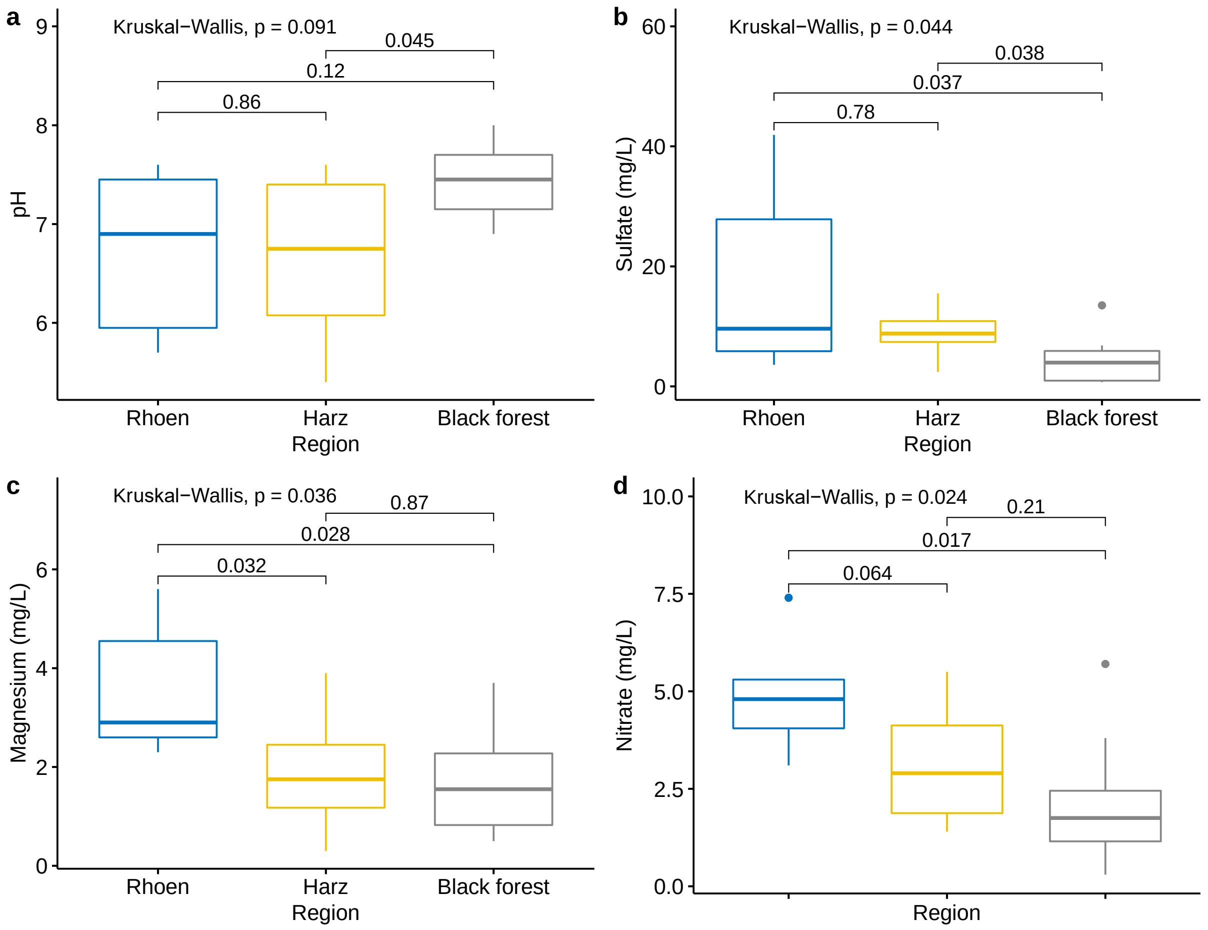


**SI1 Figure 10 |** Boxplots and significance results of Kruskal-Wallis pairwise comparisons of 4 abiotic variables between sampling regions. (**a**) pH. (**b**) Sulfate. (**c**) Magnesium. (**d**) Nitrate.

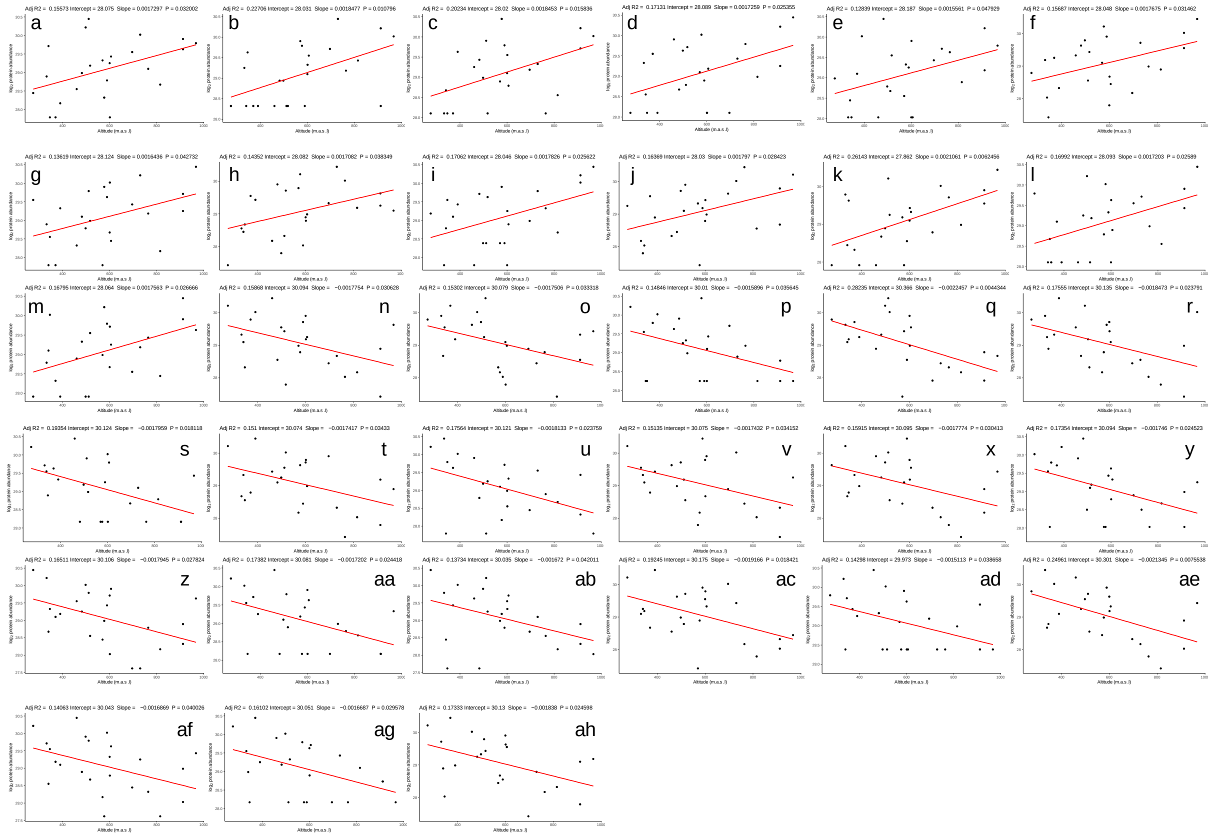

**SI1 Figure 11 | Reaction norms of 33 proteins in relation to elevation (m.a.s.l) of springs.** Of these, 13 show significantly increased abundances with increasing altitude (**a-m**) and 20 proteins significantly decreasing abundances with increasing altitude (**n-ah**).

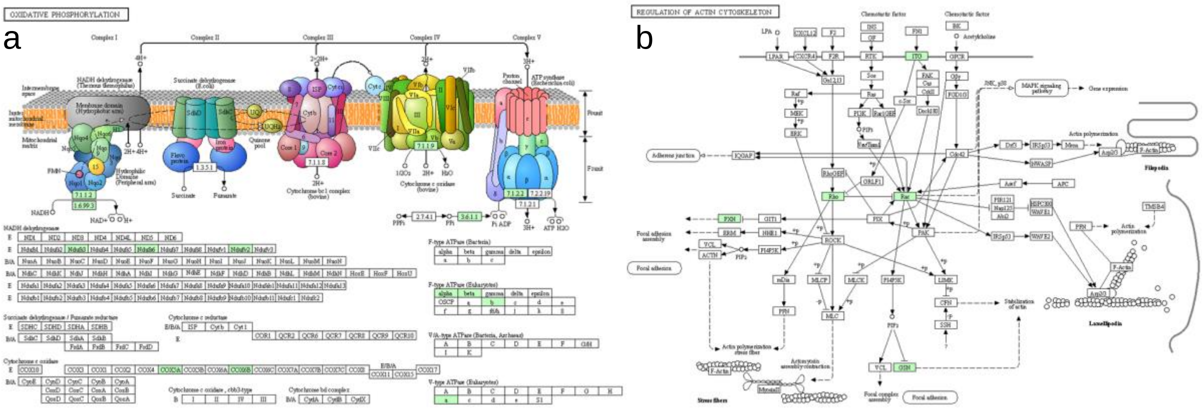


**SI1 Figure 12 | Pathways related to changes in aqueous pH.** Mapping of member proteins of the "yellow" module ($n_{yellow}$ = 110; 82.71 % annotated) to (**a**) oxidative phosphorylation ($n$ = 15 proteins (13,65 %)) and (**b**) regulation of actin cytoskeleton ($n$ = 9 proteins (8,19 %)) pathways. Green boxes represent KEGG nodes present in the module.

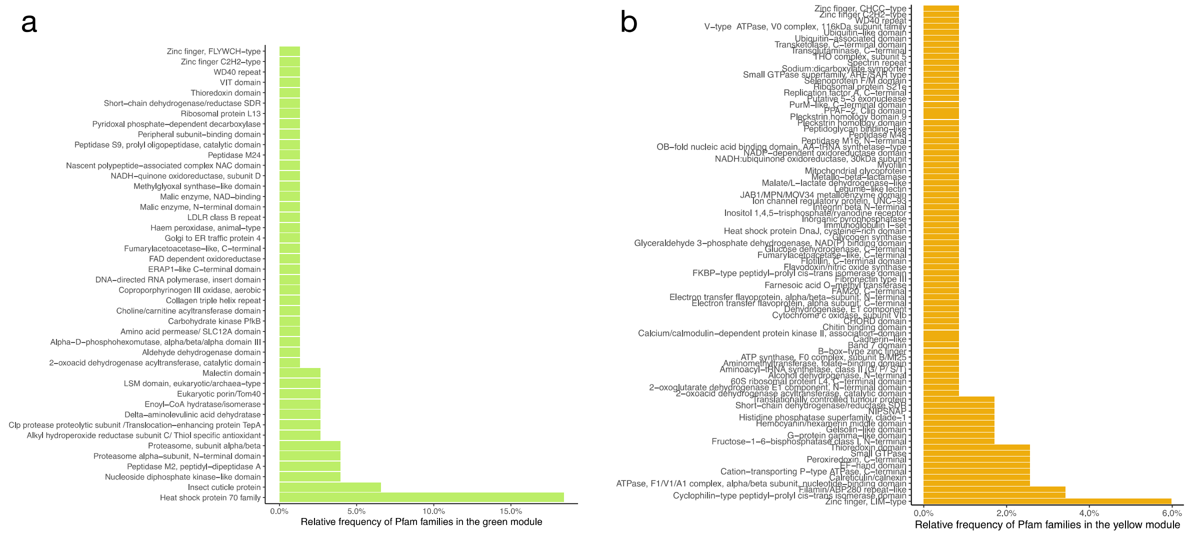


**SI1 Figure 13 |** Pfam family frequency in modules (**a**) "green" and (**b**) "yellow".

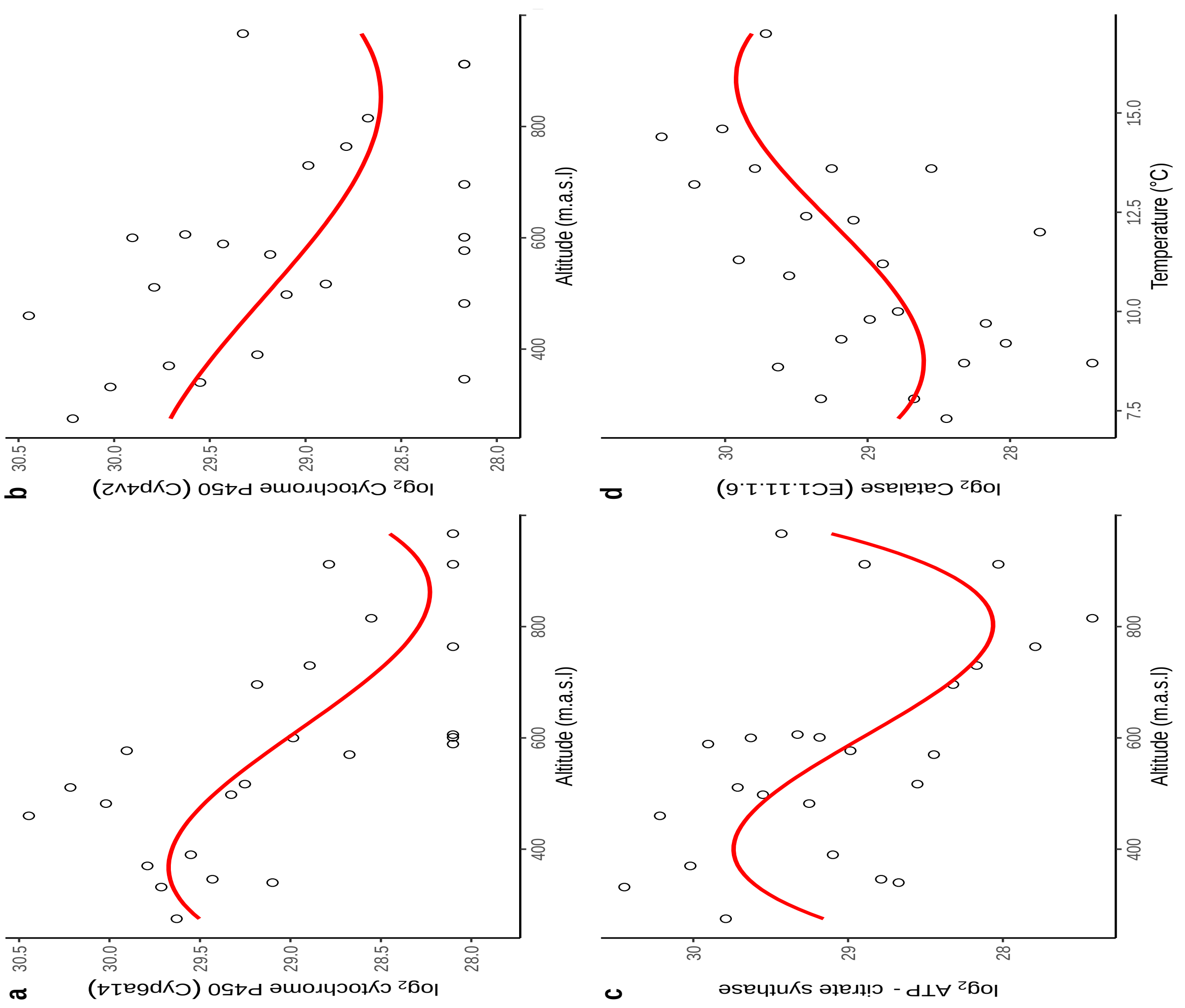


**SI1 Figure 14 | Exemplary reaction norms (polynomial regression) of population-wide protein biomarker abundances (y-axis) in relation to spring altitude and temperature (x-axis).** (**a**) Probable cytochrome P450 (*Cyp6a2*) (**b**) Cytochrome P450 4C1 (Cyp4c1). (**c**) ATP-citrate synthase. (**d**) Catalase. Shown are the fit line (red line) and 95% confidence interval (light grey area). RMSE: Root Mean Square Error; $R_2$: correlation coefficient; *p*: level of statistical significance of the model.

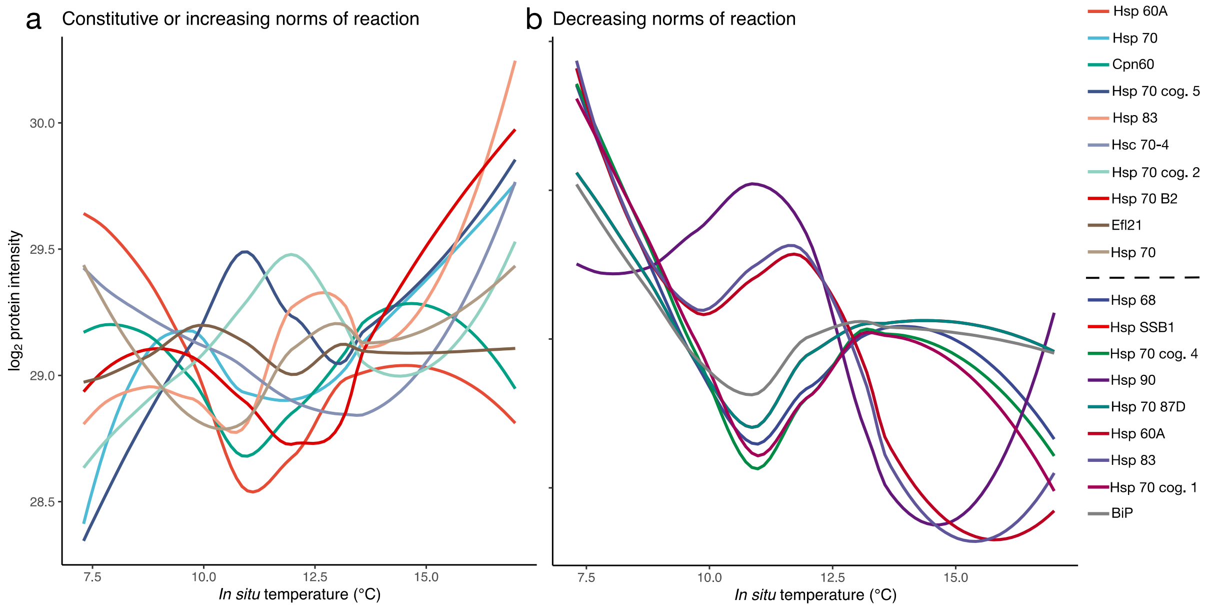


**SI1 Figure 15 | Heat-shock protein reaction norms. (a)** Constitutive or increasing norms of reaction with increasing spring *in situ* temperatures. **(b)** Decreasing norms of reaction with increasing spring *in situ* temperatures.

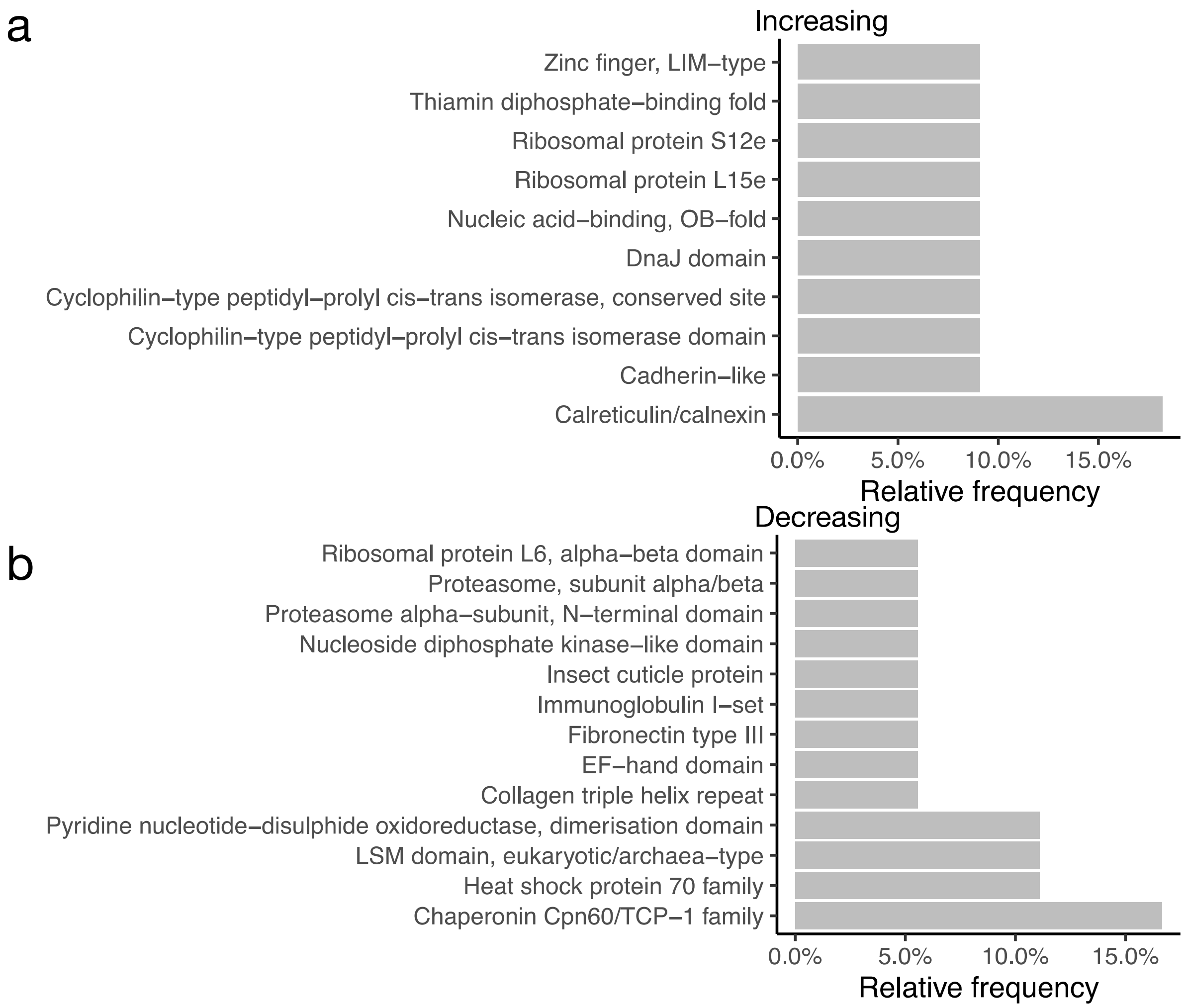

**SI1 Figure 16 | Pfam family frequencies in reaction norm proteins**. (**a**) increasing in abundance with increasing in situ temperature (°C) of springs (n = 29) and (**b**) decreasing in abundance with increasing in situ temperature (°C) of springs (n = 33).
