## Supplemental Information 4 for "Abiotic drivers of protein abundance variation among natural populations"

**Affiliations:**
[1]Geoecology Research Group, Department of Environmental Sciences, University of Basel, Basel, Switzerland
[2]Proteomics Core Facility, Biozentrum, University of Basel, Basel, Switzerland

**Supporting information for the identification of differentially abundant proteins (DAPs) using DAPAR v.1.16.11 implemented in ProStaR v.1.16.10.**

Raw intensity values were used as input, log2-transformed, LOESS-normalized (span = 0.7), partially observed values (POVs) imputed via the Structured Least Square Adaptive (SLSA) algorithm and differential abundance analysis was conducted via LIMMA. A *p*-value push was performed on the whole matrix. This functionality introduces a filtering step that only applies to each pairwise comparison, and which assigns a *p*-value of 1 to proteins that, for the considered comparison are assumed meaningless due to too many missing values (before imputation; $n$ = 12). Proteins were deemed significantly differentially abundant in pairwise comparisons between the three sampling regions if if their $\log_2$ fold-change (FC) was **> 3** (for higher abundant proteins) or **< -3** (for lower abundant proteins) and **p < 0.01**. P-value calibration was performed with the "numeric value" function (proportion of true null hypothesis = 0.05, $n$ bins = 80). This resulted in the following false discovery rates (FDRs): FDR = **1.12%** for R-H pairwise comparison; FDR = **1.44%** for R-BF pairwise comparison; FDR = **1.34%** for H-BF pairwise comparison.

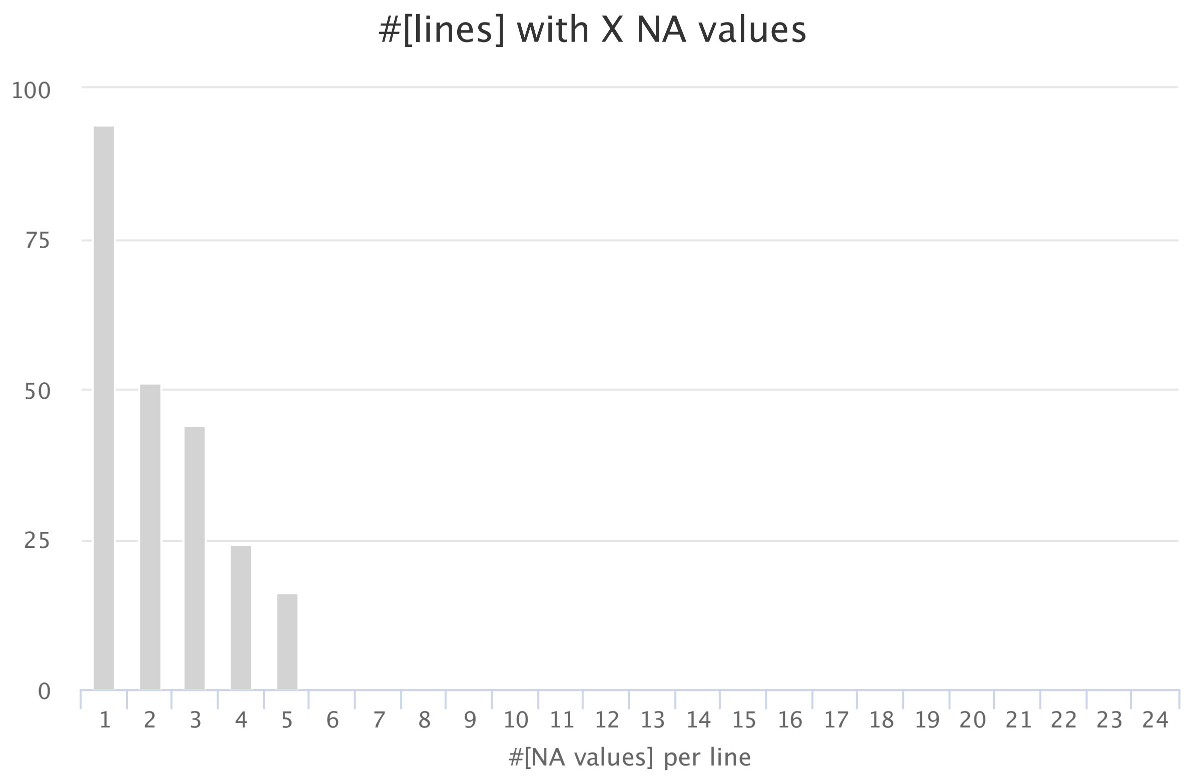


**SI4 Figure 1**: Barplot of number of lines (# of proteins; y-axis) with X missing values (NAs; x-axis).

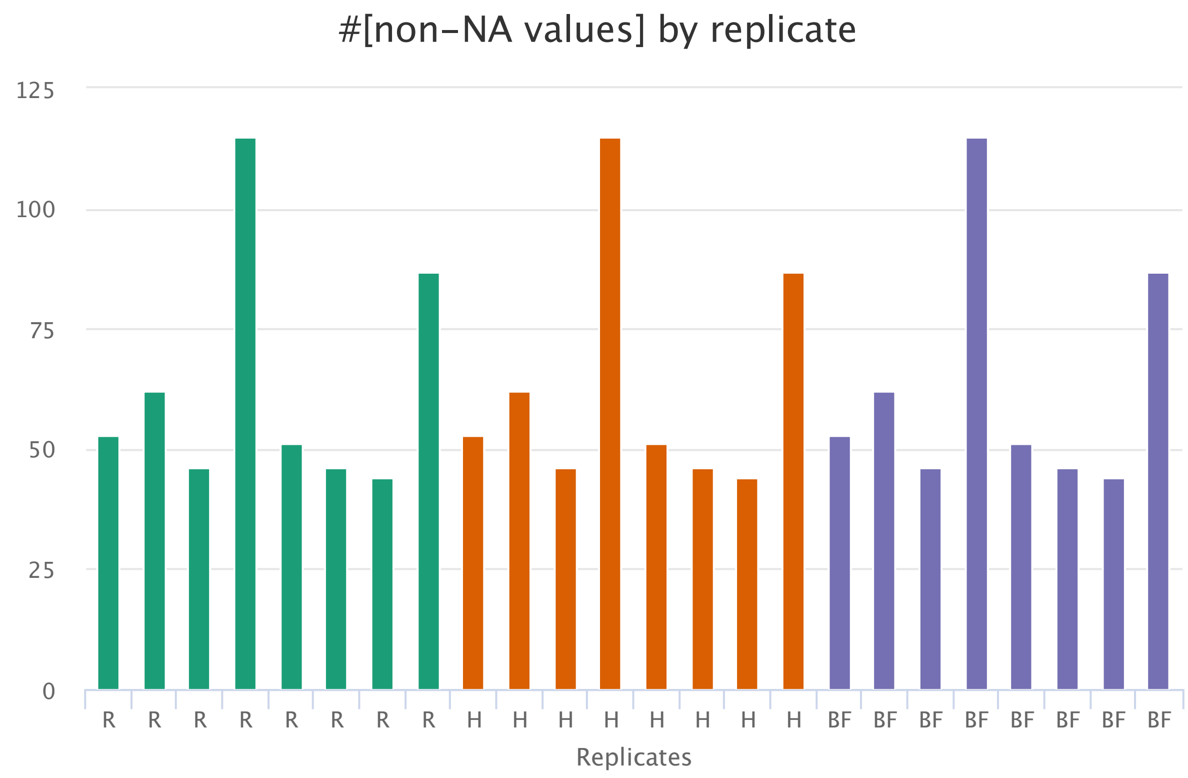


**SI4 Figure 2**: Number of missing values by sample (coloured by sampling region (condition): R = Rhoen, H = Harz, BF = Black forest.

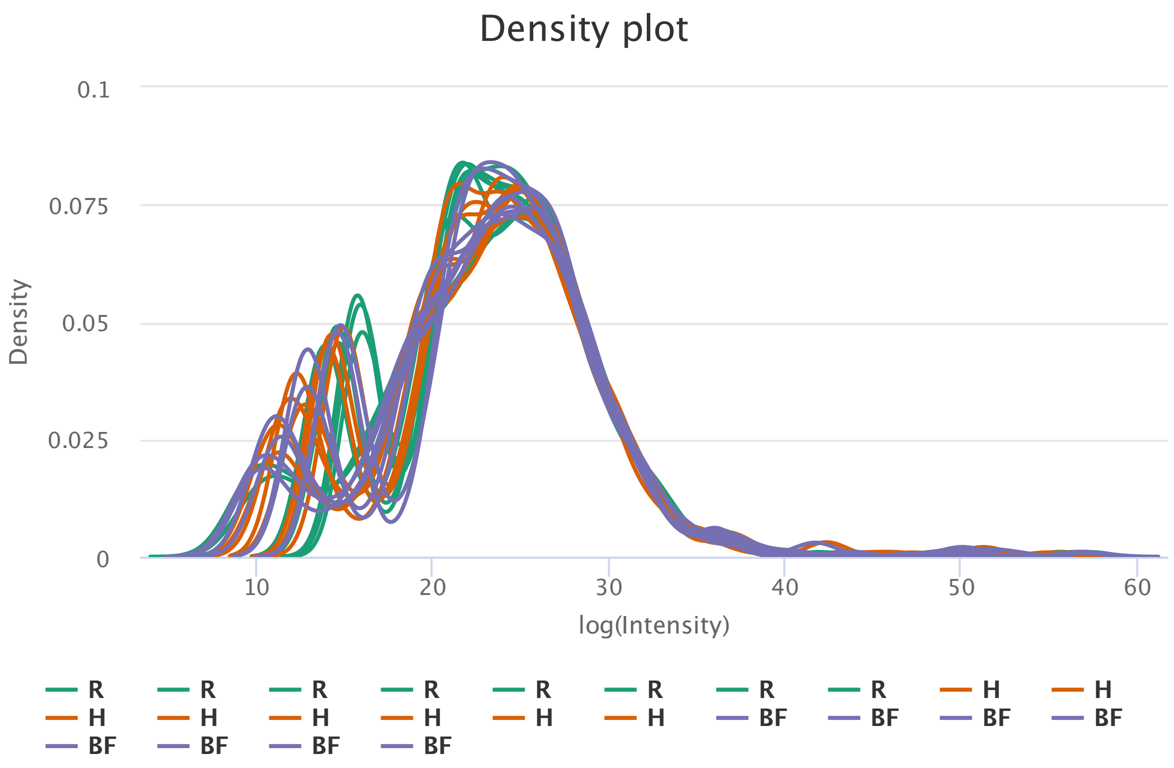


**SI4 Figure 3**: Density plot of all samples

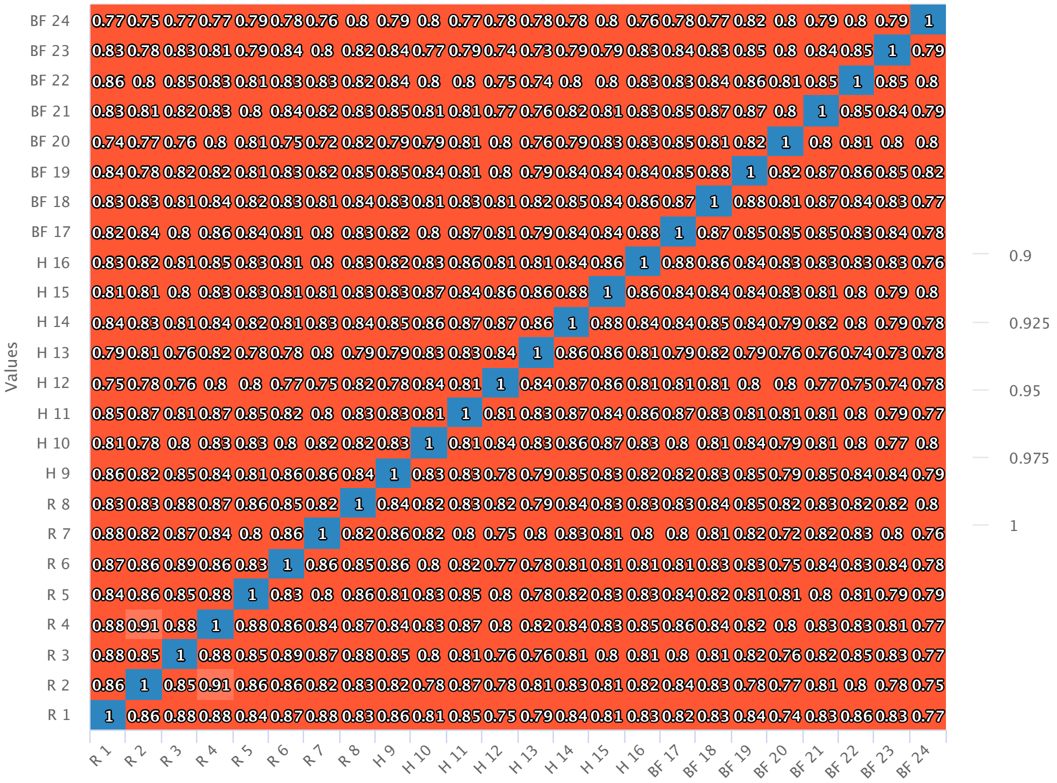


**SI4 Figure 4**: Correlation matrix between samples

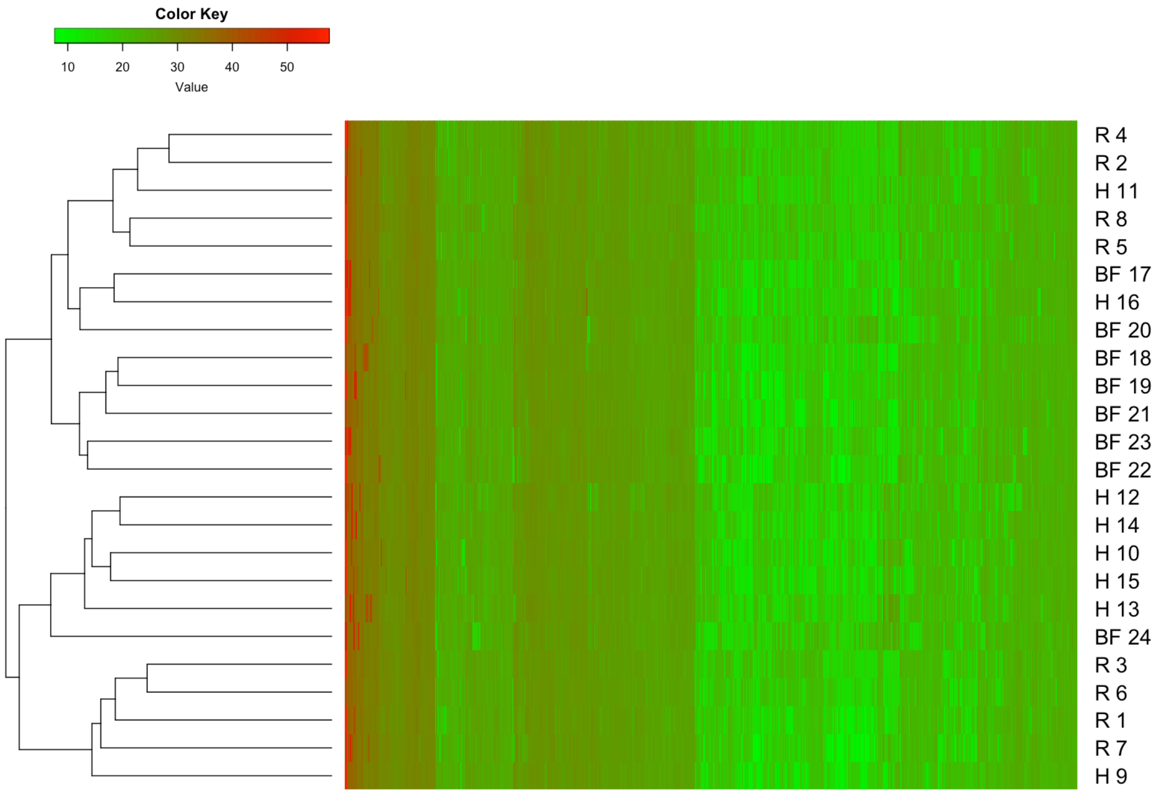


**SI4 Figure 5**: Heatmap of samples based on Minkowski distances (linkage = complete).
