## Supplemental Information 3 for "Abiotic drivers of protein abundance variation among natural populations": QC-ID-Overlap.pdf

### Pairwise peptide identification overlap (only peptides with at least 1 peptide detected and identified)

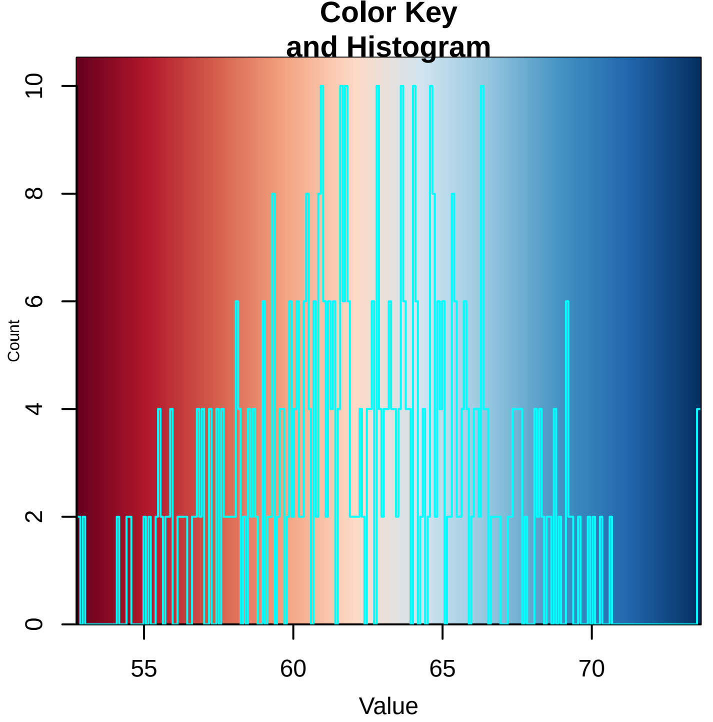


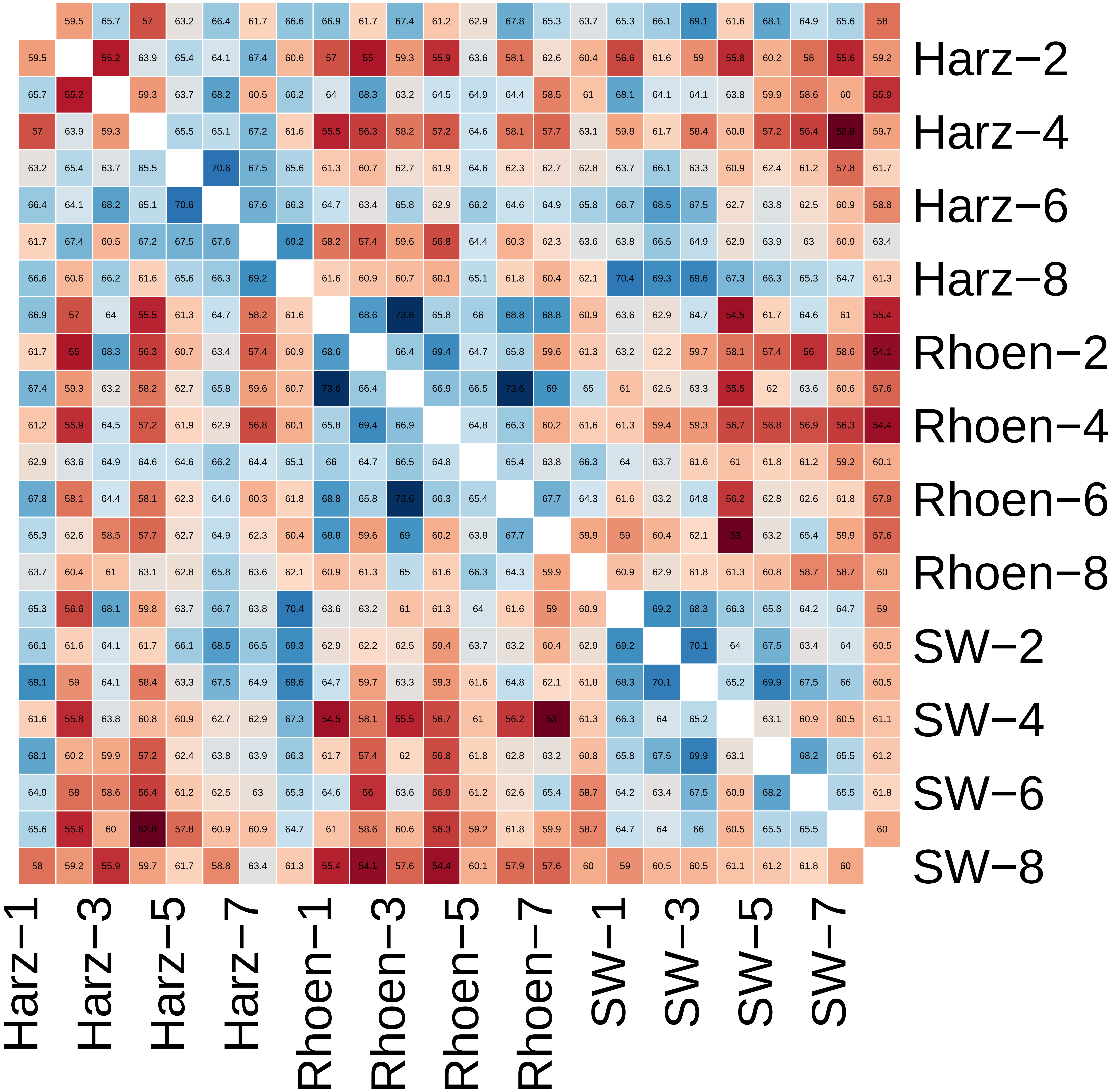

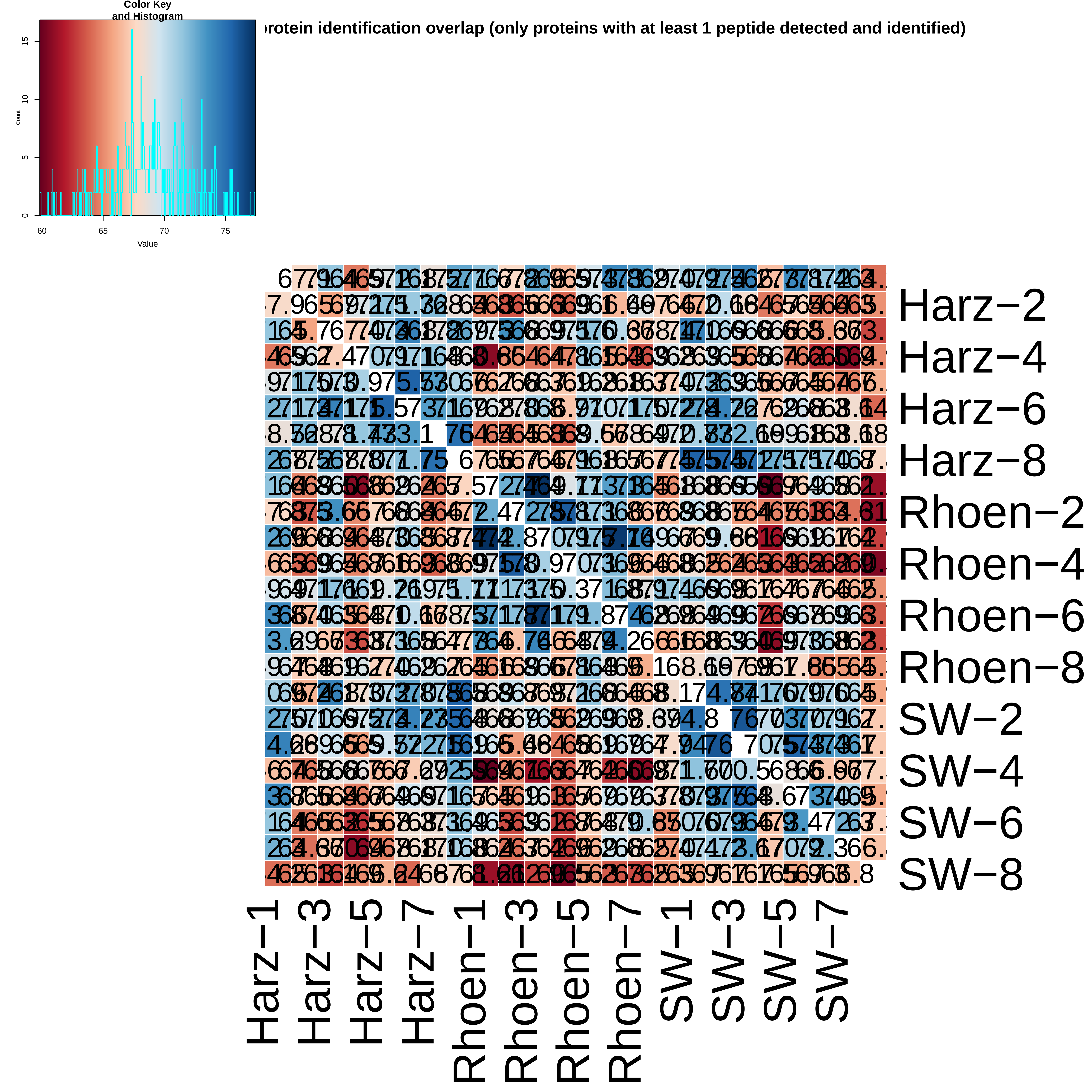

rotein identification overlap (only proteins with at least 1 peptide detected and identified)
Color Key and Histogram
Count
Value
0
5
10
15
60
65
70
75
Harz−2
Harz−4
Harz−6
Harz−8
Rhoen−2
Rhoen−4
Rhoen−6
Rhoen−8
SW−2
SW−4
SW−6
SW−8
Harz−1
Harz−3
Harz−5
Harz−7
Rhoen−1
Rhoen−3
Rhoen−5
Rhoen−7
SW−1
SW−3
SW−5
SW−7
