## Supplemental Information 3 for "Abiotic drivers of protein abundance variation among natural populations": QC_Plots_PEPINT.pdf

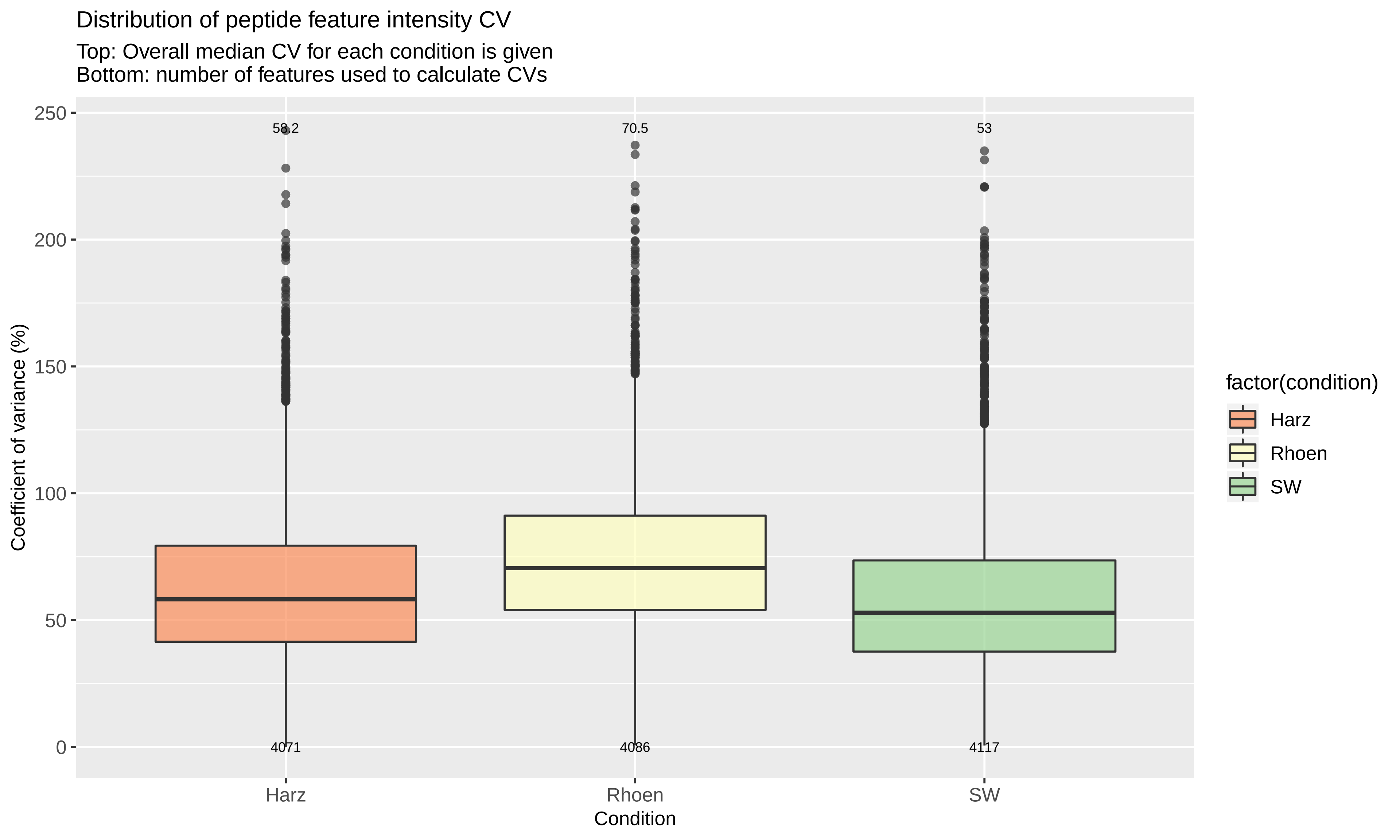
Distribution of peptide feature intensity CV
Top: Overall median CV for each condition is given
Bottom: number of features used to calculate CVs
58.2
70.5
53
4071
4086
4117
250
200
150
100
50
0
Coefficient of variance (%)
Harz
Rhoen
SW
Condition
factor(condition)
Harz
Rhoen
SW

### Distribution of peptide feature intensity CV

For each condition, peptides were ranked by summed intensity and the CV for each peptide was calculated, therefore each condition shows 4 distribution (box) for low (1) to high (4) intensity peptides.
Overall median CV within each bin/condition is shown on the top and number of features used to calculate CV is given on the bottom

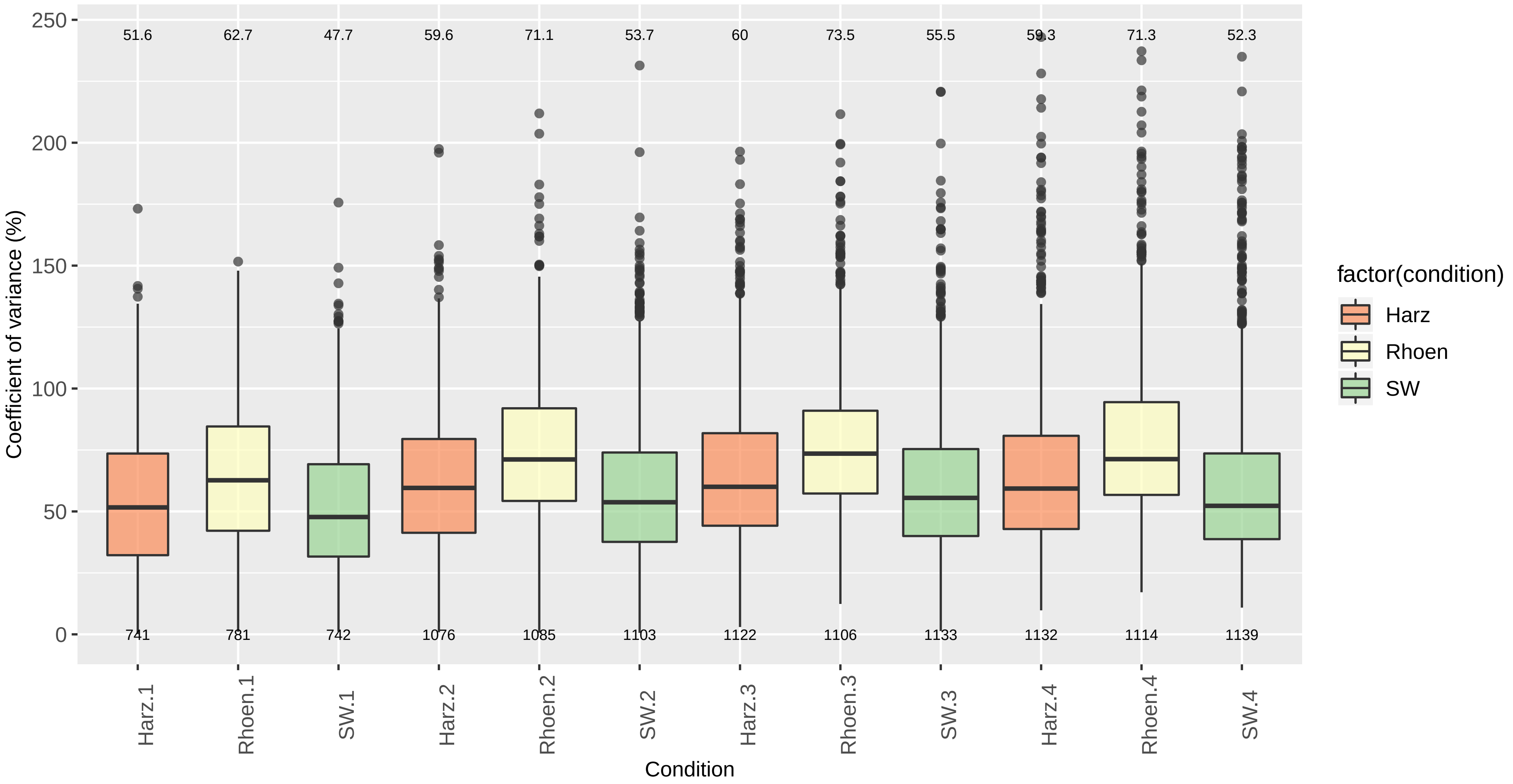
