## Supplemental Information 3 for "Abiotic drivers of protein abundance variation among natural populations": QC_Plots_ProtInt.pdf

Distribution of Protein intensity CV

Top: Overall median CV for each condition.
Bottom: number of proteins used to calculate CVs

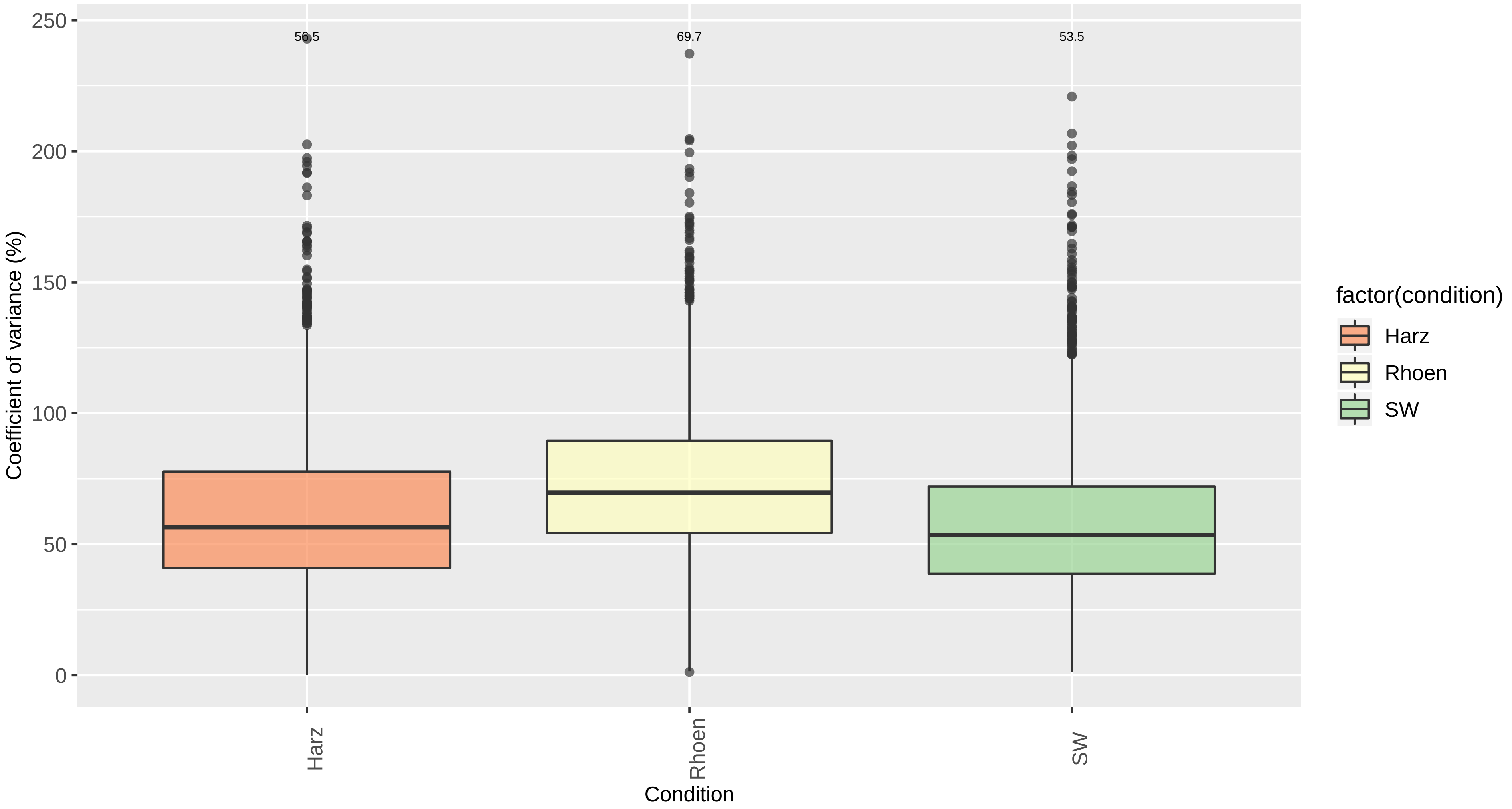

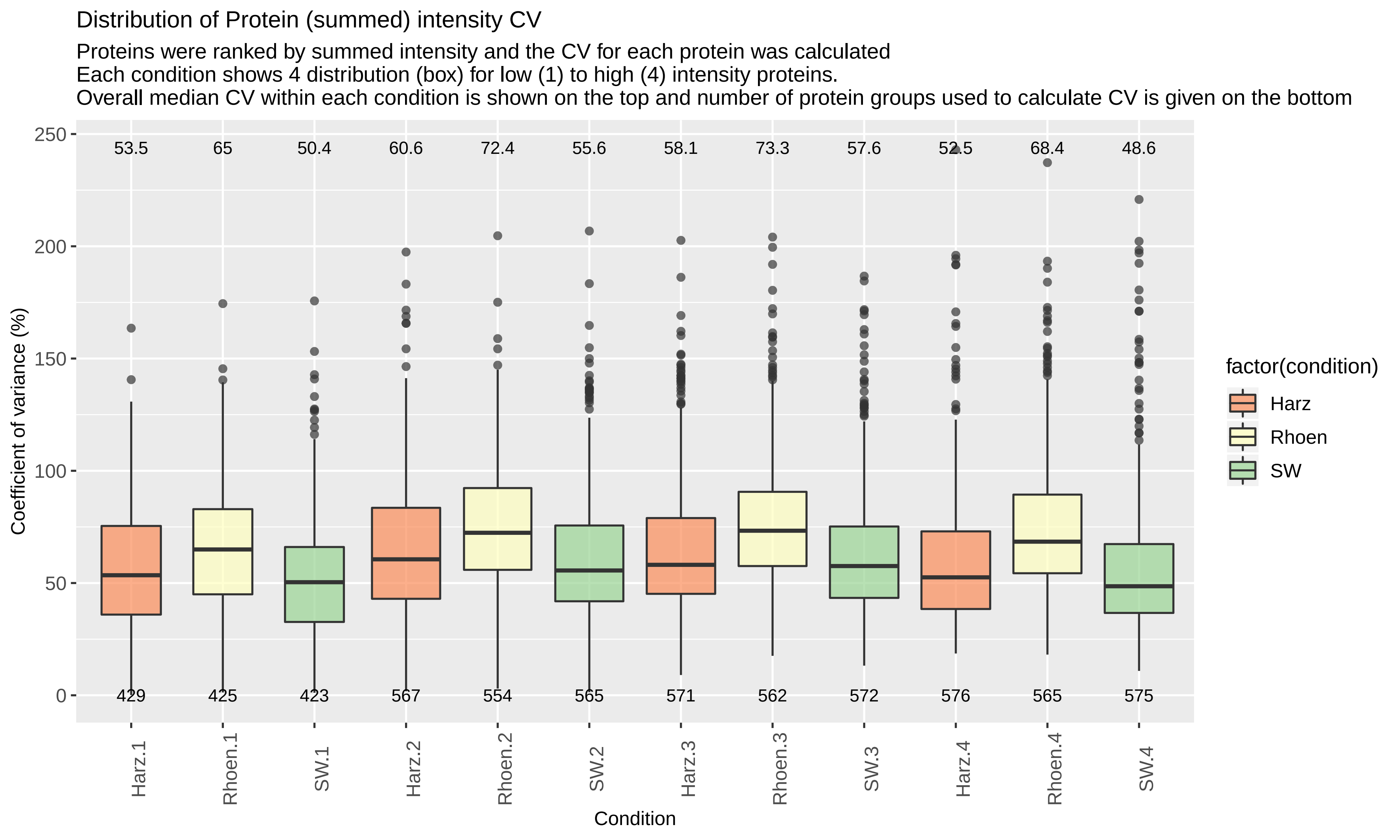

Distribution of Protein (summed) intensity CV
Proteins were ranked by summed intensity and the CV for each protein was calculated
Each condition shows 4 distribution (box) for low (1) to high (4) intensity proteins.
Overall median CV within each condition is shown on the top and number of protein groups used to calculate CV is given on the bottom
Coefficient of variance (%)
250
200
150
100
50
0
53.5
65
50.4
60.6
72.4
55.6
58.1
73.3
57.6
52.5
68.4
48.6
429
425
423
567
554
565
571
562
572
576
565
575
Harz.1
Rhoen.1
SW.1
Harz.2
Rhoen.2
SW.2
Harz.3
Rhoen.3
SW.3
Harz.4
Rhoen.4
SW.4
Condition
factor(condition)
Harz
Rhoen
SW
