## Supplemental Information 3 for "Abiotic drivers of protein abundance variation among natural populations": qcPlots_evidence.qcplot.basicReproducibility.pdf

Peptide Reproducibility between Bioreplicas
(condition: Rhoen ) Rhoen−1 vs Rhoen−2
(n = 5763  r =  0.79 )

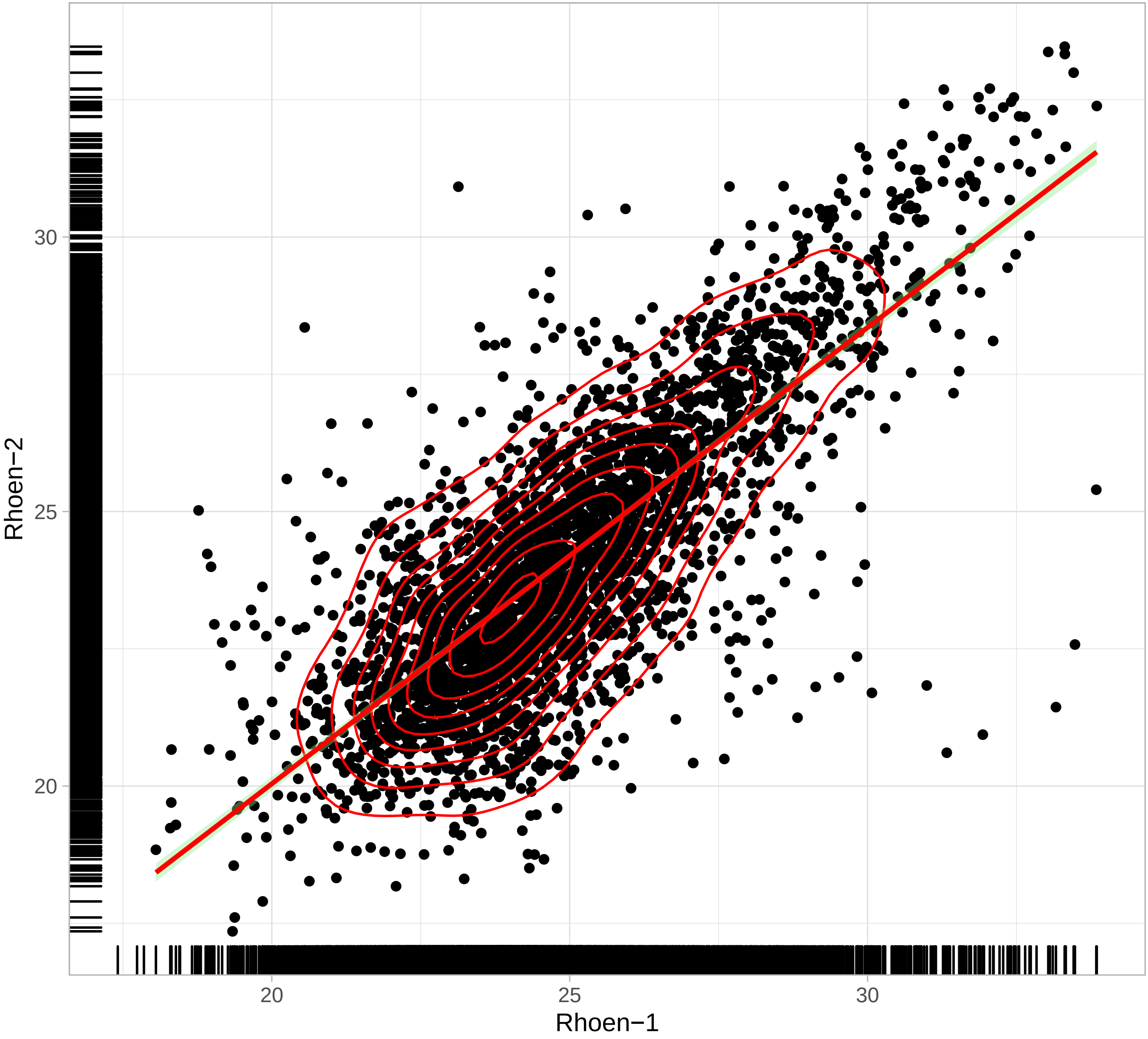

Peptide Reproducibility between Bioreplicas
(condition: Rhoen ) Rhoen−1 vs Rhoen−3
(n = 5763 r = 0.87 )

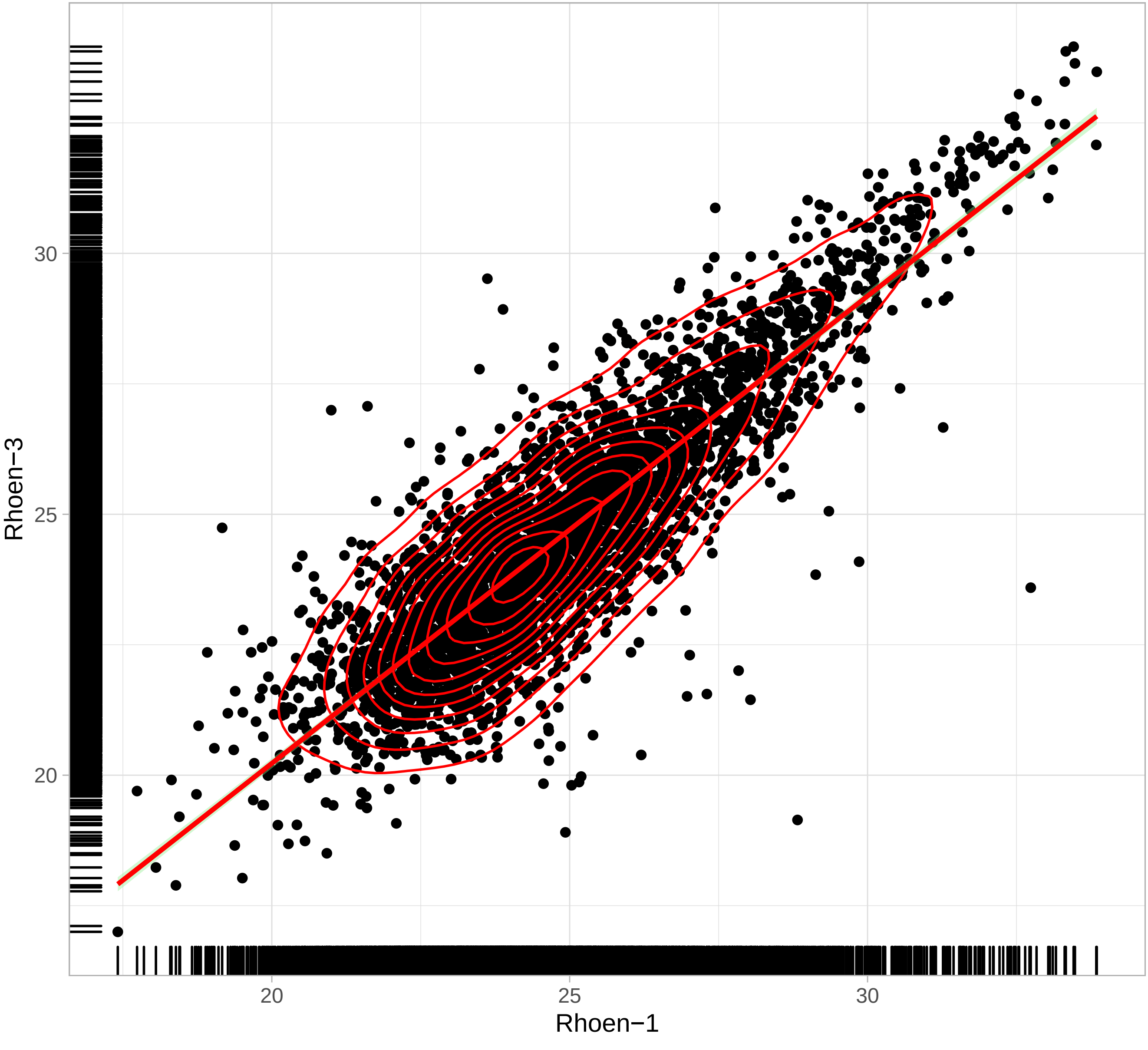

Peptide Reproducibility between Bioreplicas
 (condition: Rhoen ) Rhoen−1 vs Rhoen−4
(n = 5763  r =  0.8 )

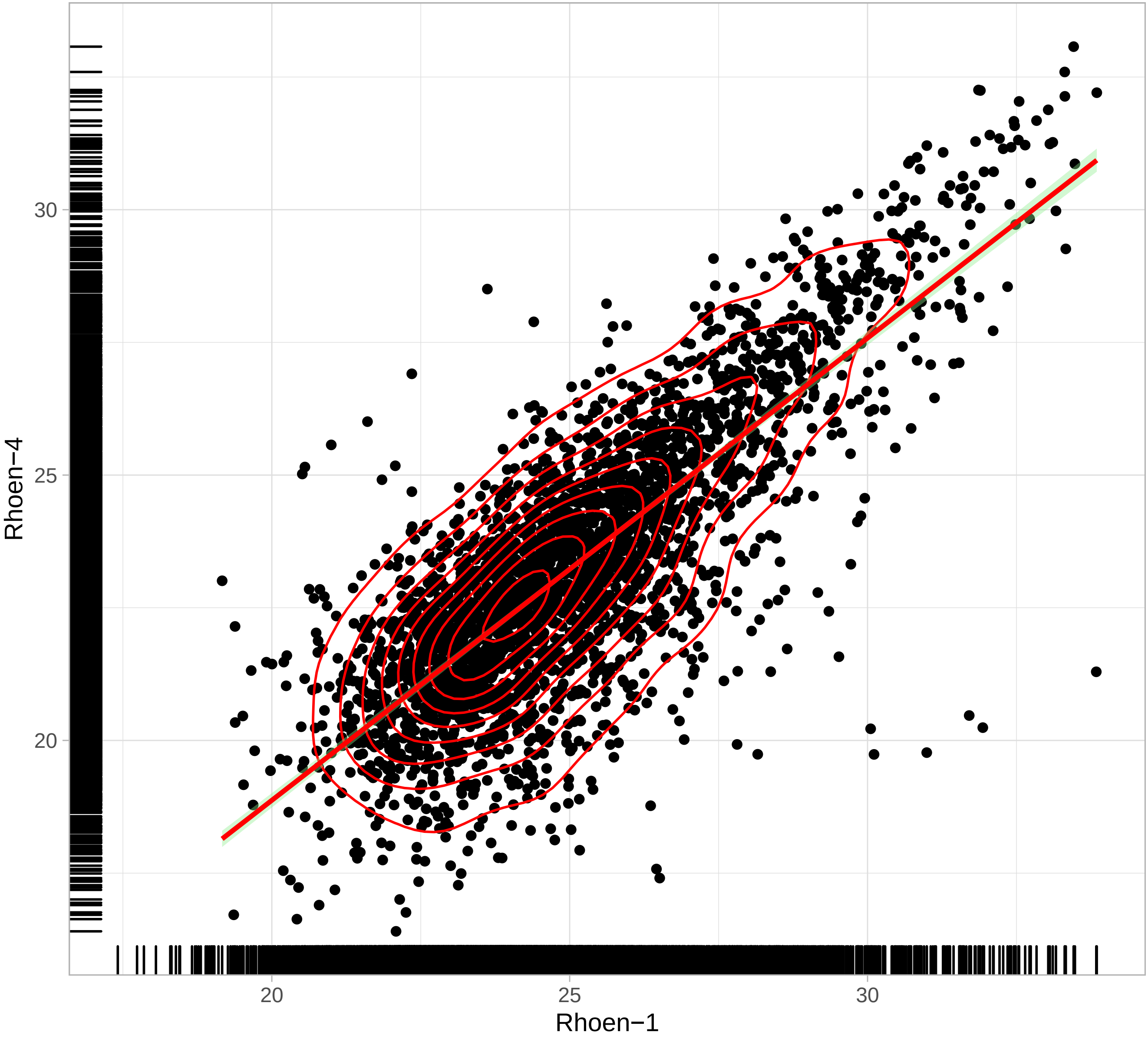

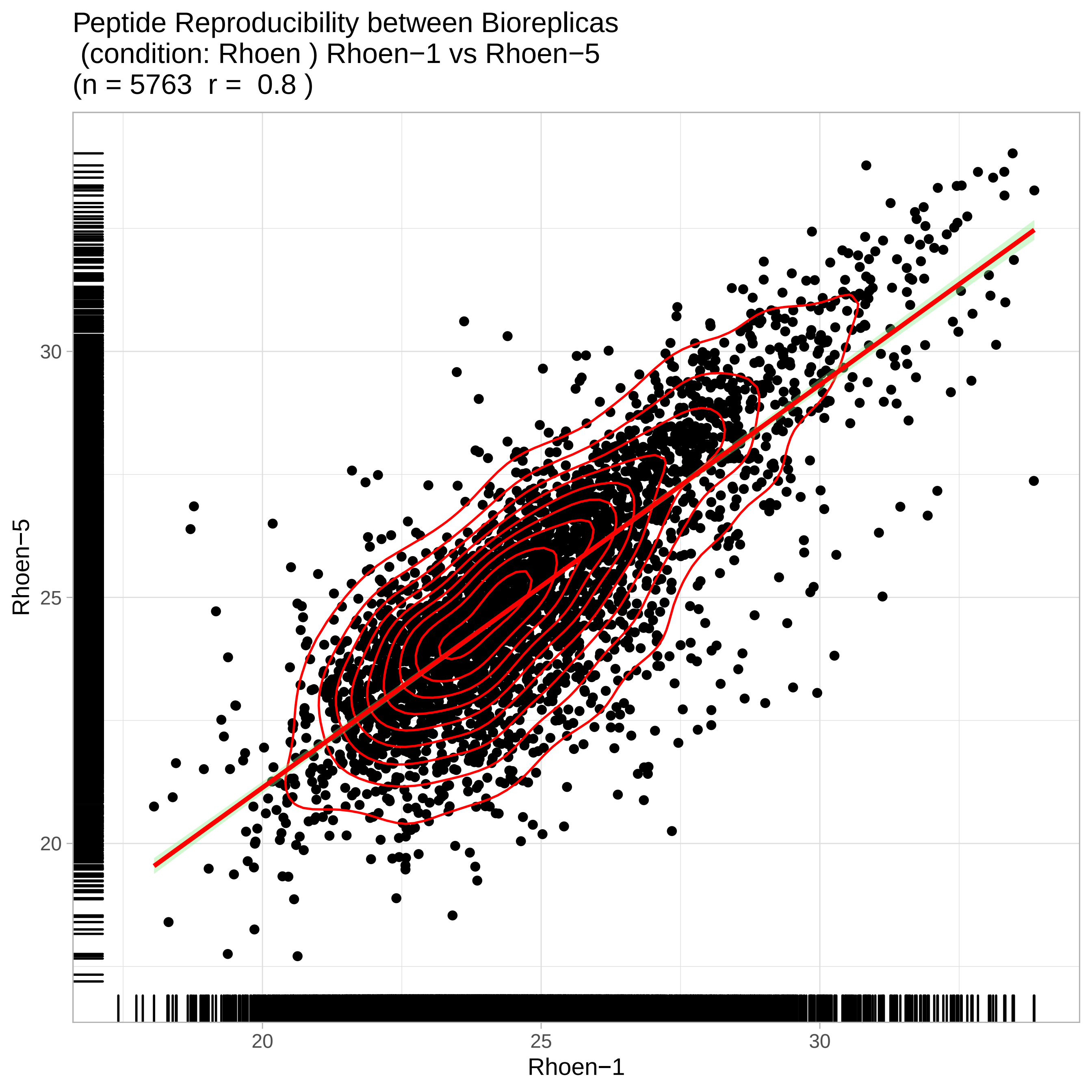
Peptide Reproducibility between Bioreplicas
(condition: Rhoen ) Rhoen−1 vs Rhoen−5
(n = 5763 r = 0.8 )
Rhoen−5
30
25
20
20
25
30
Rhoen−1

Peptide Reproducibility between Bioreplicas
(condition: Rhoen ) Rhoen−1 vs Rhoen−6
(n = 5763 r = 0.84 )

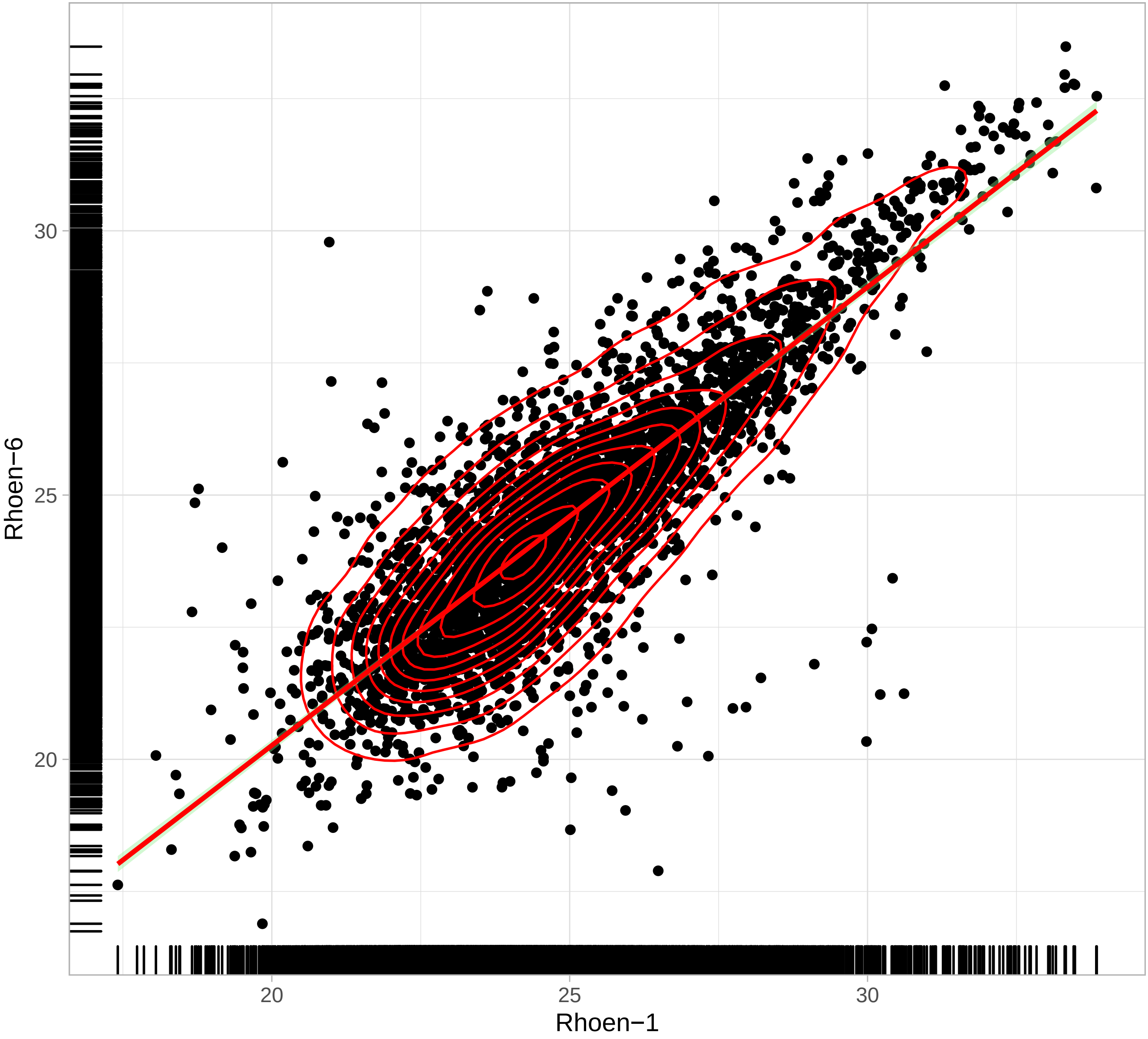

Peptide Reproducibility between Bioreplicas
(condition: Rhoen ) Rhoen−1 vs Rhoen−7
(n = 5763  r =  0.8 )

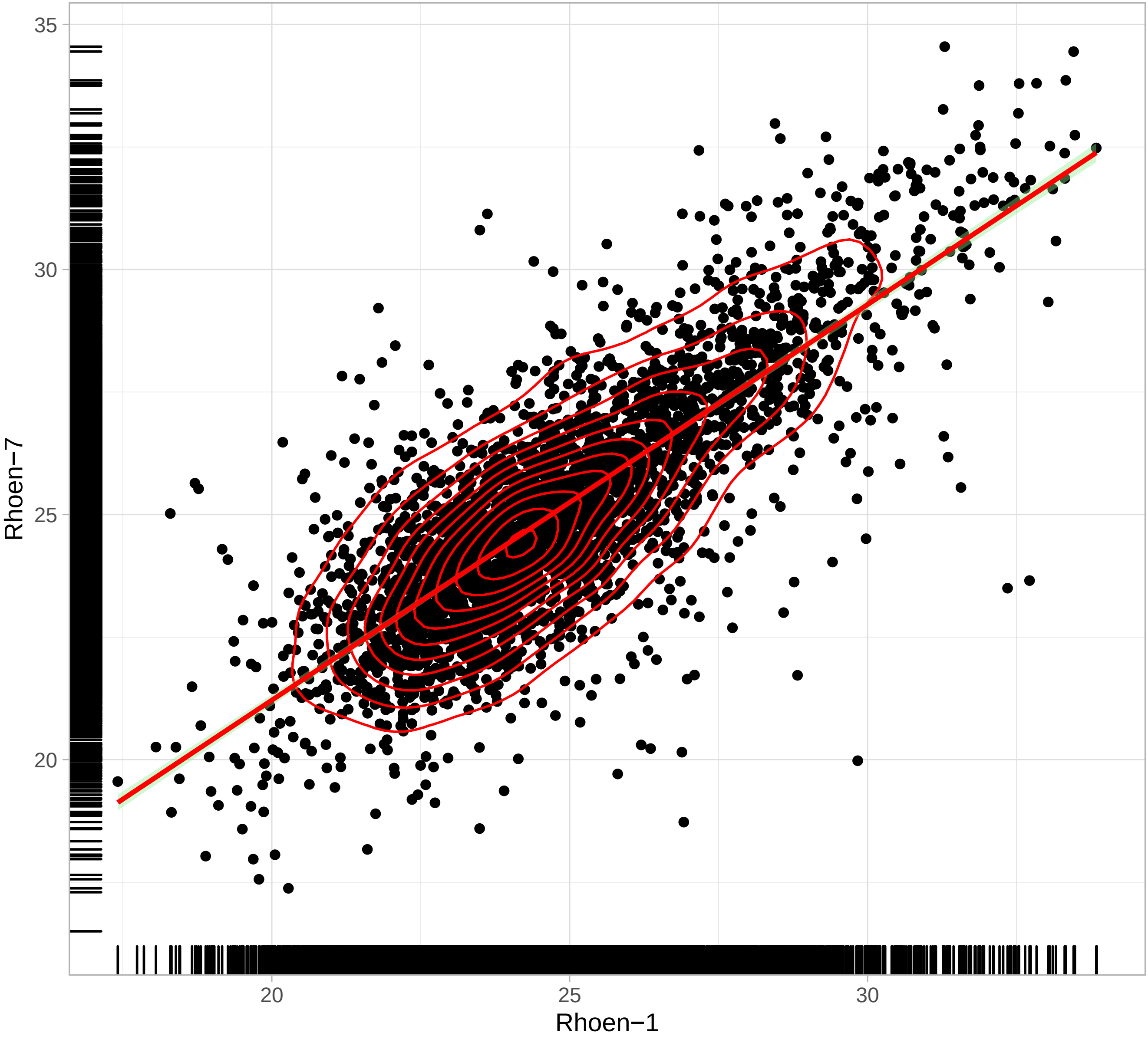

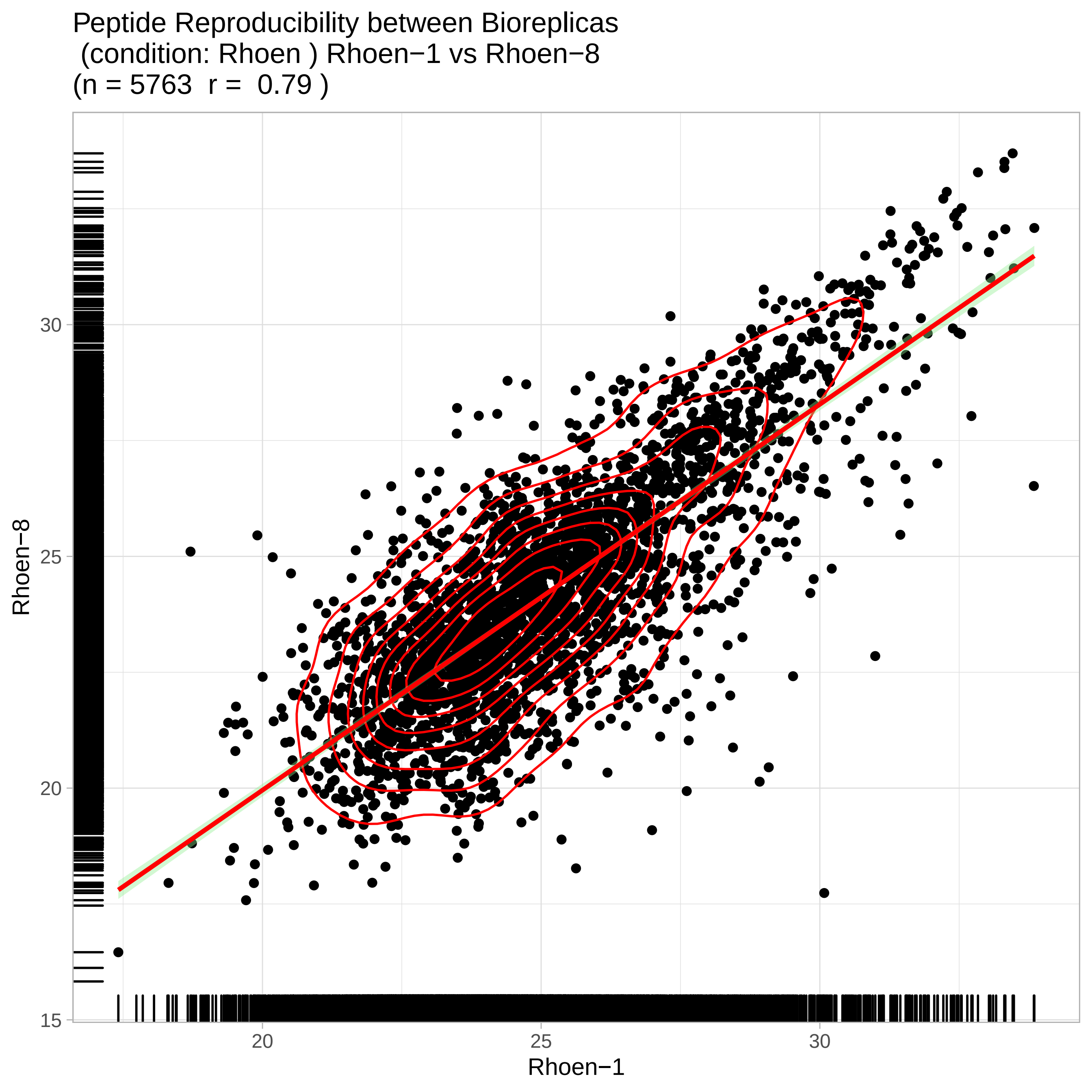
Peptide Reproducibility between Bioreplicas
(condition: Rhoen ) Rhoen−1 vs Rhoen−8
(n = 5763 r = 0.79 )
Rhoen−8
Rhoen−1
15
20
25
30
20
25
30

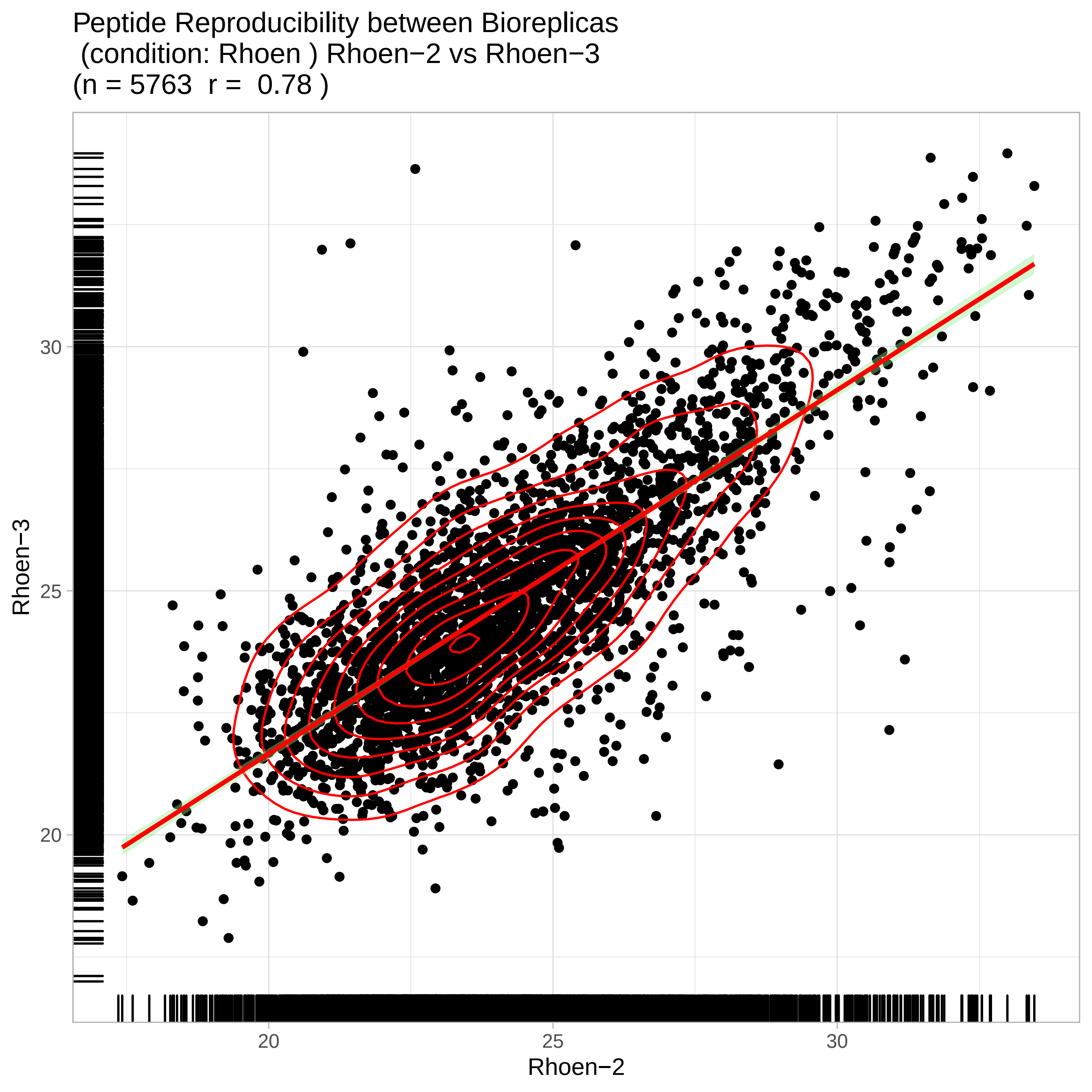
Peptide Reproducibility between Bioreplicas
(condition: Rhoen ) Rhoen−2 vs Rhoen−3
(n = 5763 r = 0.78 )
Rhoen−3
30
25
20
Rhoen−2
20
25
30

Peptide Reproducibility between Bioreplicas
(condition: Rhoen ) Rhoen−2 vs Rhoen−4
(n = 5763 r = 0.86 )

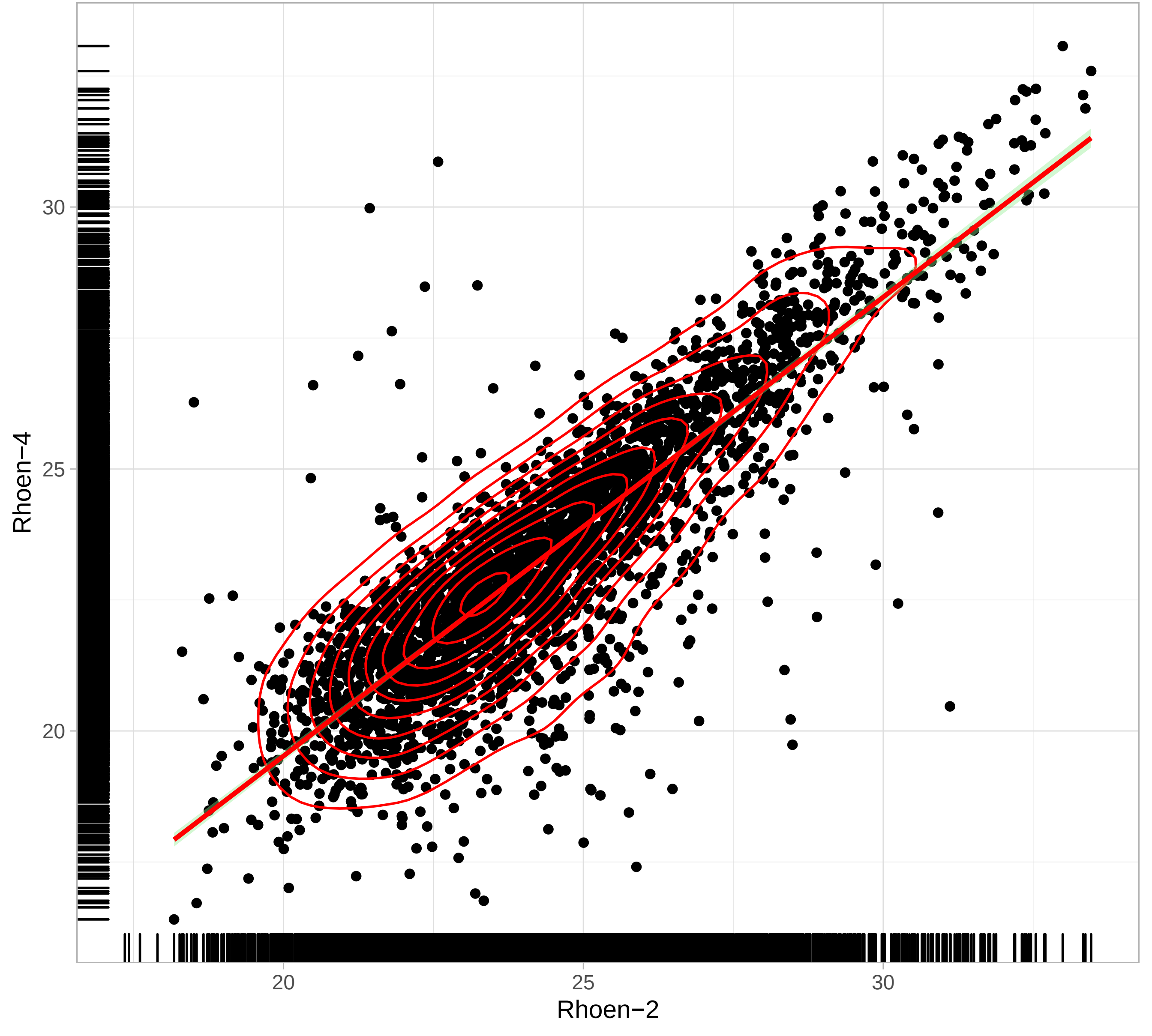

Peptide Reproducibility between Bioreplicas
(condition: Rhoen ) Rhoen−2 vs Rhoen−5
(n = 5763  r =  0.8 )

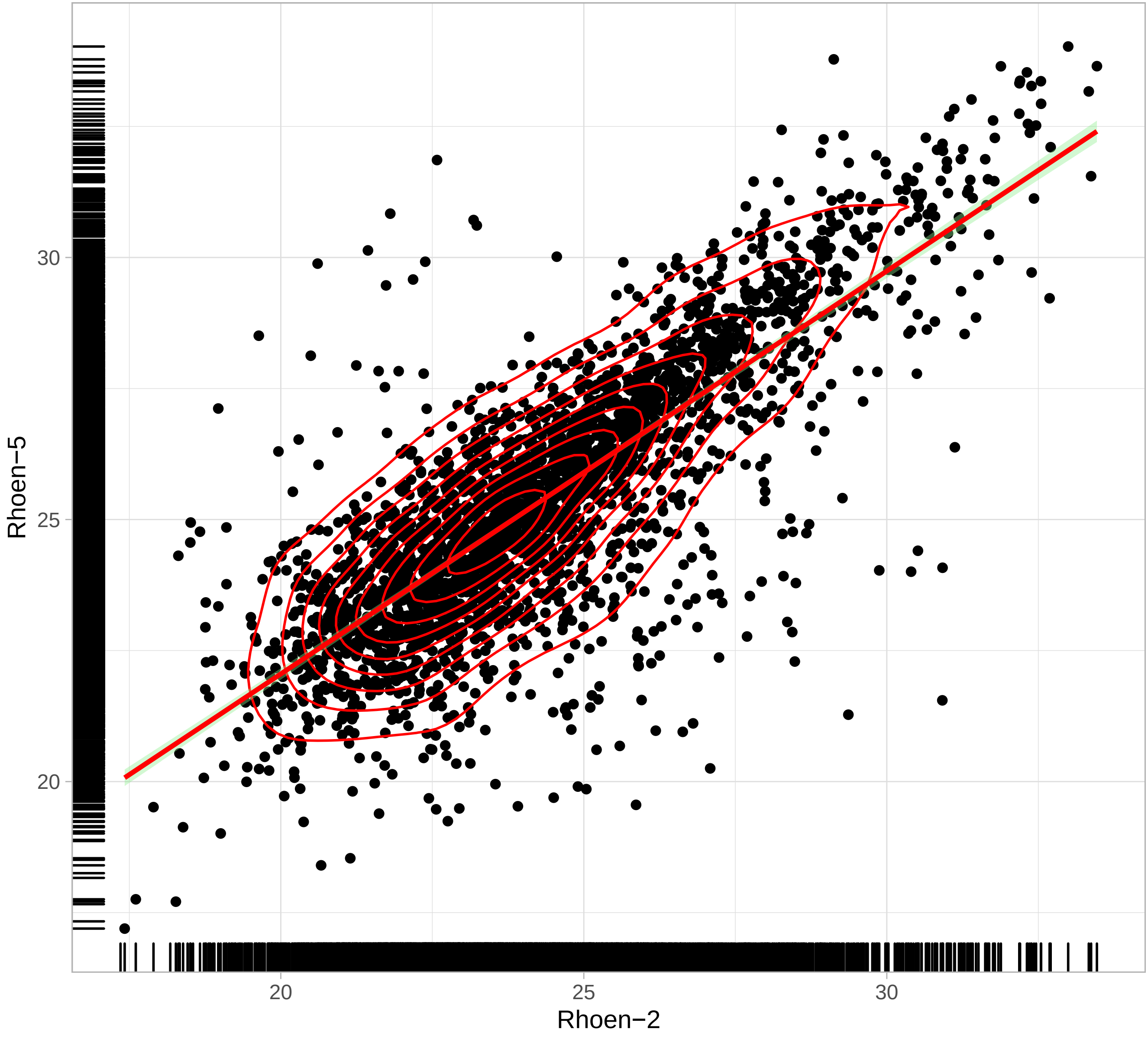

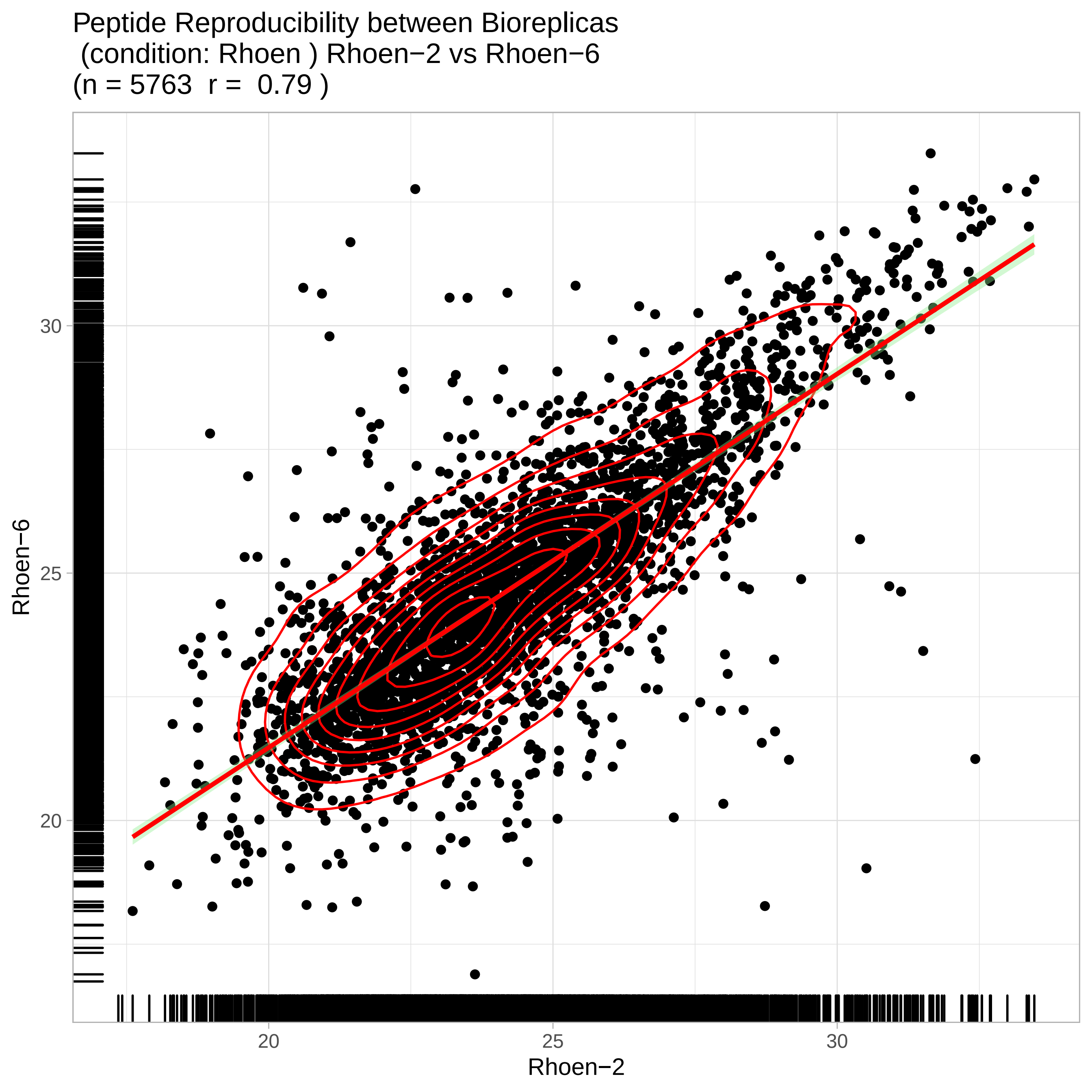
Peptide Reproducibility between Bioreplicas
(condition: Rhoen ) Rhoen−2 vs Rhoen−6
(n = 5763 r = 0.79 )
Rhoen−6
30
25
20
20
25
30
Rhoen−2

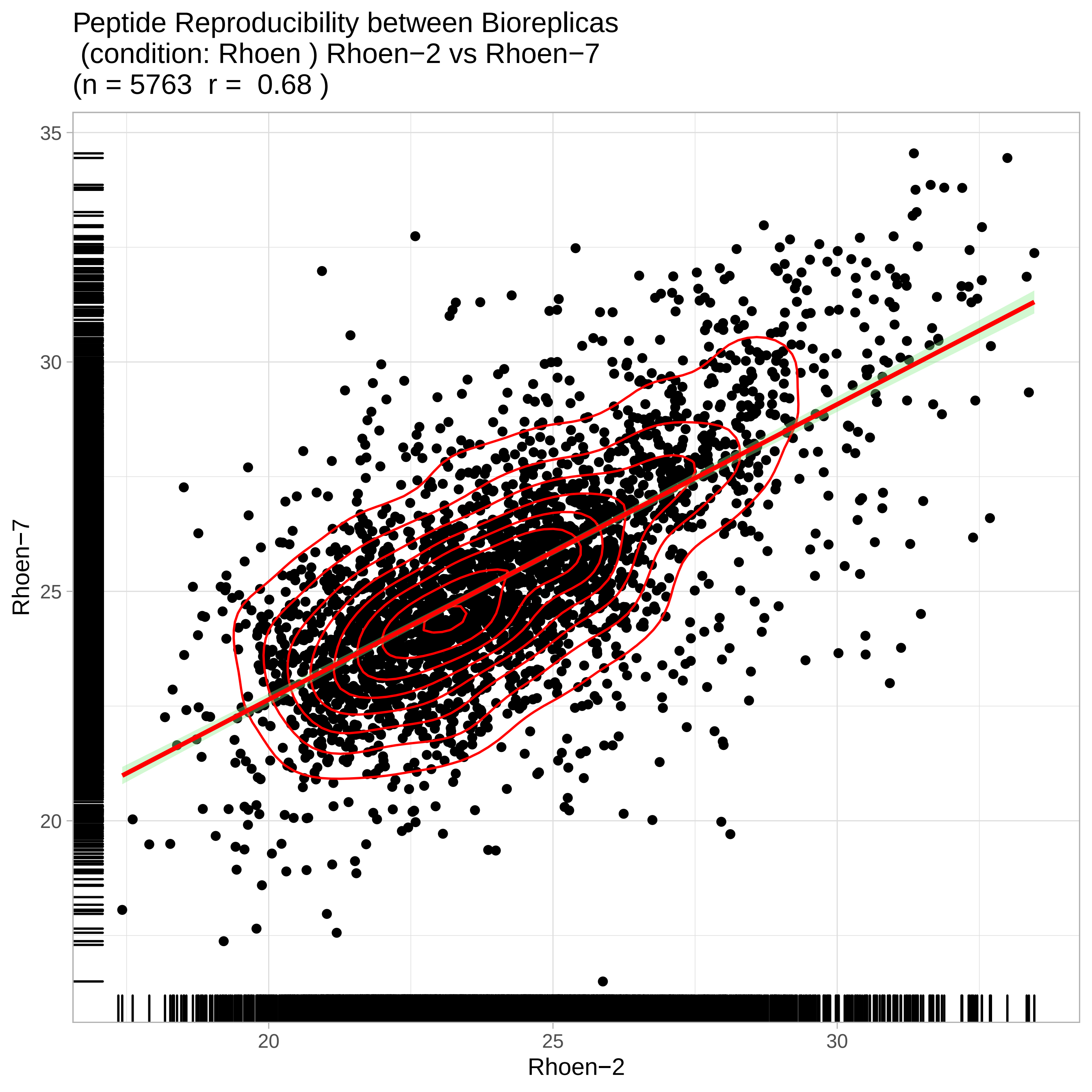

Peptide Reproducibility between Bioreplicas
(condition: Rhoen ) Rhoen−2 vs Rhoen−7
(n = 5763 r = 0.68 )
Rhoen−7
35
30
25
20
Rhoen−2
20
25
30

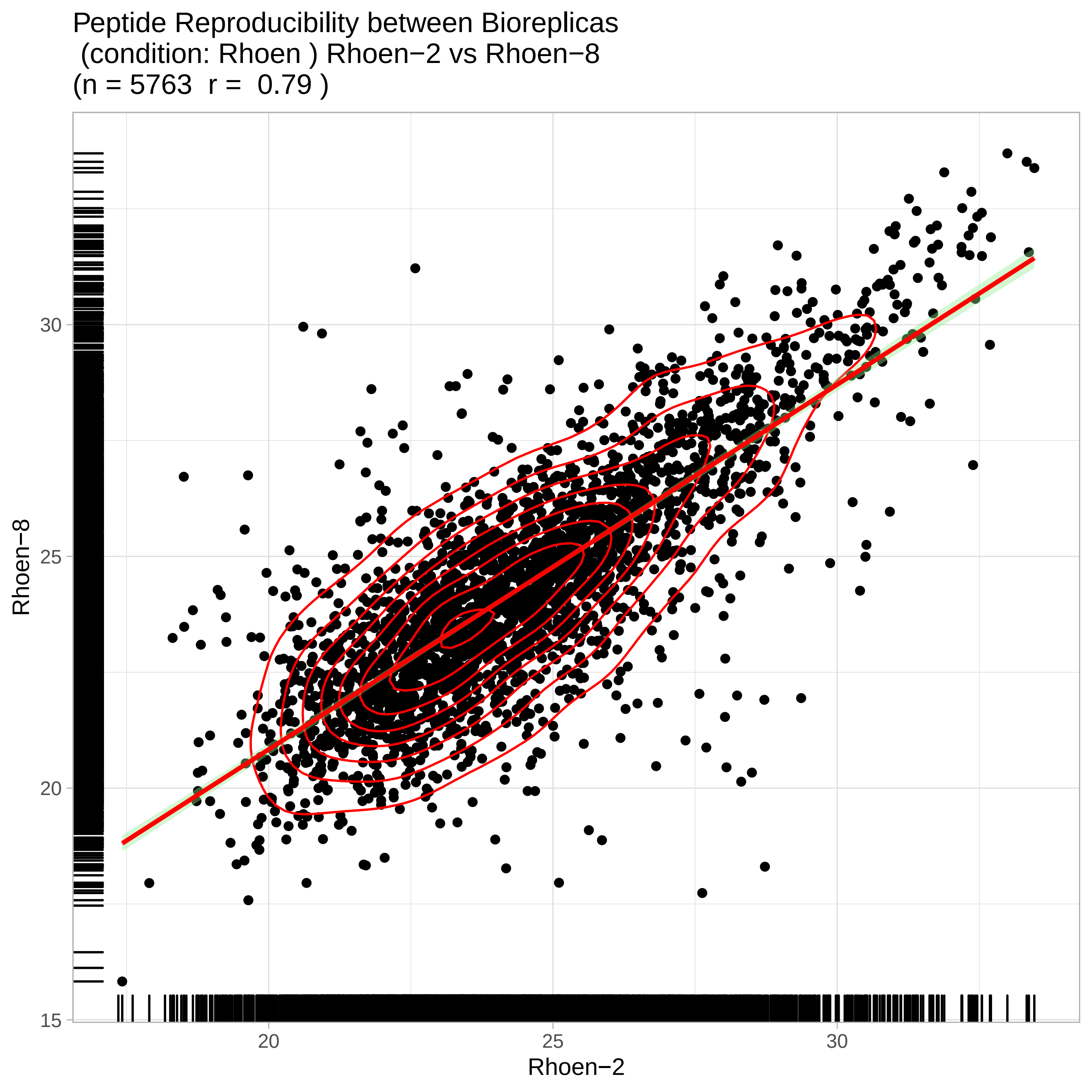
Peptide Reproducibility between Bioreplicas
(condition: Rhoen ) Rhoen−2 vs Rhoen−8
(n = 5763 r = 0.79 )
Rhoen−8
30
25
20
15
20
25
30
Rhoen−2

Peptide Reproducibility between Bioreplicas
(condition: Rhoen ) Rhoen−3 vs Rhoen−4
(n = 5763  r =  0.79 )

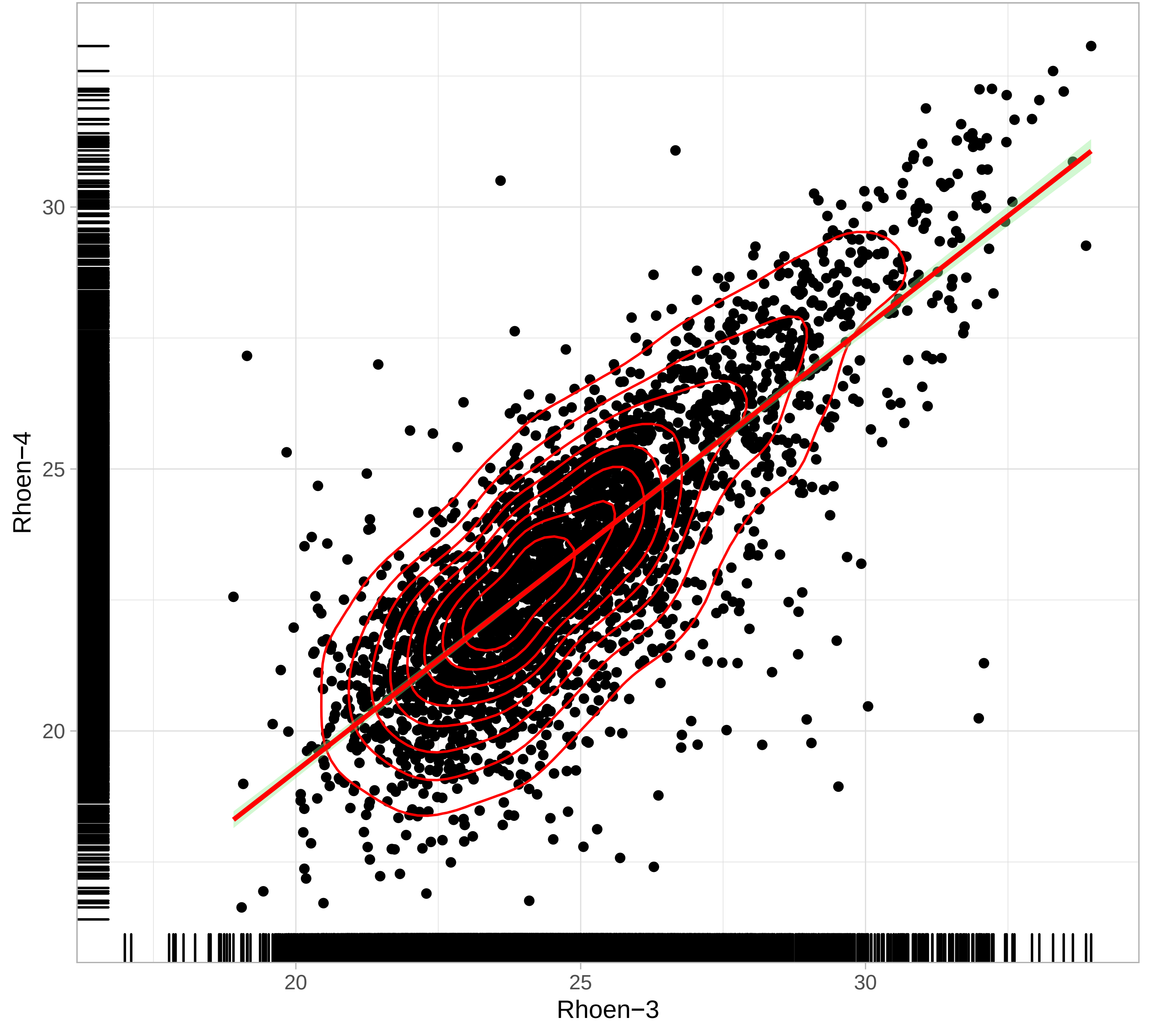

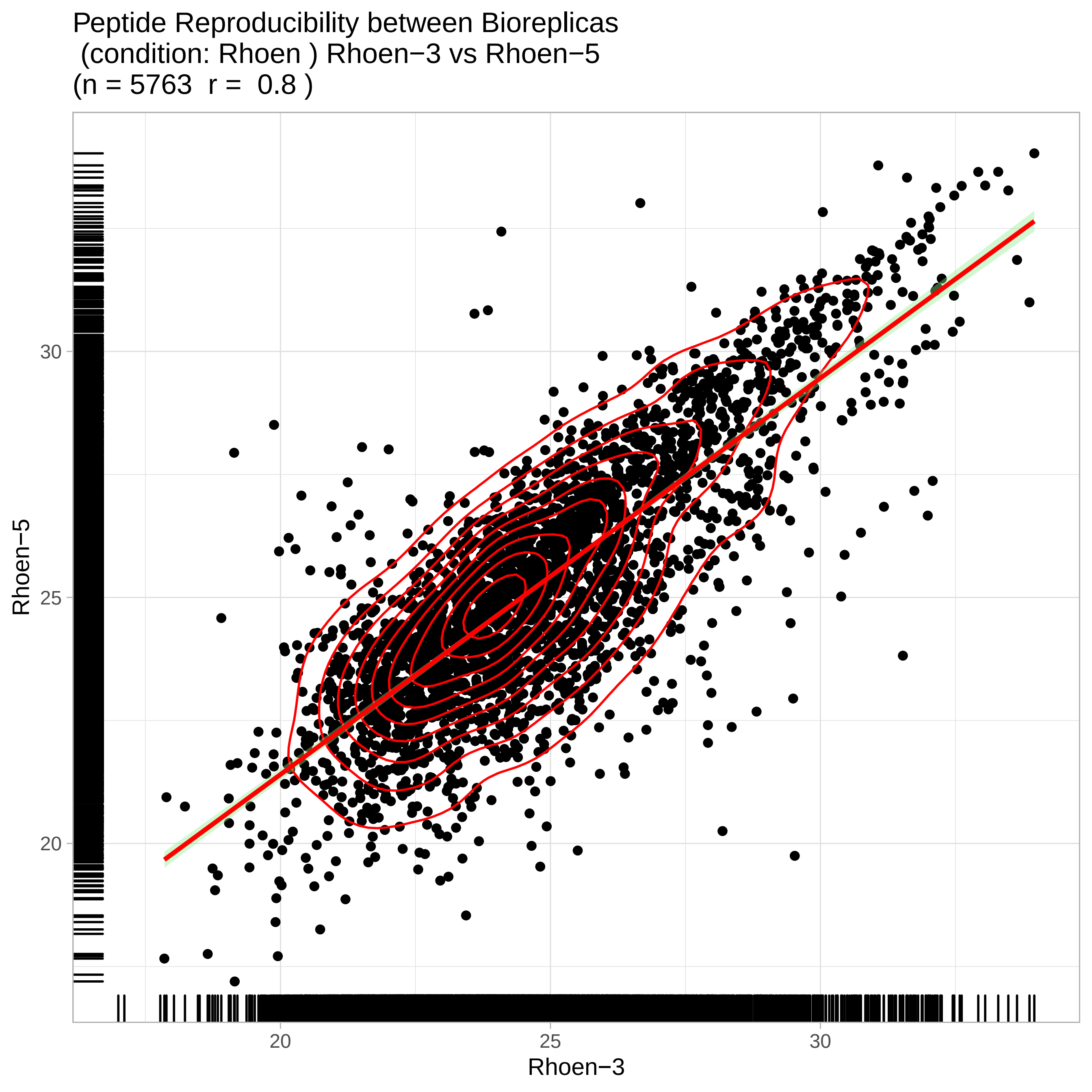
Peptide Reproducibility between Bioreplicas
(condition: Rhoen ) Rhoen−3 vs Rhoen−5
(n = 5763 r = 0.8 )
Rhoen−5
30
25
20
20
25
30
Rhoen−3

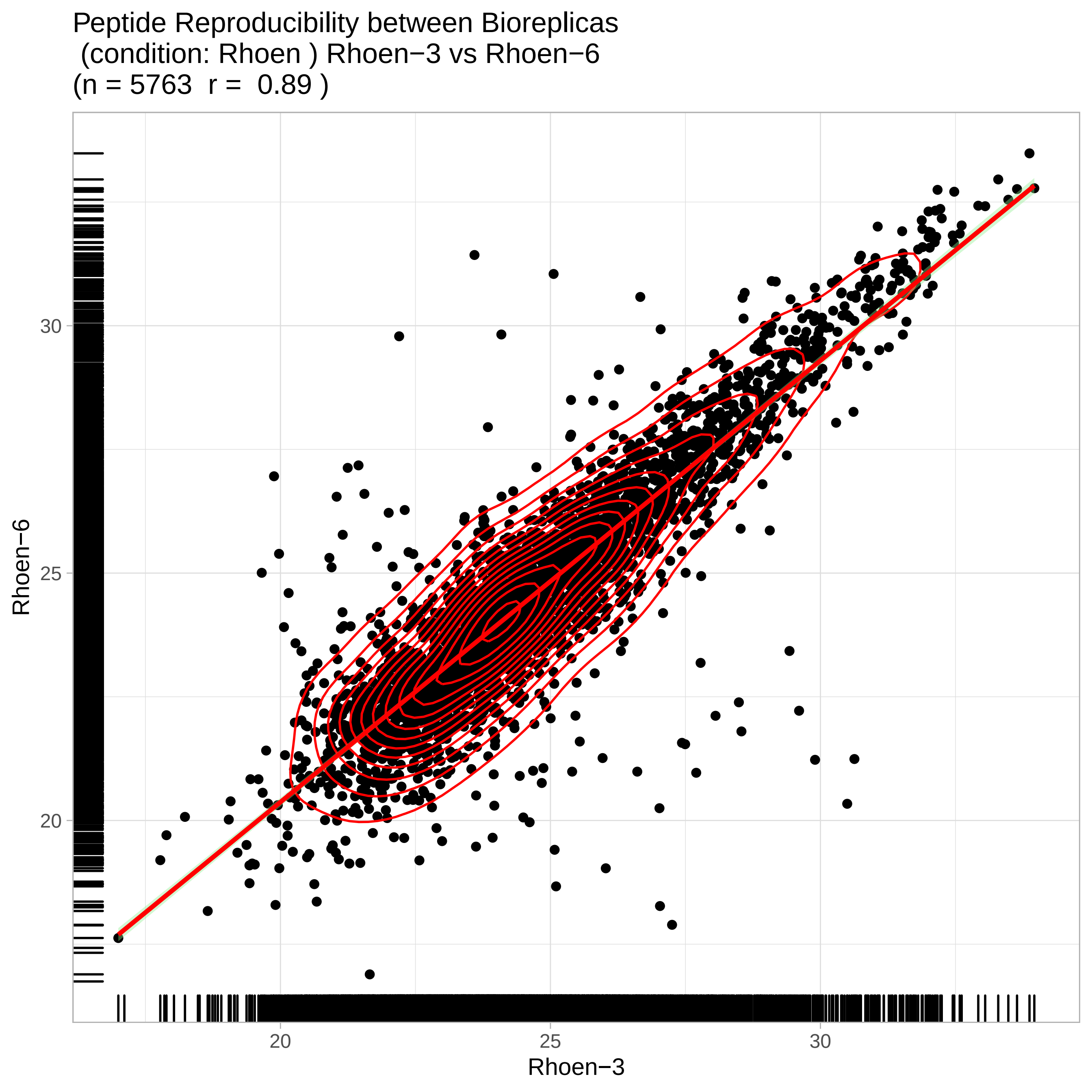

Peptide Reproducibility between Bioreplicas
(condition: Rhoen ) Rhoen−3 vs Rhoen−6
(n = 5763 r = 0.89 )
Rhoen−6
30
25
20
20
25
30
Rhoen−3

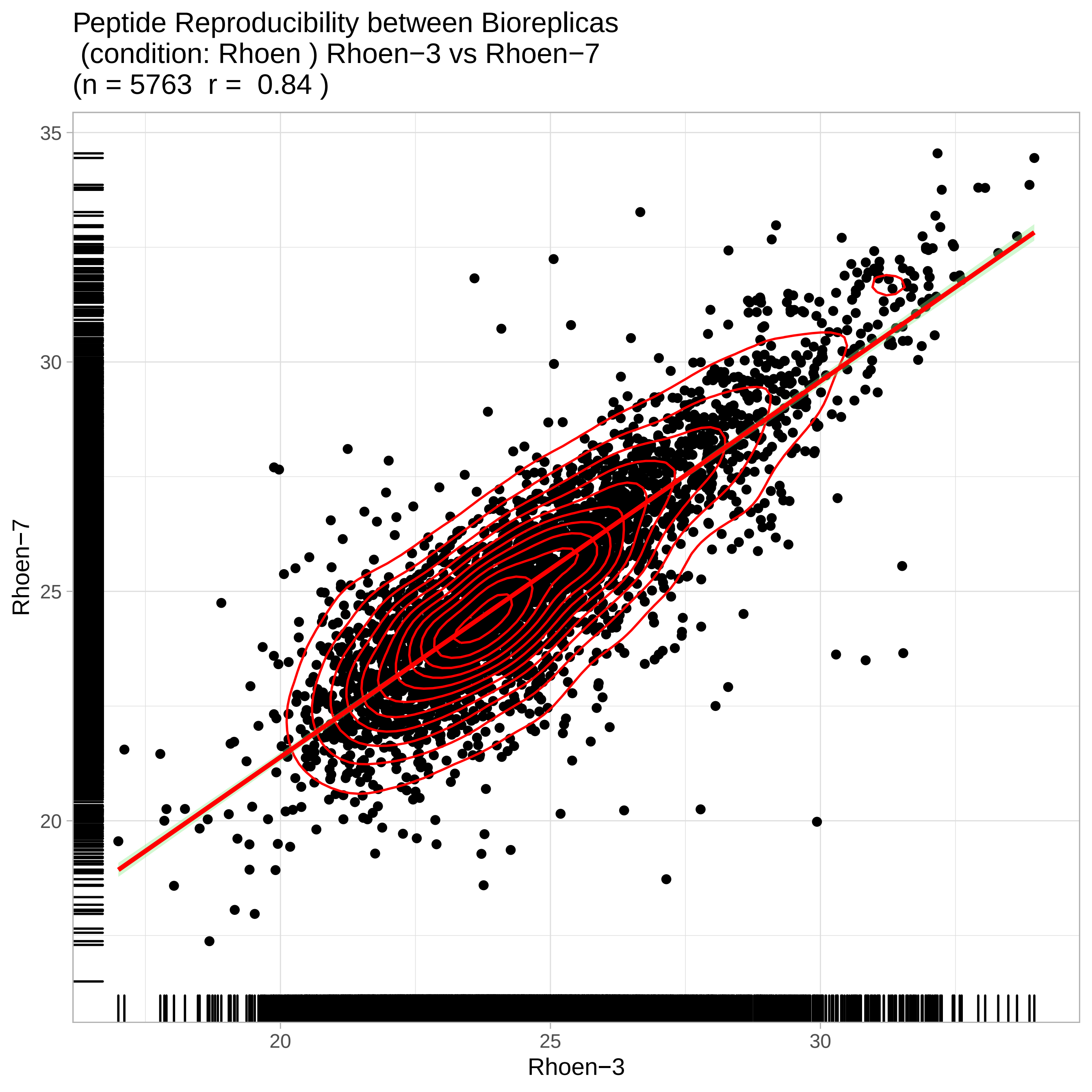

Peptide Reproducibility between Bioreplicas
(condition: Rhoen ) Rhoen−3 vs Rhoen−7
(n = 5763  r =  0.84 )
Rhoen−7
35
30
25
20
Rhoen−3
20
25
30

Peptide Reproducibility between Bioreplicas
(condition: Rhoen ) Rhoen−3 vs Rhoen−8
(n = 5763 r = 0.84 )

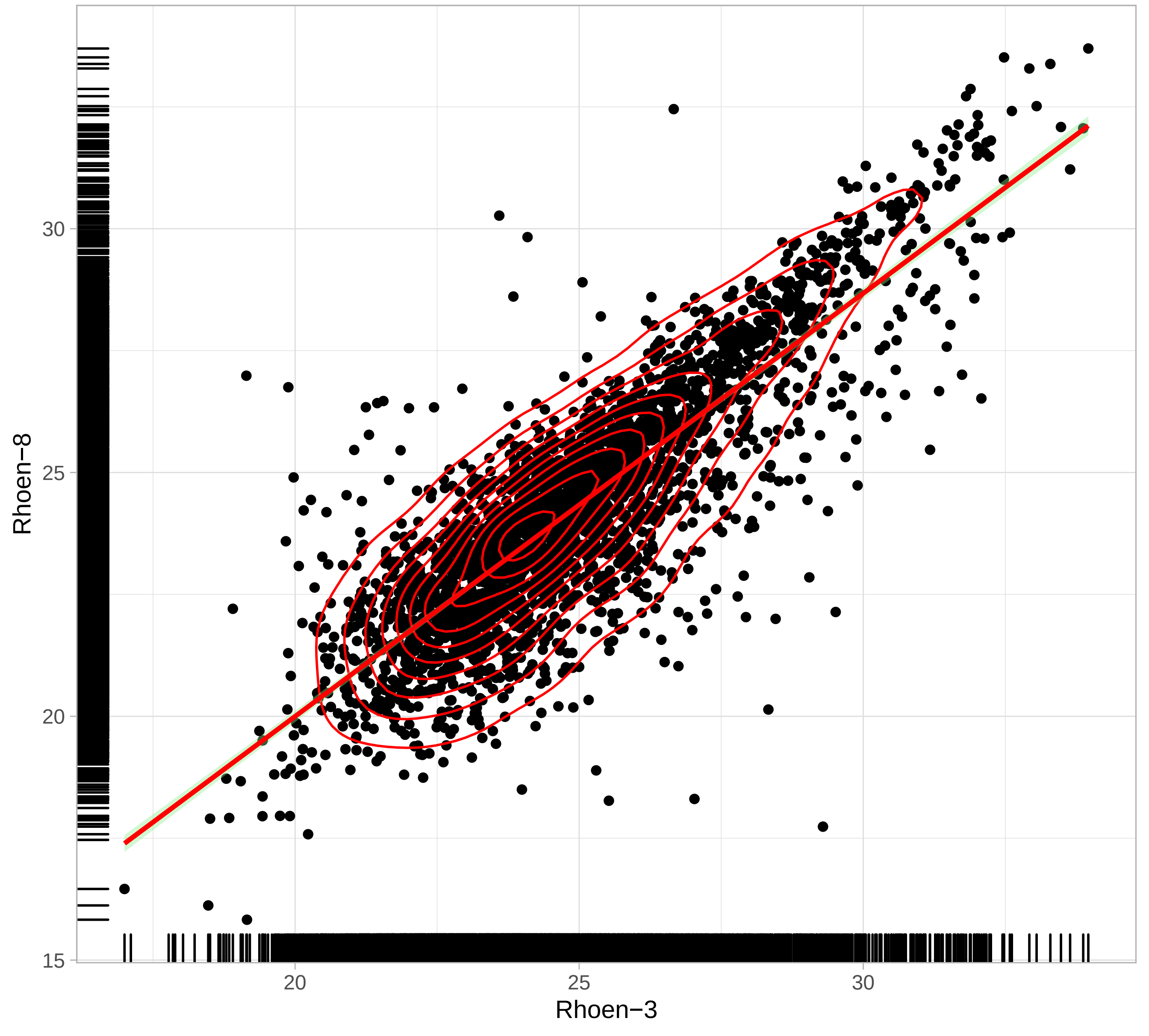

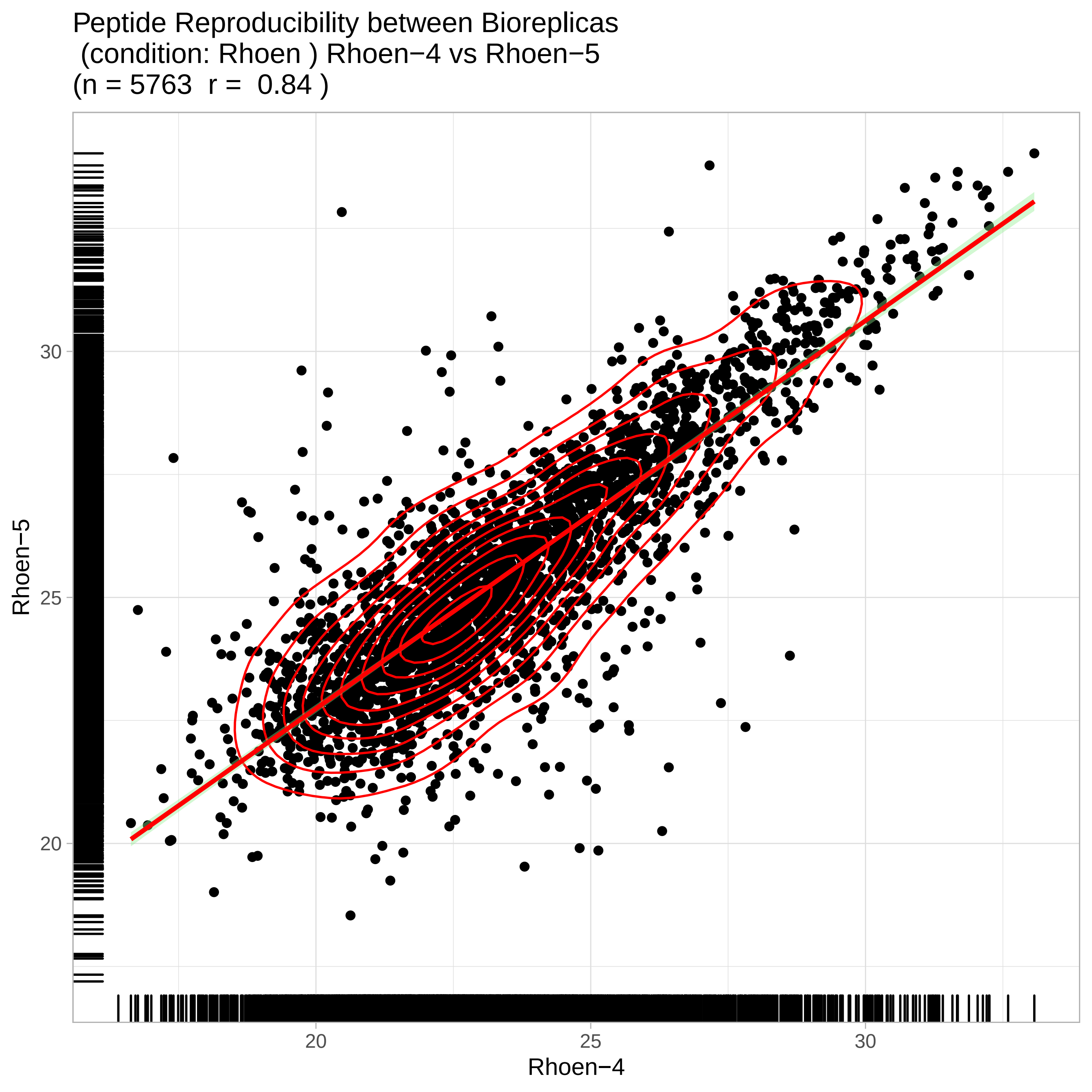
Peptide Reproducibility between Bioreplicas
(condition: Rhoen ) Rhoen−4 vs Rhoen−5
(n = 5763 r = 0.84 )
Rhoen−5
30
25
20
20
25
30
Rhoen−4

Peptide Reproducibility between Bioreplicas
(condition: Rhoen ) Rhoen−4 vs Rhoen−6
(n = 5763 r = 0.81 )

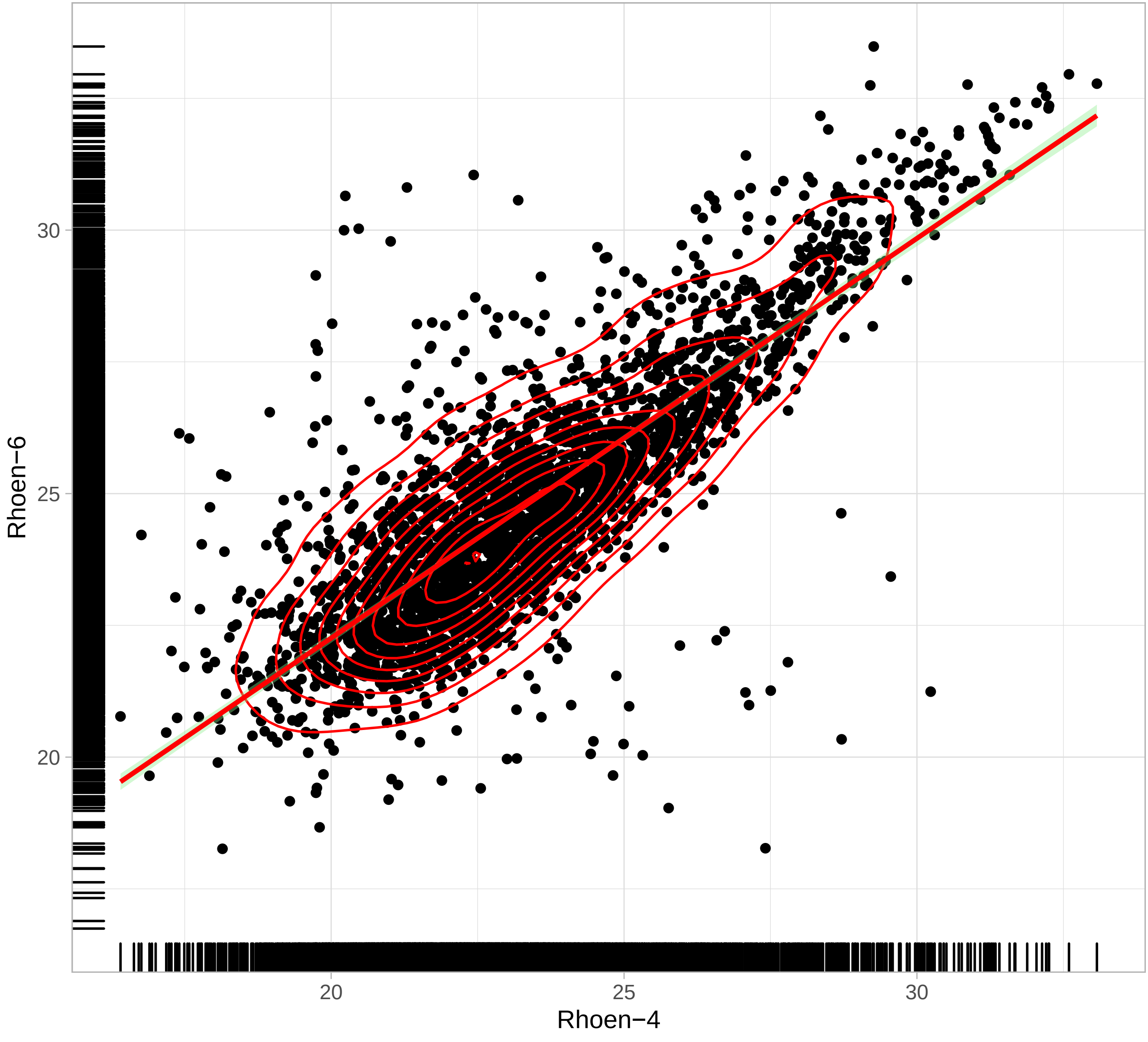

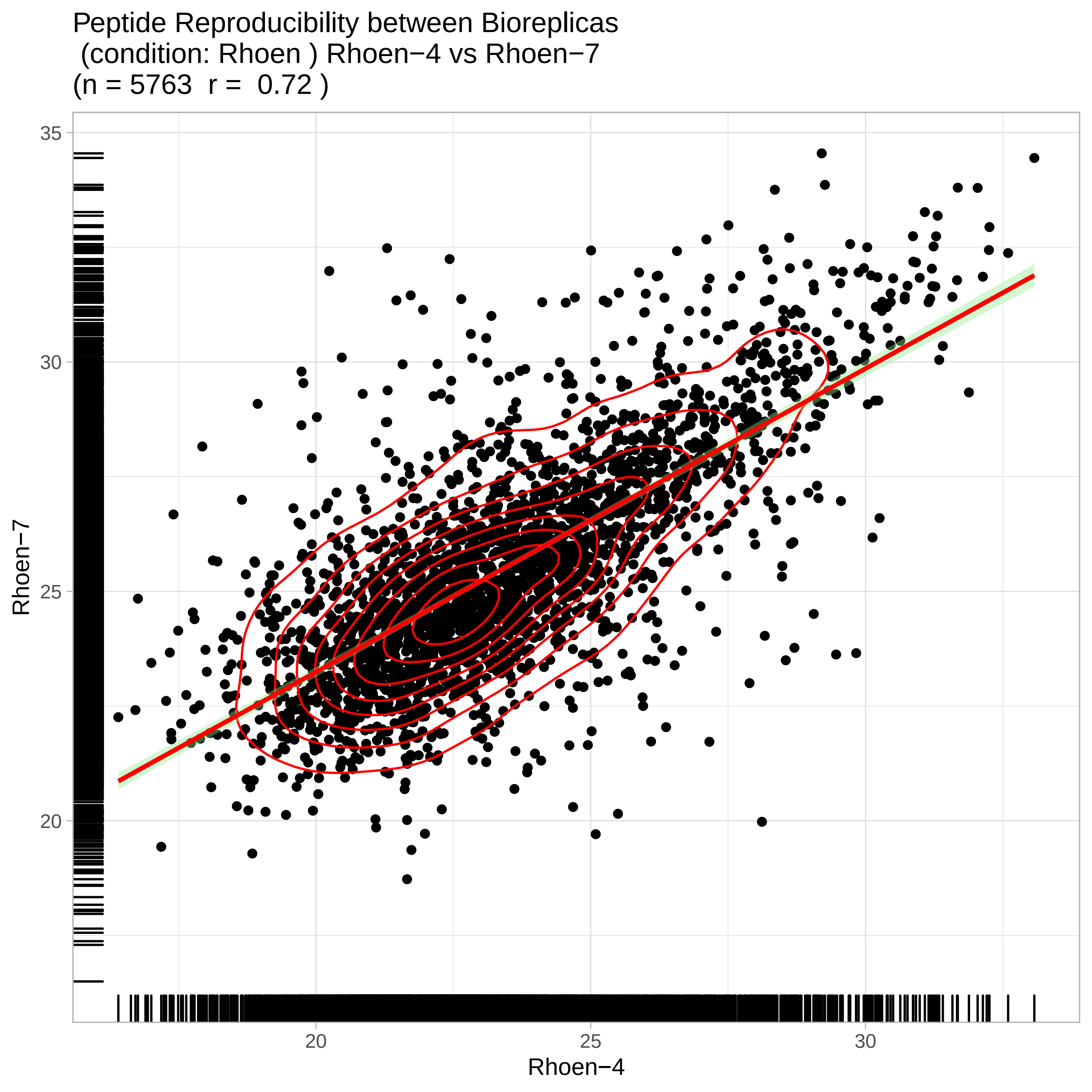
Peptide Reproducibility between Bioreplicas
(condition: Rhoen ) Rhoen−4 vs Rhoen−7
(n = 5763 r = 0.72 )
35
30
25
20
Rhoen−7
20
25
30
Rhoen−4

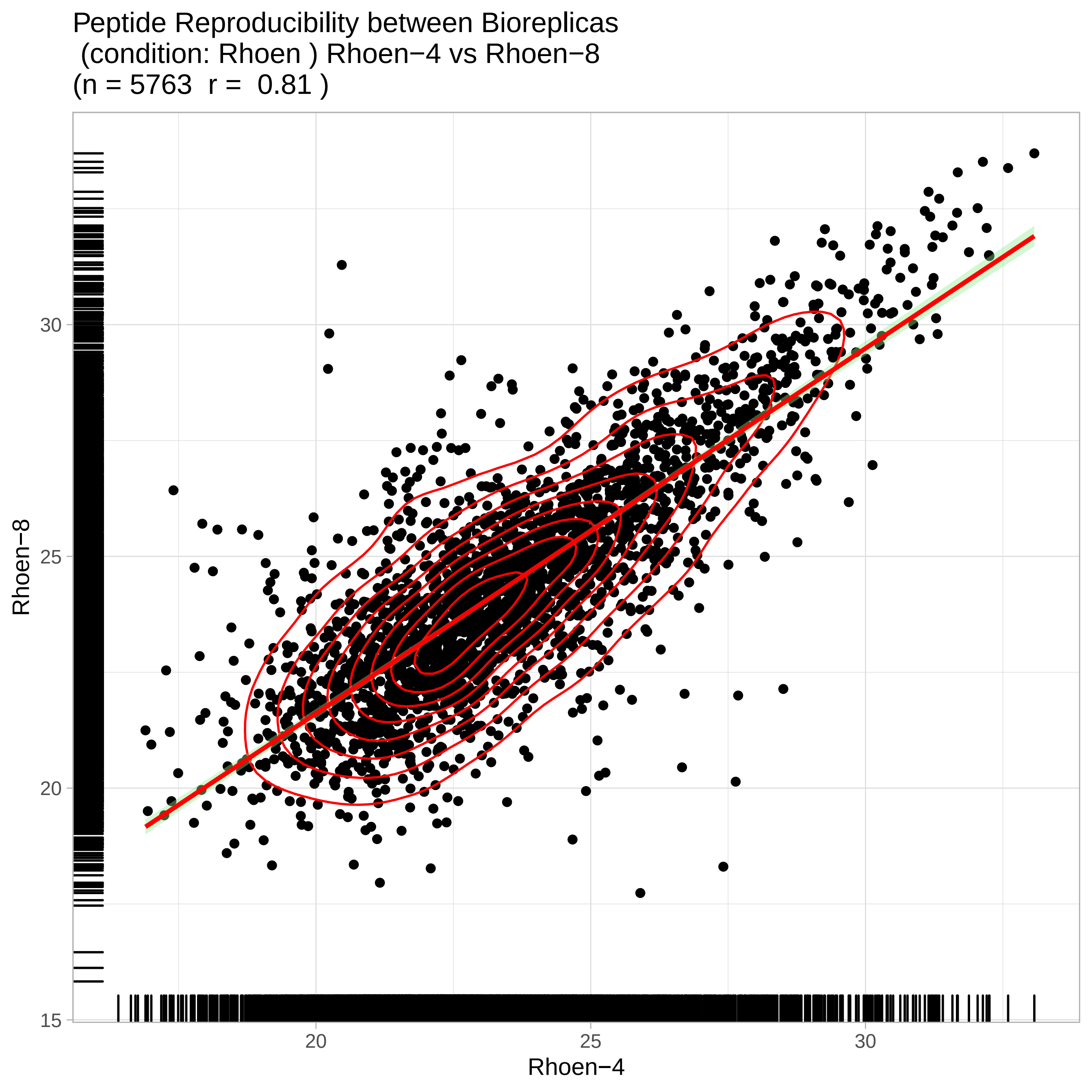

Peptide Reproducibility between Bioreplicas
(condition: Rhoen ) Rhoen−4 vs Rhoen−8
(n = 5763 r = 0.81 )
Rhoen−8
Rhoen−4
30
25
20
15
20
25
30

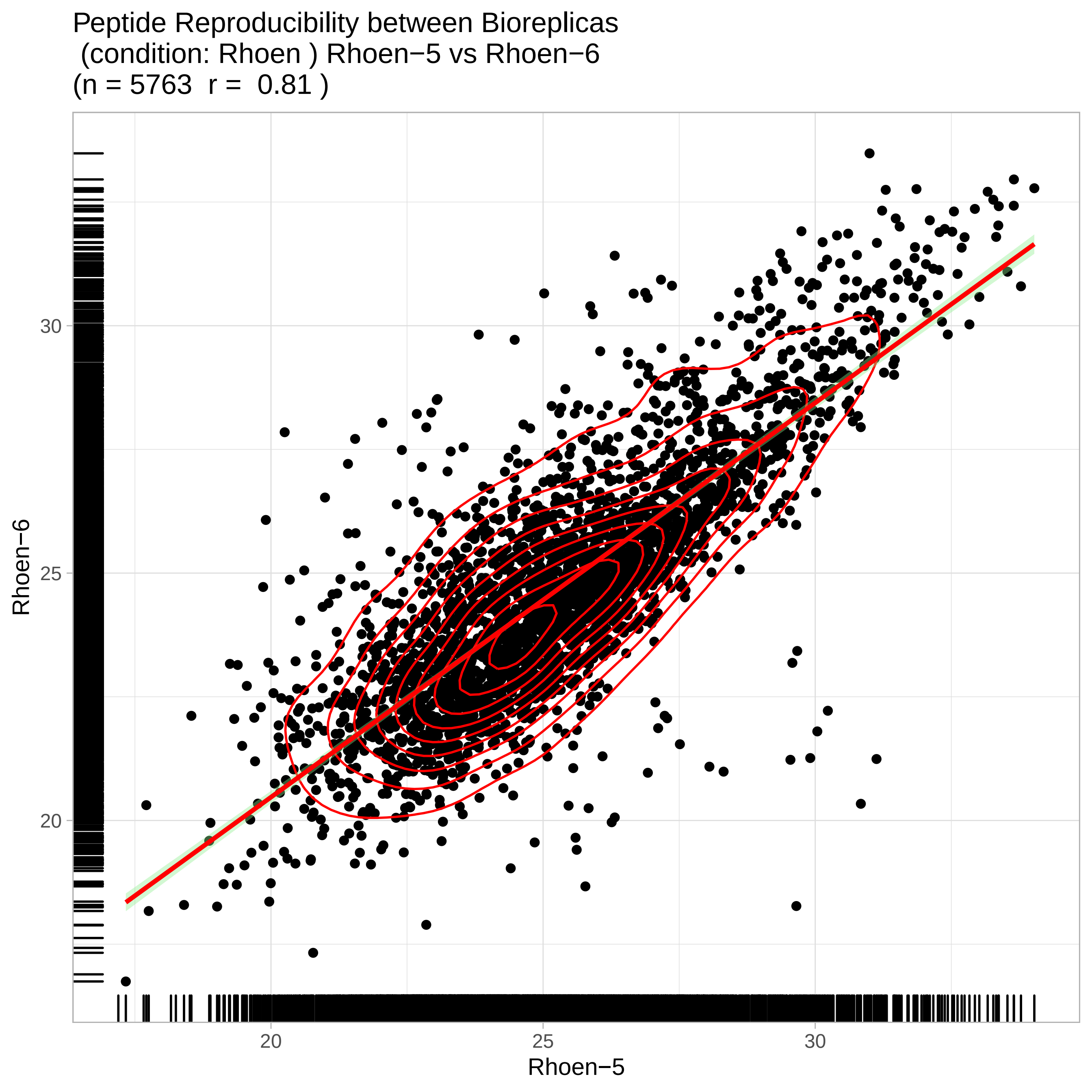

Peptide Reproducibility between Bioreplicas
(condition: Rhoen ) Rhoen−5 vs Rhoen−6
(n = 5763  r =  0.81 )
Rhoen−6
Rhoen−5
30
25
20
20
25
30

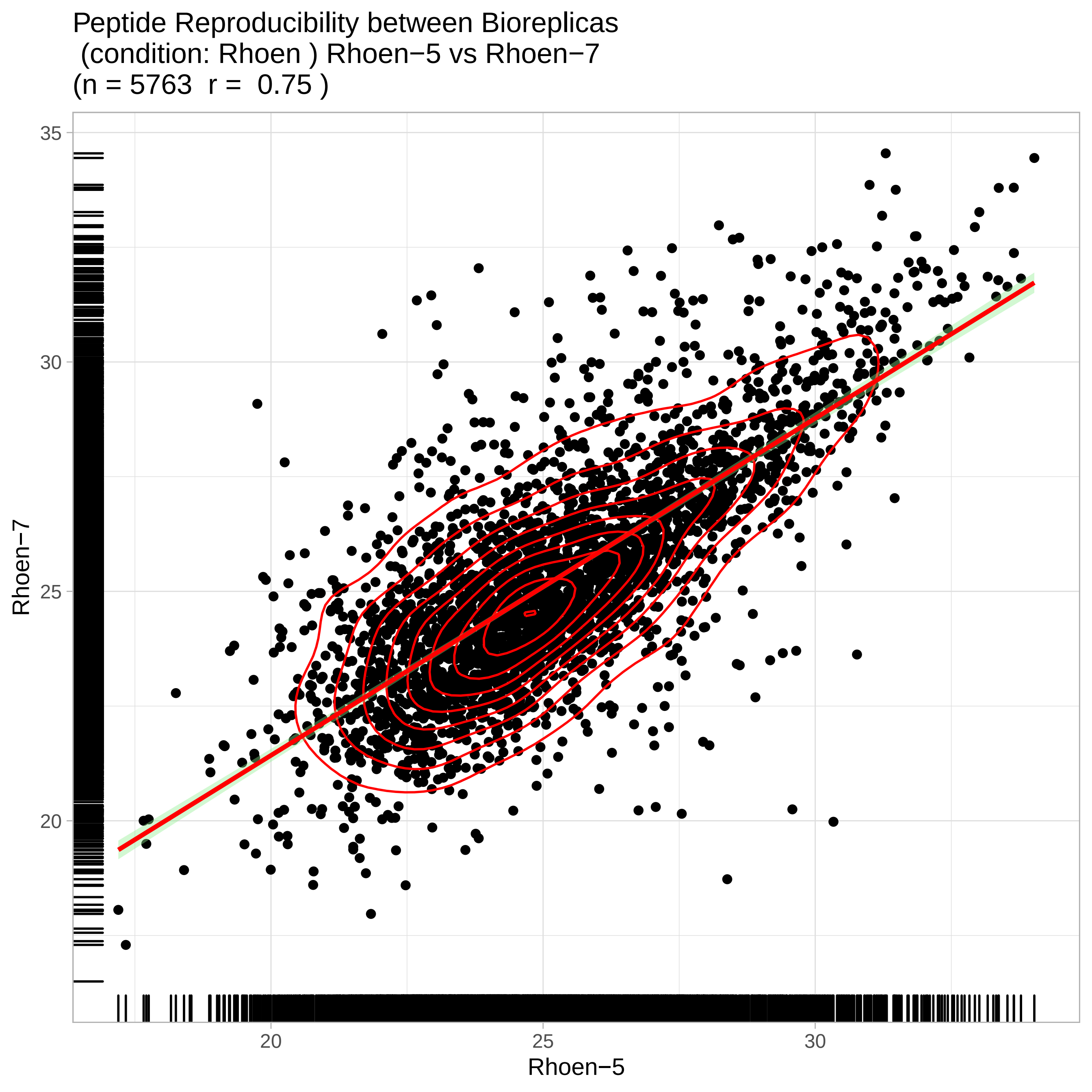
Peptide Reproducibility between Bioreplicas
(condition: Rhoen ) Rhoen−5 vs Rhoen−7
(n = 5763 r = 0.75 )
Rhoen−7
35
30
25
20
20
25
30
Rhoen−5

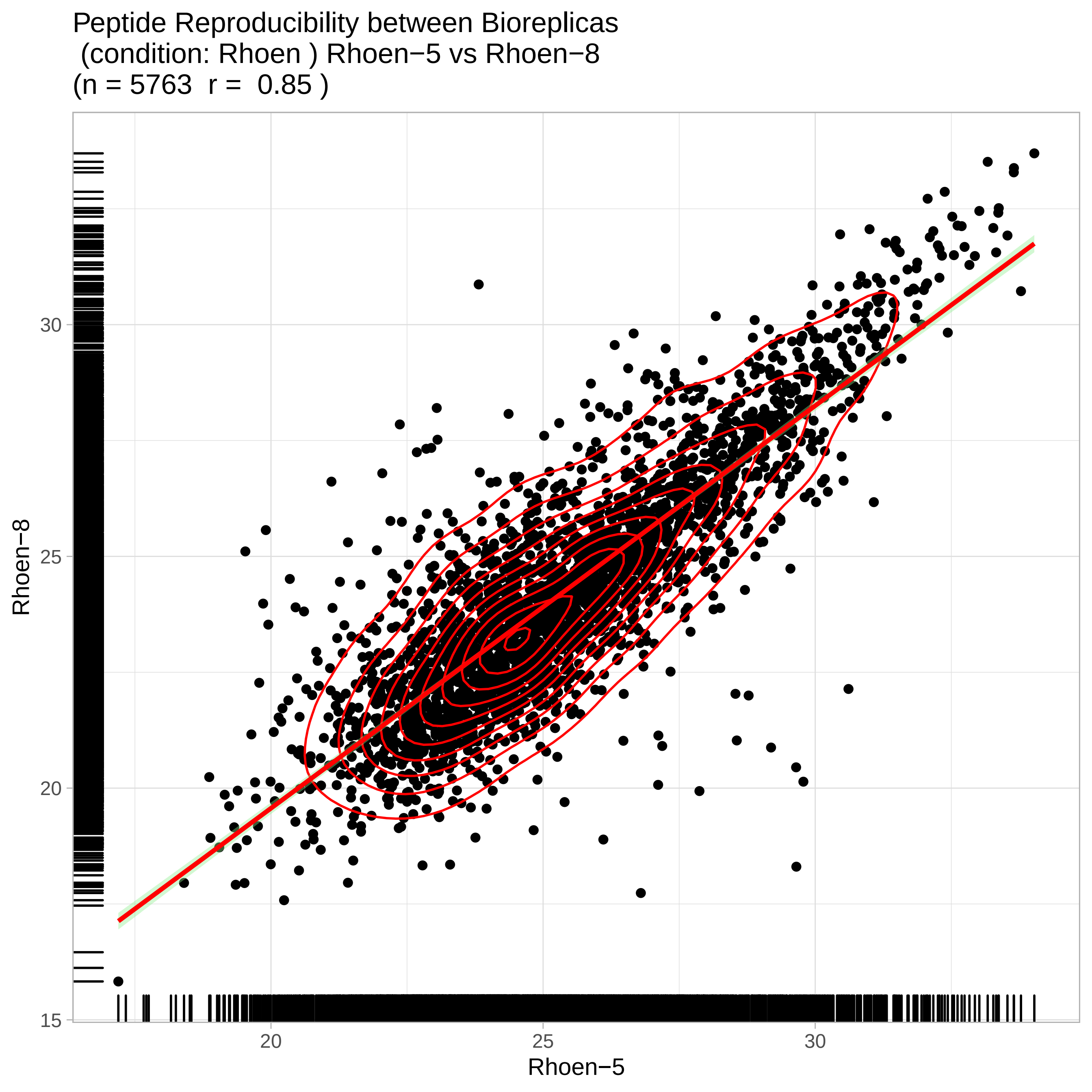
Peptide Reproducibility between Bioreplicas
(condition: Rhoen ) Rhoen−5 vs Rhoen−8
(n = 5763 r = 0.85 )
Rhoen−8
30
25
20
15
20
25
30
Rhoen−5

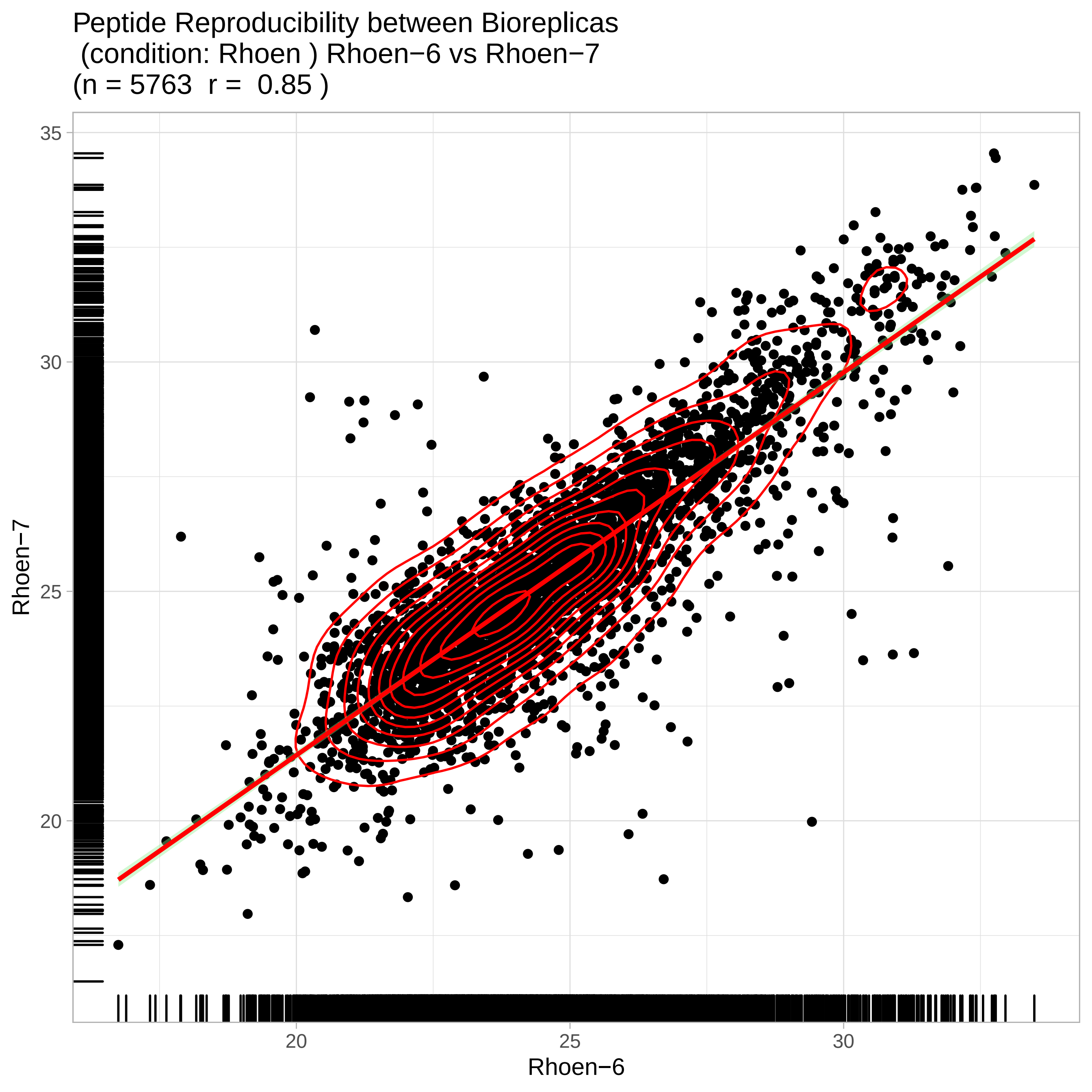
Peptide Reproducibility between Bioreplicas
(condition: Rhoen ) Rhoen−6 vs Rhoen−7
(n = 5763 r = 0.85 )
Rhoen−7
Rhoen−6
35
30
25
20
20
25
30

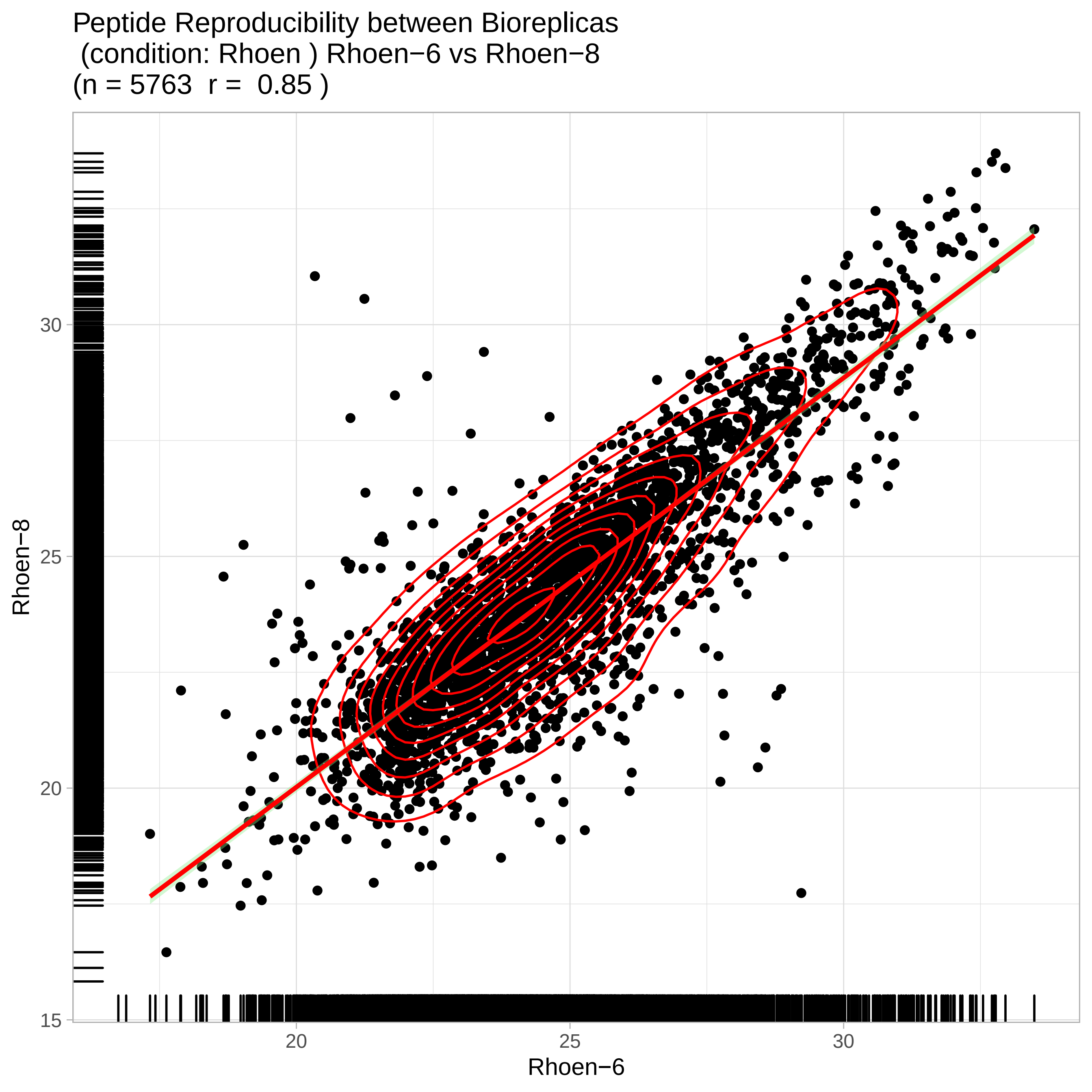
Peptide Reproducibility between Bioreplicas
(condition: Rhoen ) Rhoen−6 vs Rhoen−8
(n = 5763 r = 0.85 )
Rhoen−8
Rhoen−6
15
20
25
30
20
25
30

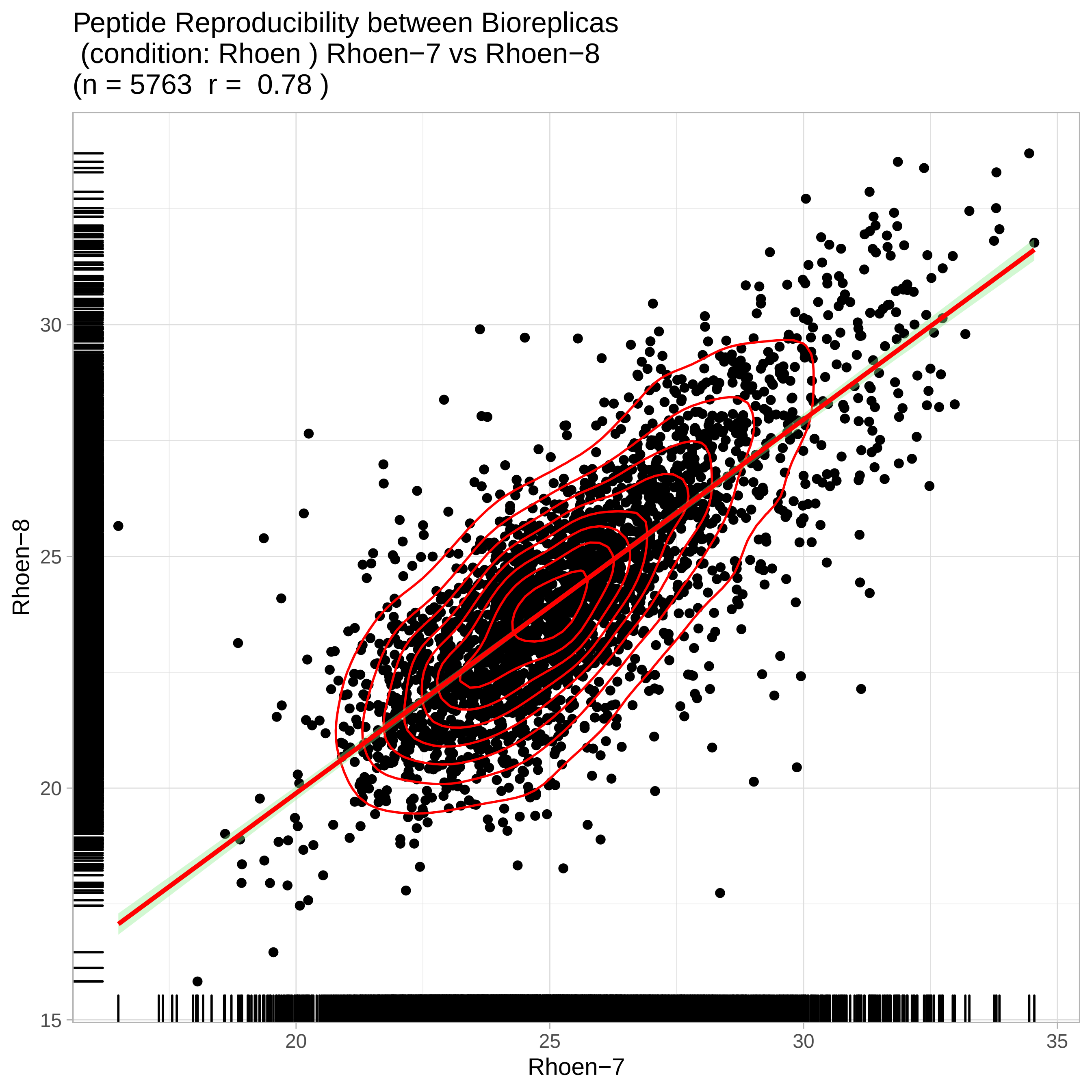
Peptide Reproducibility between Bioreplicas
(condition: Rhoen ) Rhoen−7 vs Rhoen−8
(n = 5763 r = 0.78 )
Rhoen−8
Rhoen−7
15
20
25
30
35

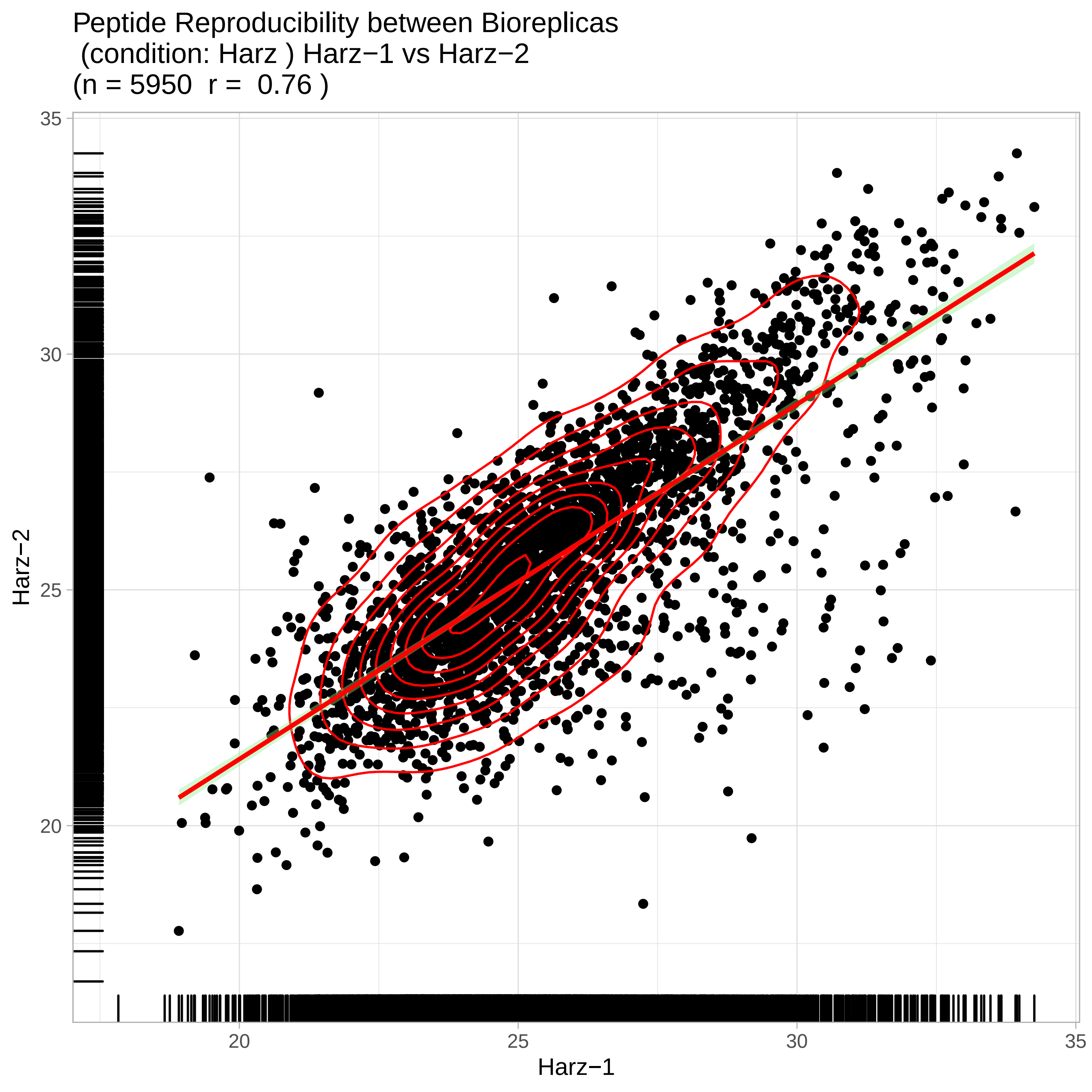
Peptide Reproducibility between Bioreplicas
(condition: Harz ) Harz−1 vs Harz−2
(n = 5950  r =  0.76 )
Harz−2
Harz−1
20
25
30
35
20
25
30
35

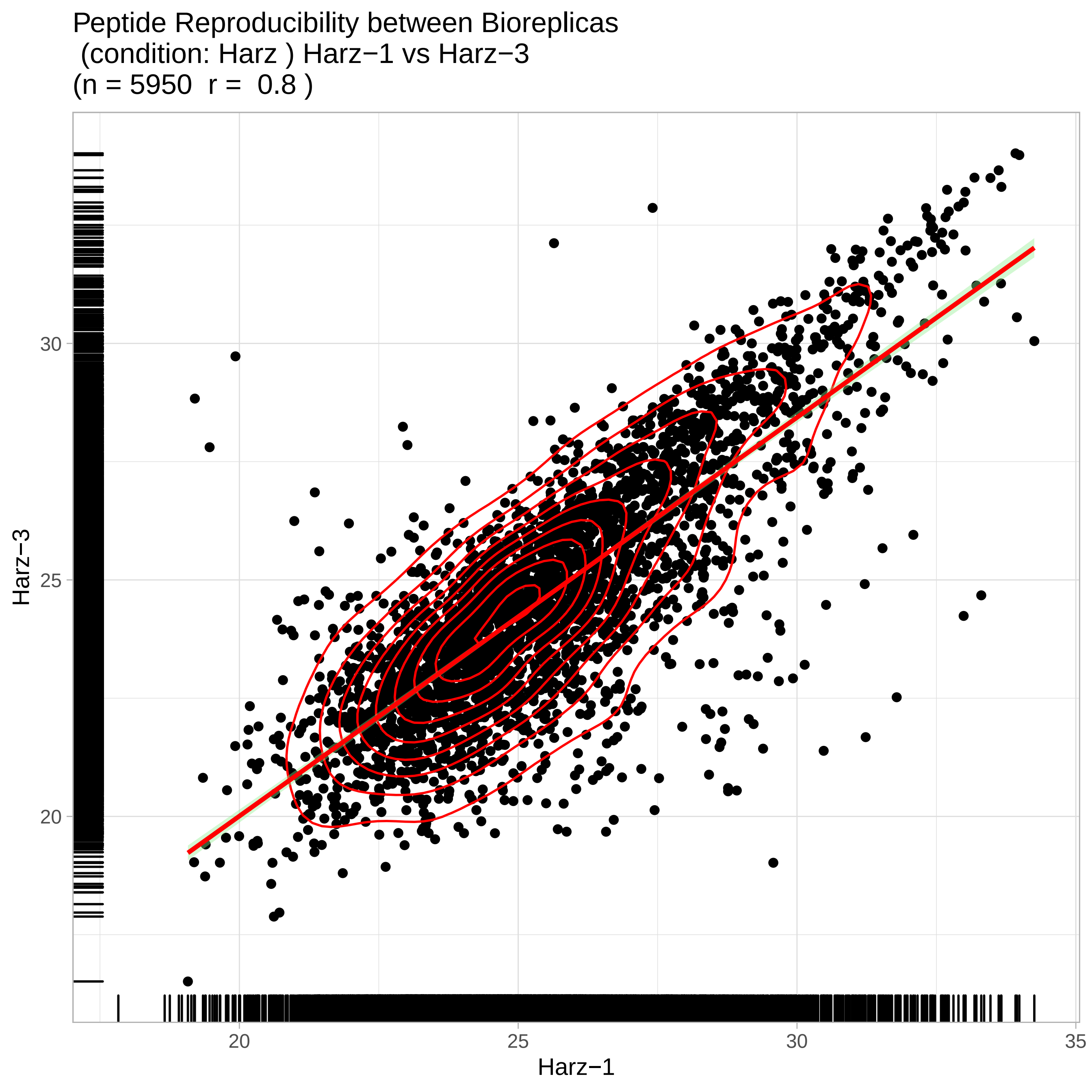
Peptide Reproducibility between Bioreplicas
(condition: Harz ) Harz−1 vs Harz−3
(n = 5950 r = 0.8 )
Harz−3
Harz−1
20
25
30
35

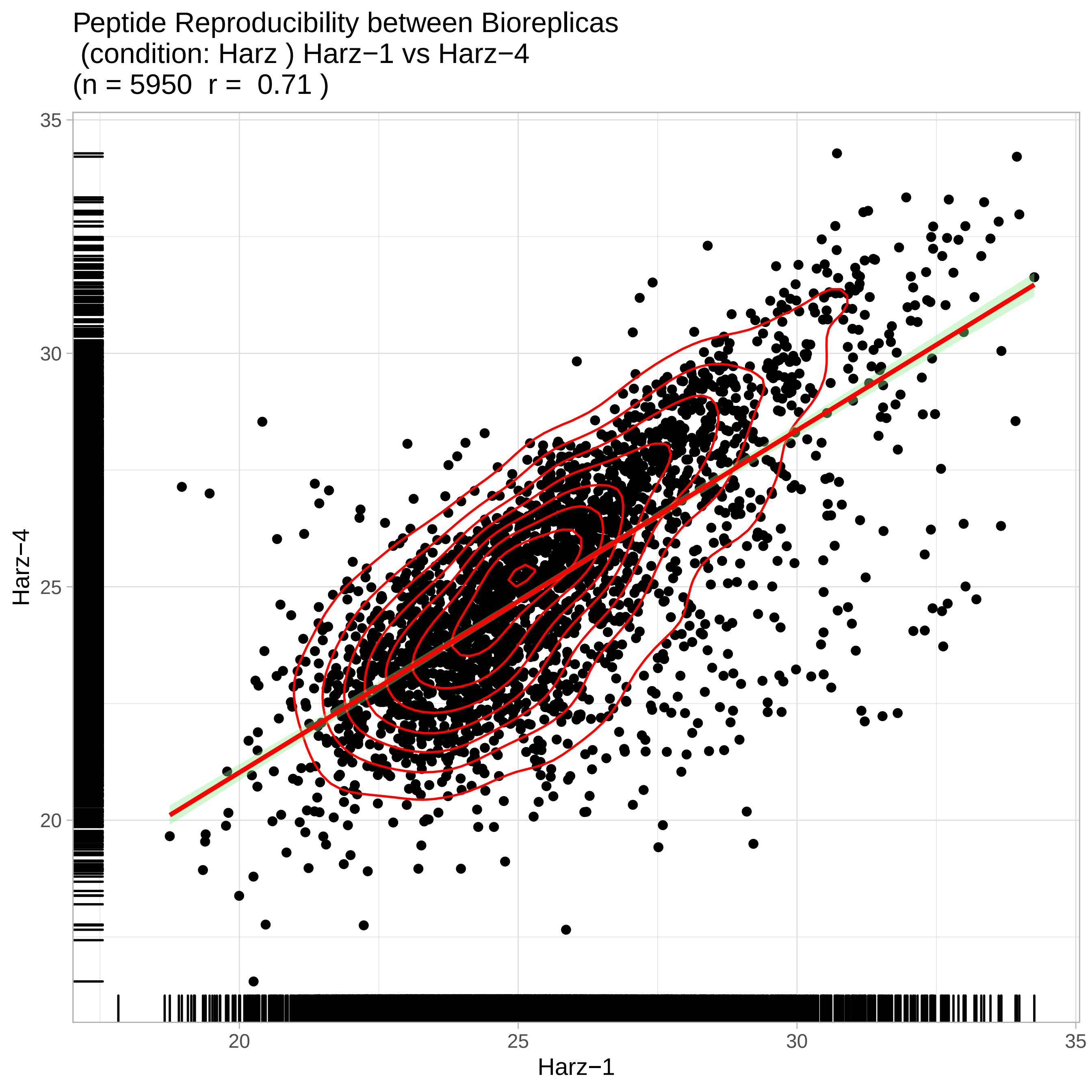
Peptide Reproducibility between Bioreplicas
(condition: Harz ) Harz−1 vs Harz−4
(n = 5950 r = 0.71 )
Harz−4
Harz−1
35
30
25
20
20
25
30
35

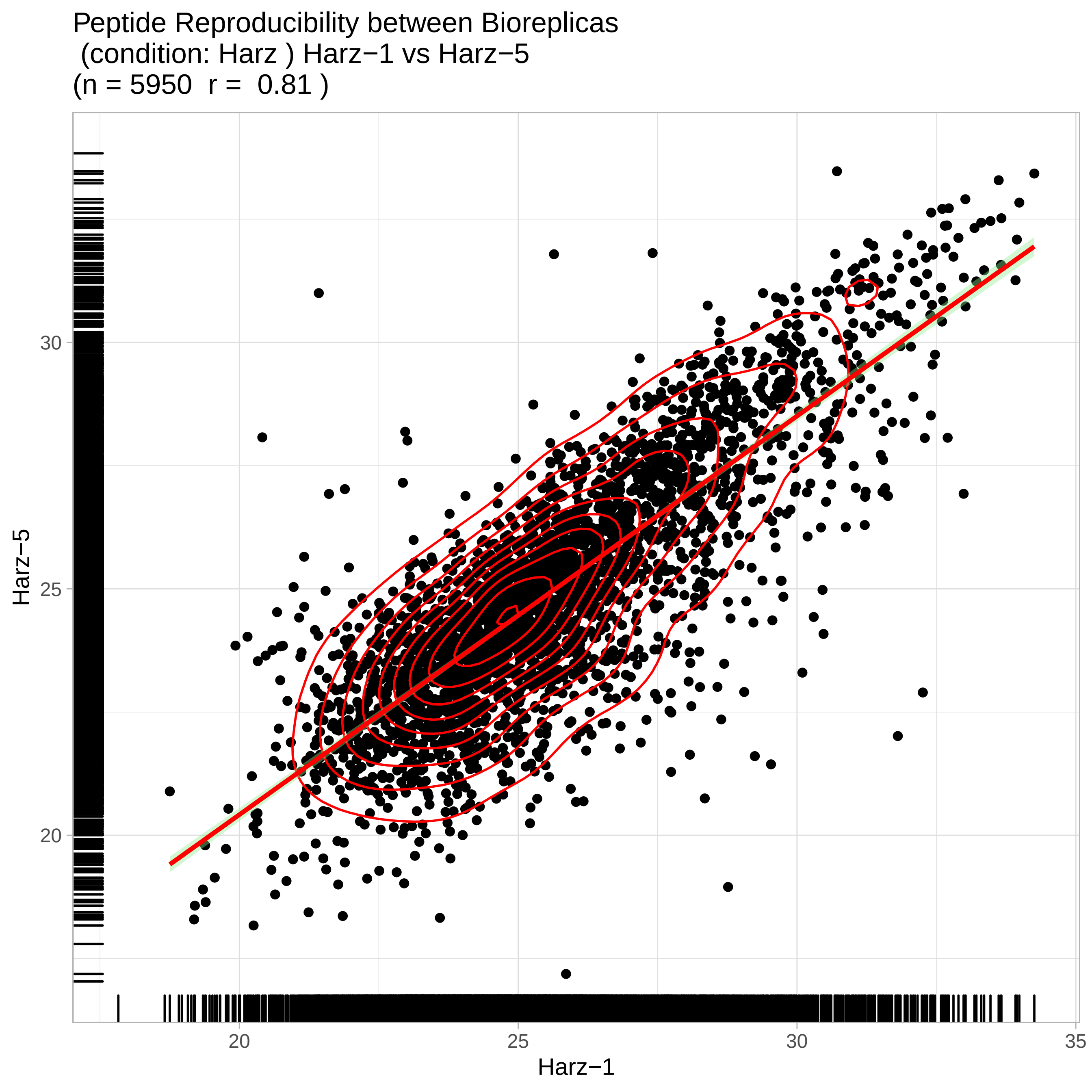
Peptide Reproducibility between Bioreplicas
(condition: Harz ) Harz−1 vs Harz−5
(n = 5950 r = 0.81 )
Harz−5
Harz−1
20
25
30
35

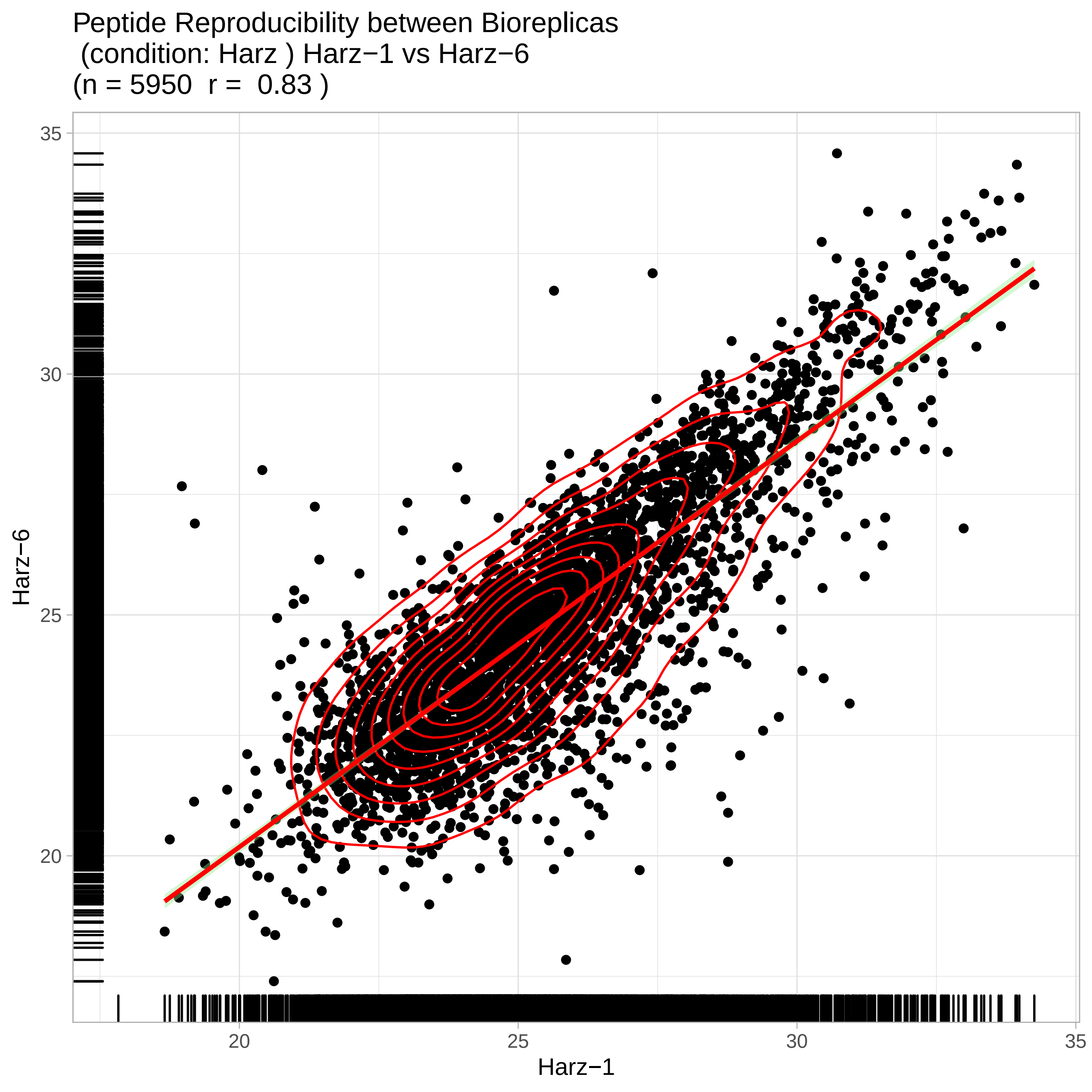

Peptide Reproducibility between Bioreplicas
(condition: Harz ) Harz−1 vs Harz−6
(n = 5950 r = 0.83 )
Harz−6
Harz−1
35
30
25
20
20
25
30
35

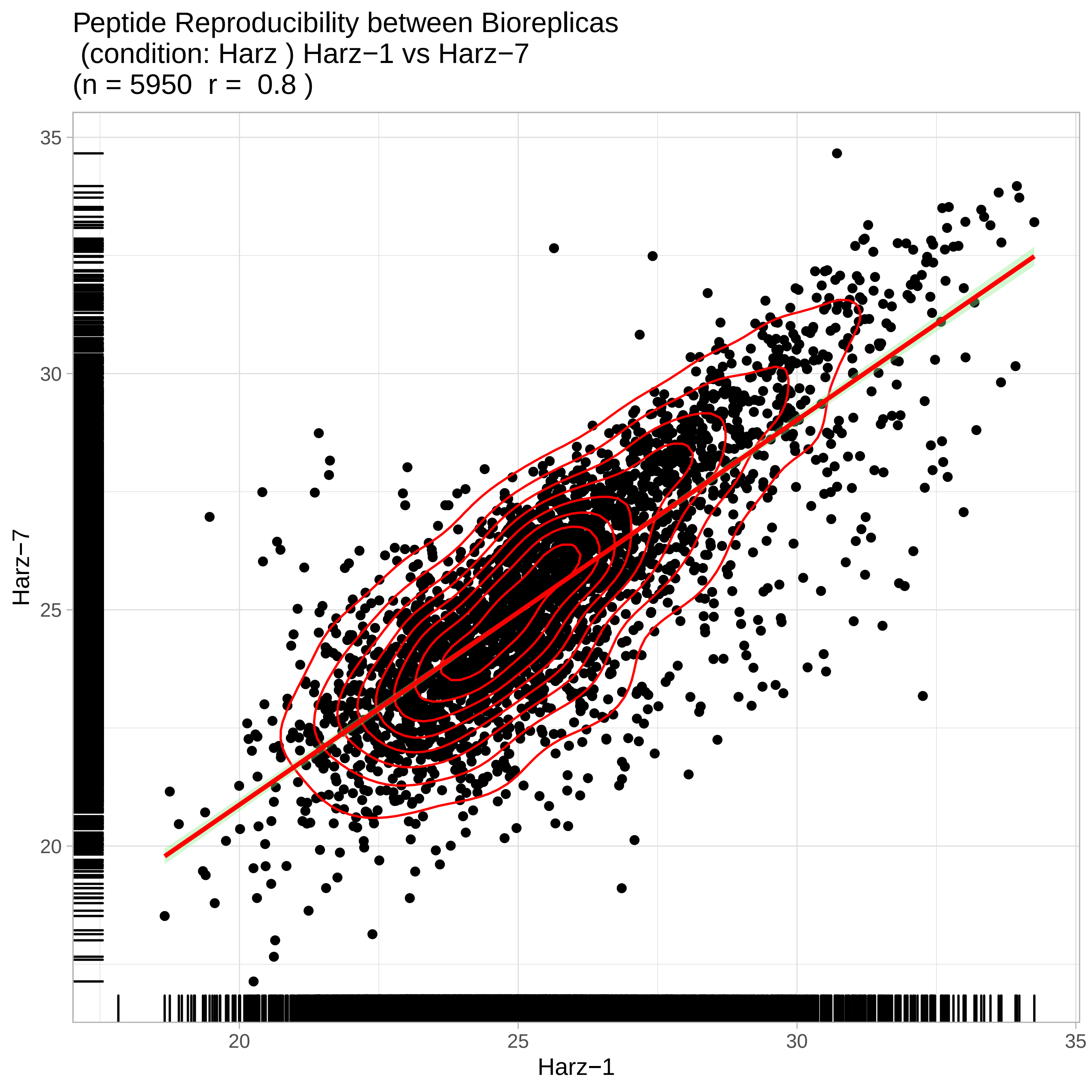
Peptide Reproducibility between Bioreplicas
(condition: Harz ) Harz−1 vs Harz−7
(n = 5950 r = 0.8 )
Harz−7
Harz−1
35
30
25
20
20
25
30
35

Peptide Reproducibility between Bioreplicas
(condition: Harz ) Harz−1 vs Harz−8
(n = 5950  r =  0.84 )

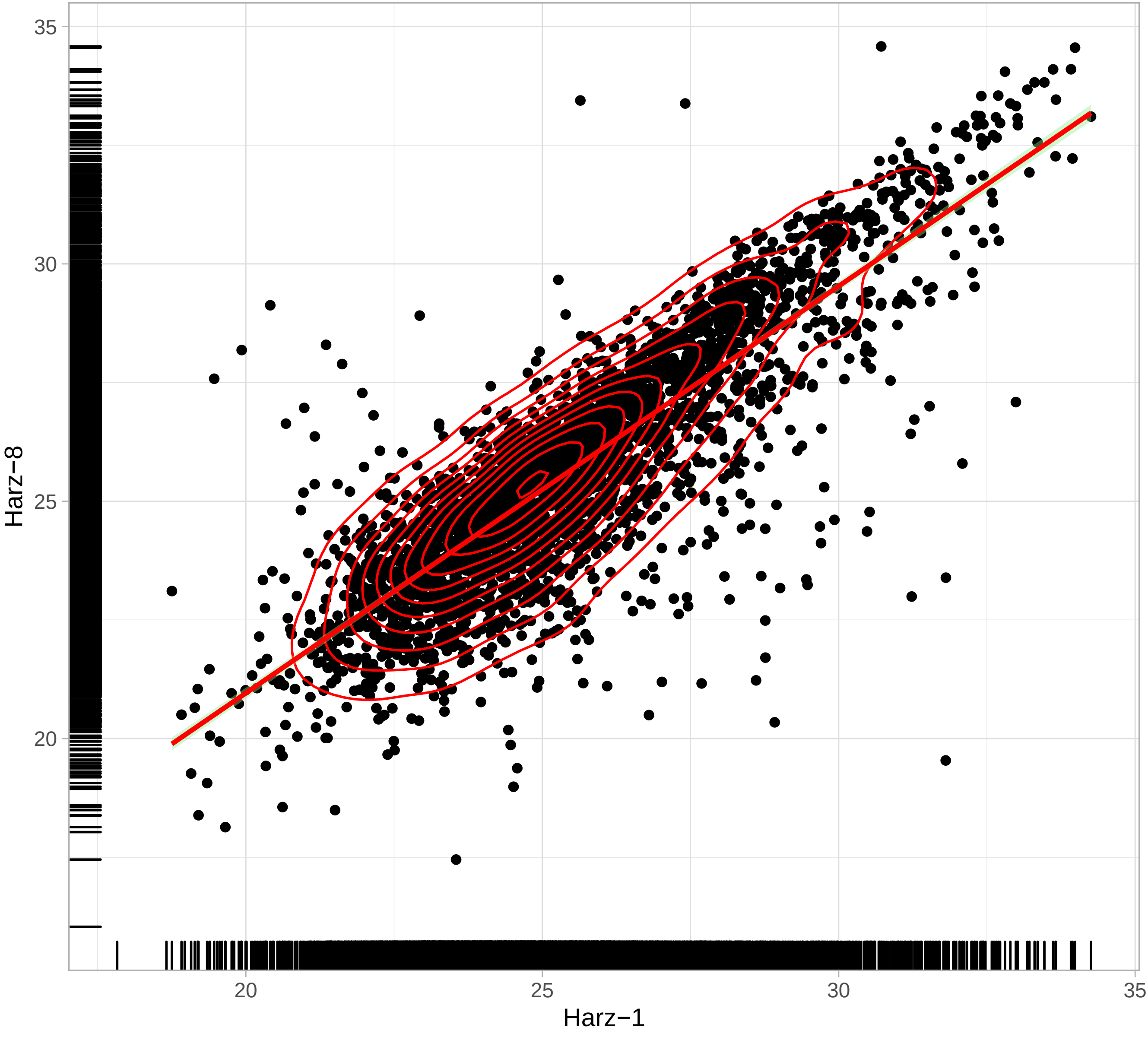

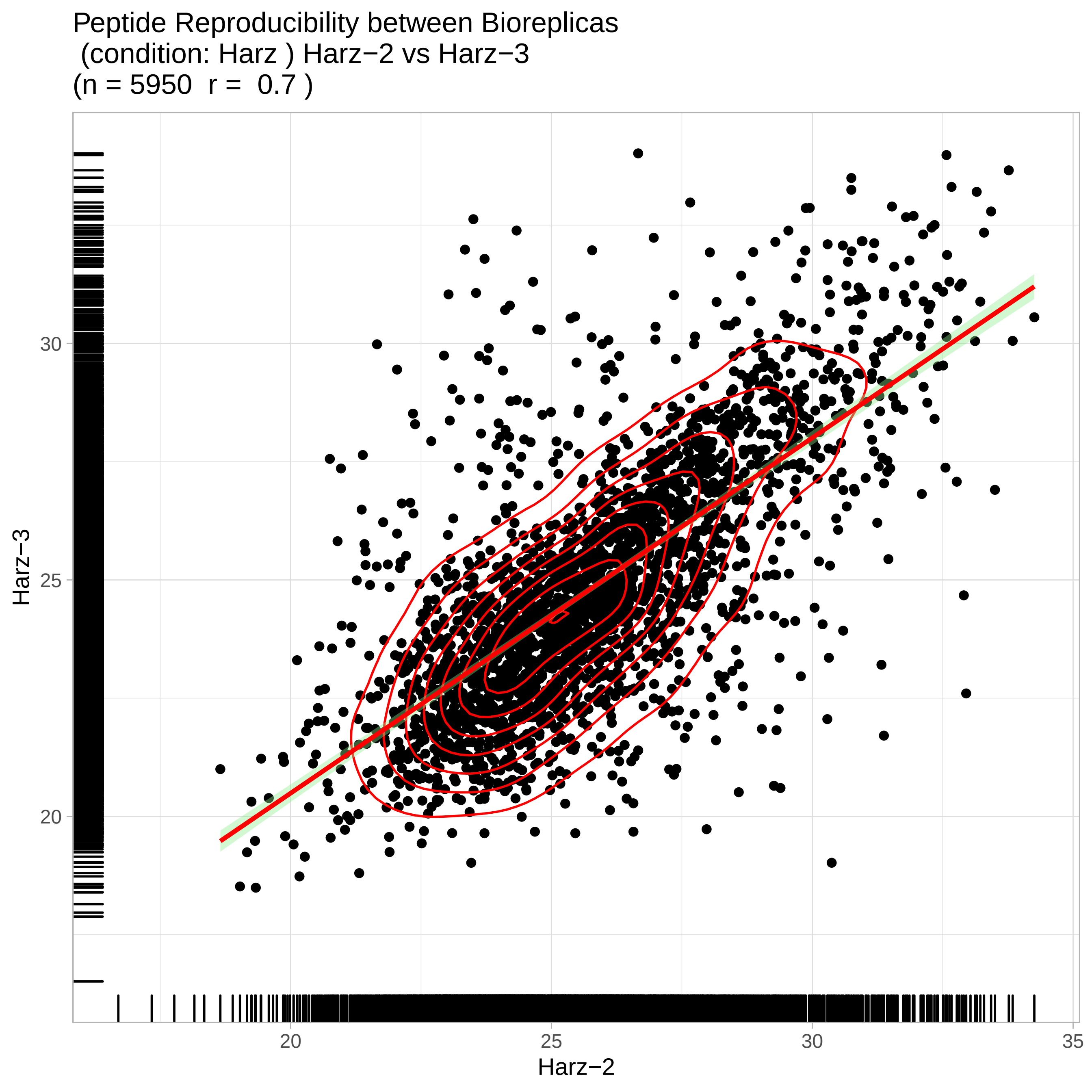
Peptide Reproducibility between Bioreplicas
(condition: Harz ) Harz−2 vs Harz−3
(n = 5950 r = 0.7 )
Harz−3
30
25
20
Harz−2
20
25
30
35

Peptide Reproducibility between Bioreplicas
(condition: Harz ) Harz−2 vs Harz−4
(n = 5950 r = 0.85 )

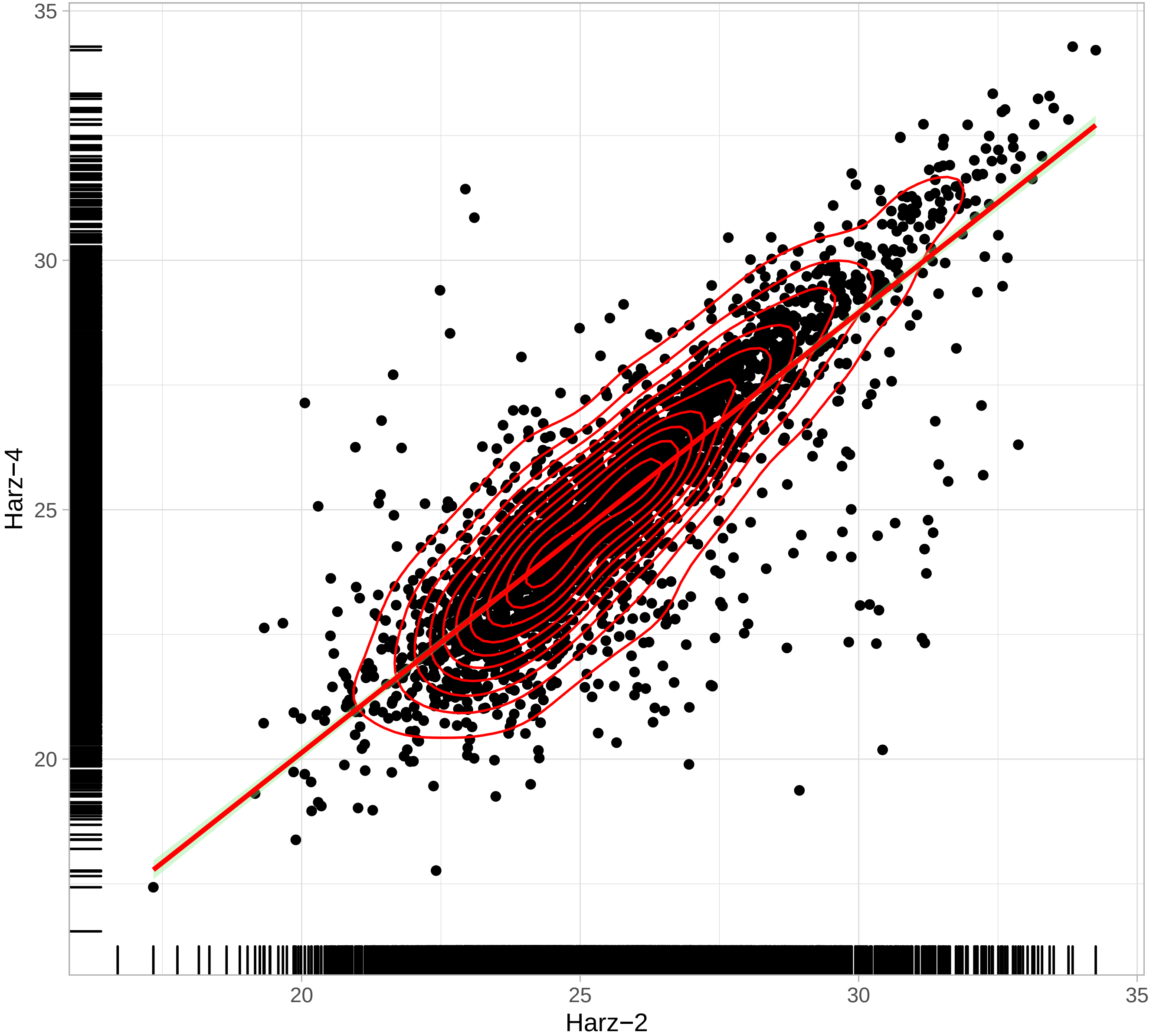

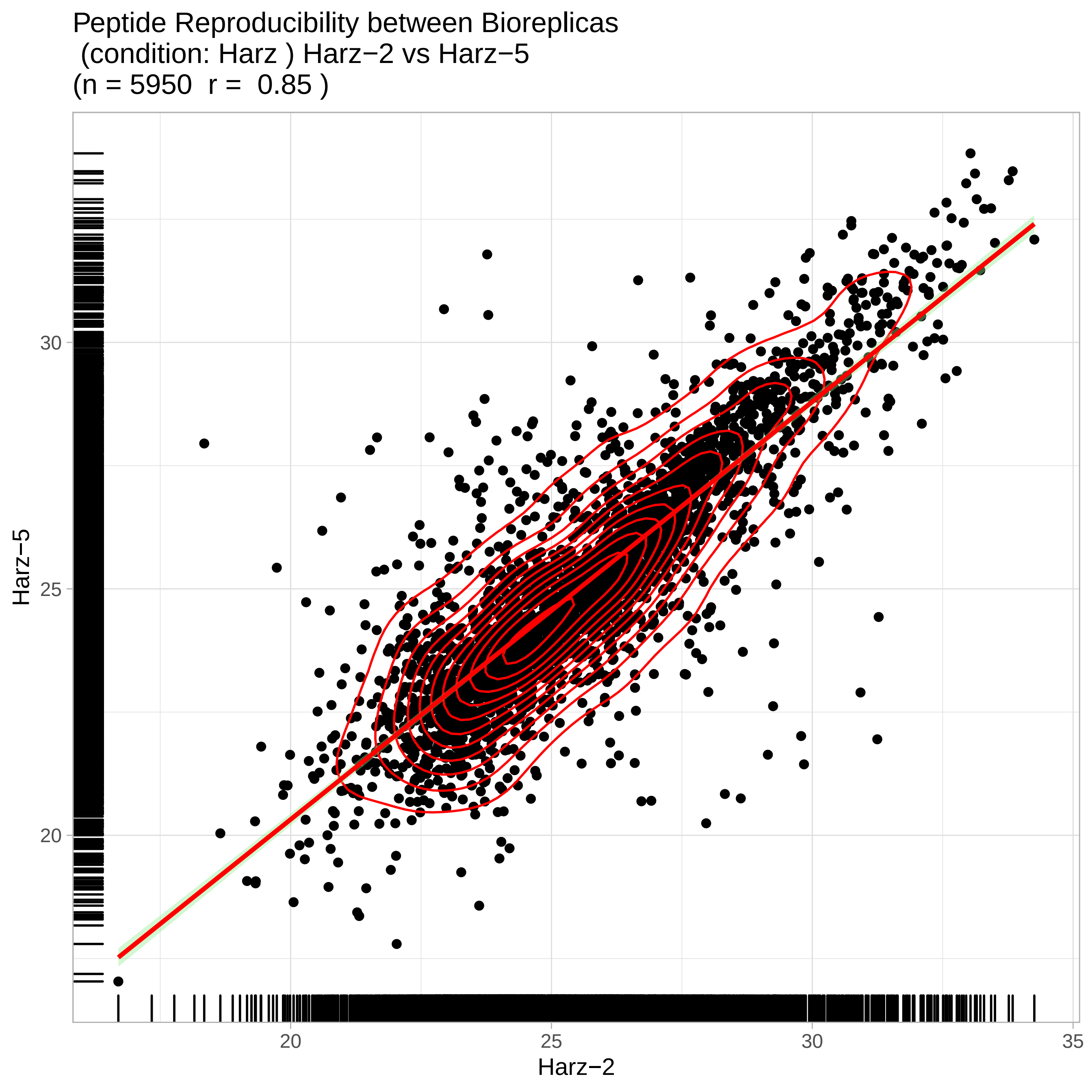

Peptide Reproducibility between Bioreplicas
(condition: Harz ) Harz−2 vs Harz−5
(n = 5950 r = 0.85 )
Harz−5
30
25
20
Harz−2
20
25
30
35

Peptide Reproducibility between Bioreplicas
(condition: Harz ) Harz−2 vs Harz−6
(n = 5950  r =  0.85 )

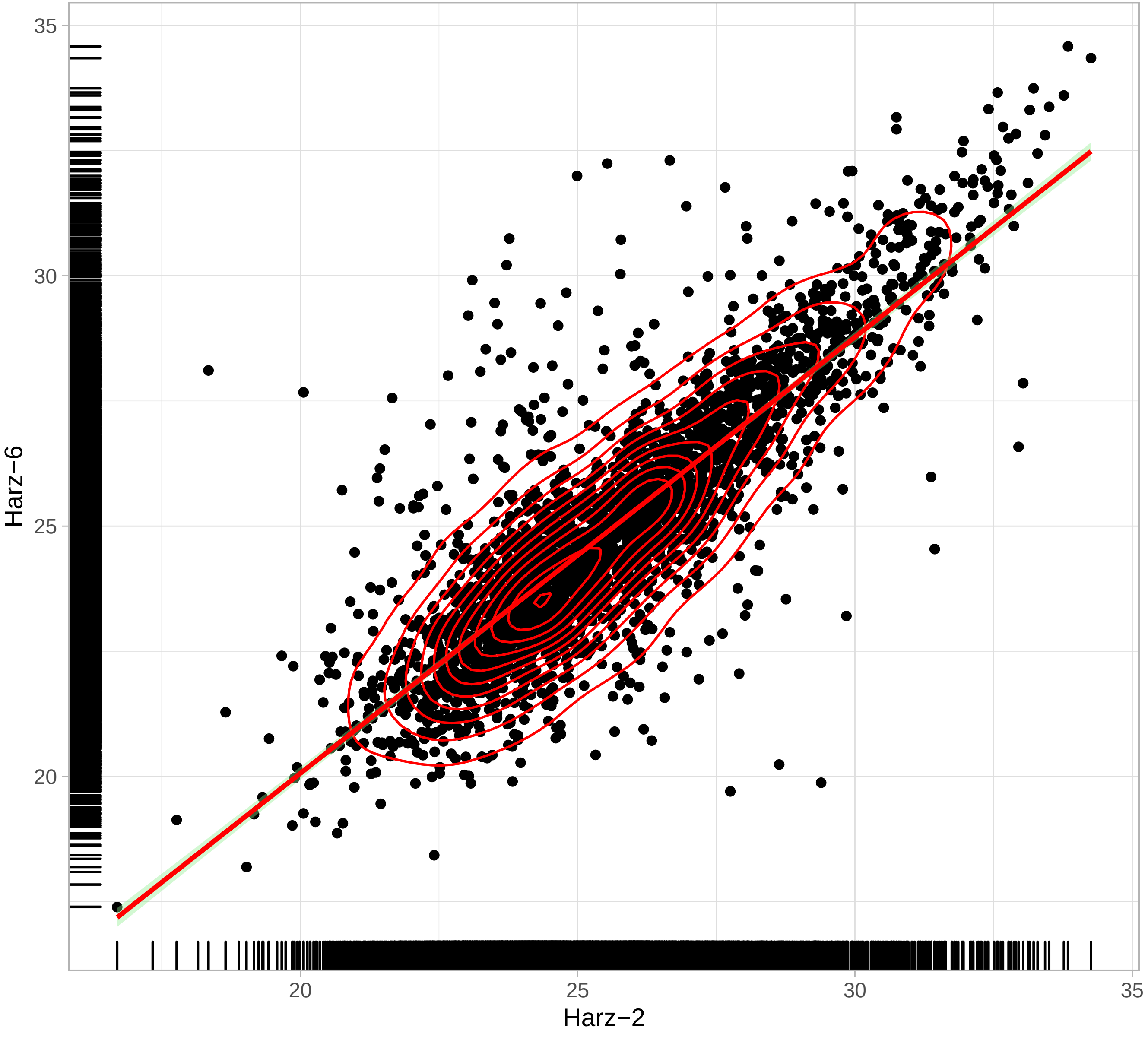

Peptide Reproducibility between Bioreplicas
(condition: Harz ) Harz−2 vs Harz−7
(n = 5950 r = 0.87 )

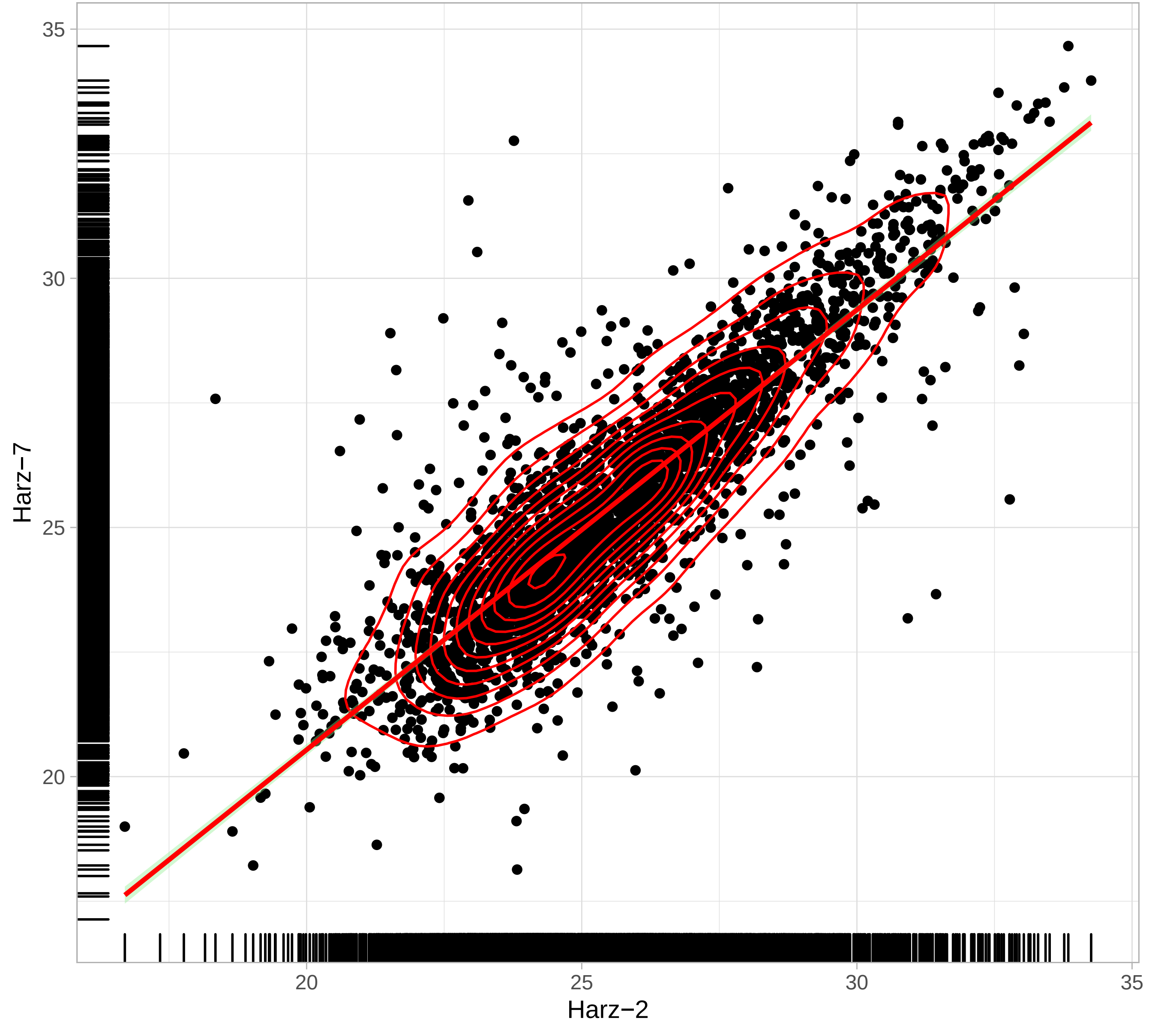

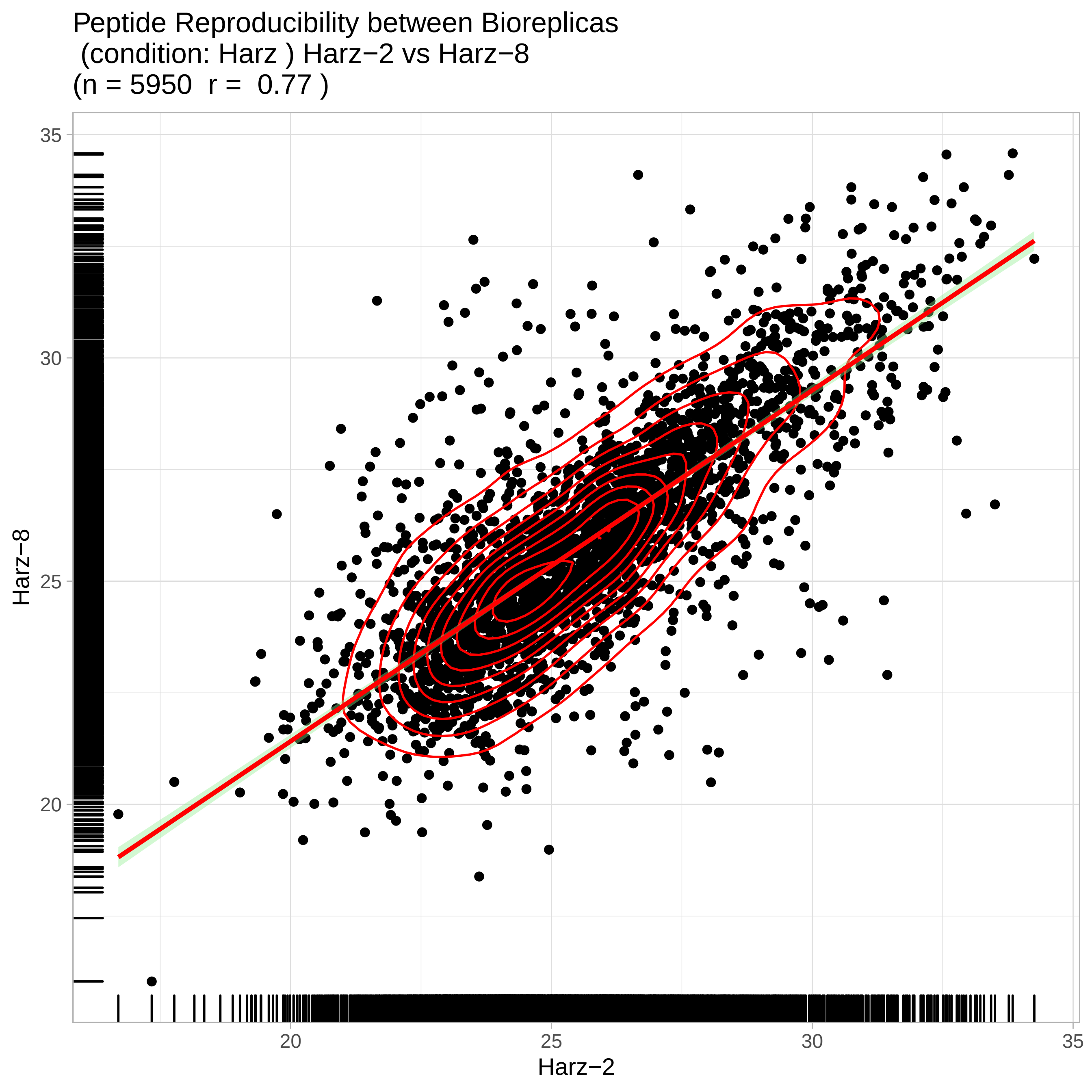
Peptide Reproducibility between Bioreplicas
(condition: Harz ) Harz−2 vs Harz−8
(n = 5950 r = 0.77 )
Harz−8
Harz−2
35
30
25
20
20
25
30
35

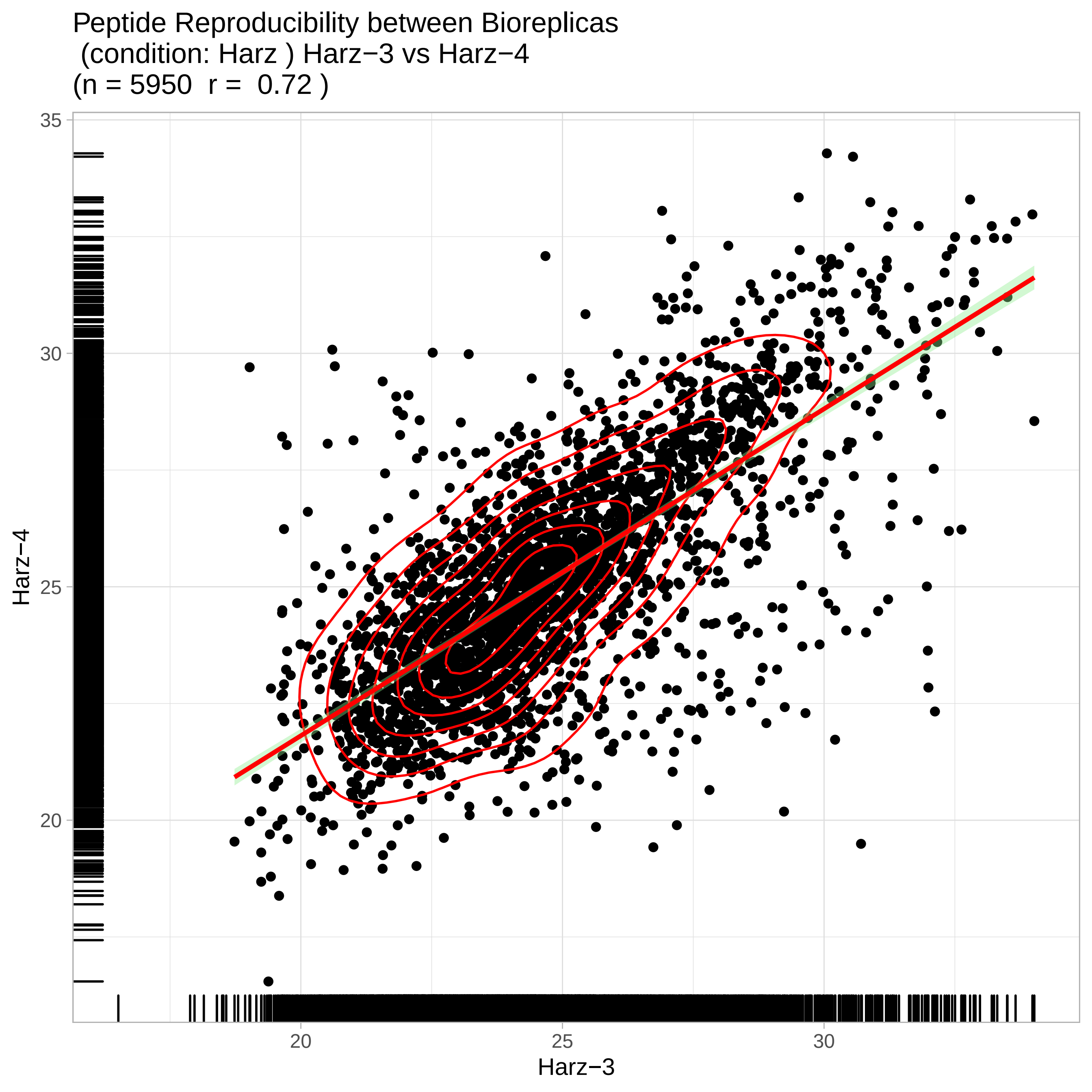

Peptide Reproducibility between Bioreplicas
(condition: Harz ) Harz−3 vs Harz−4
(n = 5950 r = 0.72 )
35
30
25
20
Harz−4
20
25
30
Harz−3

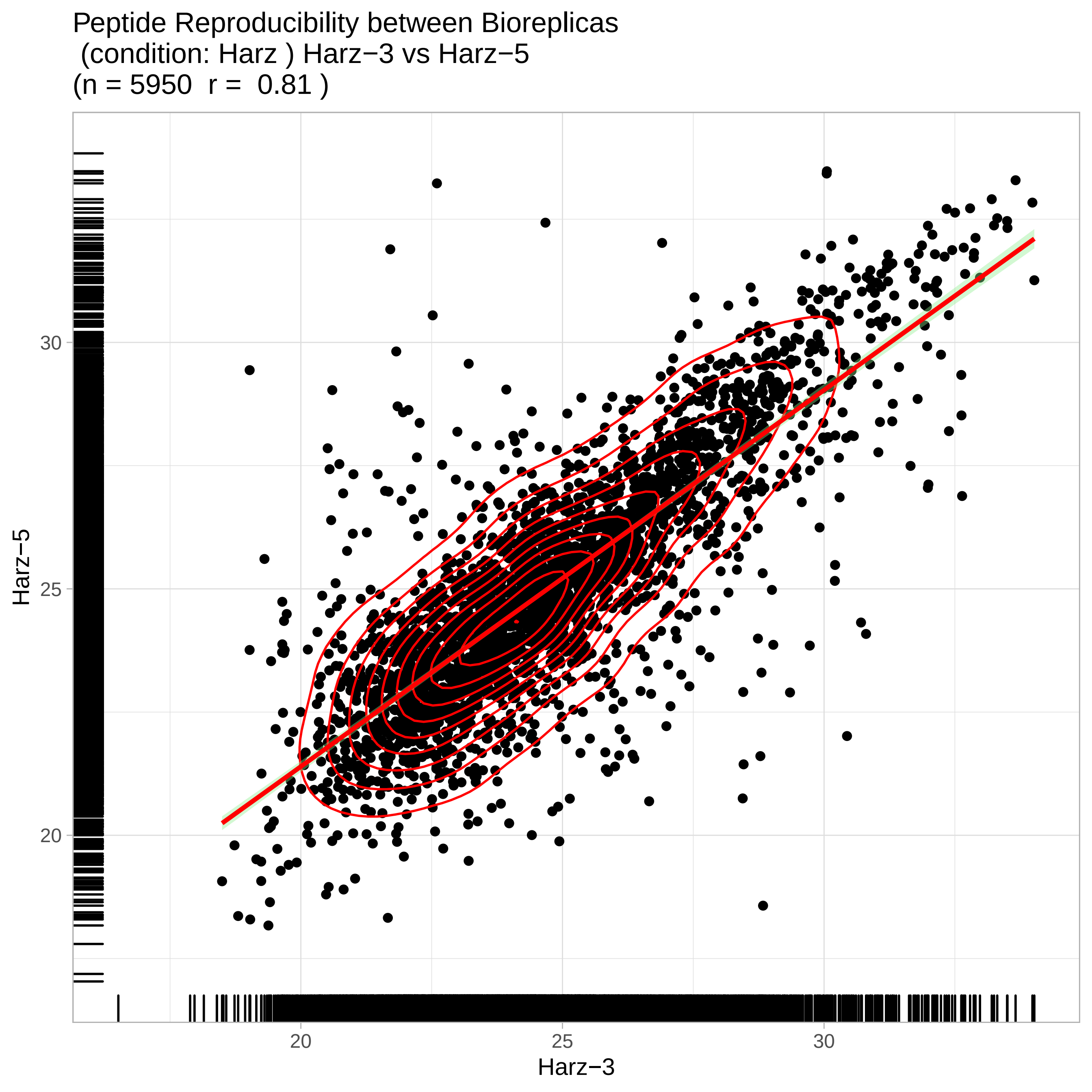
Peptide Reproducibility between Bioreplicas
(condition: Harz ) Harz−3 vs Harz−5
(n = 5950 r = 0.81 )
Harz−5
Harz−3
20
25
30
20
25
30

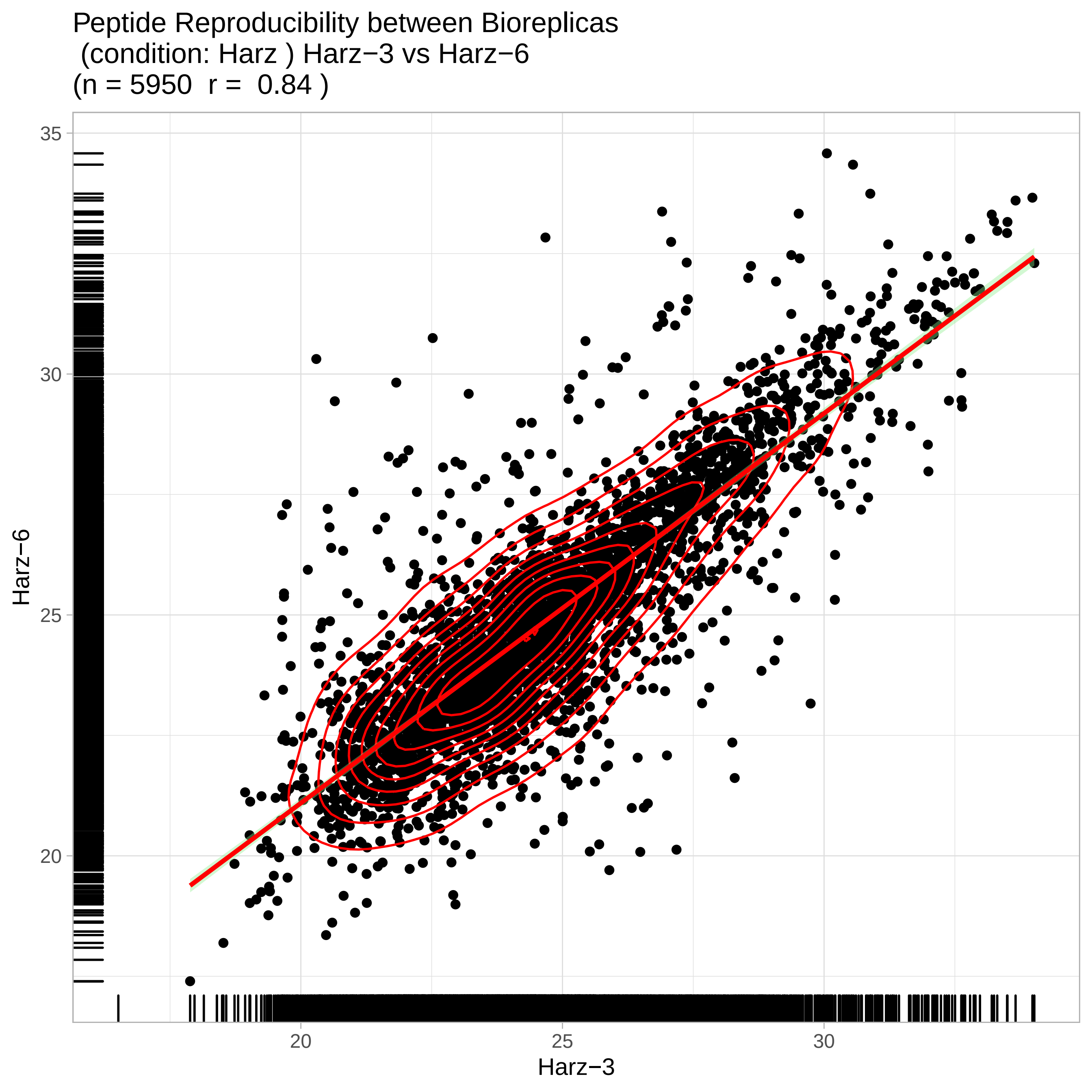
Peptide Reproducibility between Bioreplicas
(condition: Harz ) Harz−3 vs Harz−6
(n = 5950 r = 0.84 )
35
30
25
20
Harz−6
20
25
30
Harz−3

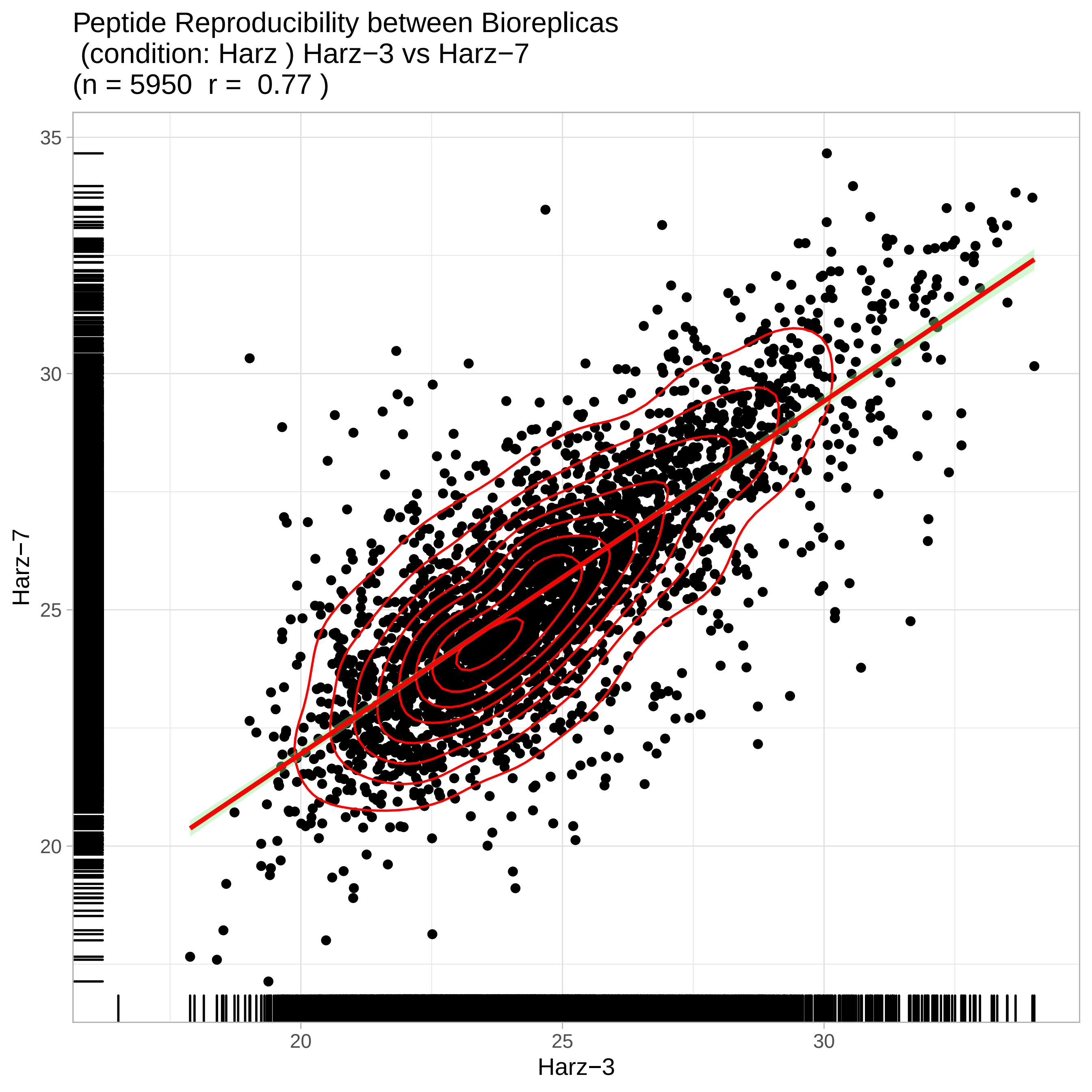

Peptide Reproducibility between Bioreplicas
(condition: Harz ) Harz−3 vs Harz−7
(n = 5950 r = 0.77 )
Harz−7
35
30
25
20
Harz−3
20
25
30

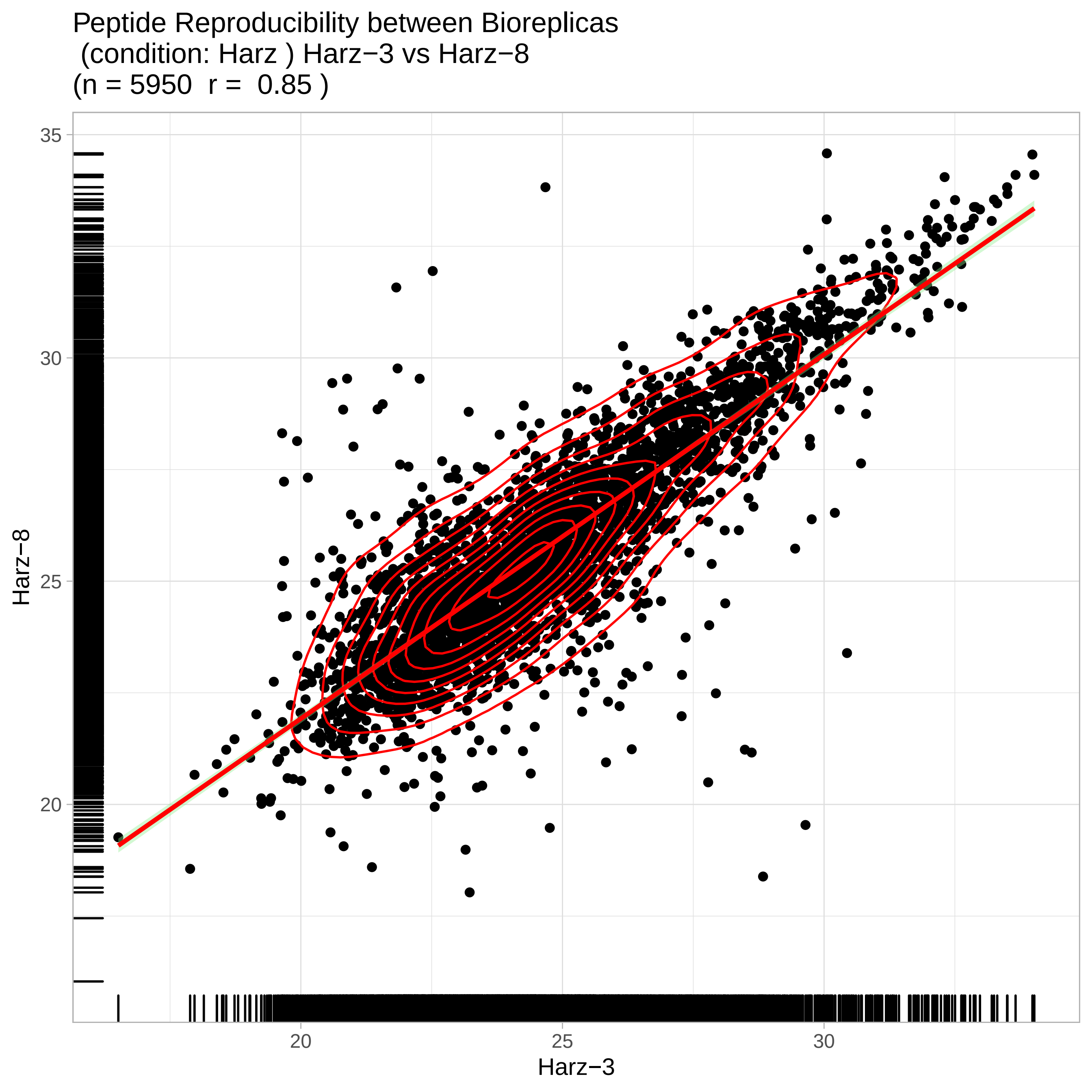

Peptide Reproducibility between Bioreplicas
(condition: Harz ) Harz−3 vs Harz−8
(n = 5950 r = 0.85 )
Harz−8
35
30
25
20
20
25
30
Harz−3

Peptide Reproducibility between Bioreplicas
(condition: Harz ) Harz−4 vs Harz−5
(n = 5950 r = 0.84 )

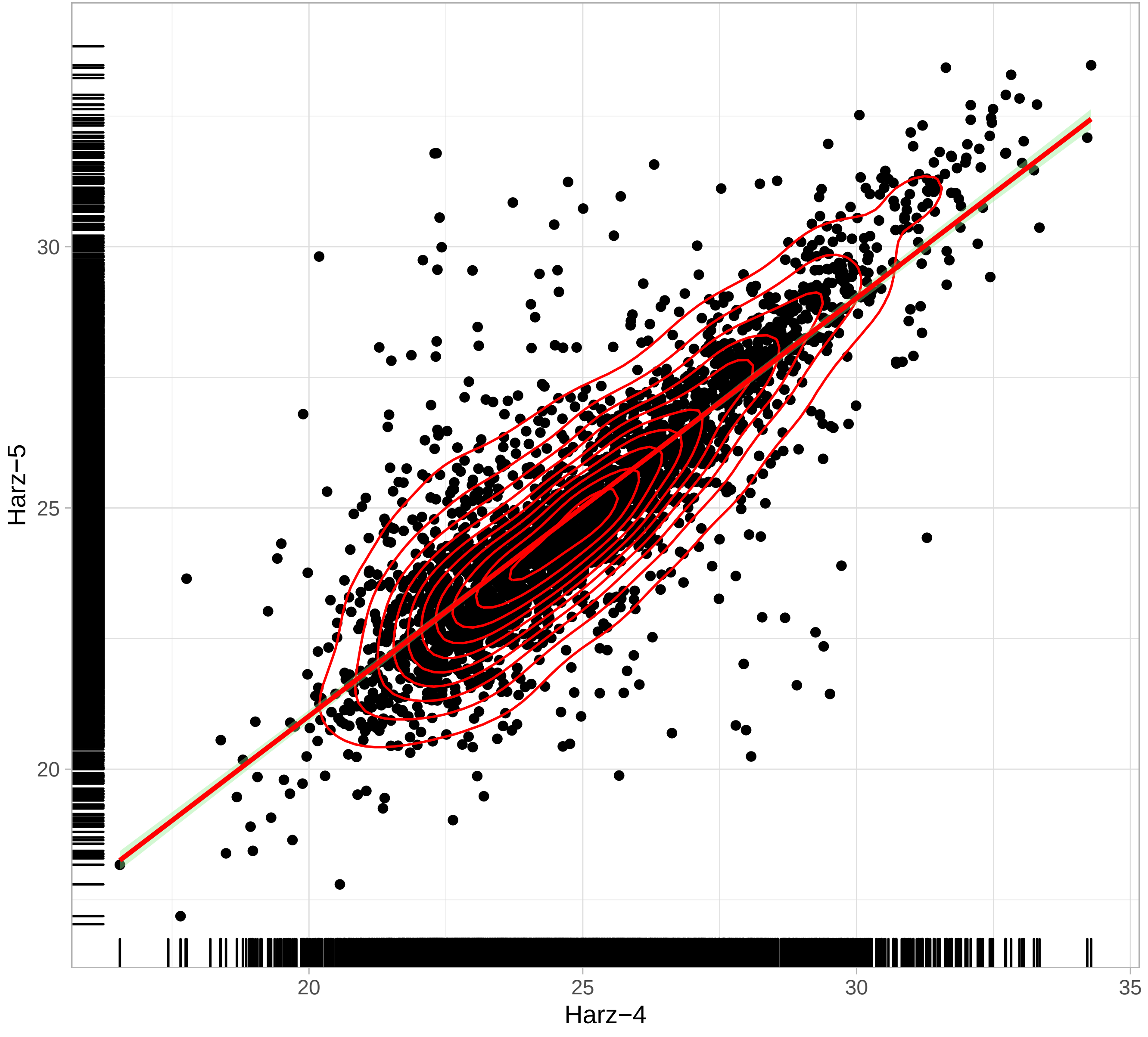

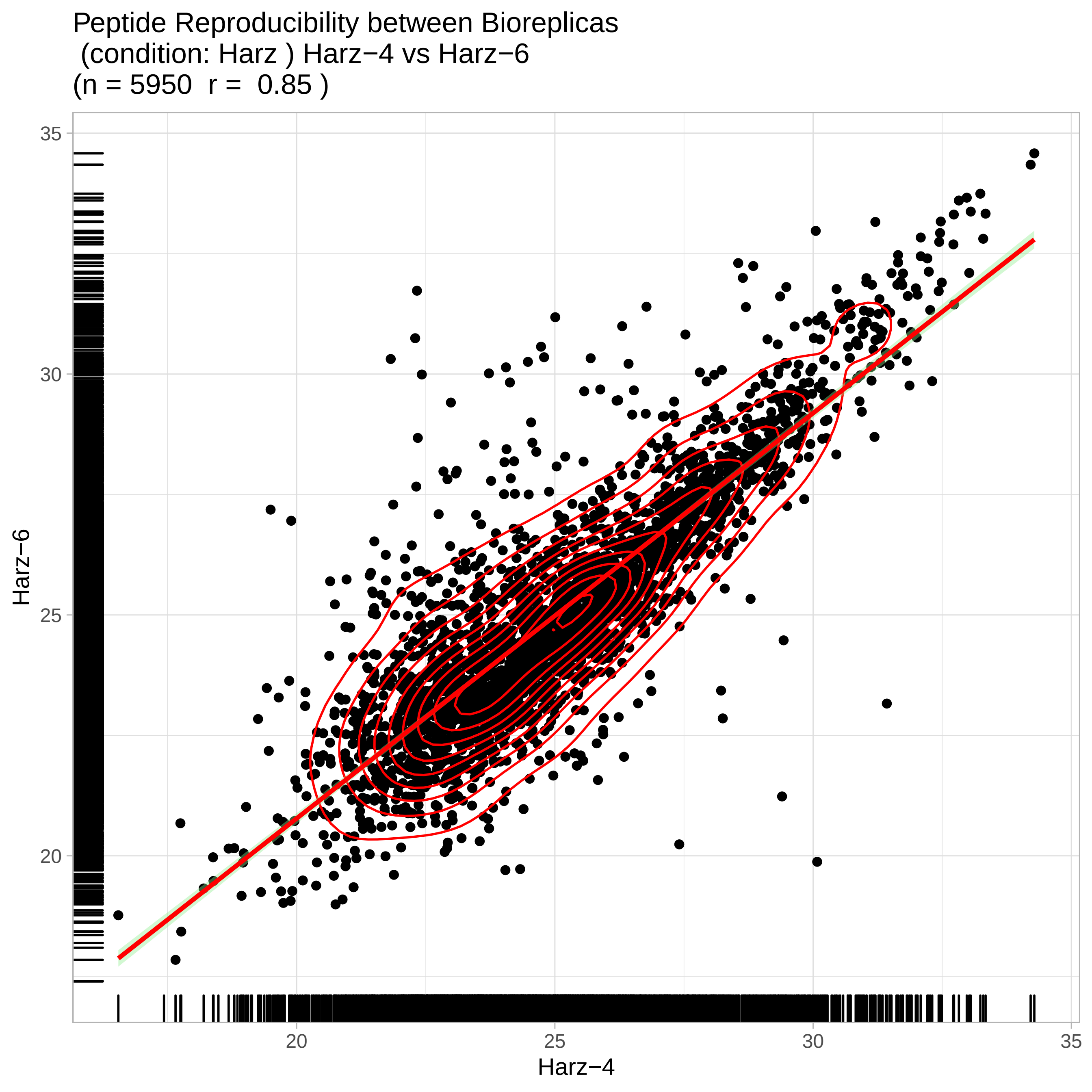

Peptide Reproducibility between Bioreplicas
(condition: Harz ) Harz−4 vs Harz−6
(n = 5950 r = 0.85 )
Harz−6
Harz−4
35
30
25
20
20
25
30
35

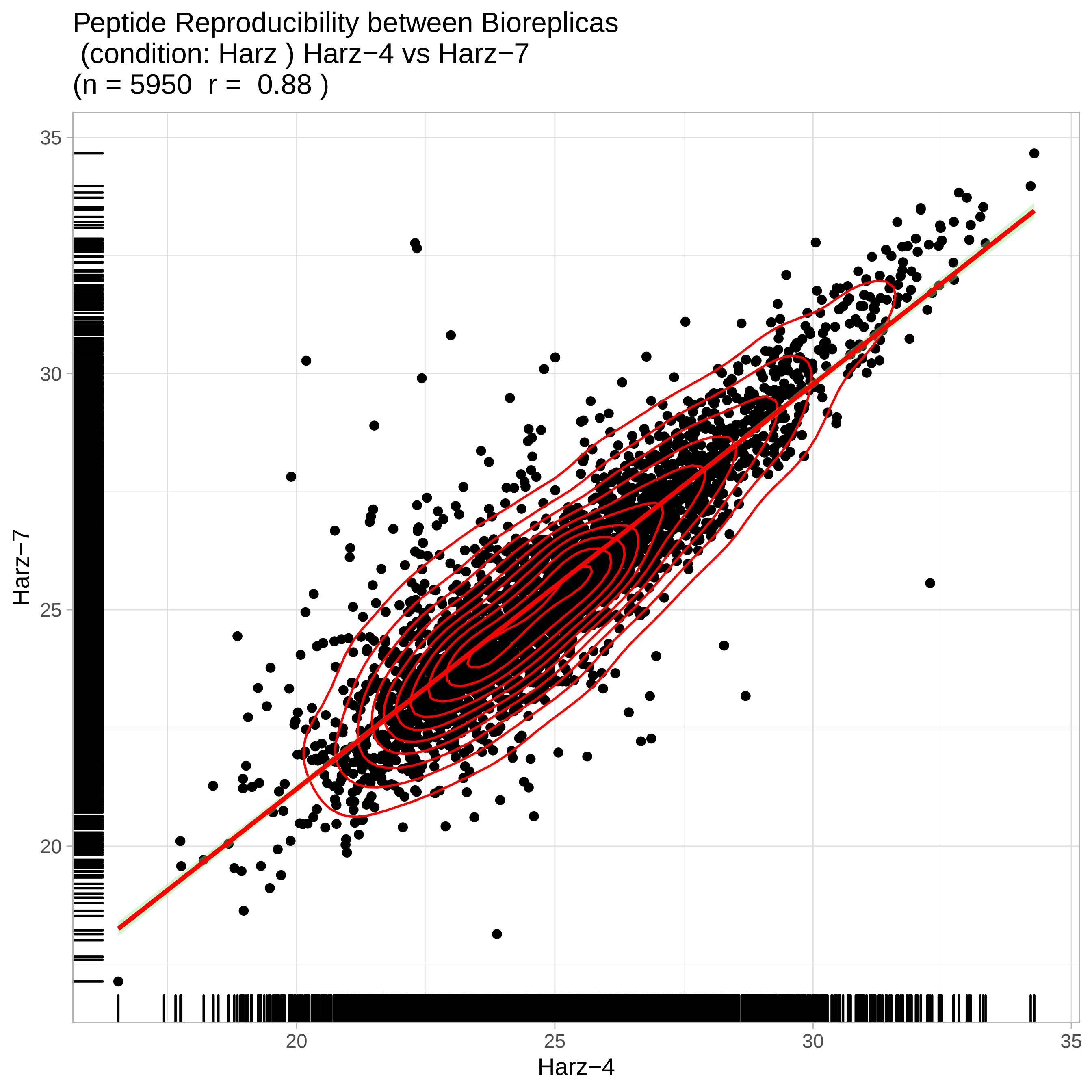
Peptide Reproducibility between Bioreplicas
(condition: Harz ) Harz−4 vs Harz−7
(n = 5950 r = 0.88 )
Harz−7
Harz−4
35
30
25
20
20
25
30
35

Peptide Reproducibility between Bioreplicas
(condition: Harz ) Harz−4 vs Harz−8
(n = 5950 r = 0.79 )

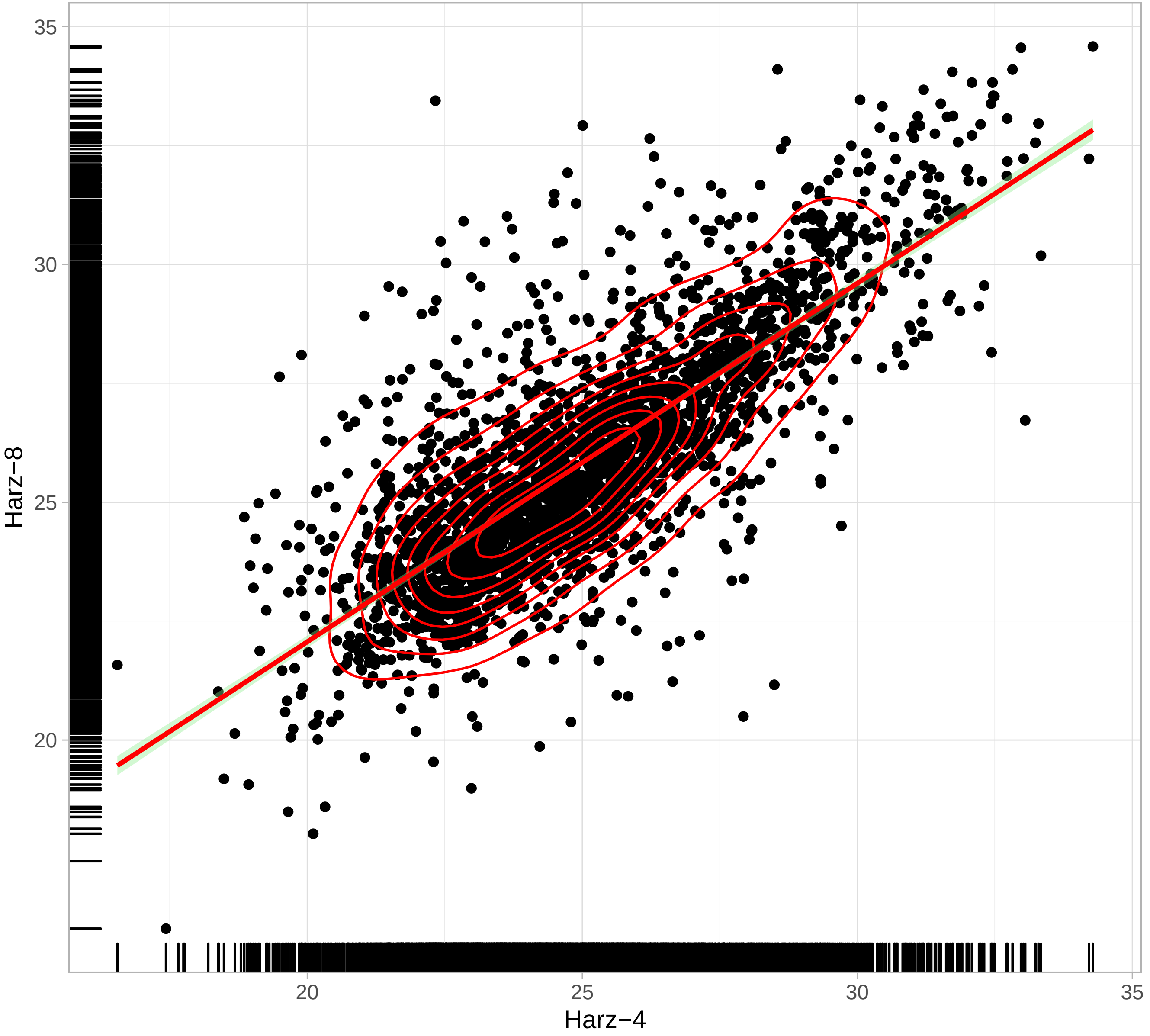

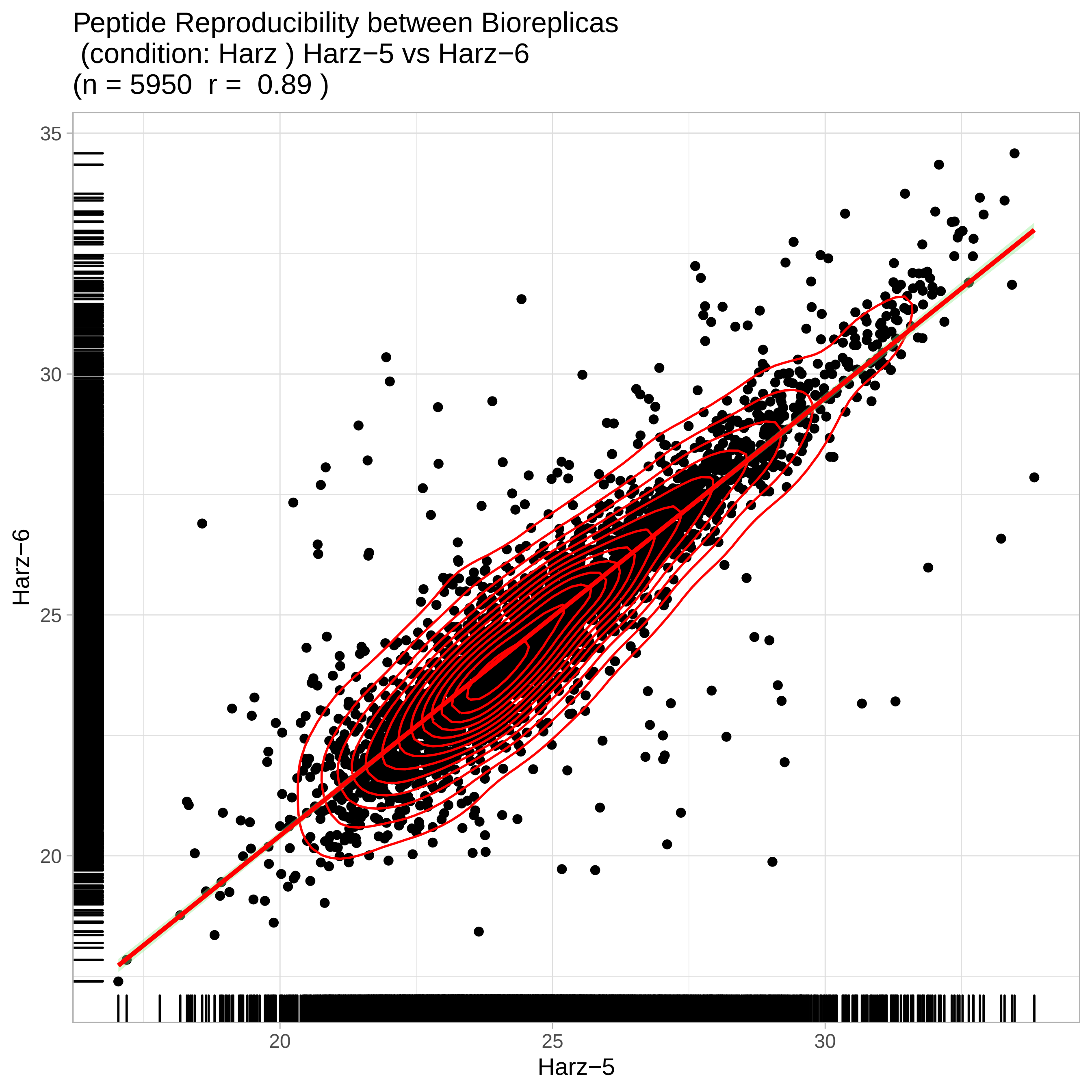
Peptide Reproducibility between Bioreplicas
(condition: Harz ) Harz−5 vs Harz−6
(n = 5950 r = 0.89 )
Harz−6
Harz−5
35
30
25
20
20
25
30

Peptide Reproducibility between Bioreplicas
(condition: Harz ) Harz−5 vs Harz−7
(n = 5950 r = 0.87 )

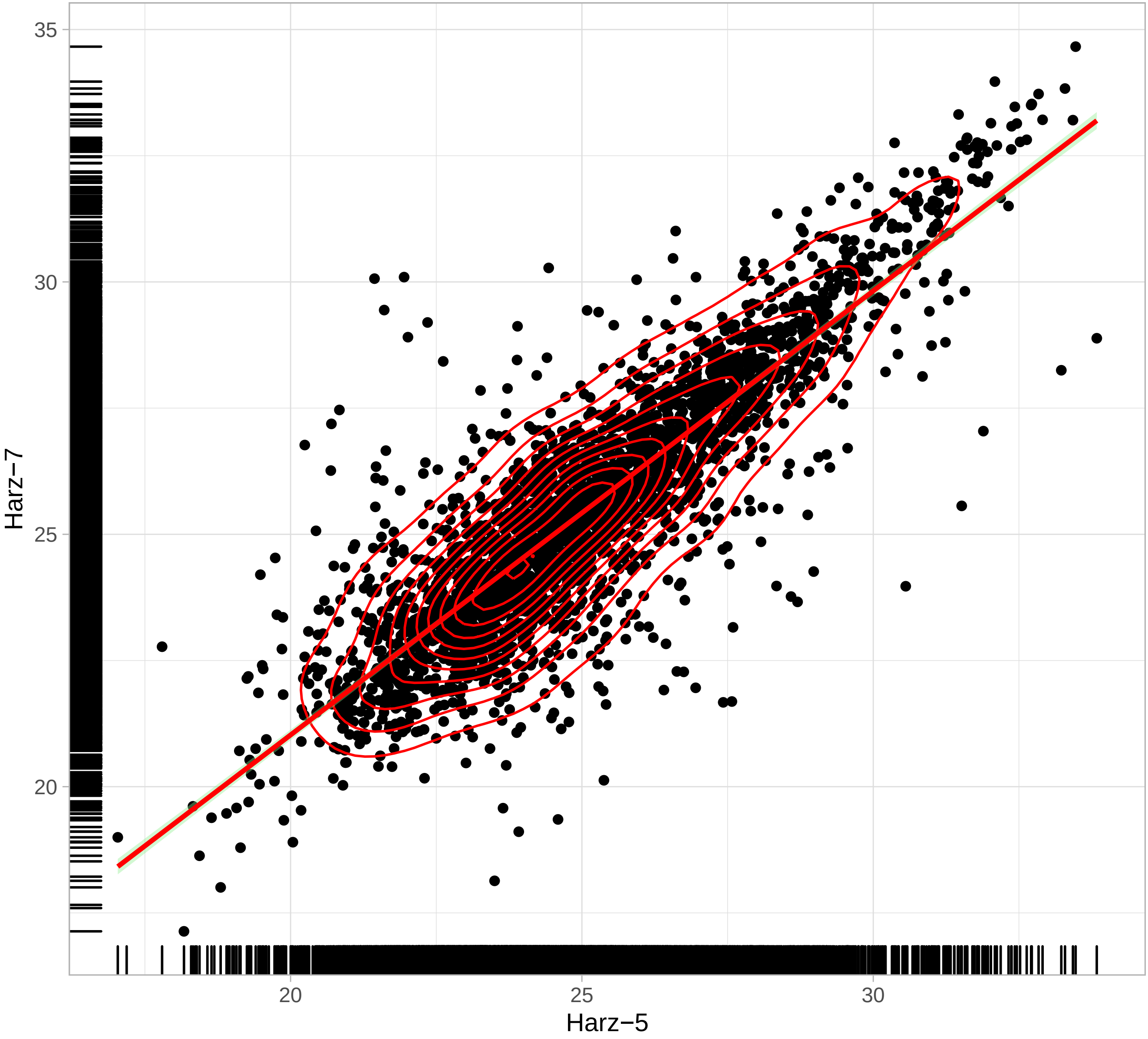

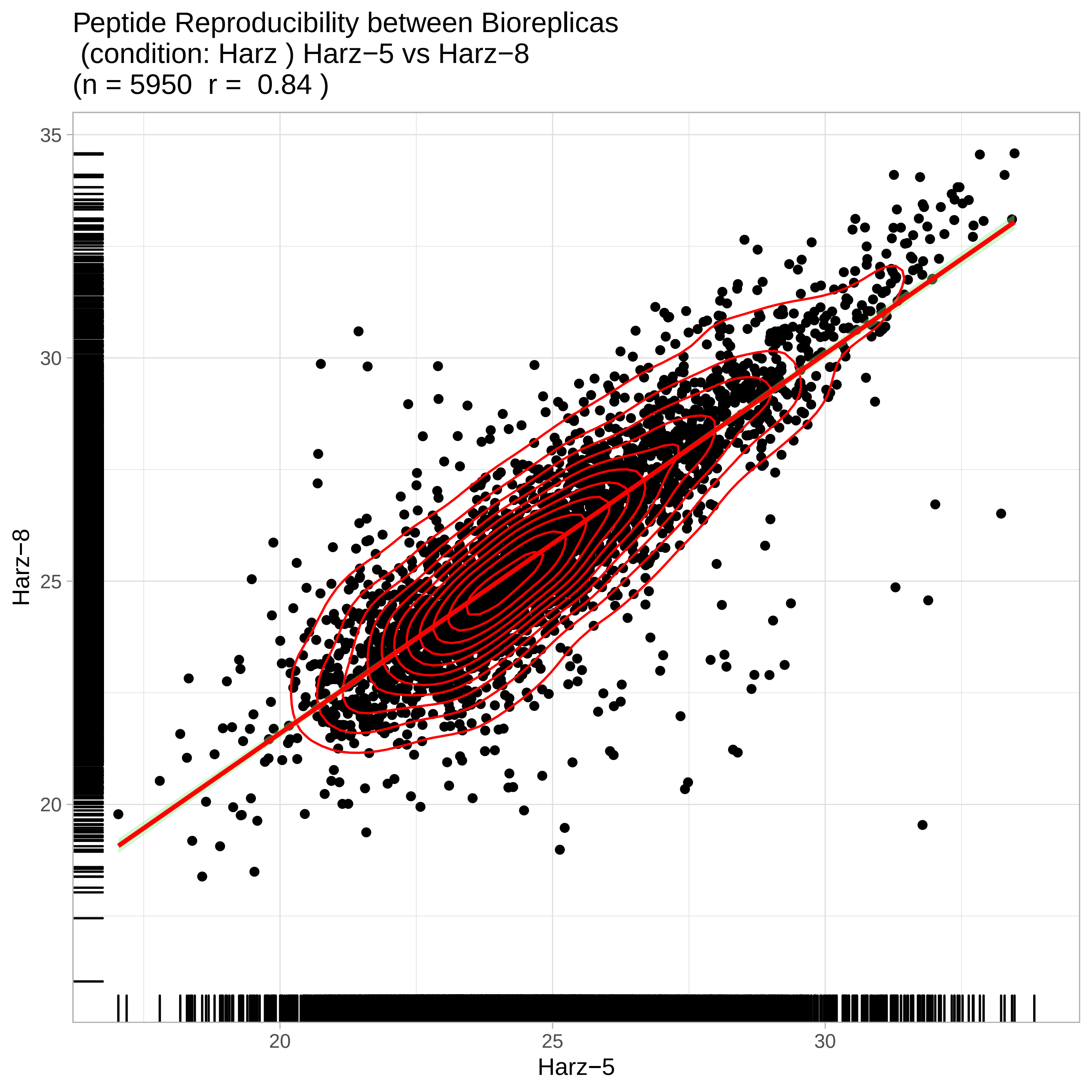
Peptide Reproducibility between Bioreplicas
(condition: Harz ) Harz−5 vs Harz−8
(n = 5950 r = 0.84 )
35
30
25
20
Harz−8
20
25
30
Harz−5

Peptide Reproducibility between Bioreplicas
(condition: Harz ) Harz−6 vs Harz−7
(n = 5950  r =  0.89 )

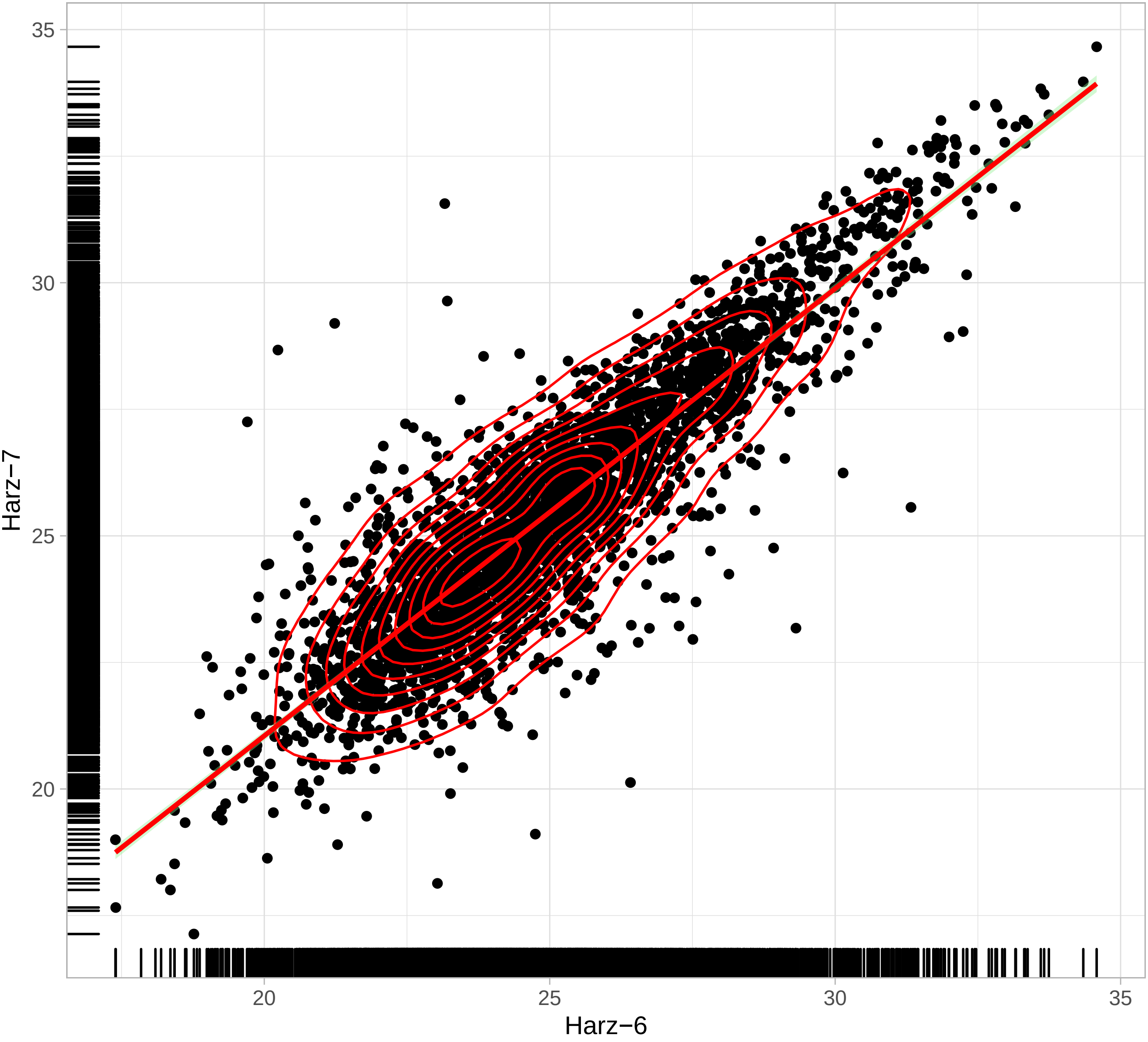

Peptide Reproducibility between Bioreplicas
(condition: Harz ) Harz−6 vs Harz−8
(n = 5950 r = 0.85 )

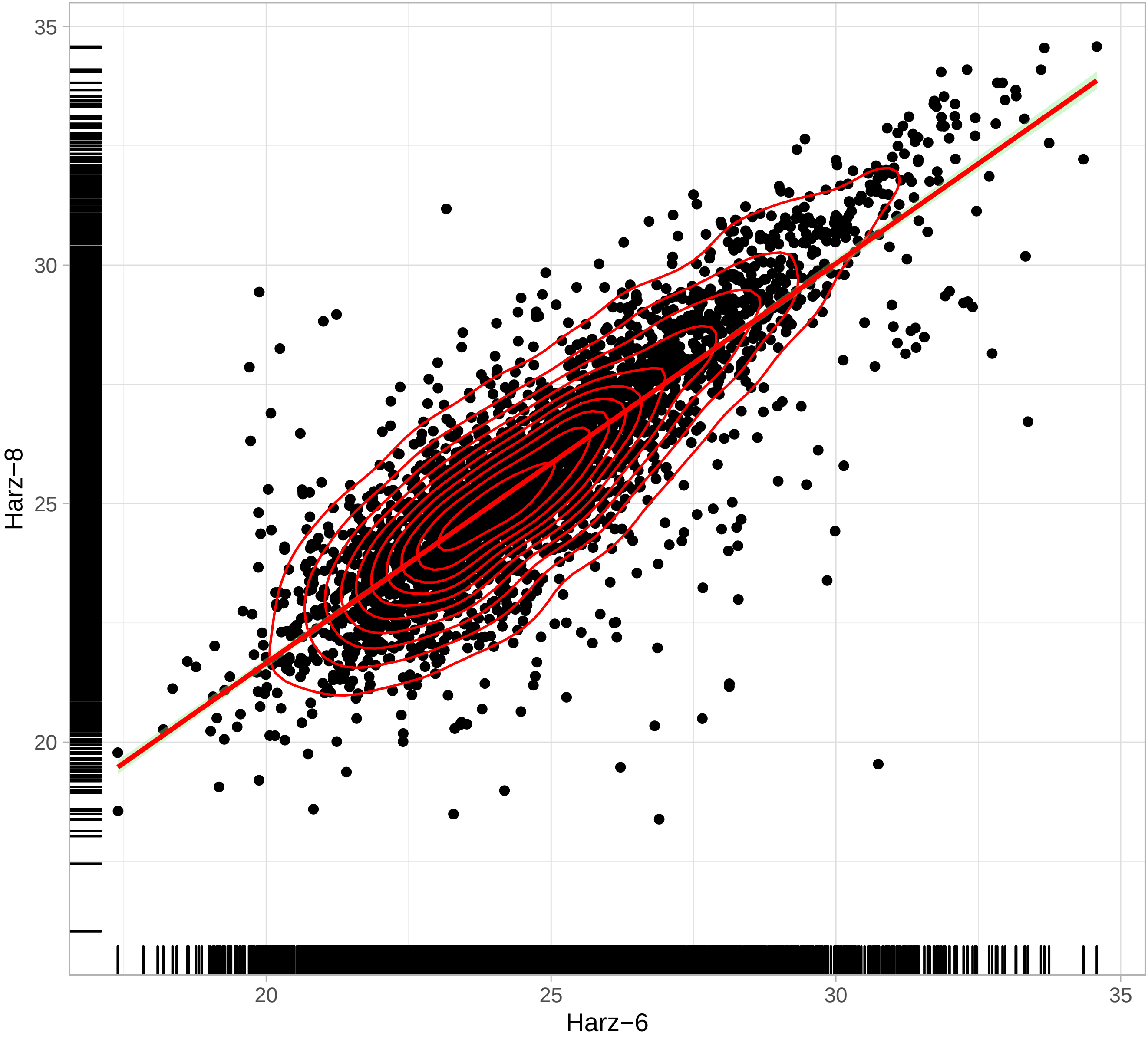

Peptide Reproducibility between Bioreplicas
(condition: Harz ) Harz−7 vs Harz−8
(n = 5950 r = 0.86 )

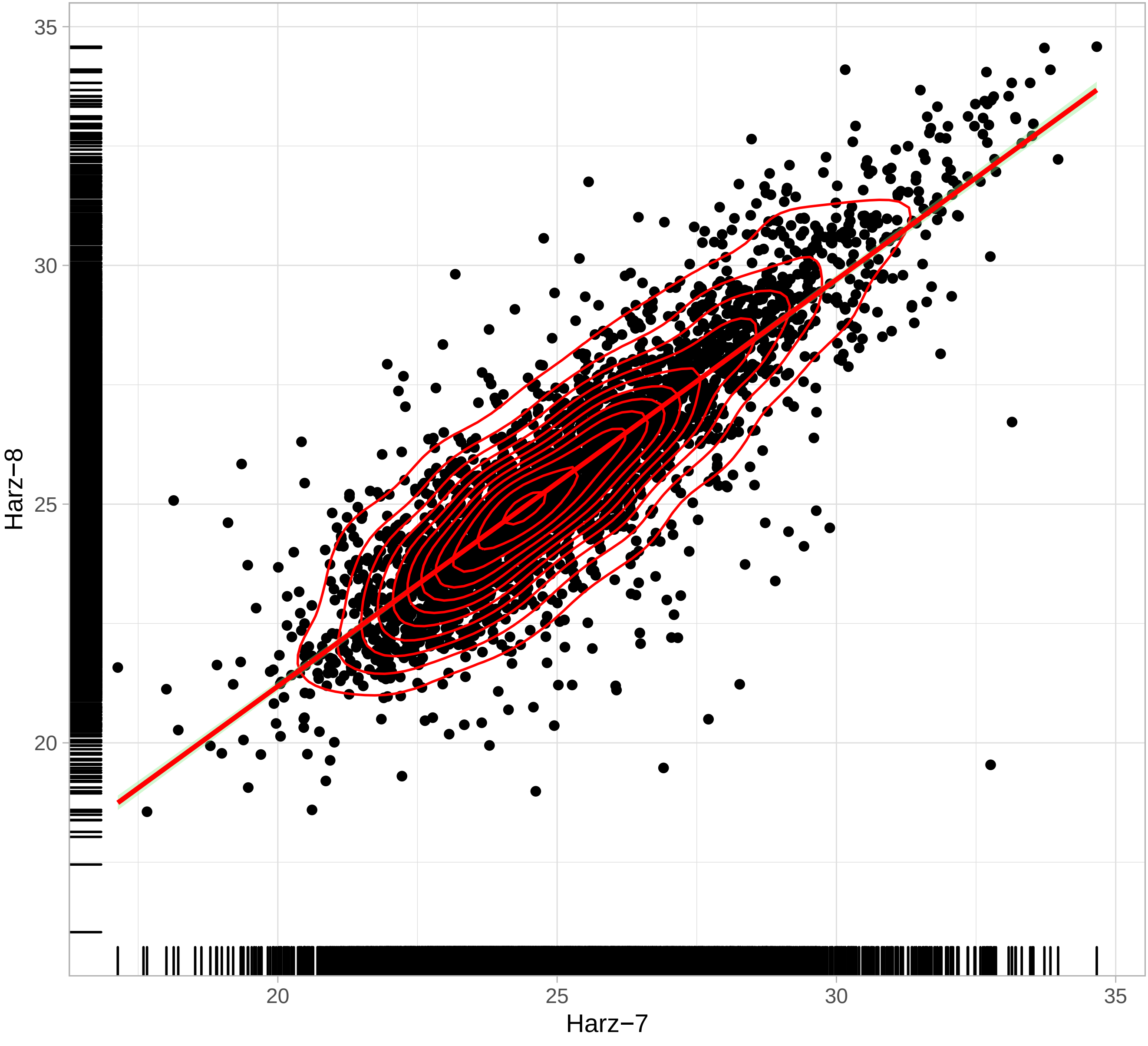

Peptide Reproducibility between Bioreplicas
(condition: SW ) SW−1 vs SW−2
(n = 6055  r =  0.87 )

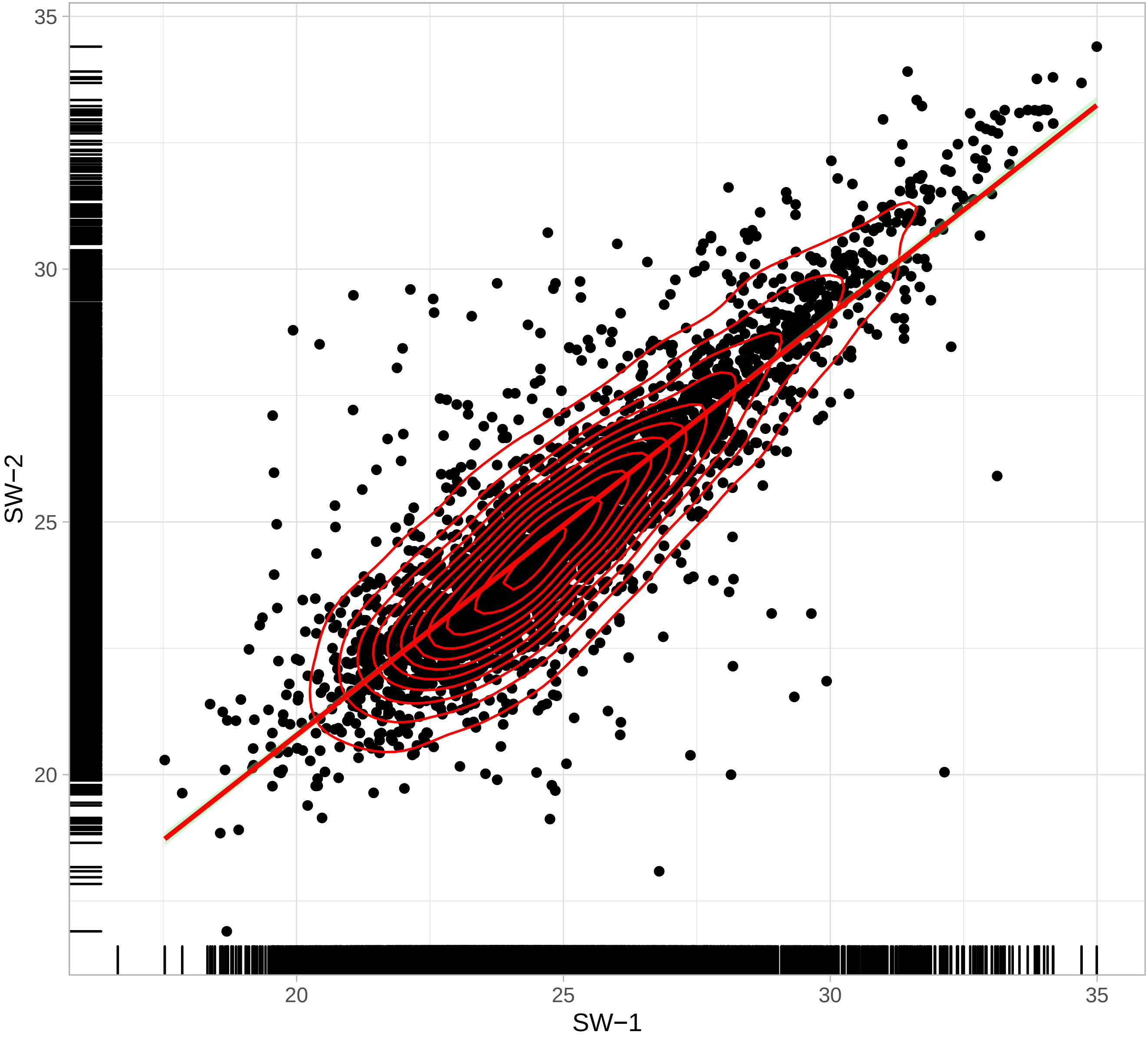

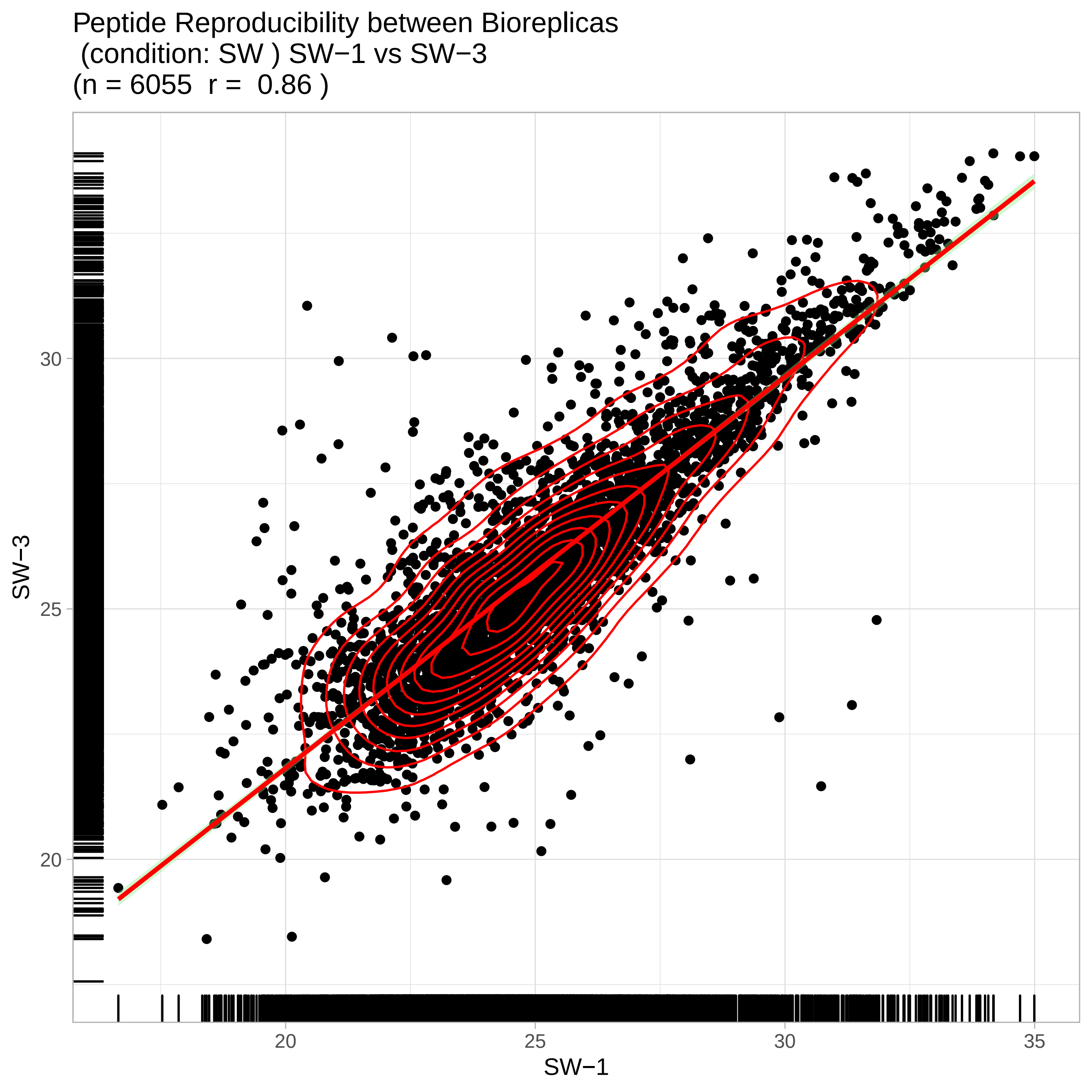
Peptide Reproducibility between Bioreplicas
(condition: SW ) SW−1 vs SW−3
(n = 6055 r = 0.86 )
SW−3
SW−1
20
25
30
35

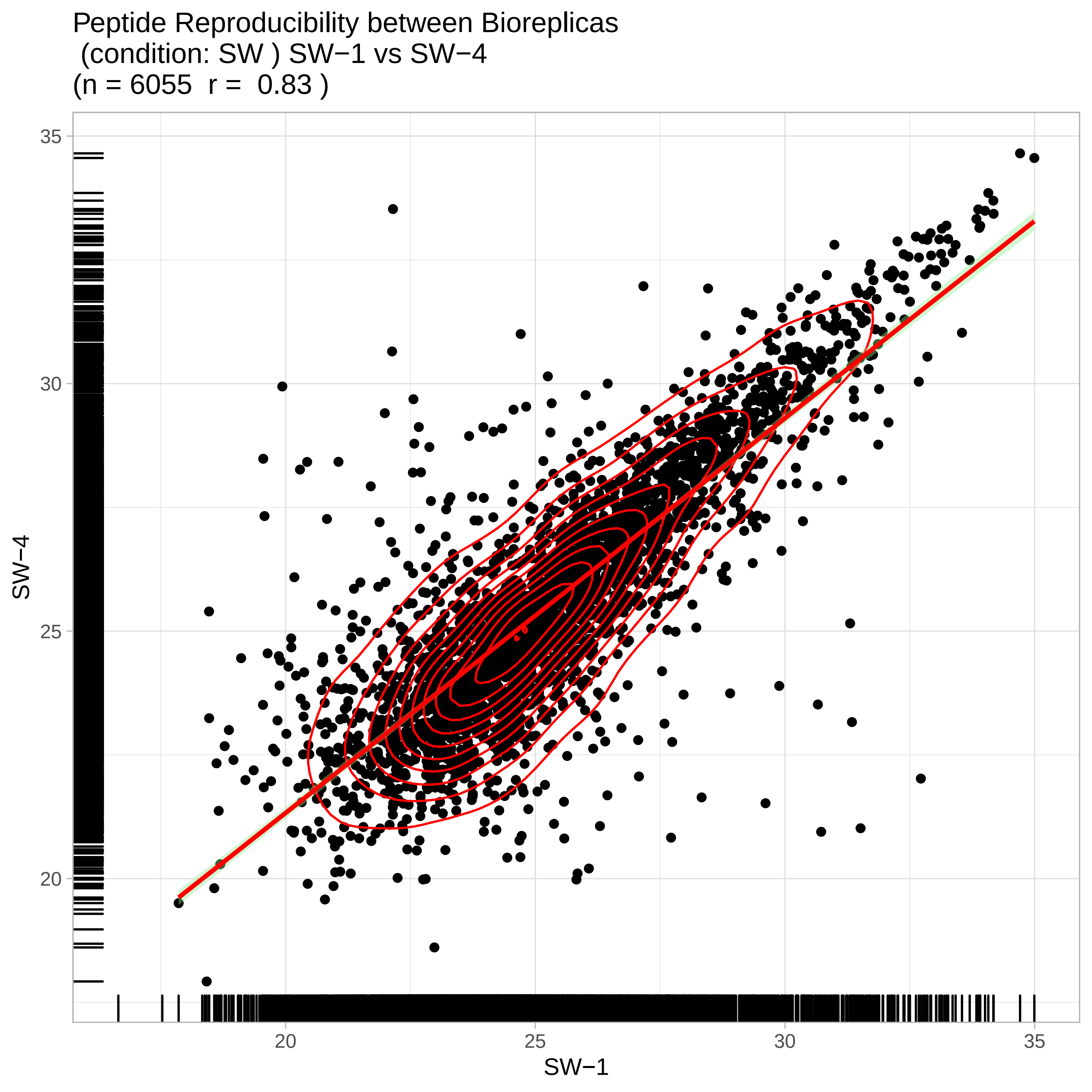
Peptide Reproducibility between Bioreplicas
(condition: SW ) SW−1 vs SW−4
(n = 6055 r = 0.83 )
SW−4
35
30
25
20
SW−1
20
25
30
35

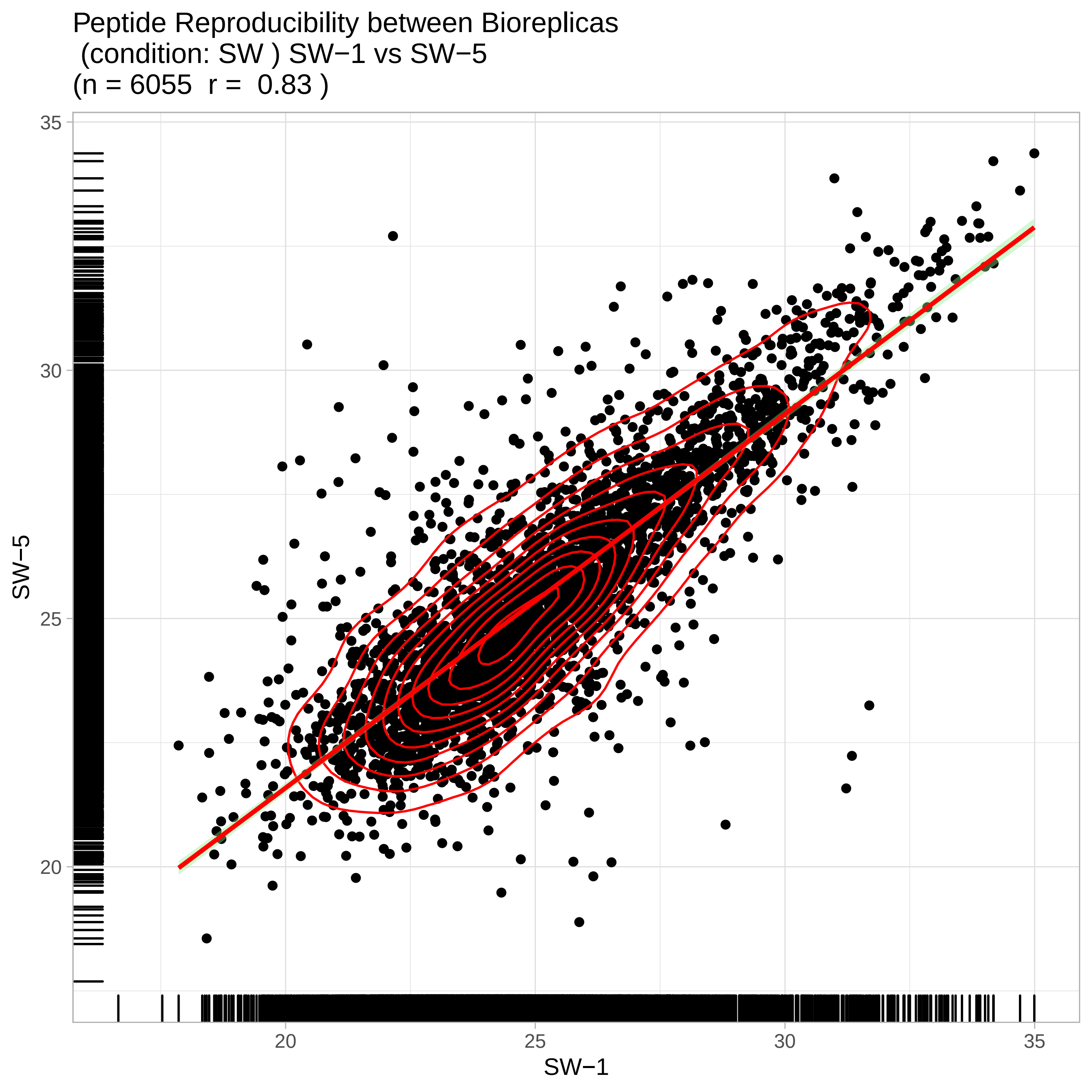

Peptide Reproducibility between Bioreplicas
(condition: SW ) SW−1 vs SW−5
(n = 6055 r = 0.83 )
SW−5
35
30
25
20
SW−1
20
25
30
35

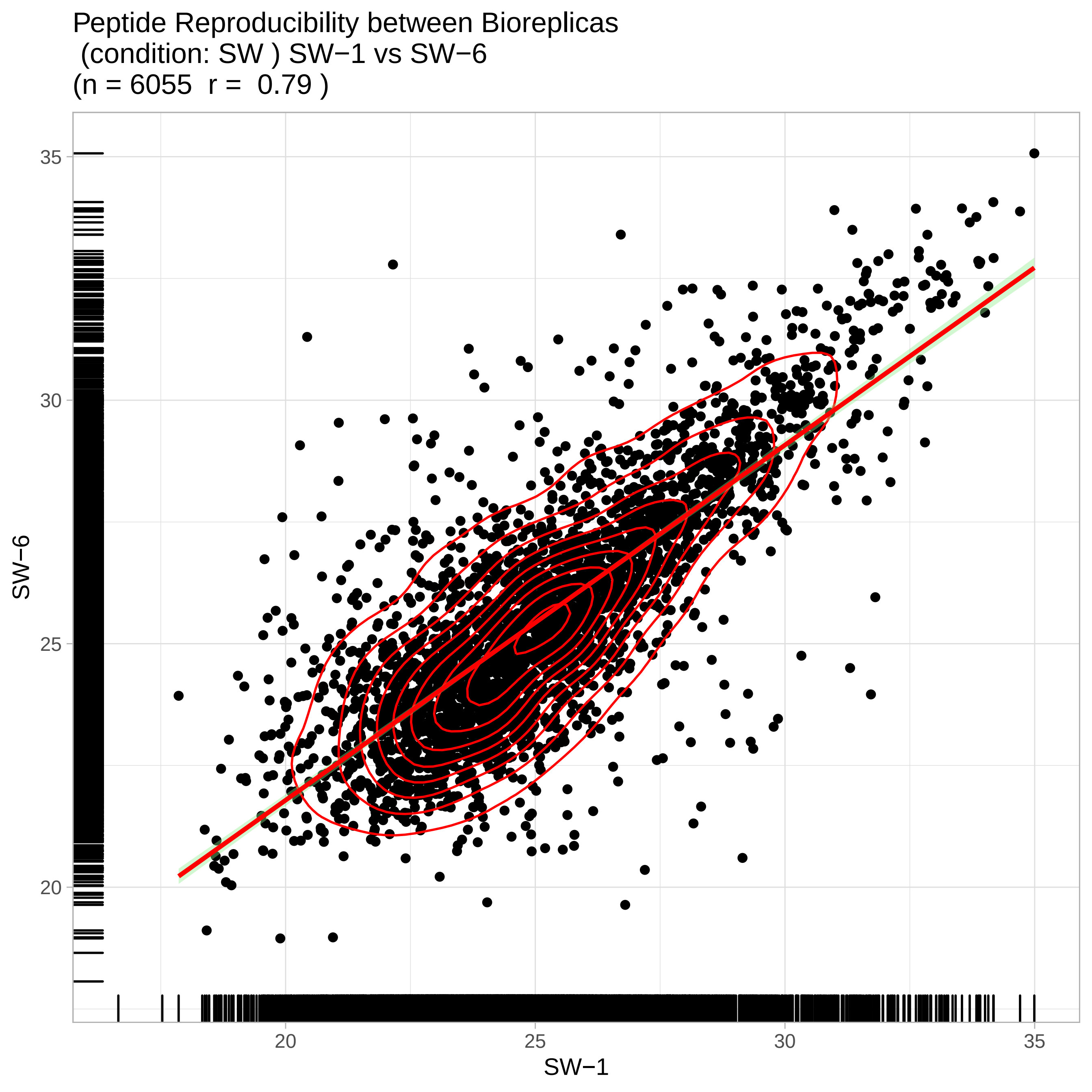
Peptide Reproducibility between Bioreplicas
(condition: SW ) SW−1 vs SW−6
(n = 6055 r = 0.79 )
SW−6
SW−1
20
25
30
35
20
25
30
35

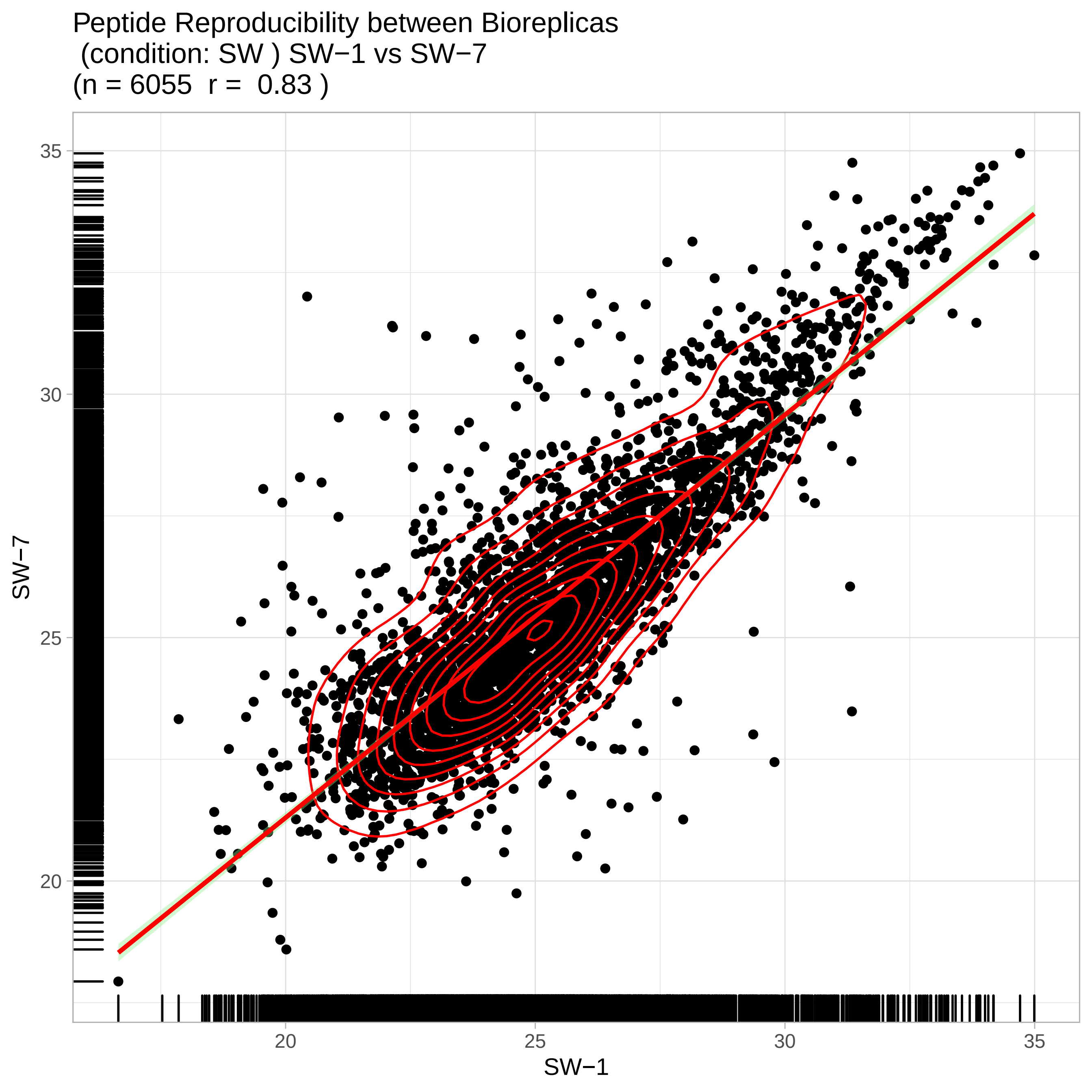
Peptide Reproducibility between Bioreplicas
(condition: SW ) SW−1 vs SW−7
(n = 6055 r = 0.83 )
35
30
25
20
SW−7
20
25
30
35
SW−1

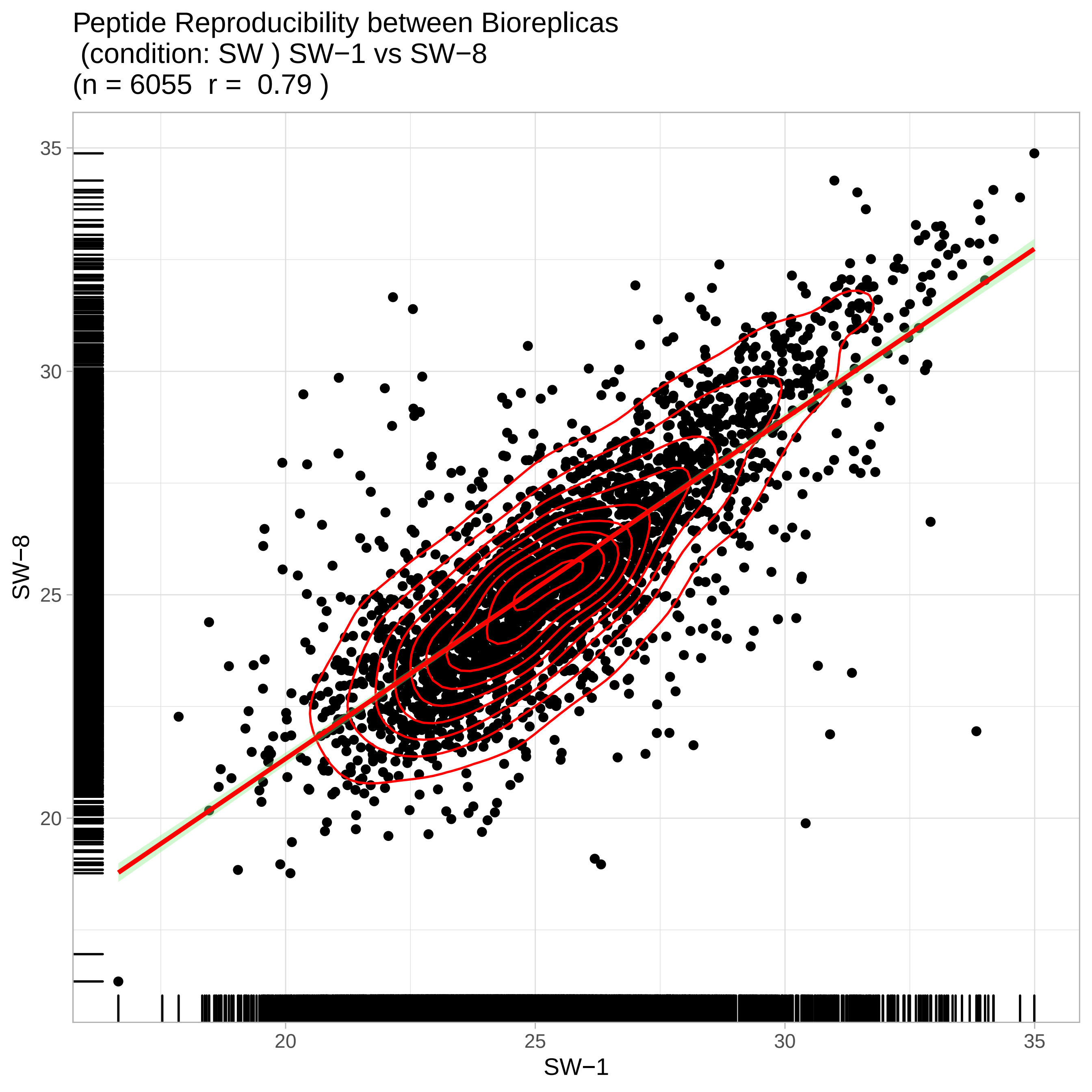

Peptide Reproducibility between Bioreplicas
(condition: SW ) SW−1 vs SW−8
(n = 6055  r =  0.79 )
35
30
25
20
SW−8
20
25
30
35
SW−1

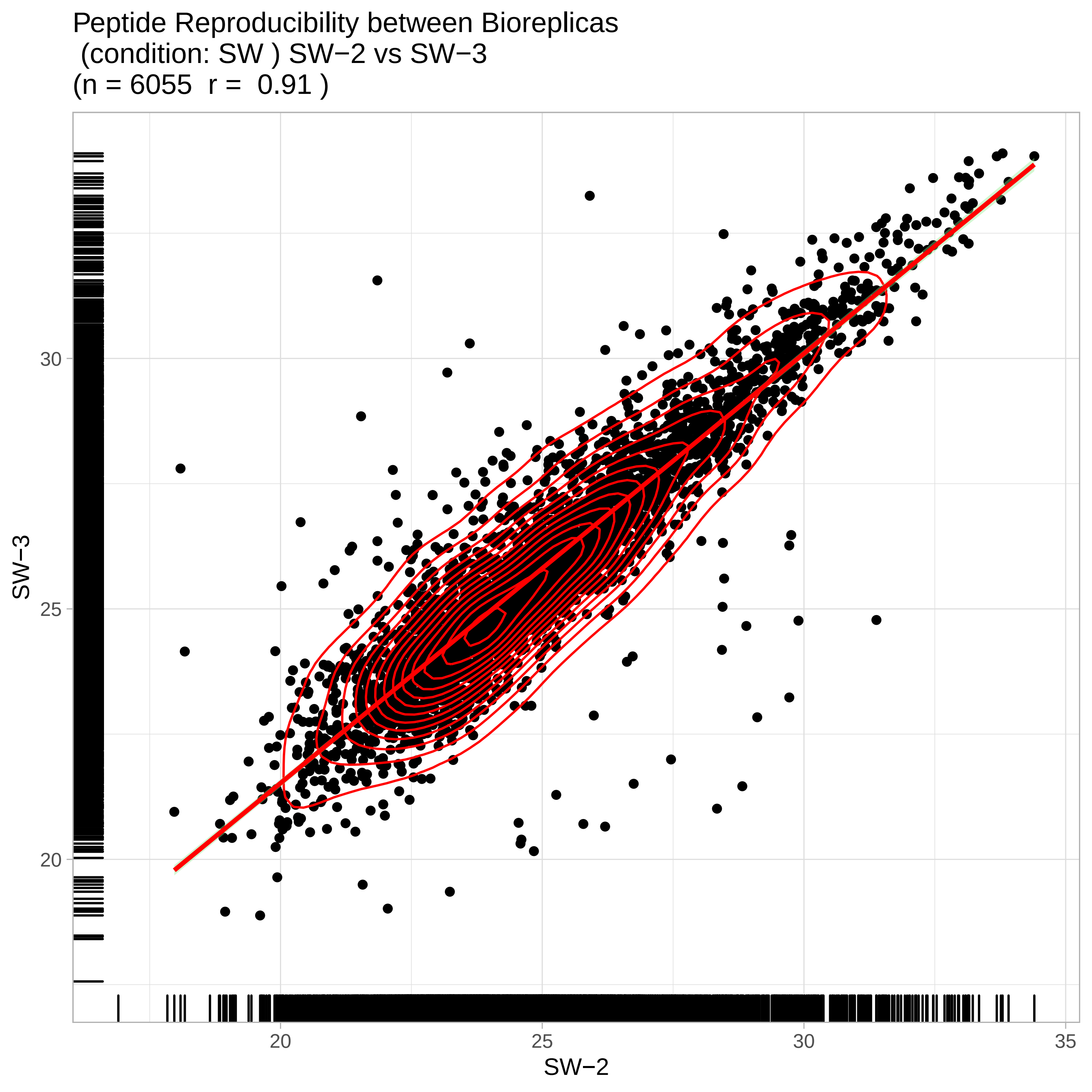

Peptide Reproducibility between Bioreplicas
(condition: SW ) SW−2 vs SW−3
(n = 6055 r = 0.91 )
SW−3
30
25
20
SW−2
20
25
30
35

Peptide Reproducibility between Bioreplicas
(condition: SW ) SW−2 vs SW−4
(n = 6055 r = 0.85 )

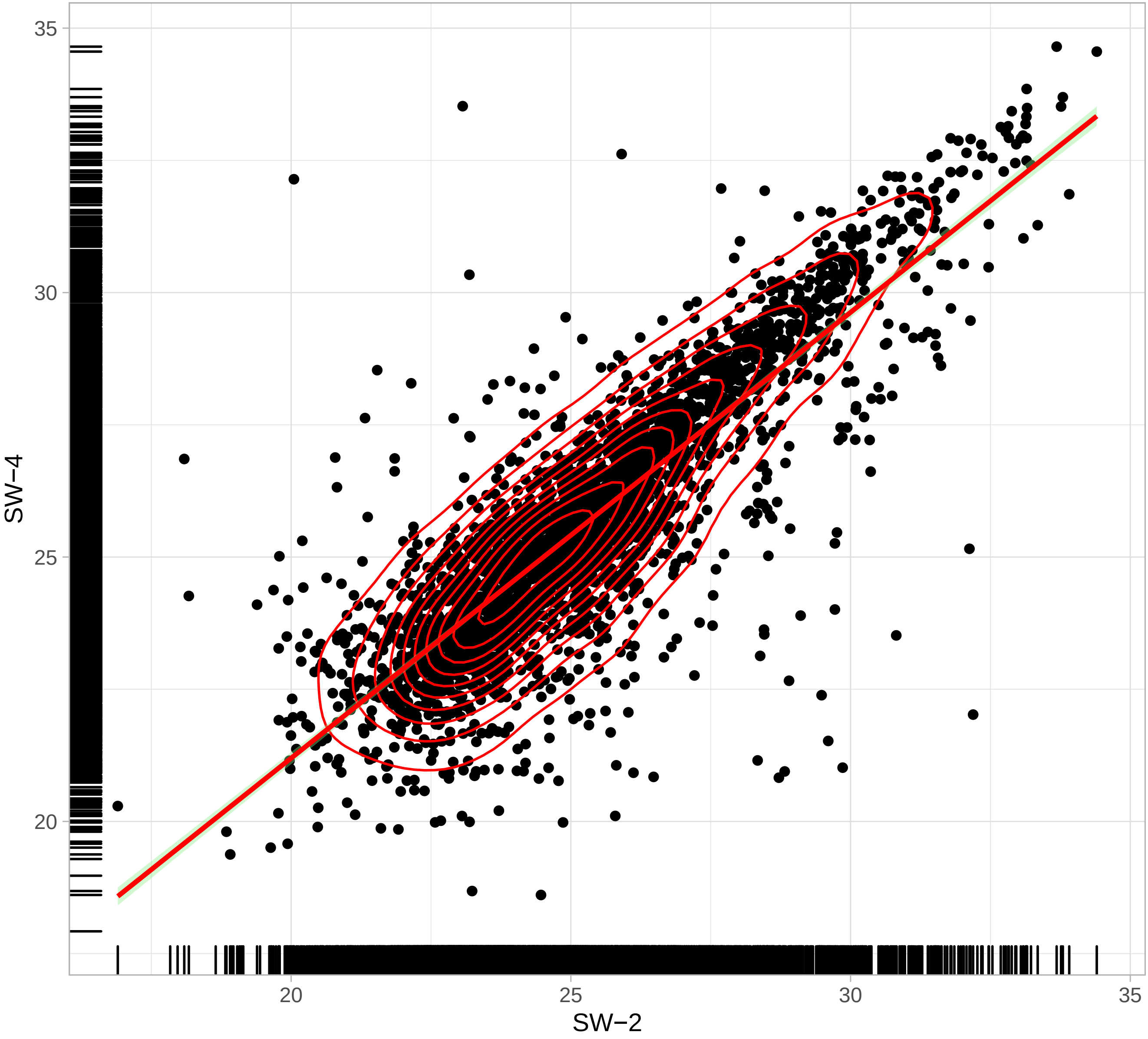

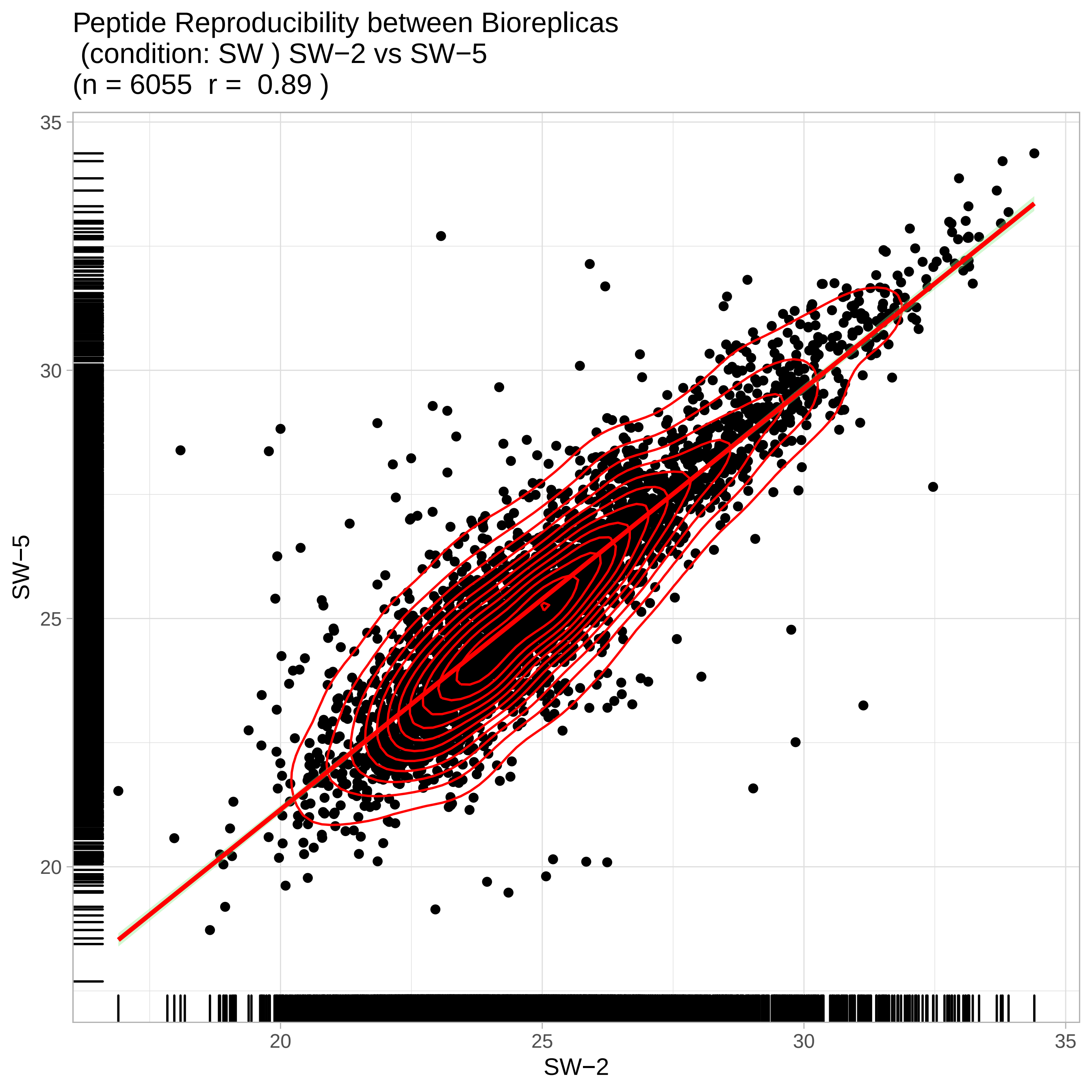
Peptide Reproducibility between Bioreplicas
(condition: SW ) SW−2 vs SW−5
(n = 6055 r = 0.89 )
35
30
25
20
SW−5
20
25
30
35
SW−2

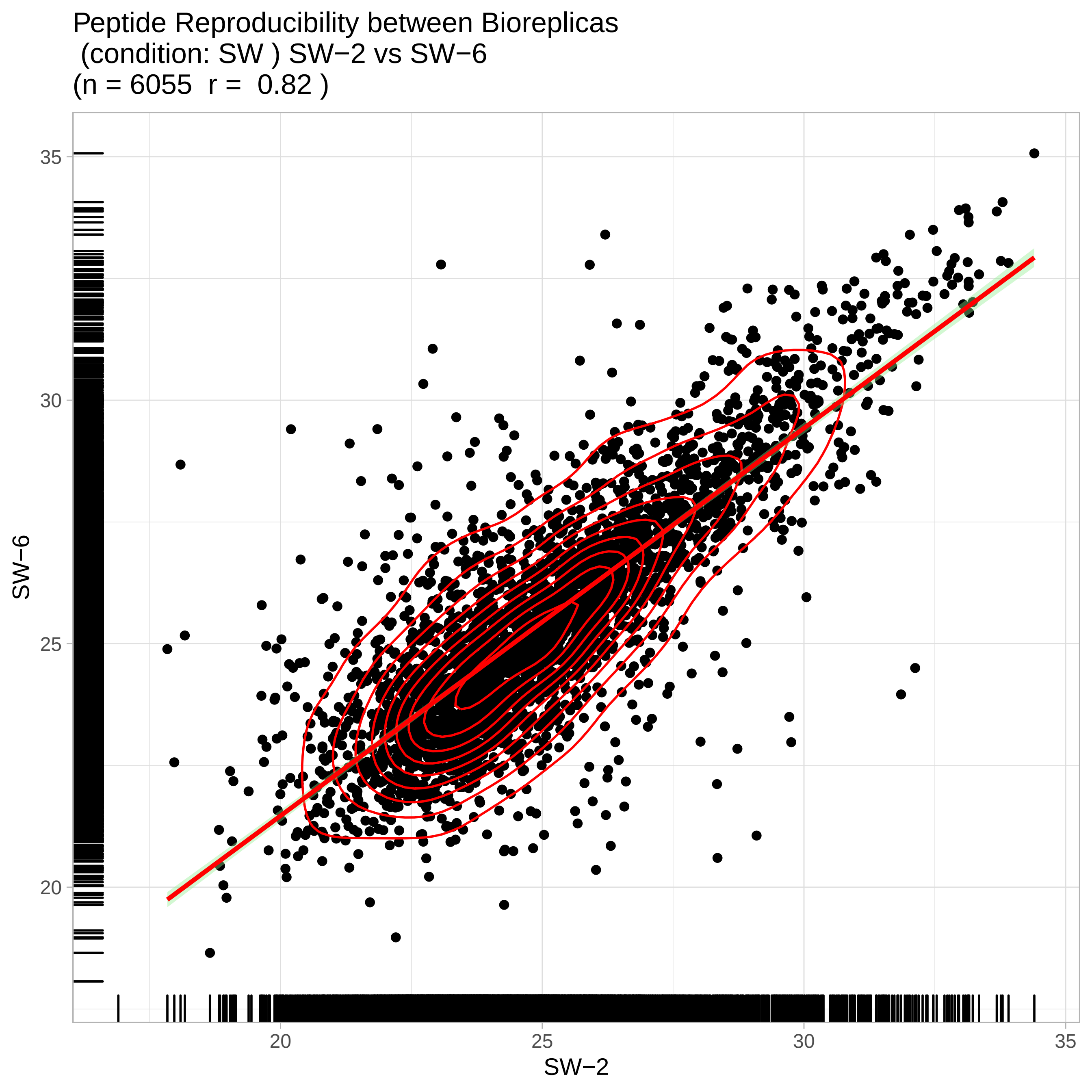

Peptide Reproducibility between Bioreplicas
(condition: SW ) SW−2 vs SW−6
(n = 6055 r = 0.82 )
SW−6
35
30
25
20
SW−2
20
25
30
35

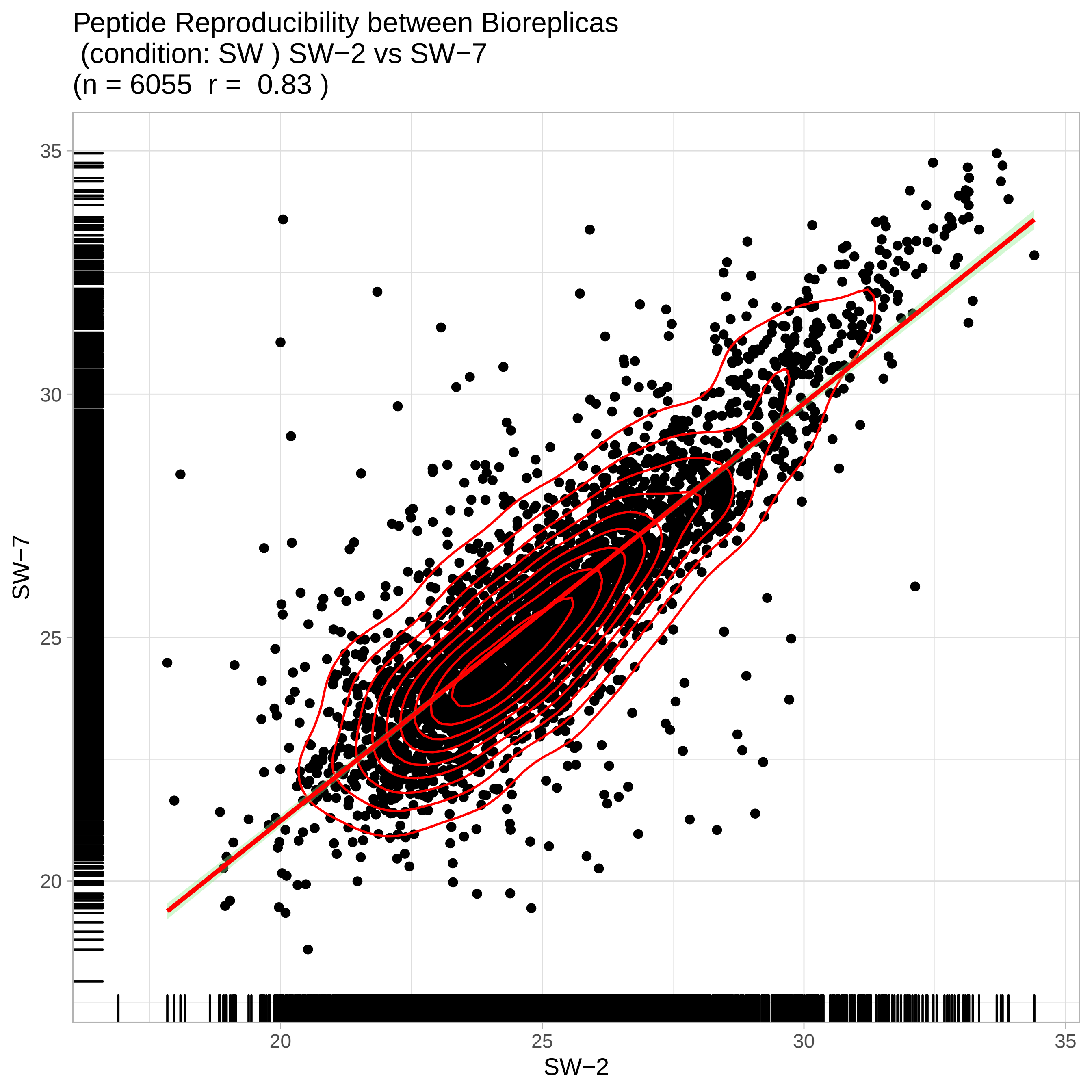
Peptide Reproducibility between Bioreplicas
(condition: SW ) SW−2 vs SW−7
(n = 6055 r = 0.83 )
SW−7
35
30
25
20
SW−2
20
25
30
35

Peptide Reproducibility between Bioreplicas
(condition: SW ) SW−2 vs SW−8
(n = 6055 r = 0.85 )

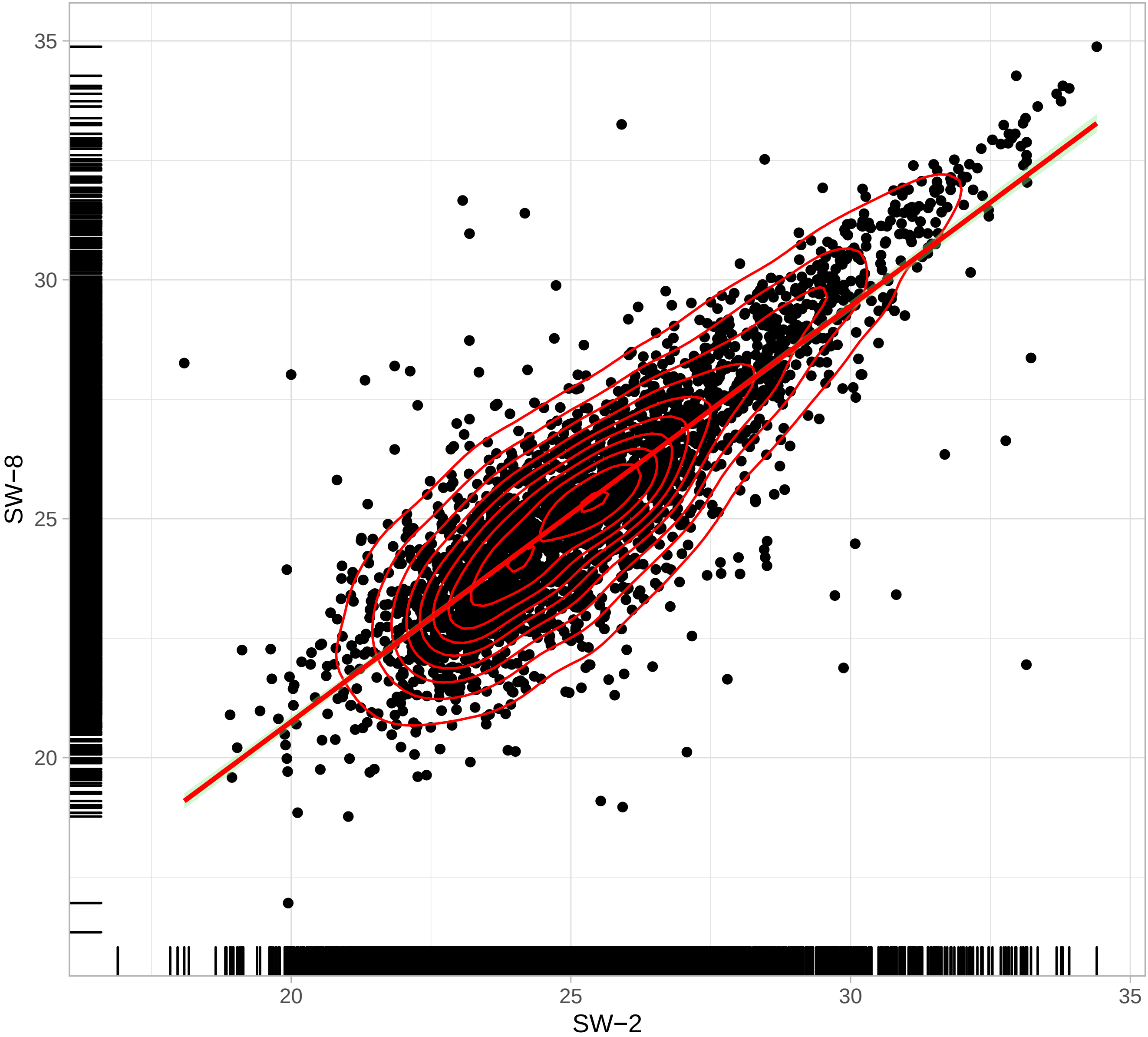

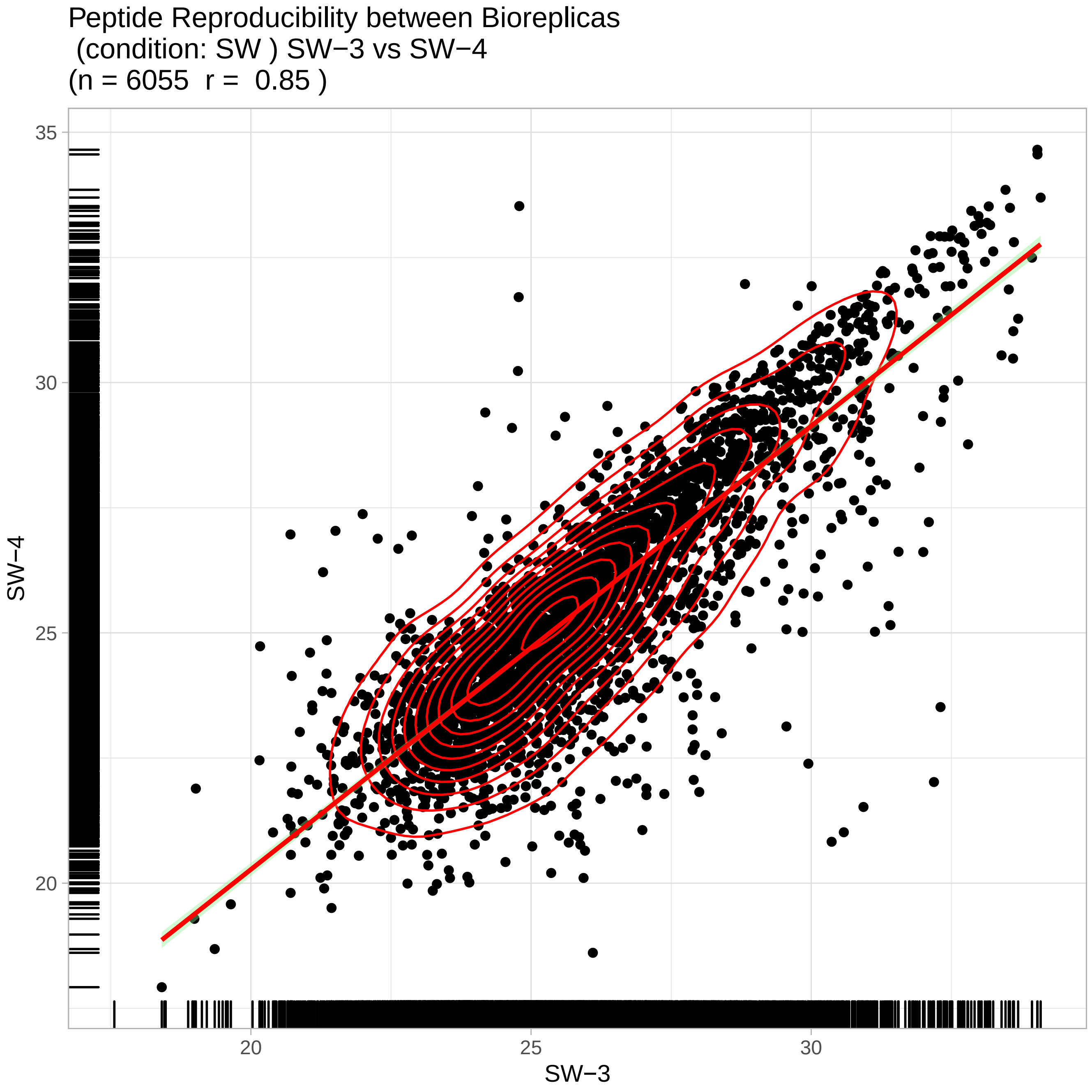
Peptide Reproducibility between Bioreplicas
(condition: SW ) SW−3 vs SW−4
(n = 6055 r = 0.85 )
SW−4
SW−3
35
30
25
20
20
25
30

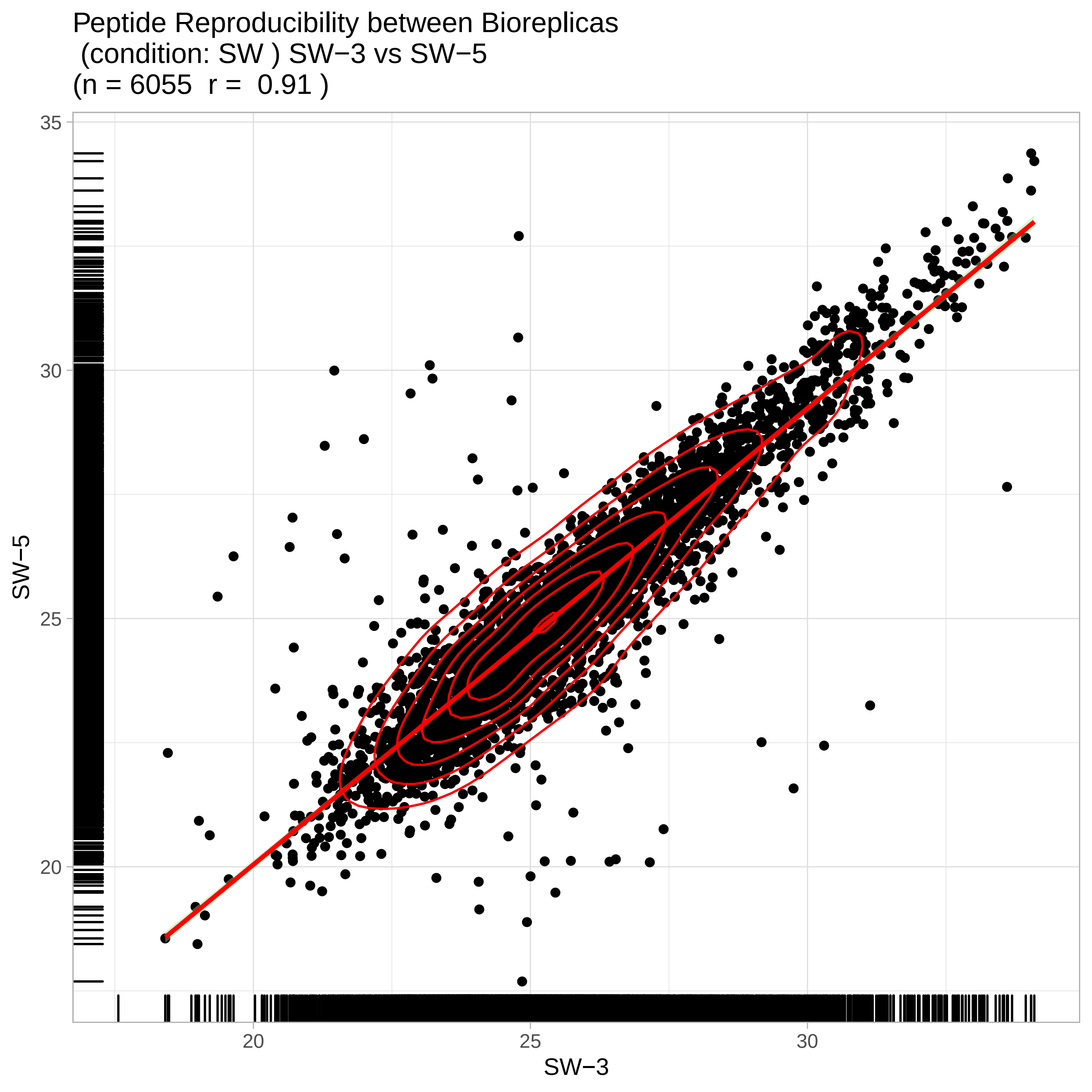
Peptide Reproducibility between Bioreplicas
(condition: SW ) SW−3 vs SW−5
(n = 6055  r =  0.91 )
35
30
25
20
SW−5
20
25
30
SW−3

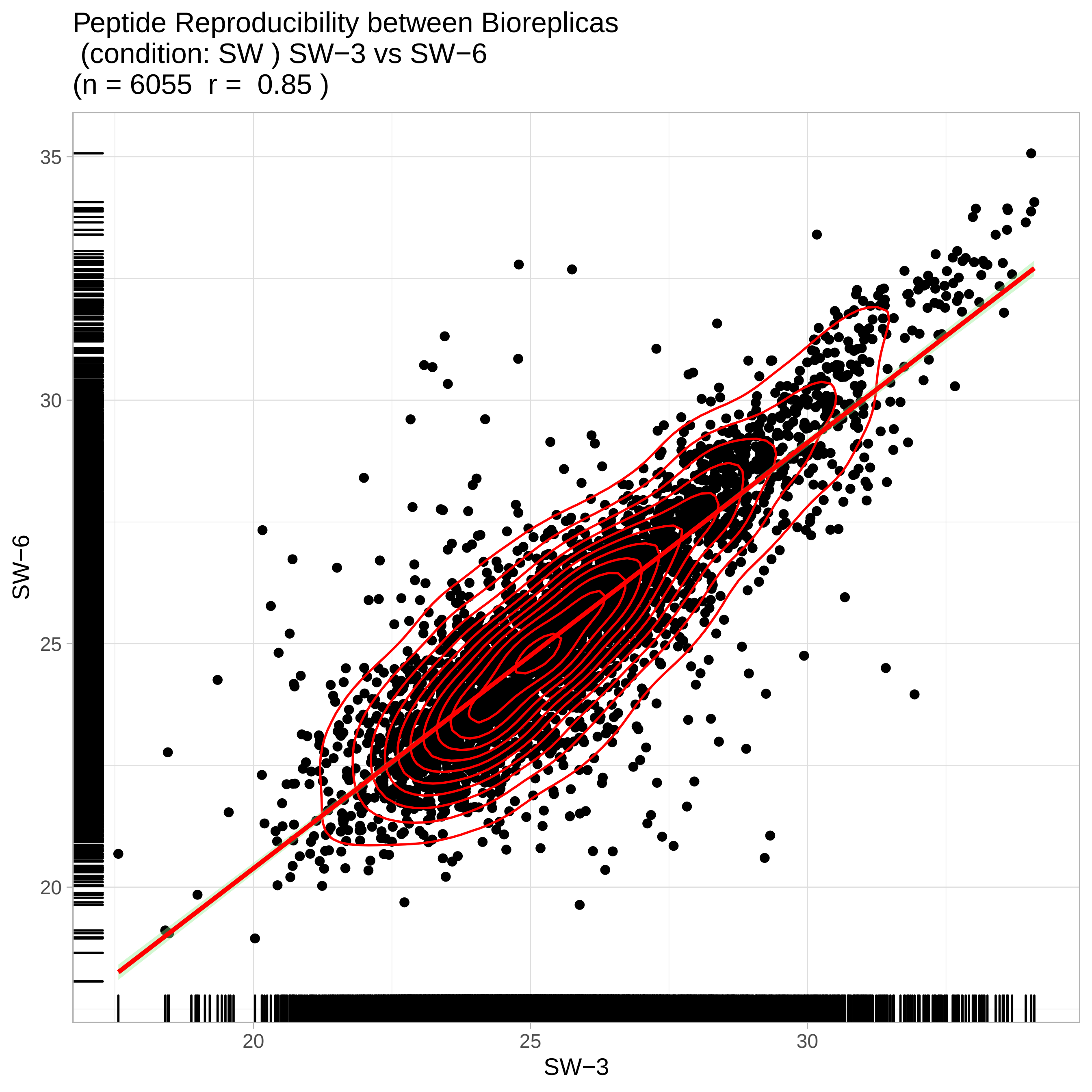
Peptide Reproducibility between Bioreplicas
(condition: SW ) SW−3 vs SW−6
(n = 6055 r = 0.85 )
SW−6
SW−3
35
30
25
20
20
25
30

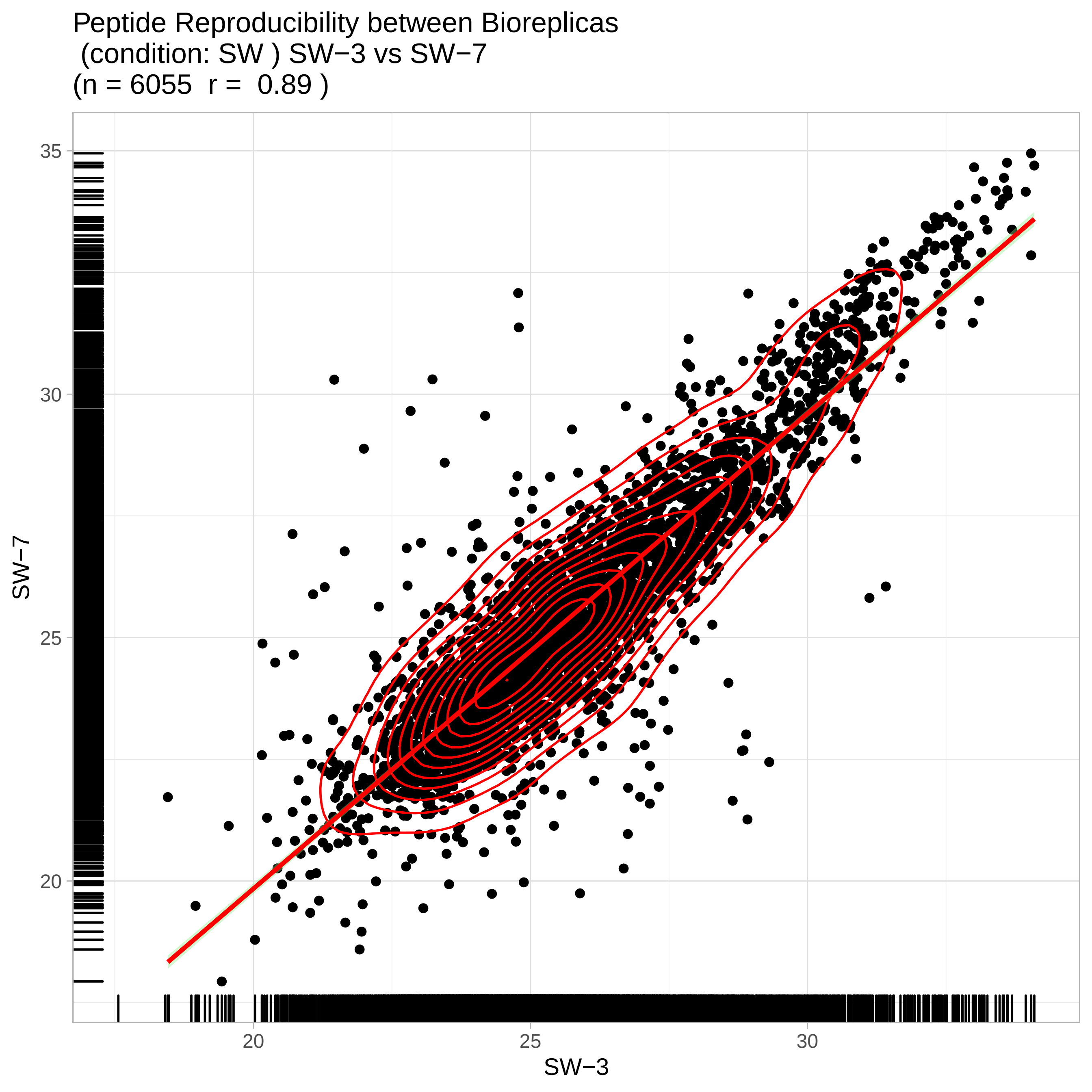
Peptide Reproducibility between Bioreplicas
(condition: SW ) SW−3 vs SW−7
(n = 6055 r = 0.89 )
35
30
25
20
SW−7
20
25
30
SW−3

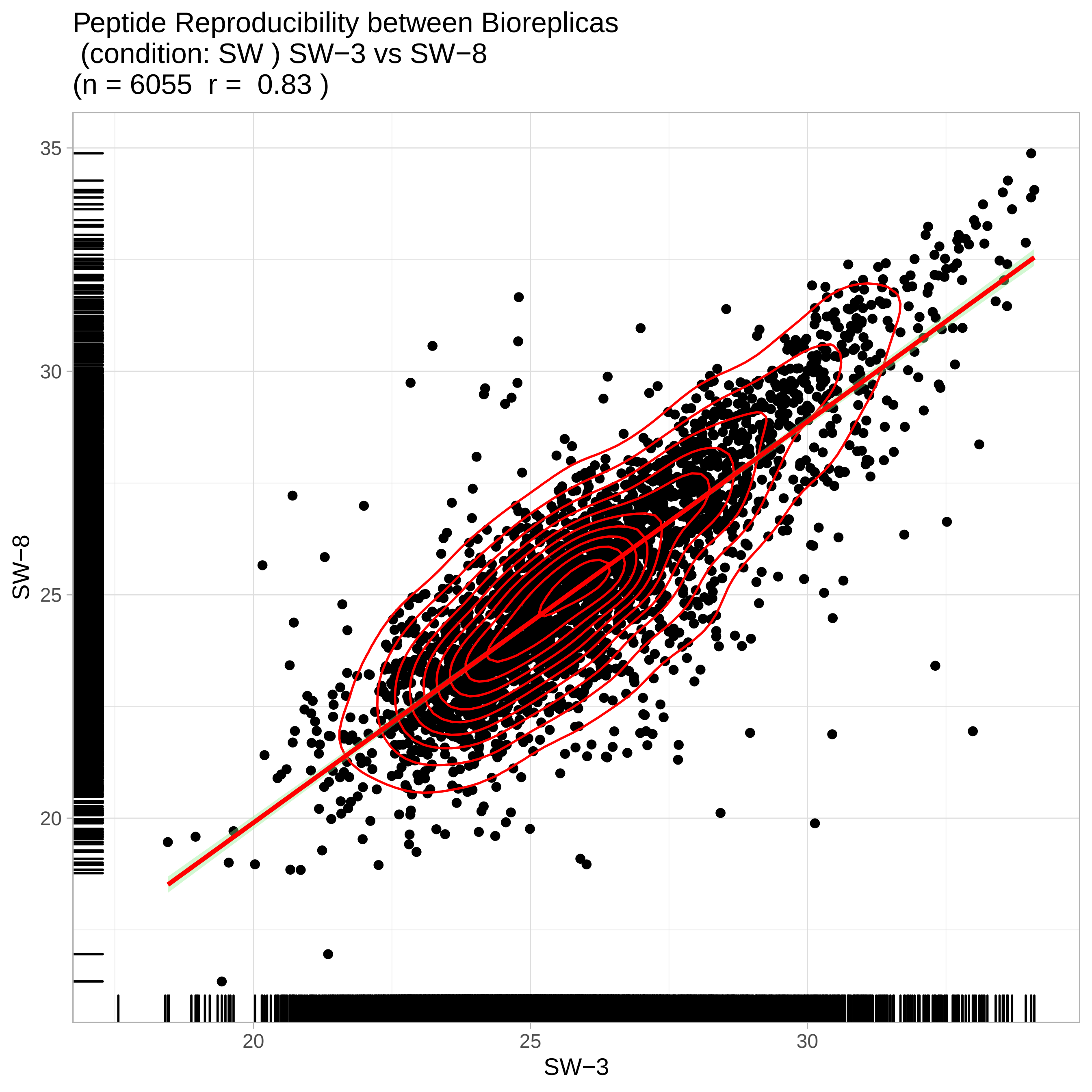

Peptide Reproducibility between Bioreplicas
(condition: SW ) SW−3 vs SW−8
(n = 6055 r = 0.83 )
SW−8
35
30
25
20
20
25
30
SW−3

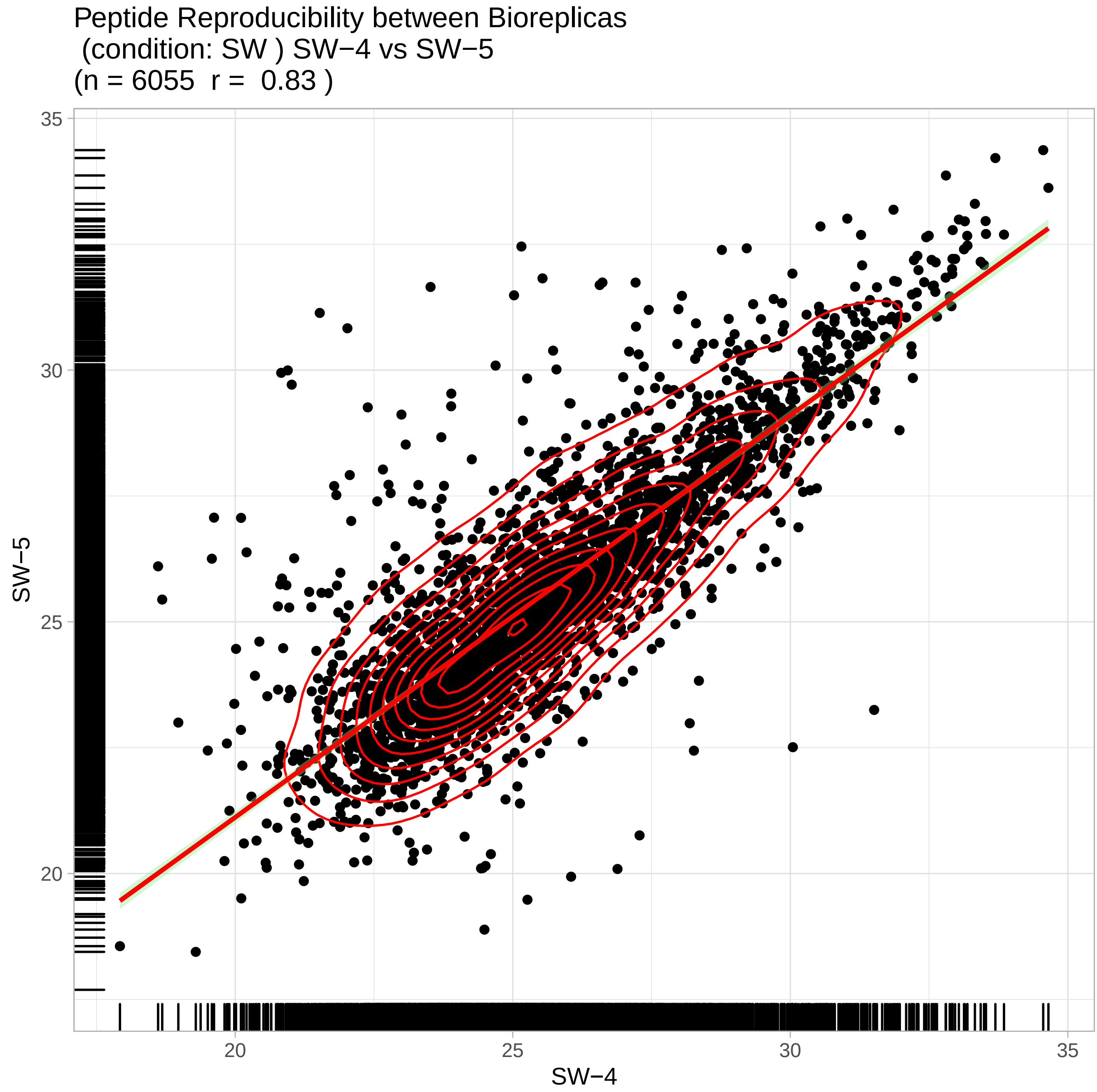
Peptide Reproducibility between Bioreplicas
(condition: SW ) SW−4 vs SW−5
(n = 6055 r = 0.83 )
SW−5
SW−4
35
30
25
20
20
25
30
35

Peptide Reproducibility between Bioreplicas
(condition: SW ) SW−4 vs SW−6
(n = 6055 r = 0.78 )

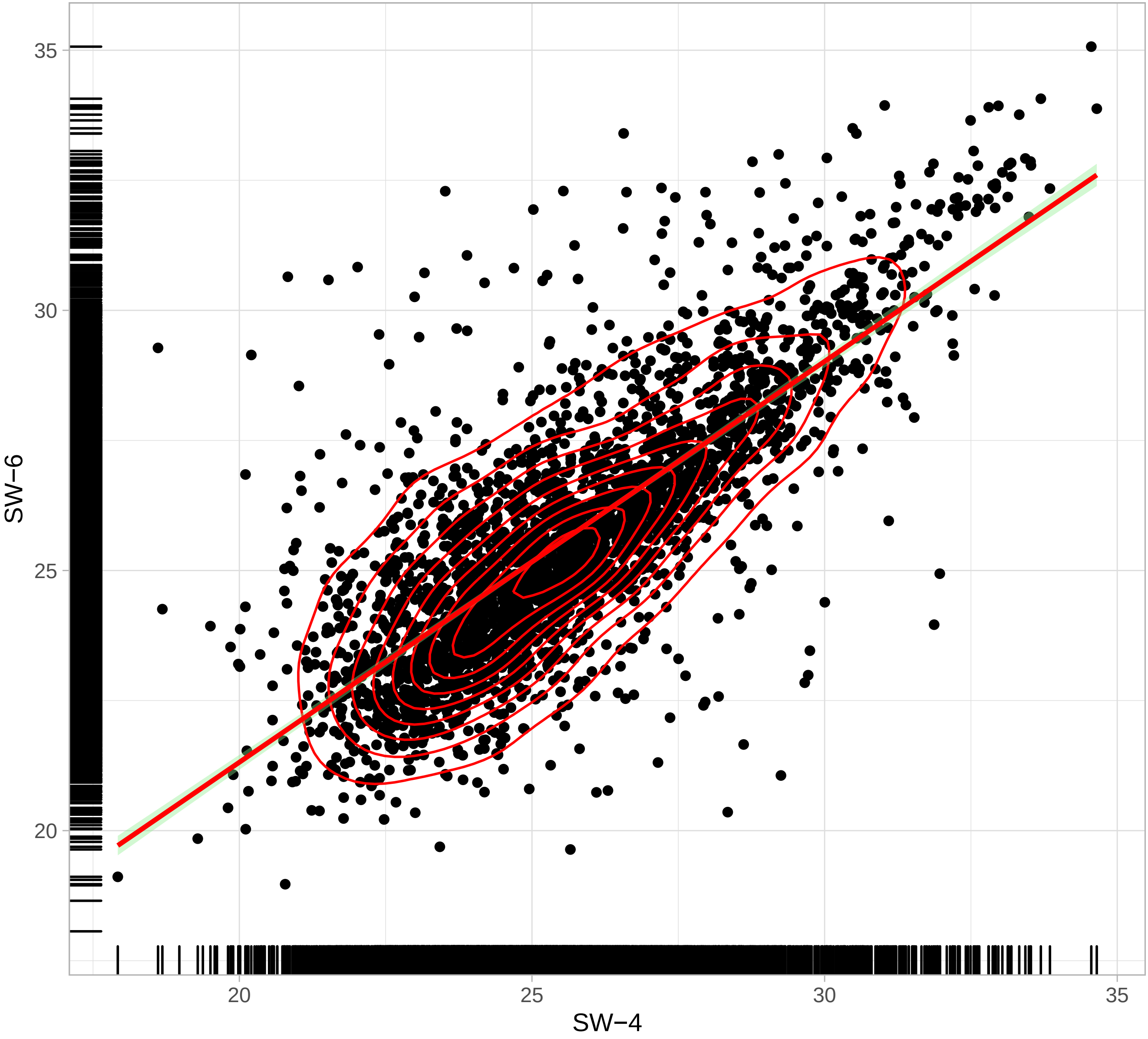

Peptide Reproducibility between Bioreplicas
(condition: SW ) SW−4 vs SW−7
(n = 6055  r =  0.78 )

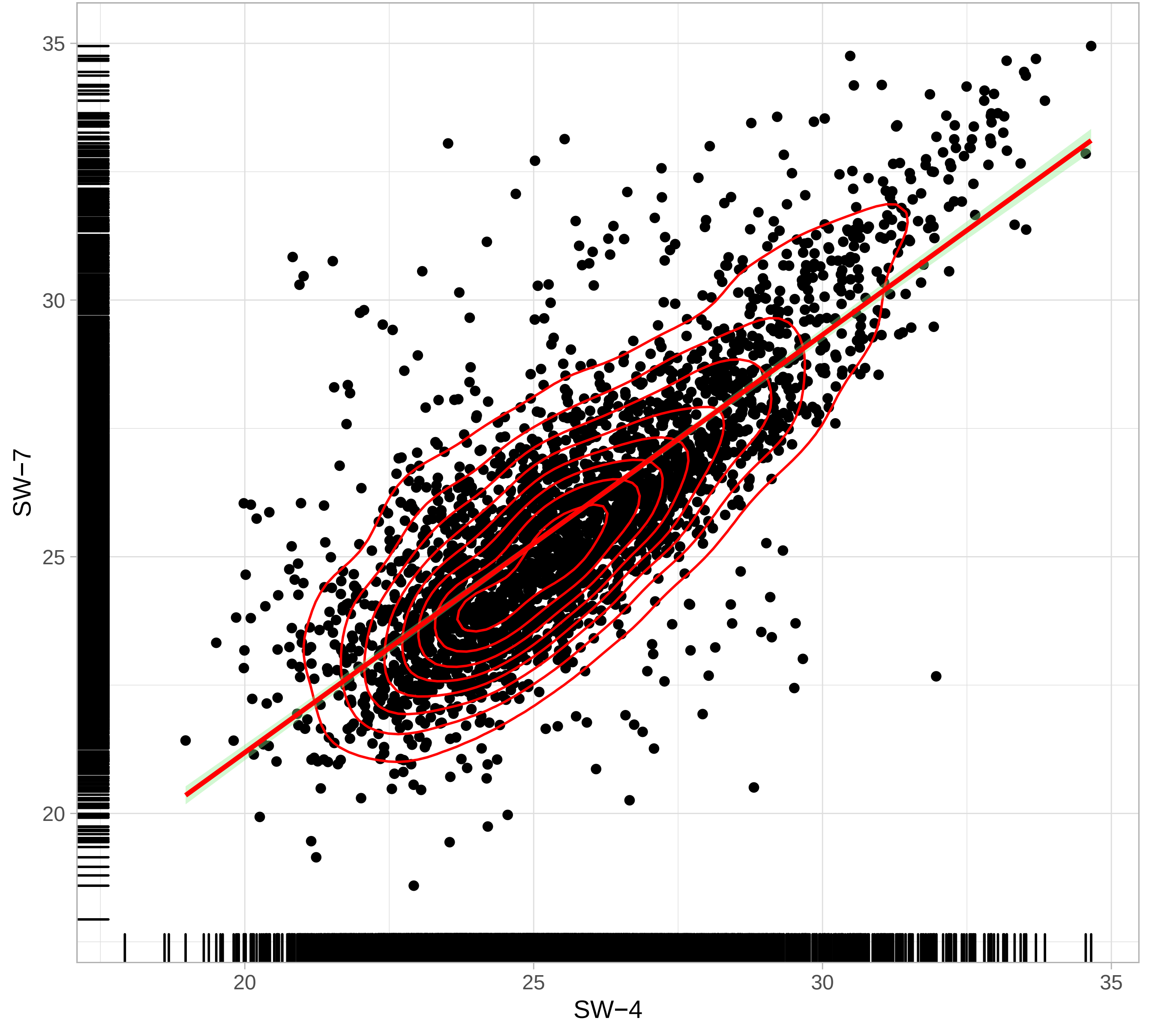

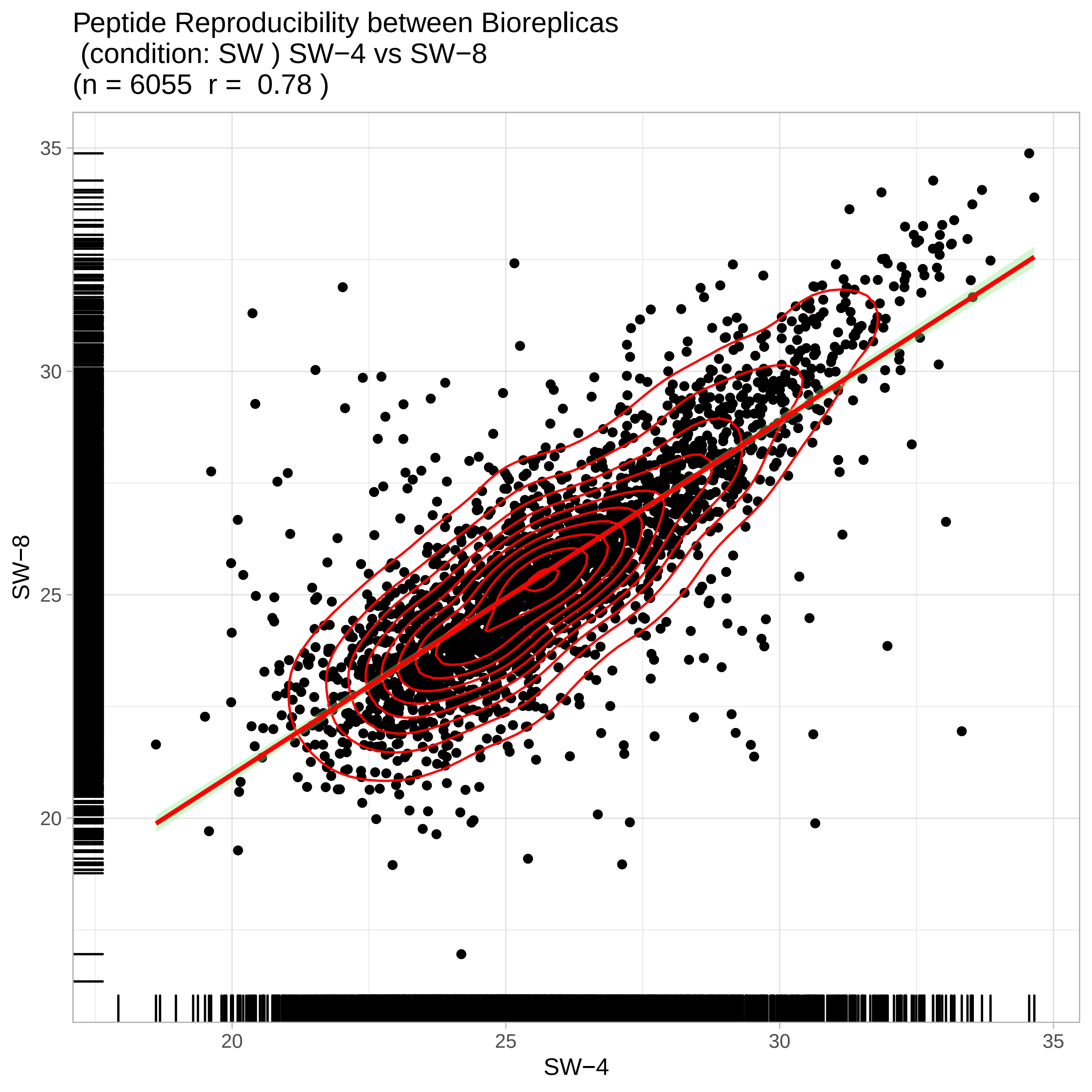

Peptide Reproducibility between Bioreplicas
(condition: SW ) SW−4 vs SW−8
(n = 6055 r = 0.78 )
35
30
25
20
SW−8
20
25
30
35
SW−4

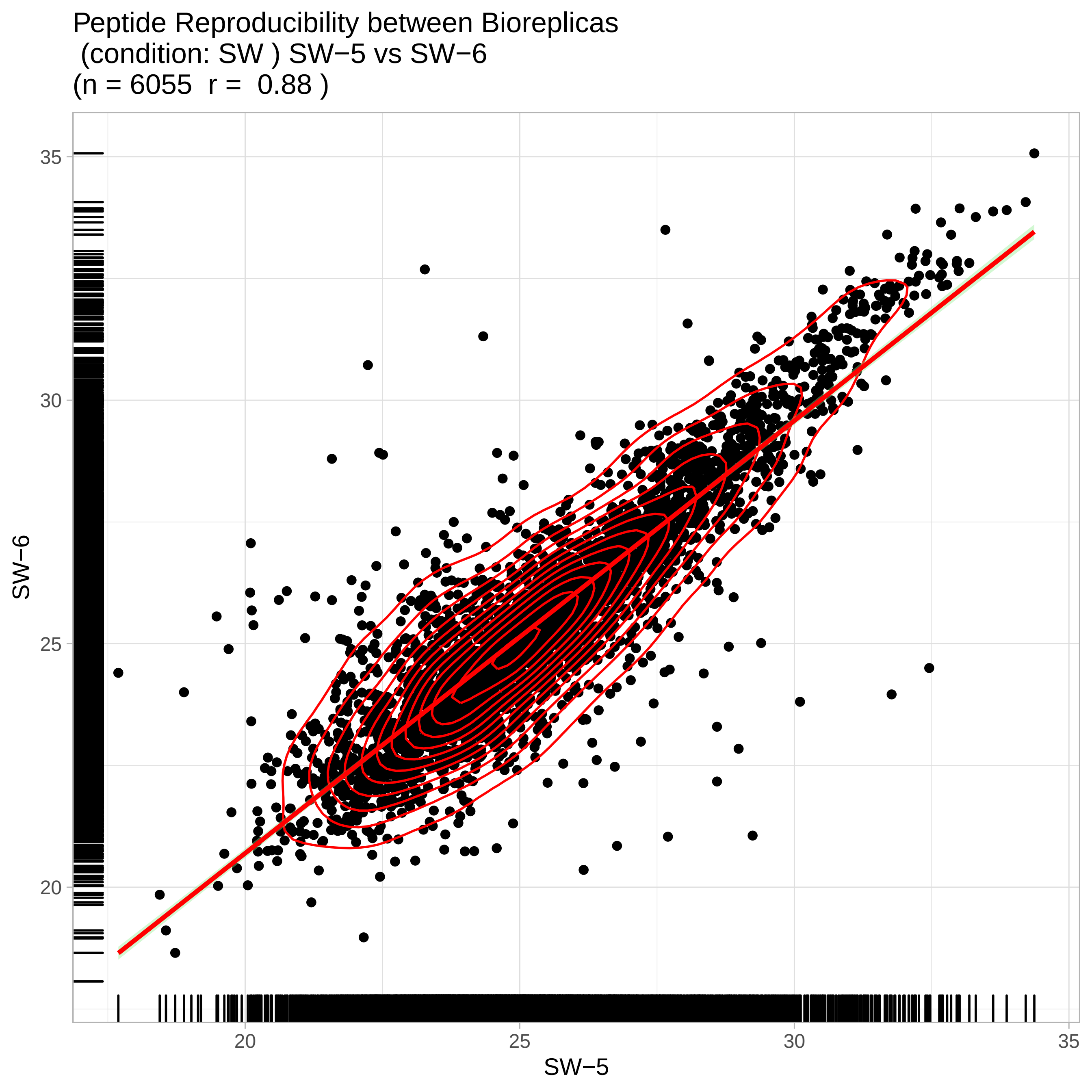
Peptide Reproducibility between Bioreplicas
(condition: SW ) SW−5 vs SW−6
(n = 6055 r = 0.88 )
SW−6
SW−5
35
30
25
20
20
25
30
35

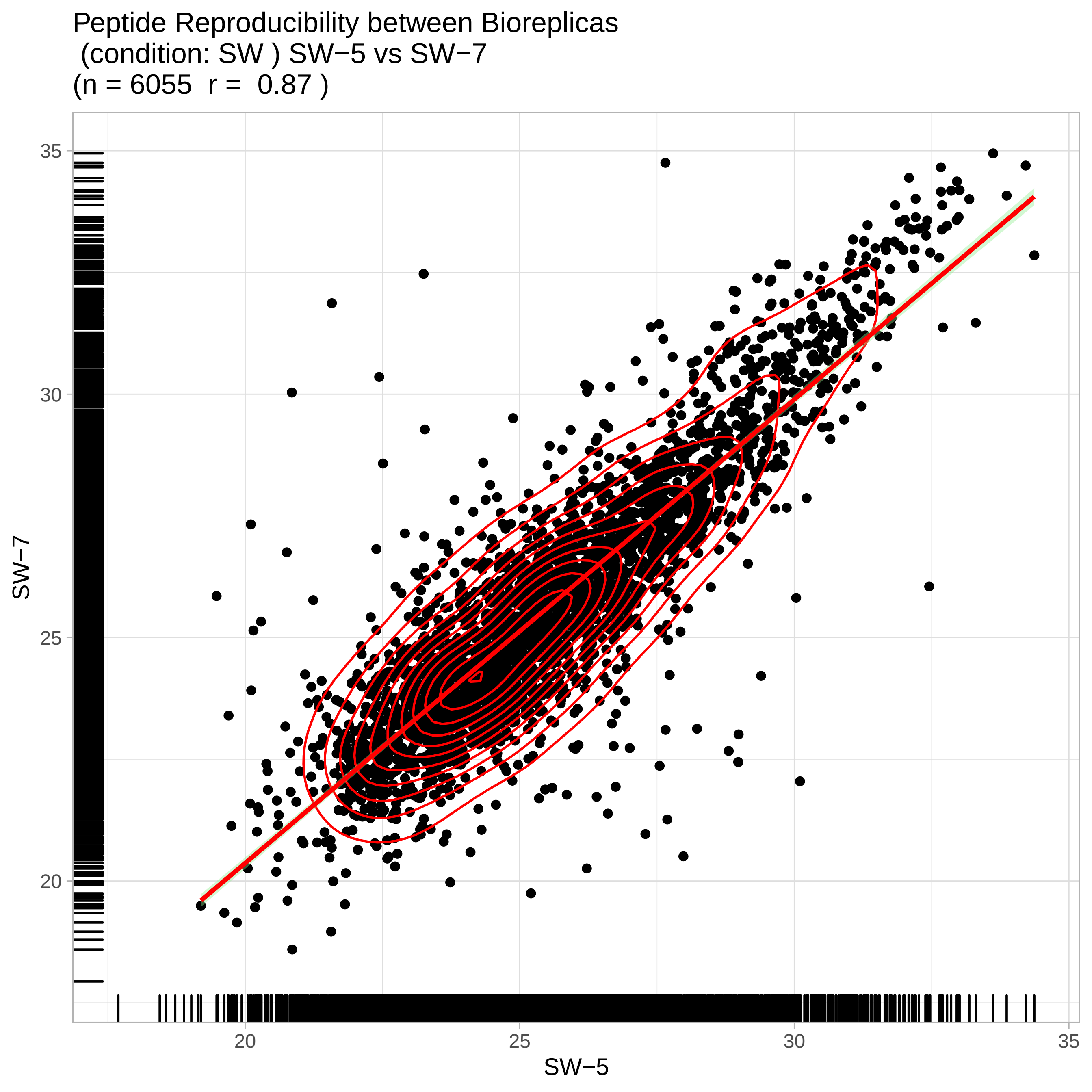
Peptide Reproducibility between Bioreplicas
(condition: SW ) SW−5 vs SW−7
(n = 6055 r = 0.87 )
SW−7
SW−5
35
30
25
20
20
25
30
35

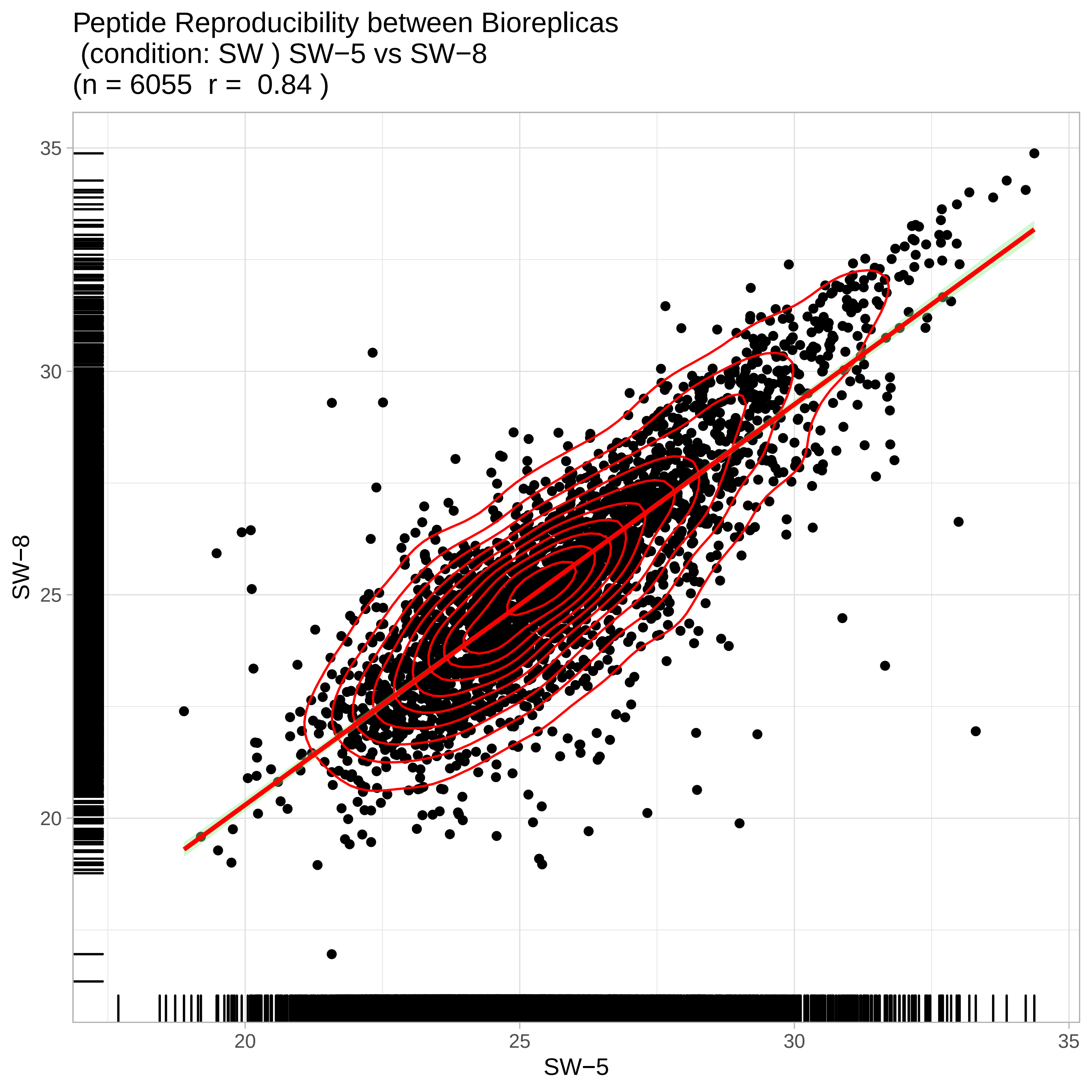

Peptide Reproducibility between Bioreplicas
(condition: SW ) SW−5 vs SW−8
(n = 6055 r = 0.84 )
SW−8
SW−5
35
30
25
20
20
25
30
35

Peptide Reproducibility between Bioreplicas
(condition: SW ) SW−6 vs SW−7
(n = 6055 r = 0.85 )

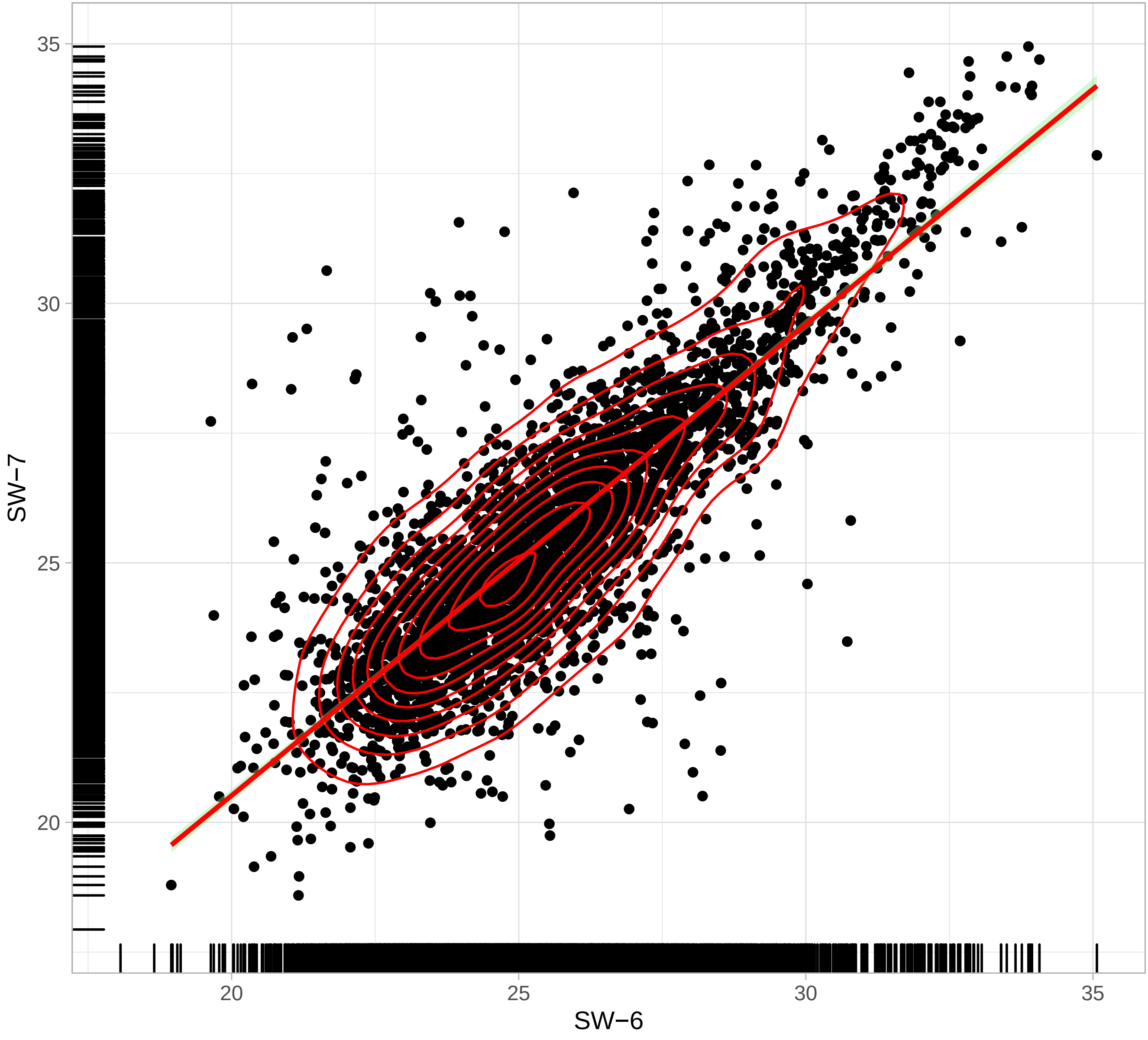

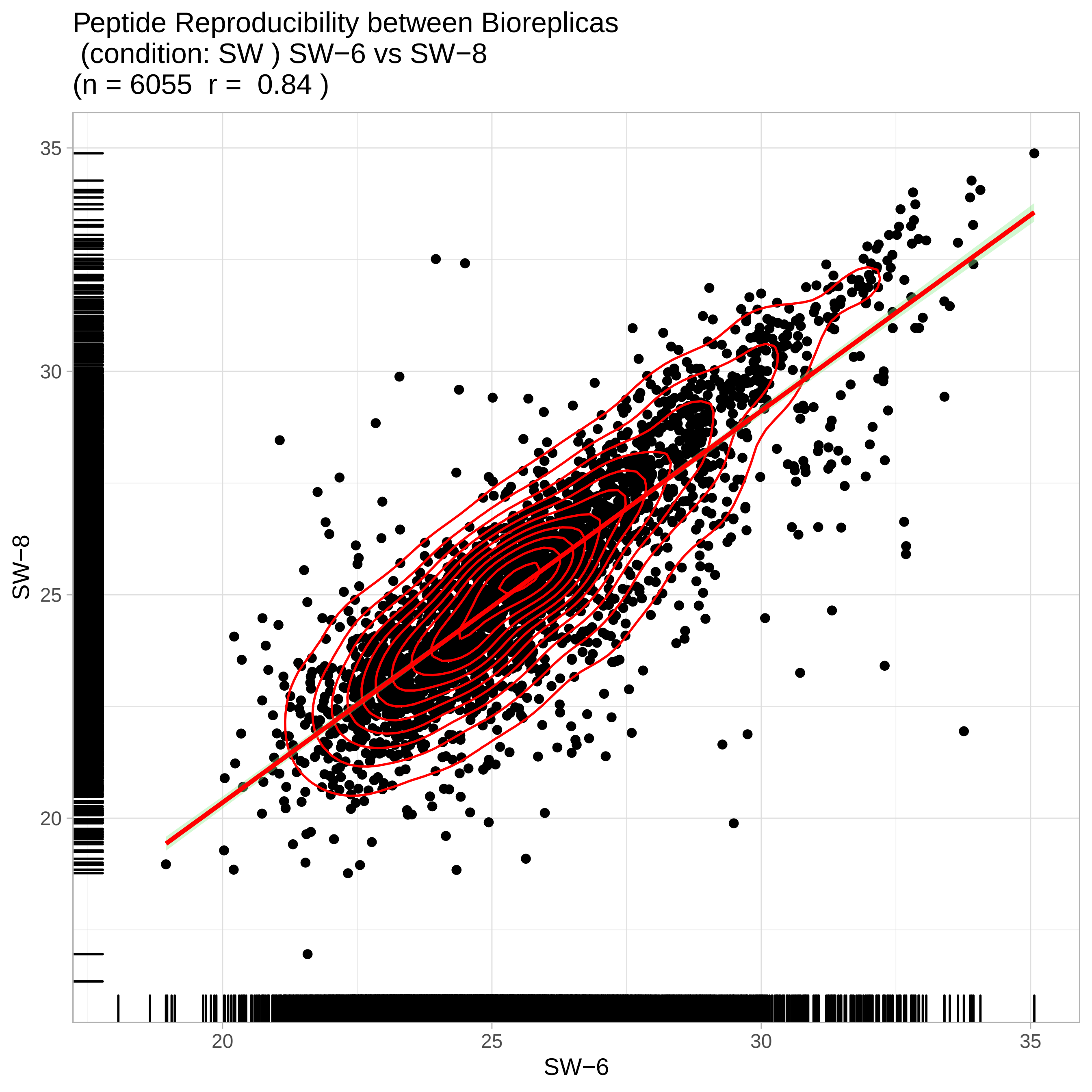
Peptide Reproducibility between Bioreplicas
(condition: SW ) SW−6 vs SW−8
(n = 6055 r = 0.84 )
SW−8
35
30
25
20
20
25
30
35
SW−6

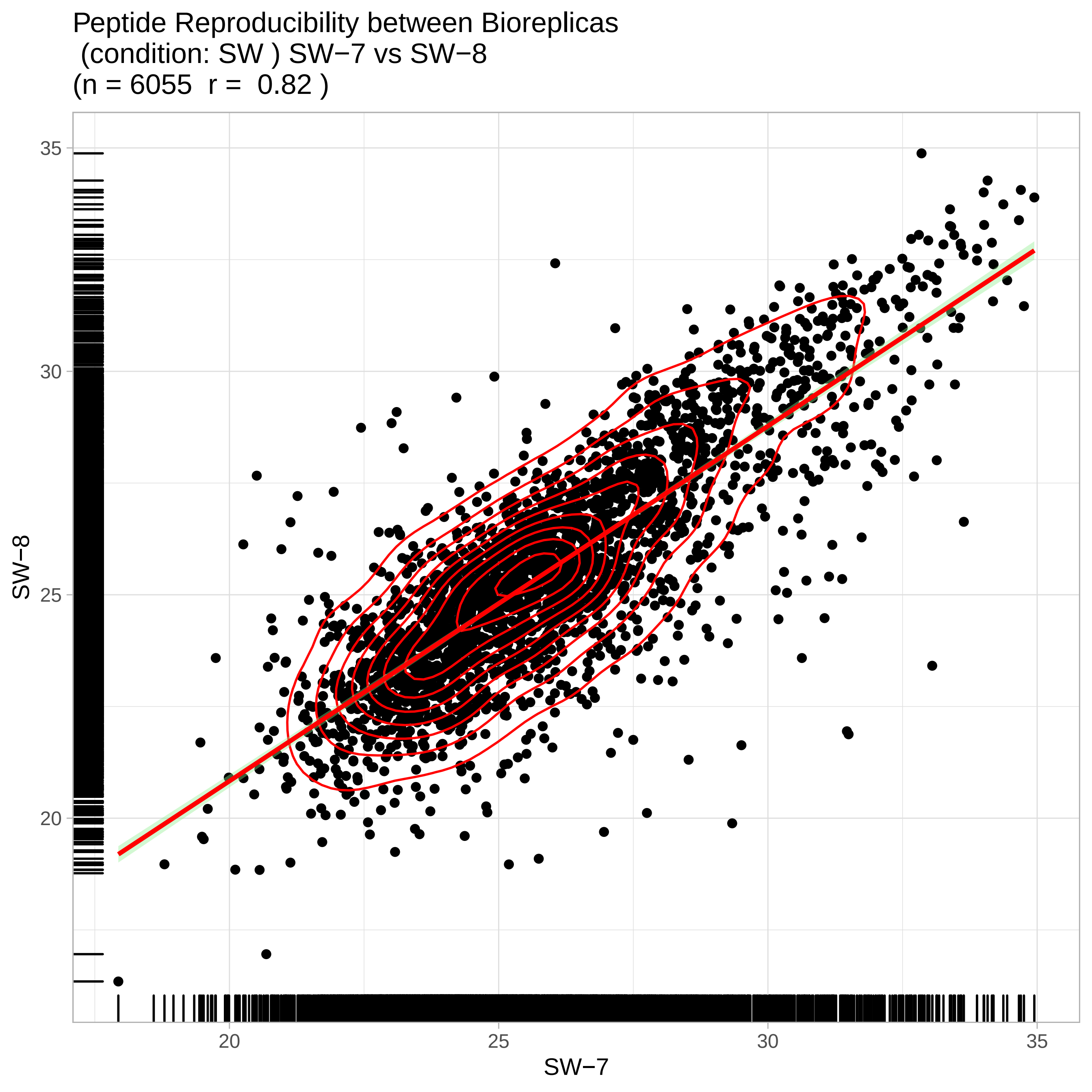

Peptide Reproducibility between Bioreplicas
(condition: SW ) SW−7 vs SW−8
(n = 6055 r = 0.82 )
35
30
25
20
SW−8
20
25
30
35
SW−7
