## Supplemental Information 3 for "Abiotic drivers of protein abundance variation among natural populations": qcPlots_evidence.qcplot.intensityStats.pdf

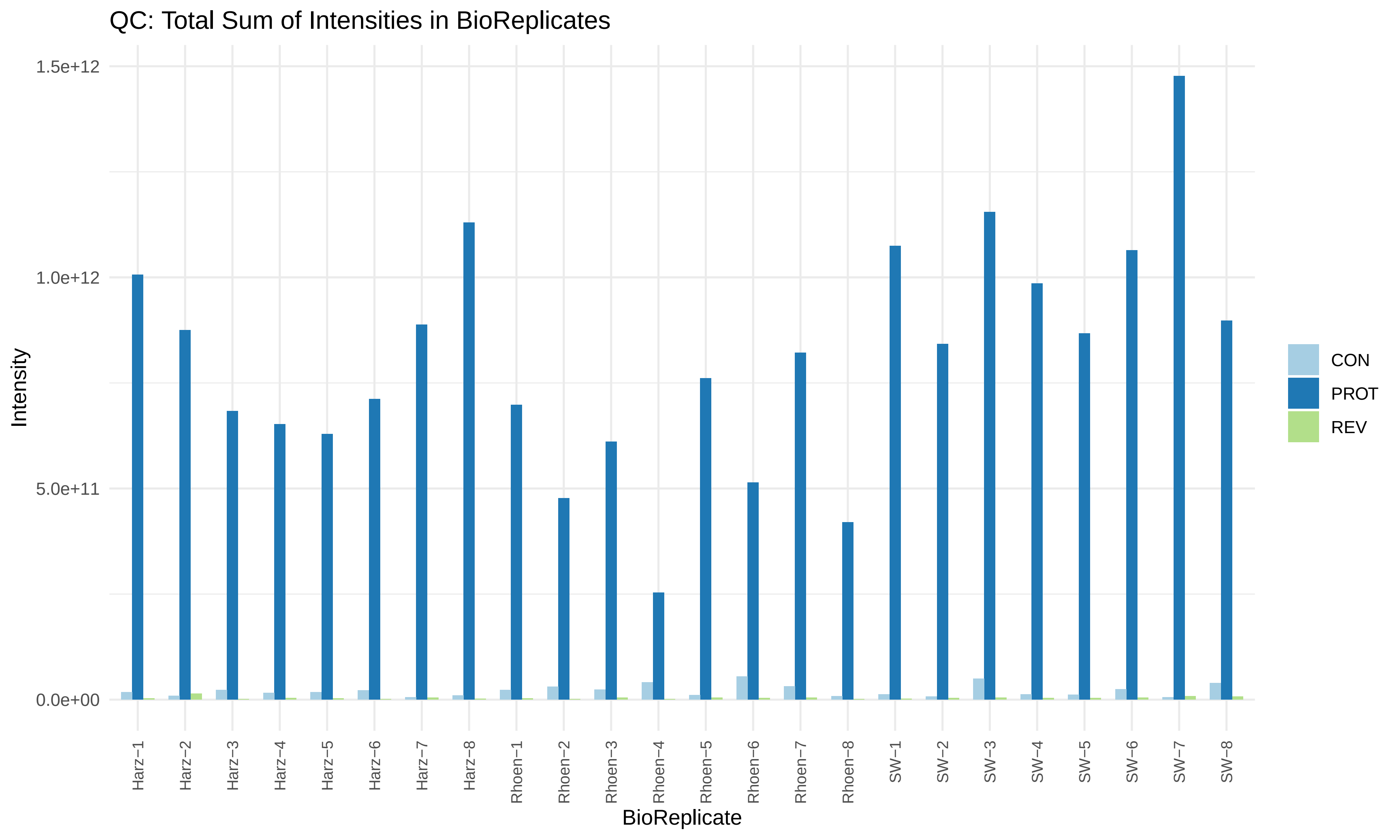
QC: Total Sum of Intensities in BioReplicates
1.5e+12
1.0e+12
5.0e+11
0.0e+00
Intensity
Harz−1
Harz−2
Harz−3
Harz−4
Harz−5
Harz−6
Harz−7
Harz−8
Rhoen−1
Rhoen−2
Rhoen−3
Rhoen−4
Rhoen−5
Rhoen−6
Rhoen−7
Rhoen−8
SW−1
SW−2
SW−3
SW−4
SW−5
SW−6
SW−7
SW−8
BioReplicate
CON
PROT
REV

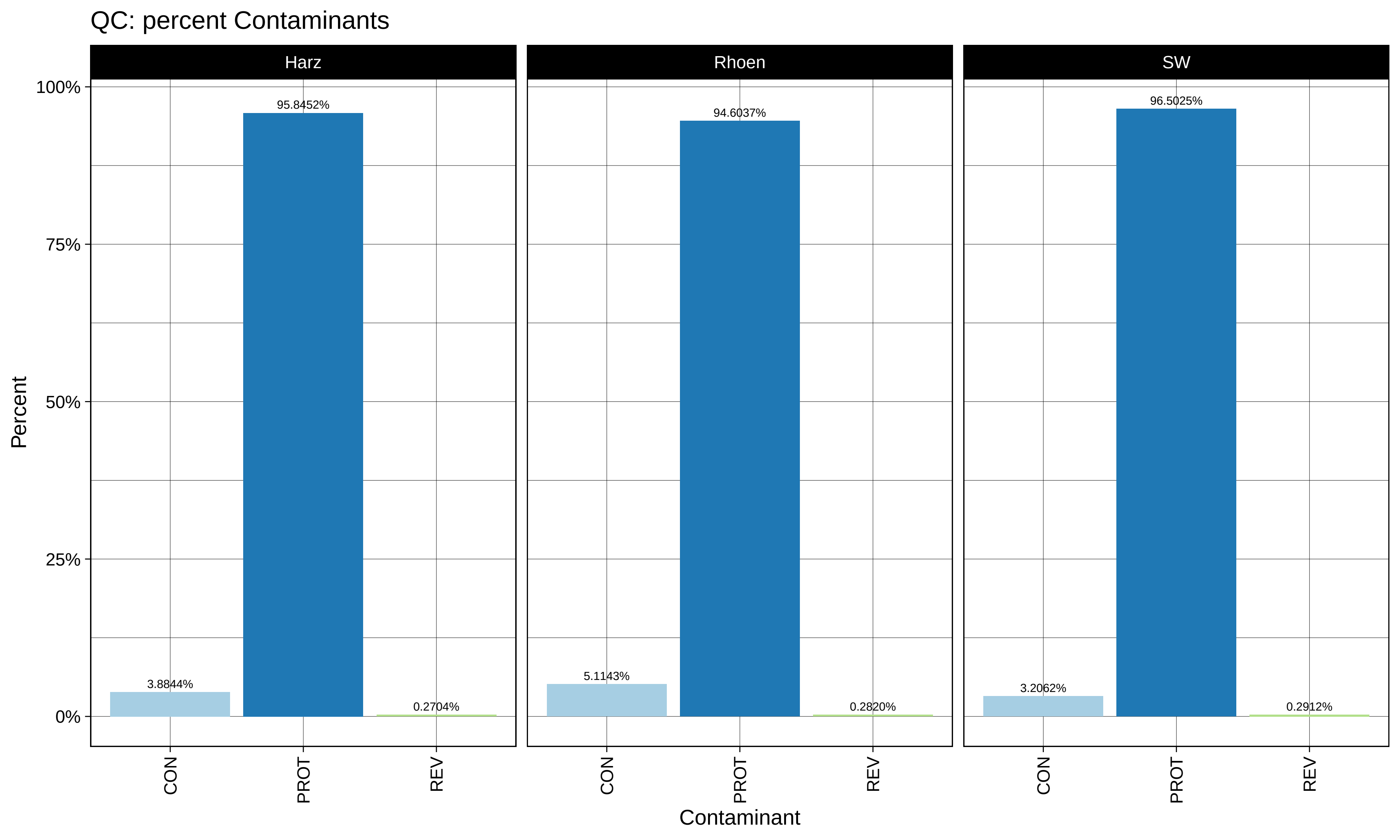
QC: percent Contaminants
Harz
Rhoen
SW
100%
75%
50%
25%
0%
Percent
95.8452%
3.8844%
0.2704%
94.6037%
5.1143%
0.2820%
96.5025%
3.2062%
0.2912%
CON
PROT
REV
Contaminant

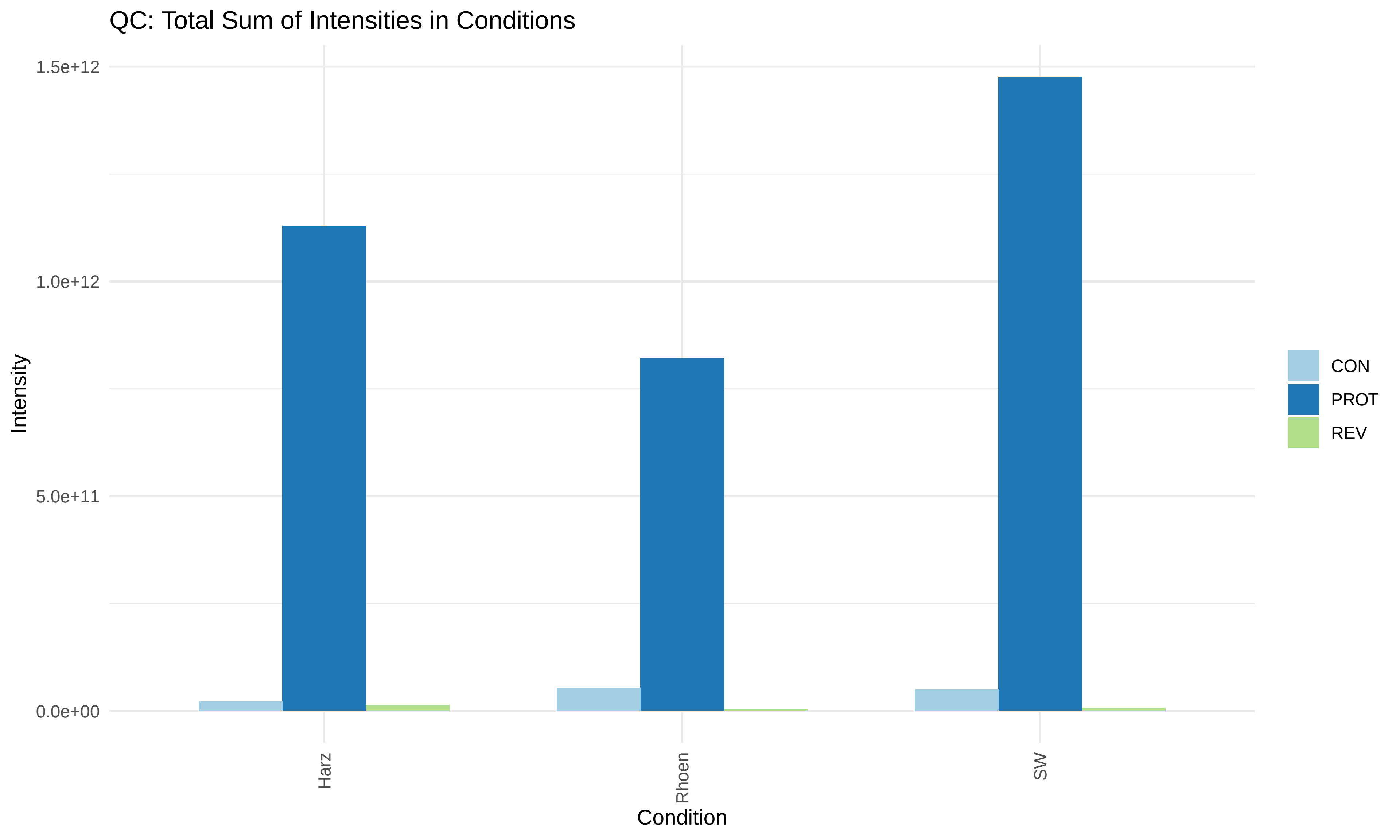
QC: Total Sum of Intensities in Conditions
1.5e+12
1.0e+12
5.0e+11
0.0e+00
Intensity
Harz
Rhoen
SW
Condition
CON
PROT
REV

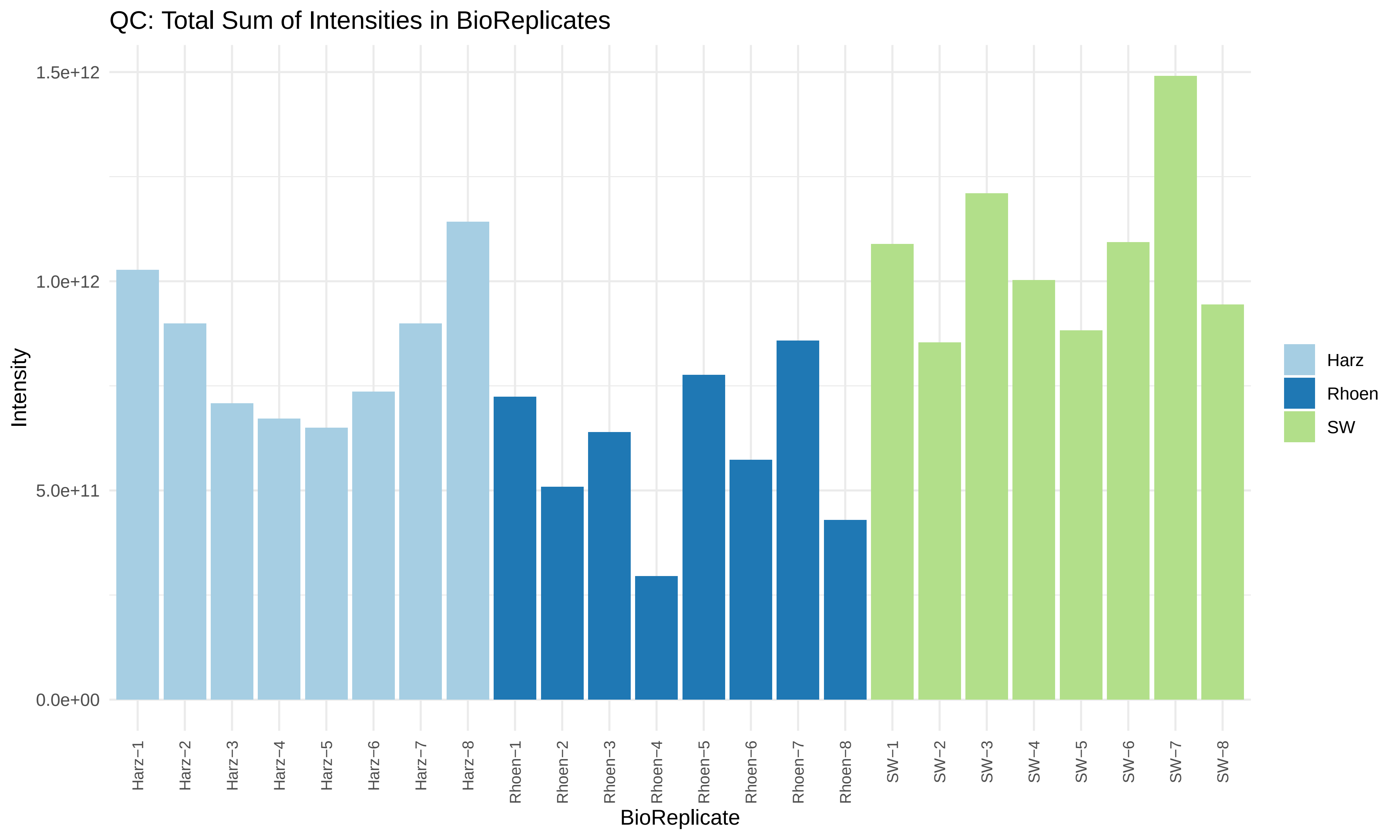
QC: Total Sum of Intensities in BioReplicates
Intensity
1.5e+12
1.0e+12
5.0e+11
0.0e+00
Harz−1
Harz−2
Harz−3
Harz−4
Harz−5
Harz−6
Harz−7
Harz−8
Rhoen−1
Rhoen−2
Rhoen−3
Rhoen−4
Rhoen−5
Rhoen−6
Rhoen−7
Rhoen−8
SW−1
SW−2
SW−3
SW−4
SW−5
SW−6
SW−7
SW−8
BioReplicate
Harz
Rhoen
SW

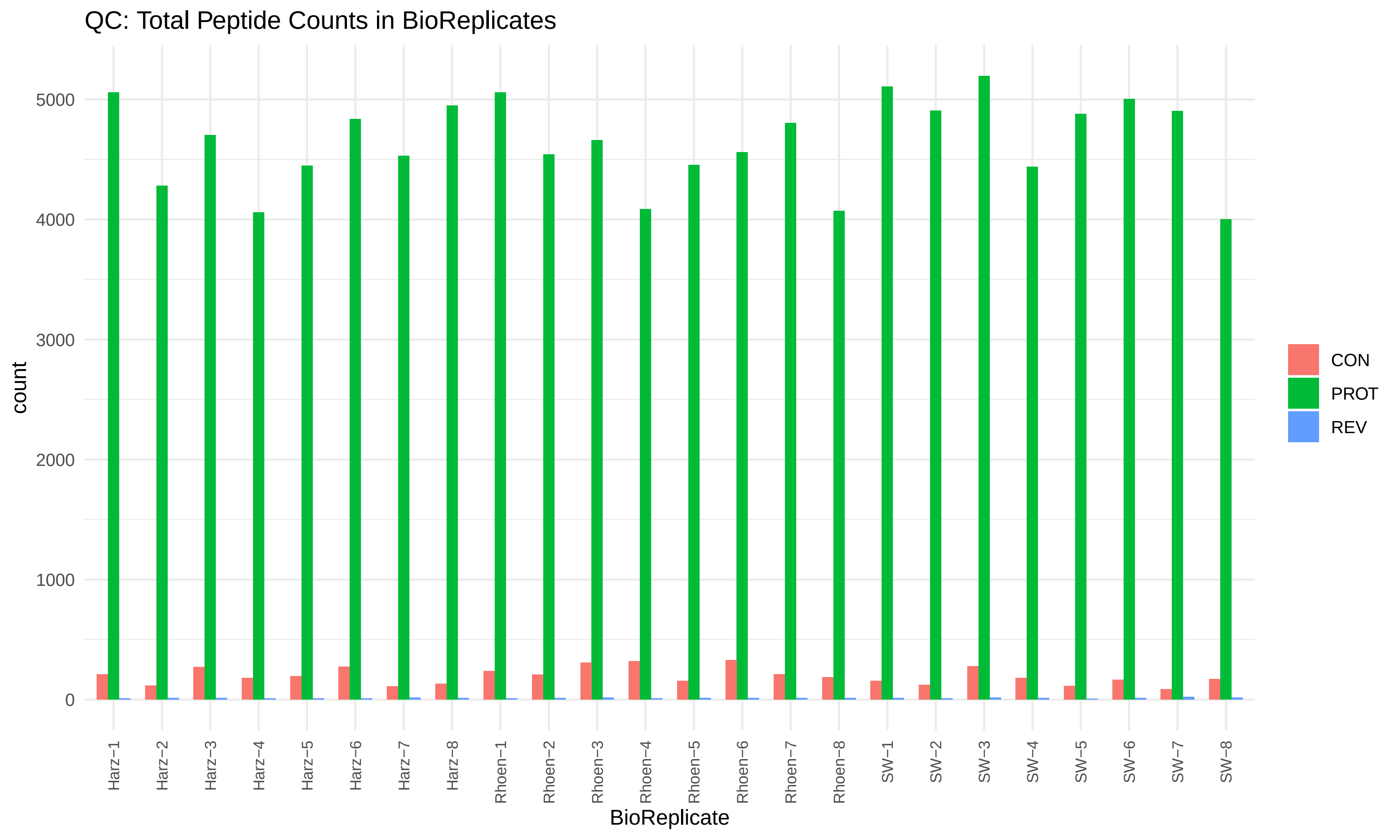
QC: Total Peptide Counts in BioReplicates
count
0
1000
2000
3000
4000
5000
Harz−1
Harz−2
Harz−3
Harz−4
Harz−5
Harz−6
Harz−7
Harz−8
Rhoen−1
Rhoen−2
Rhoen−3
Rhoen−4
Rhoen−5
Rhoen−6
Rhoen−7
Rhoen−8
SW−1
SW−2
SW−3
SW−4
SW−5
SW−6
SW−7
SW−8
BioReplicate
CON
PROT
REV

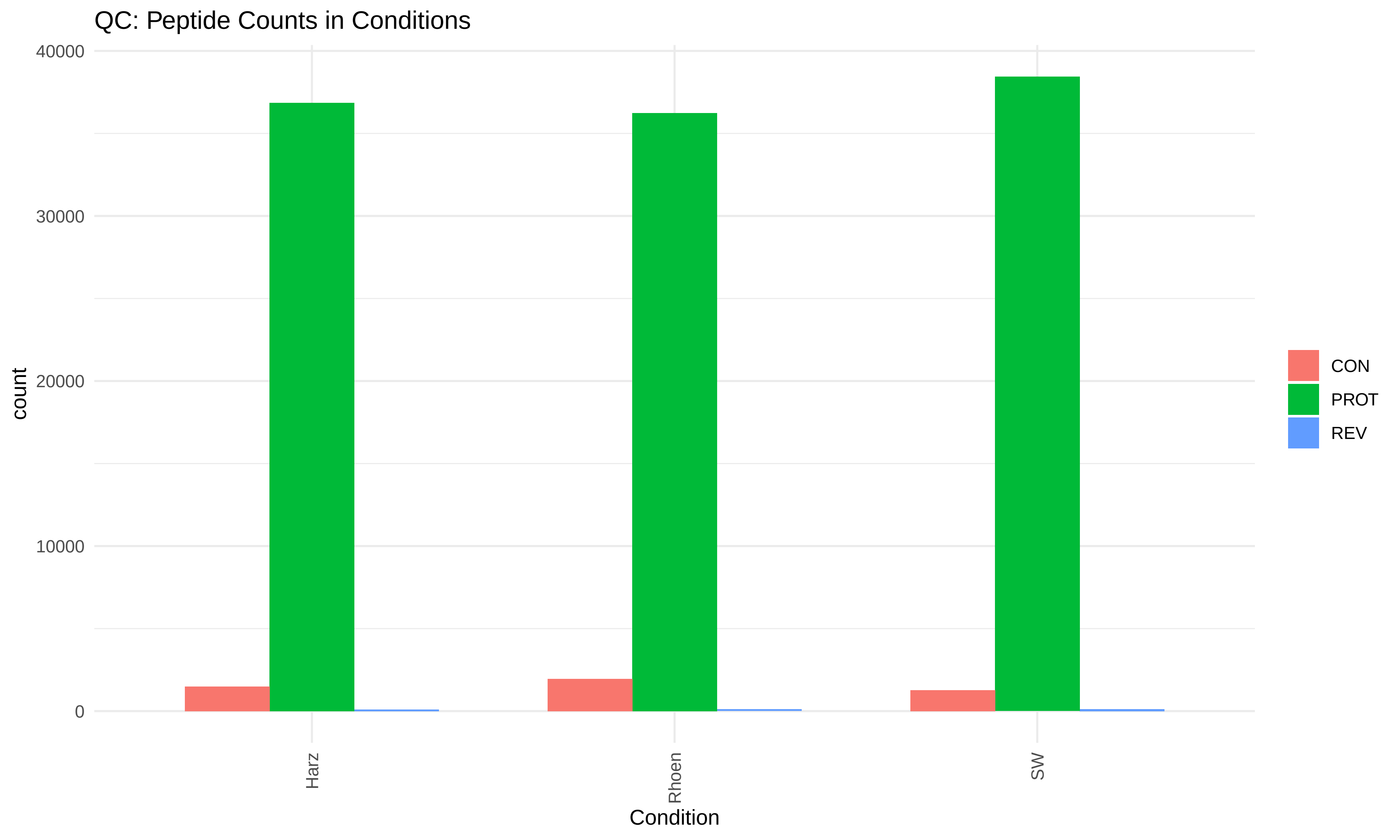
QC: Peptide Counts in Conditions
40000
30000
20000
10000
0
count
Harz
Rhoen
SW
Condition
CON
PROT
REV

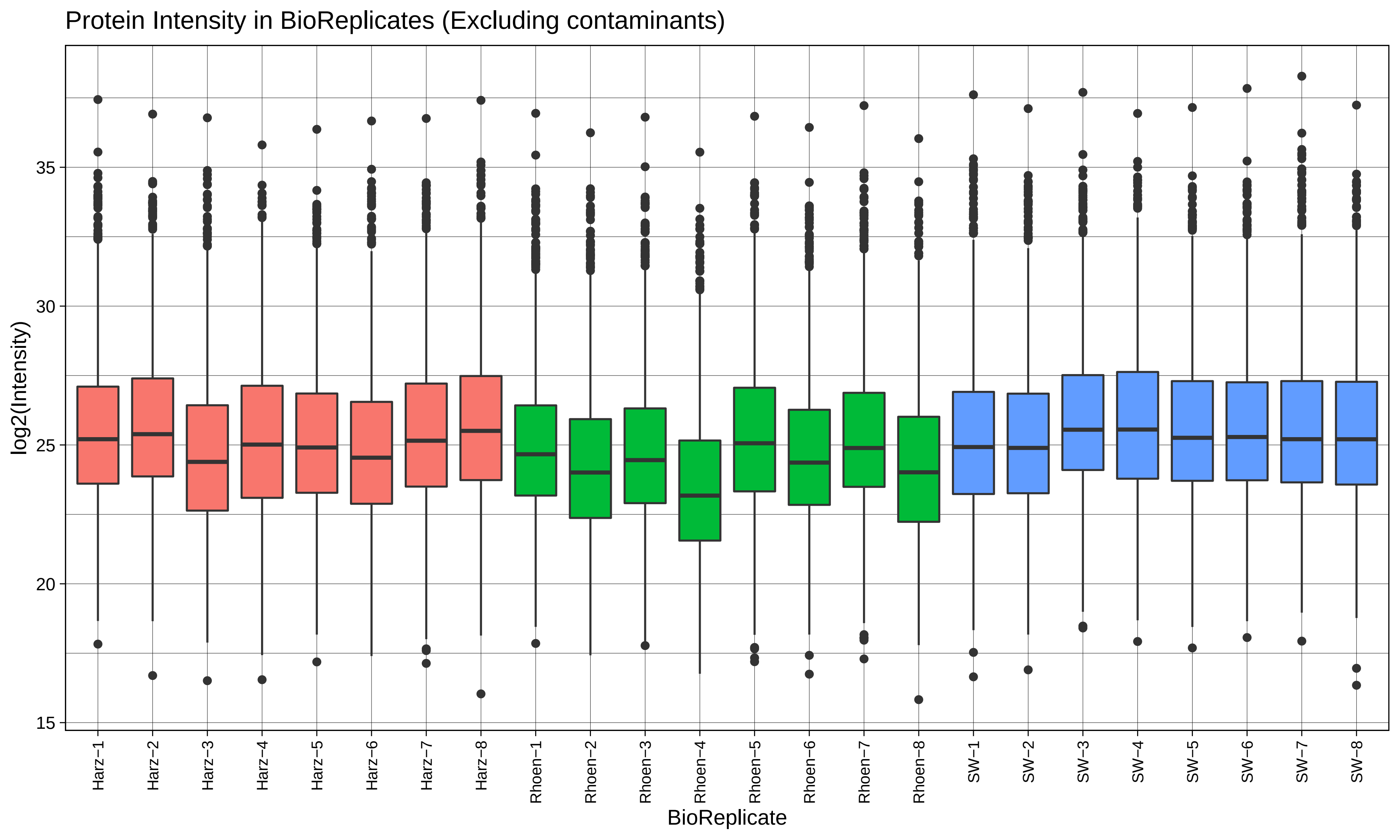
Protein Intensity in BioReplicates (Excluding contaminants)
log2(Intensity)
35
30
25
20
15
Harz−1
Harz−2
Harz−3
Harz−4
Harz−5
Harz−6
Harz−7
Harz−8
Rhoen−1
Rhoen−2
Rhoen−3
Rhoen−4
Rhoen−5
Rhoen−6
Rhoen−7
Rhoen−8
SW−1
SW−2
SW−3
SW−4
SW−5
SW−6
SW−7
SW−8
BioReplicate

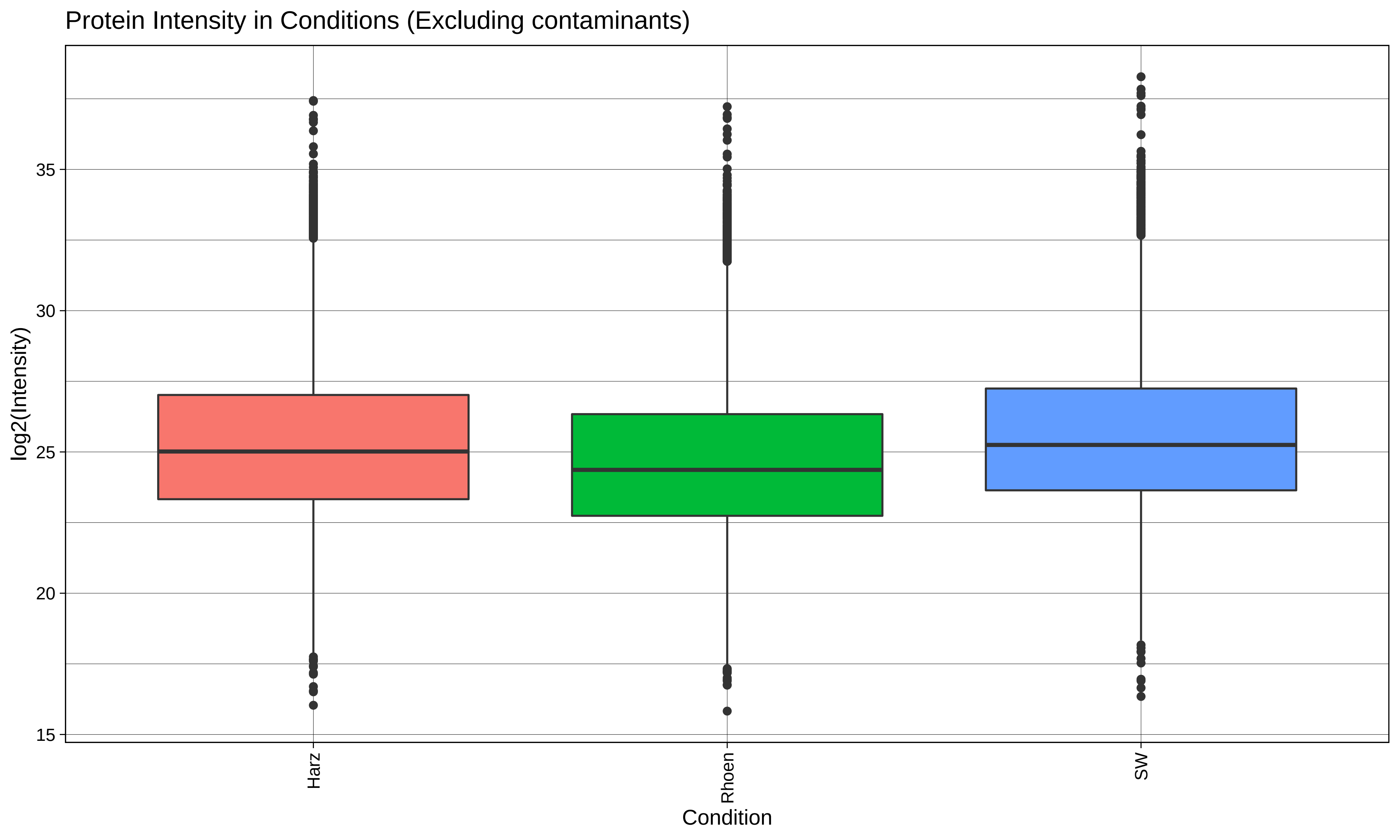
Protein Intensity in Conditions (Excluding contaminants)
log2(Intensity)
35
30
25
20
15
Harz
Rhoen
SW
Condition

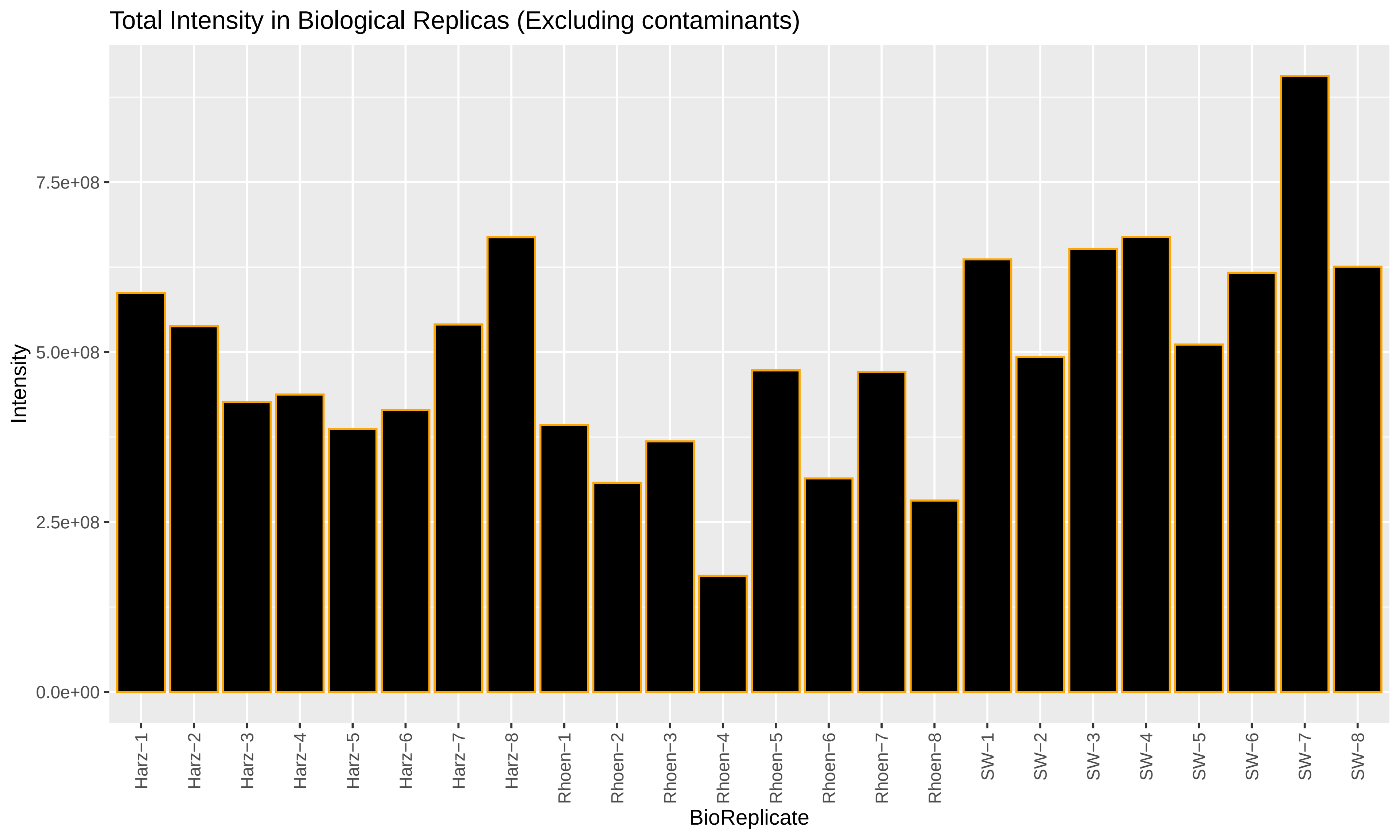
Total Intensity in Biological Replicas (Excluding contaminants)
Intensity
7.5e+08
5.0e+08
2.5e+08
0.0e+00
Harz−1
Harz−2
Harz−3
Harz−4
Harz−5
Harz−6
Harz−7
Harz−8
Rhoen−1
Rhoen−2
Rhoen−3
Rhoen−4
Rhoen−5
Rhoen−6
Rhoen−7
Rhoen−8
SW−1
SW−2
SW−3
SW−4
SW−5
SW−6
SW−7
SW−8
BioReplicate

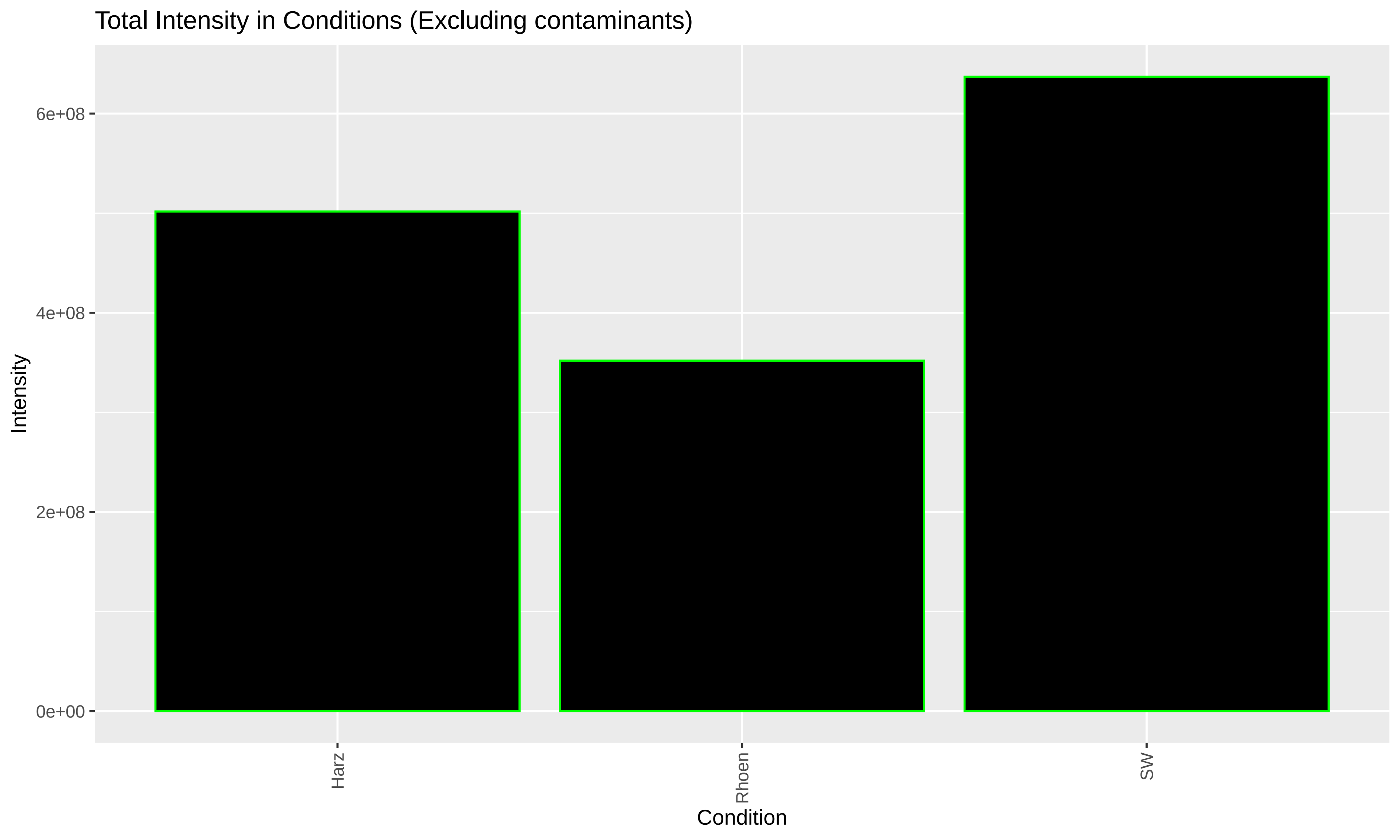
Total Intensity in Conditions (Excluding contaminants)
Intensity
6e+08
4e+08
2e+08
0e+00
Harz
Rhoen
SW
Condition

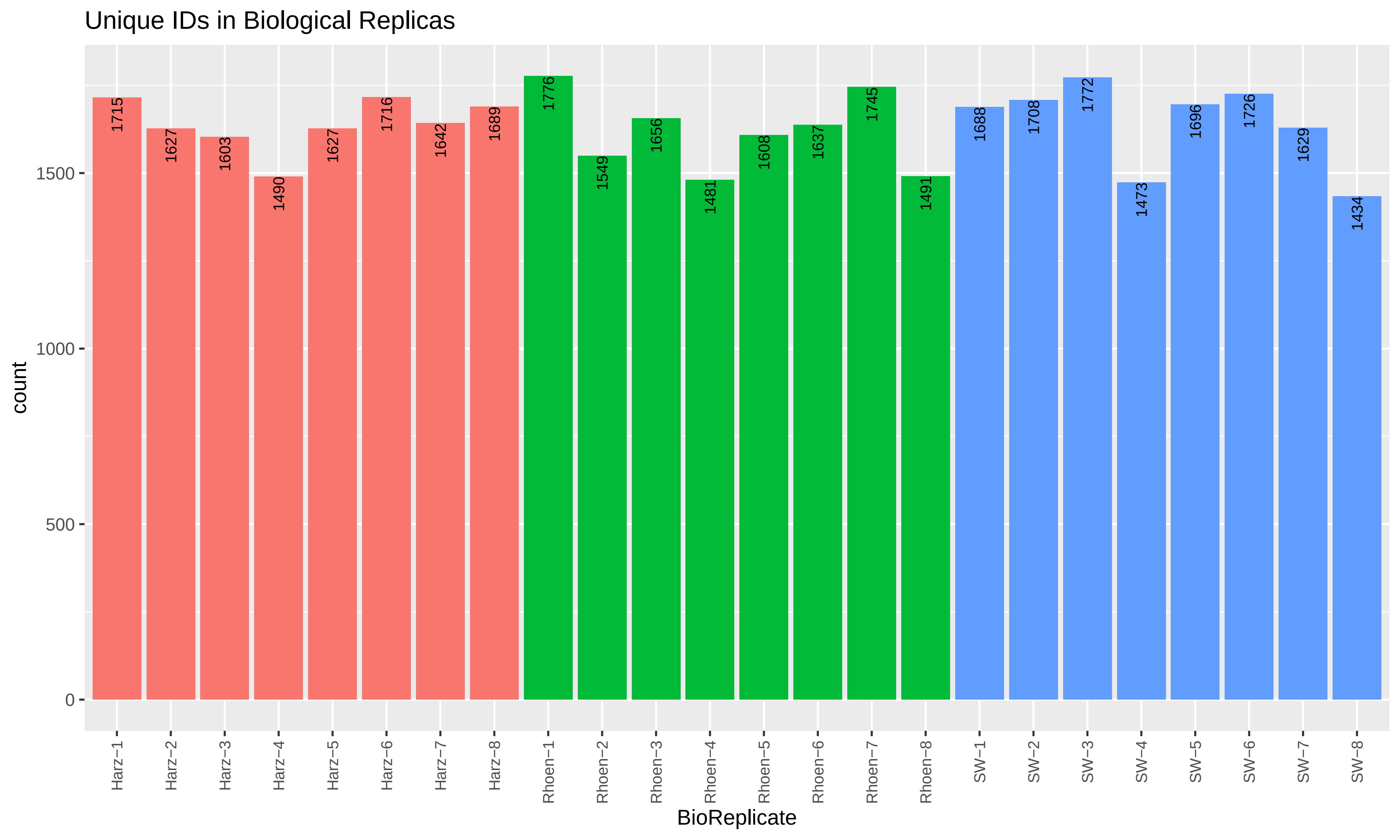

Unique IDs in Biological Replicas
count
0
500
1000
1500
1715
1627
1603
1490
1627
1716
1642
1689
1776
1549
1656
1481
1608
1637
1745
1491
1688
1708
1772
1473
1696
1726
1629
1434
Harz−1
Harz−2
Harz−3
Harz−4
Harz−5
Harz−6
Harz−7
Harz−8
Rhoen−1
Rhoen−2
Rhoen−3
Rhoen−4
Rhoen−5
Rhoen−6
Rhoen−7
Rhoen−8
SW−1
SW−2
SW−3
SW−4
SW−5
SW−6
SW−7
SW−8
BioReplicate

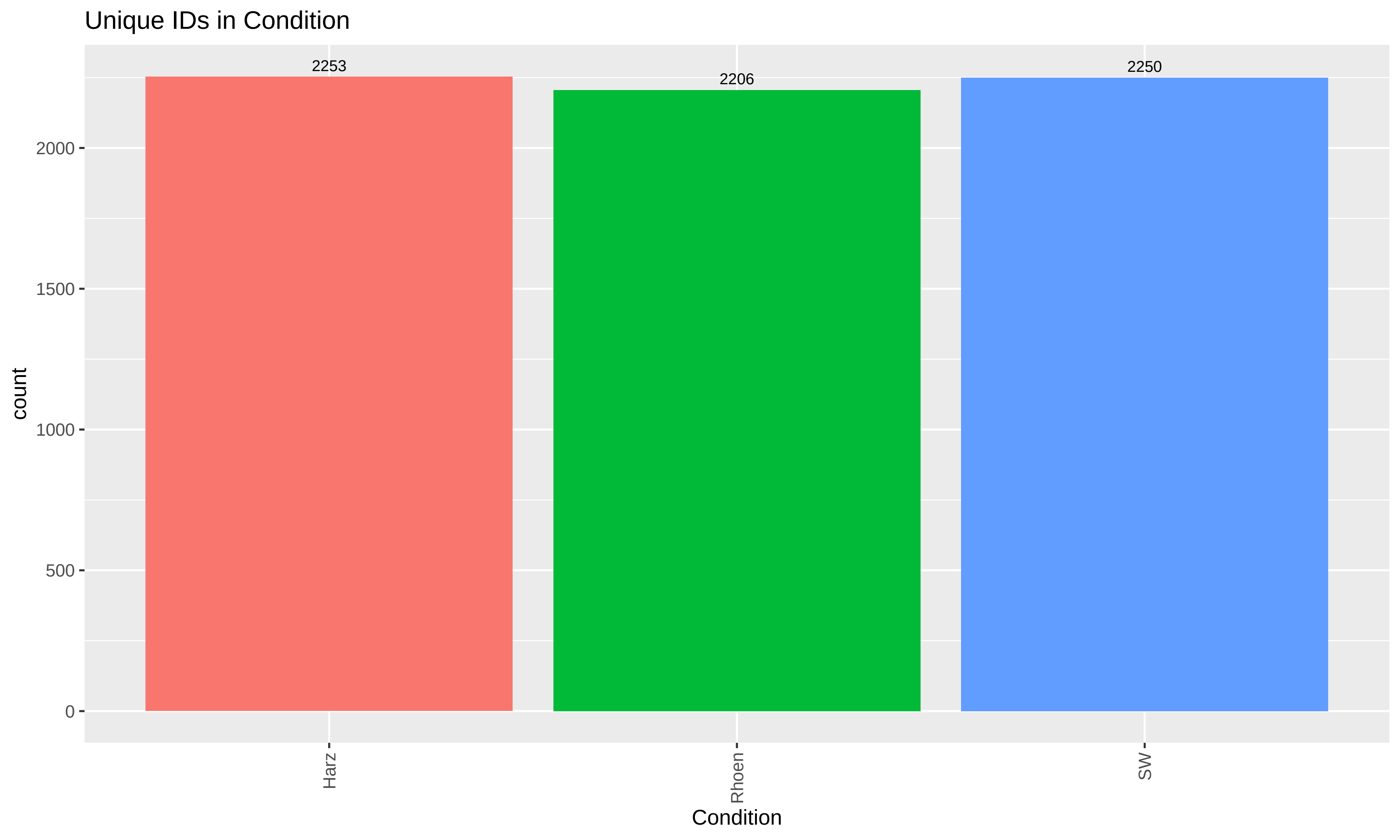

Unique IDs in Condition
2253
2206
2250
2000
1500
1000
500
0
count
Harz
Rhoen
SW
Condition
