## Supplementary figures and images for "Abiotic drivers of protein abundance variation among natural populations"

### QC-IntCorrelation.pdf

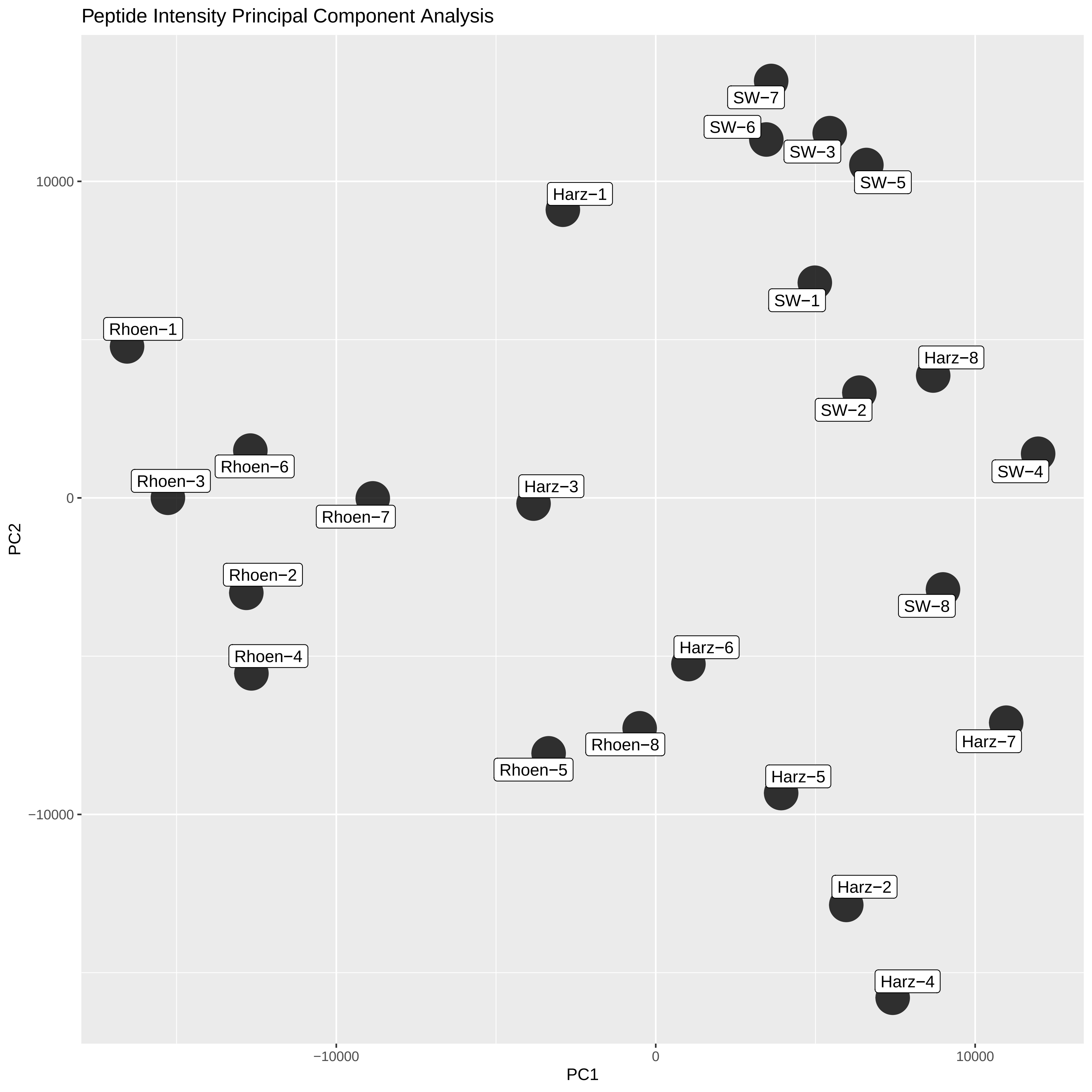

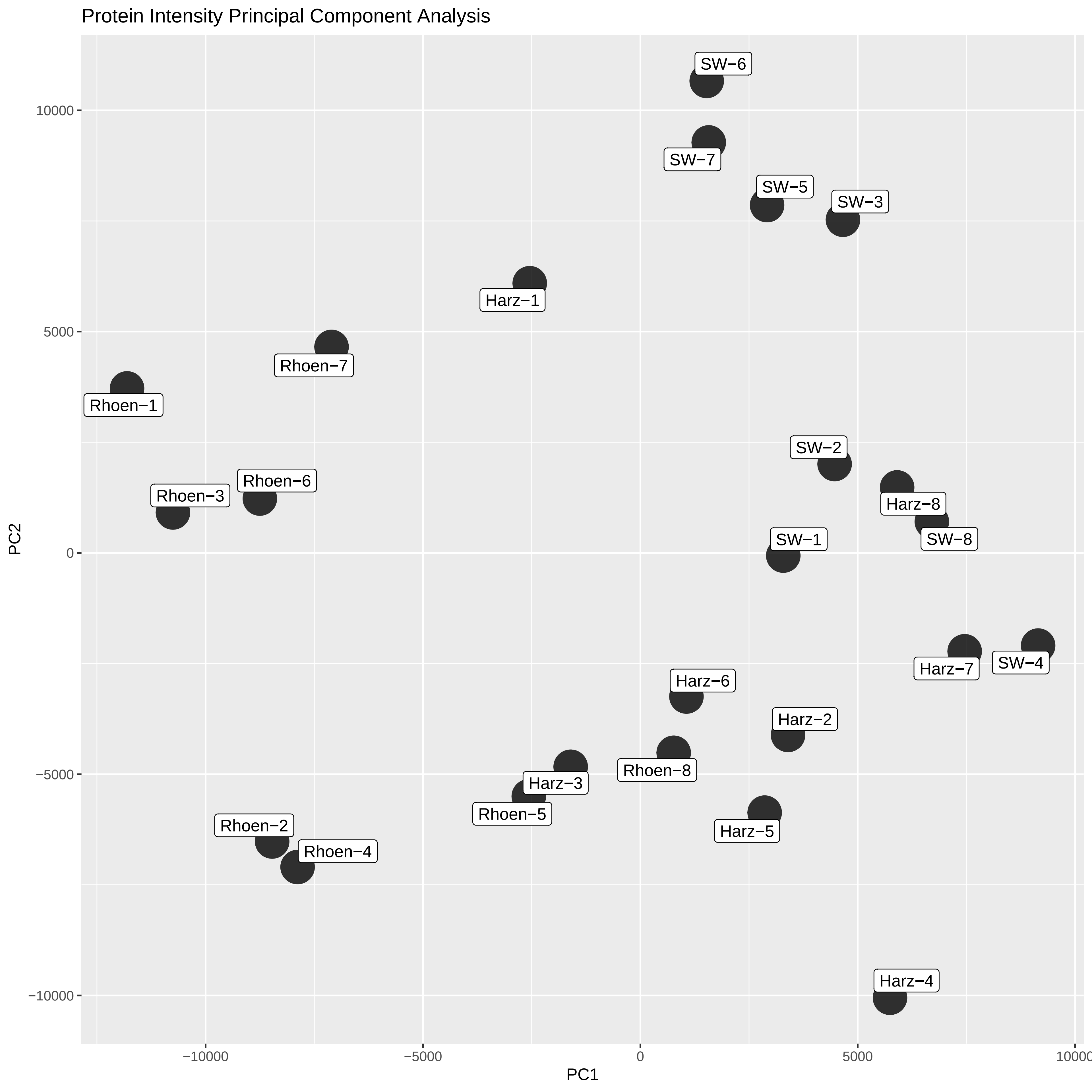

### QC-SamplePrep.pdf

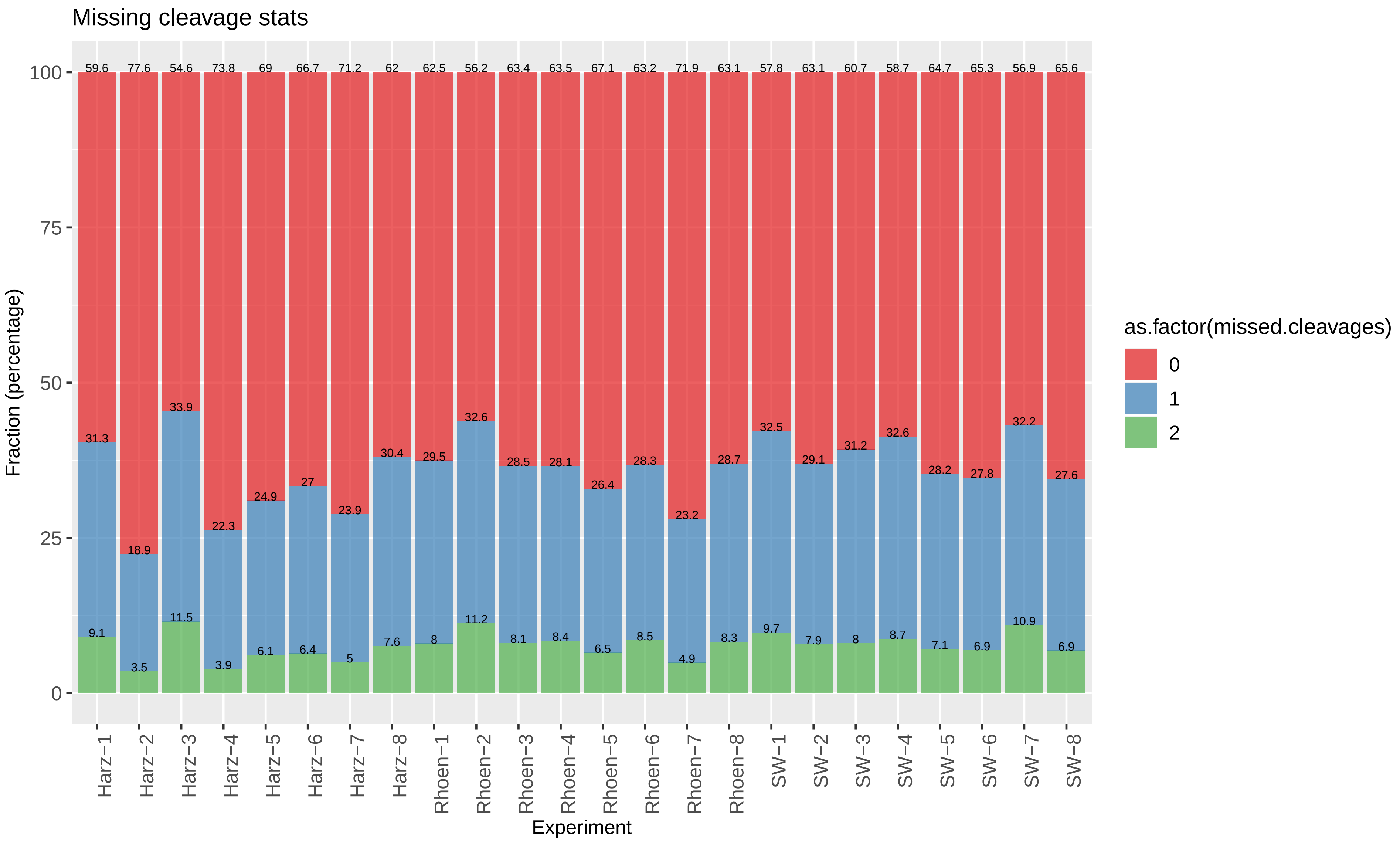

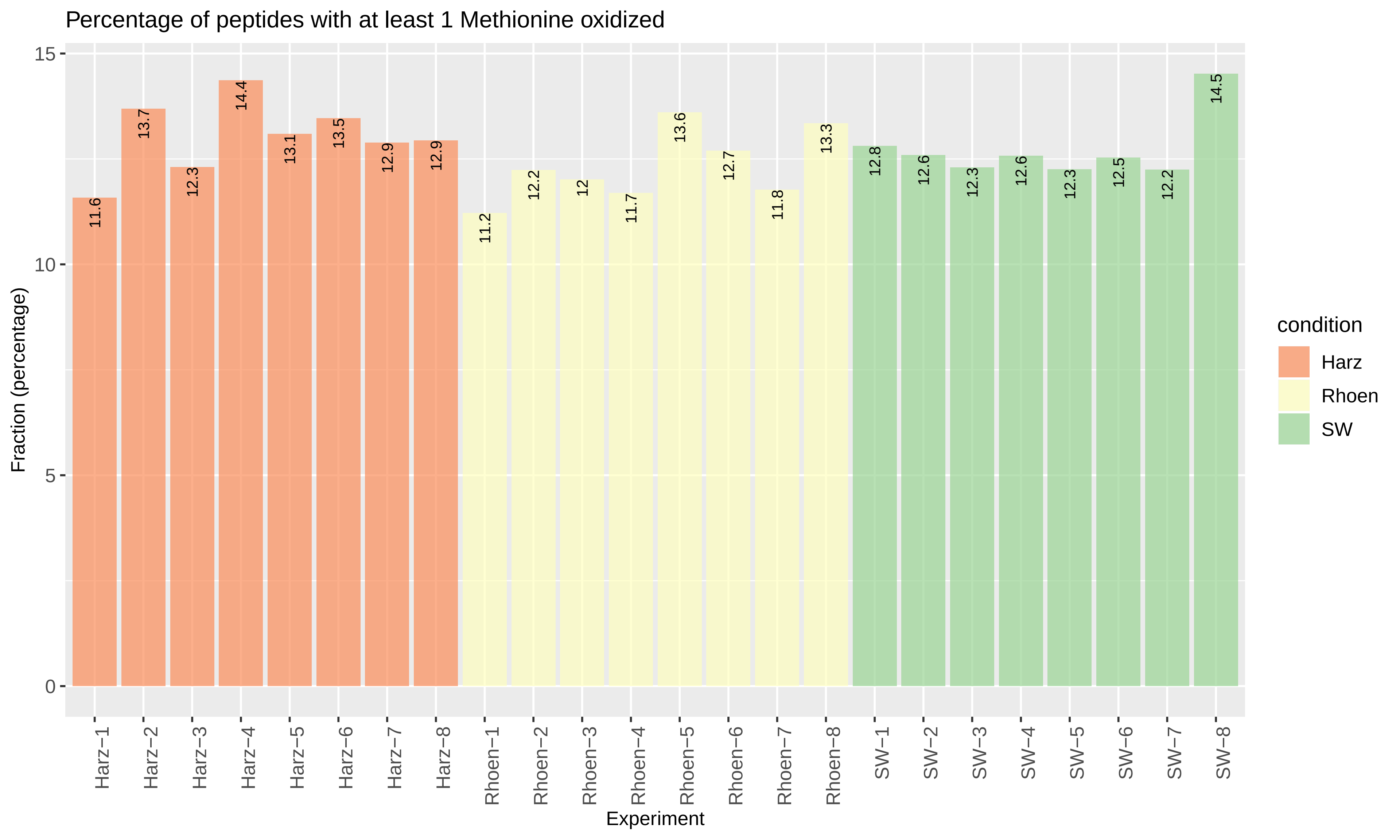

### QC_Plots_CHARGESTATE.pdf

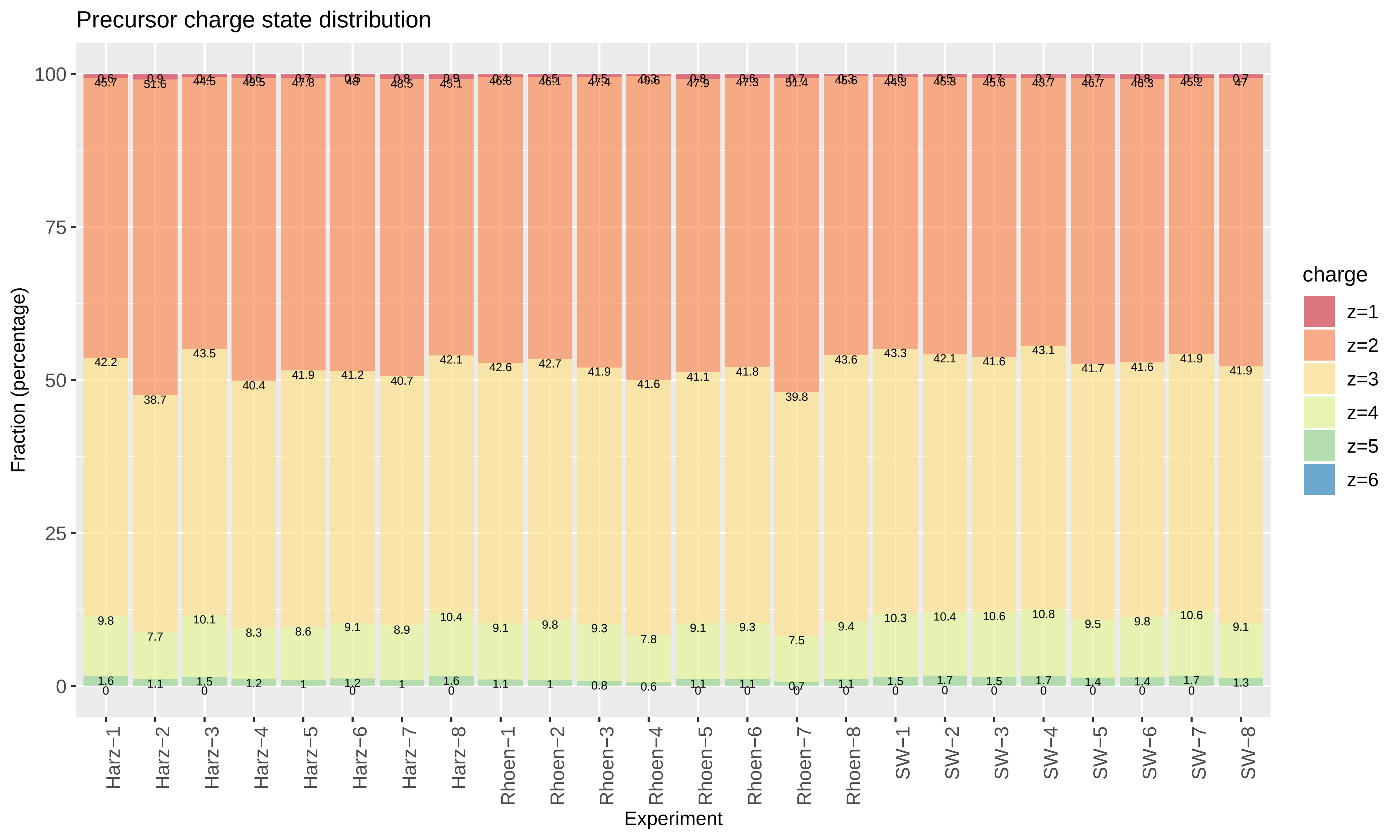

### QC_Plots_IONS.pdf

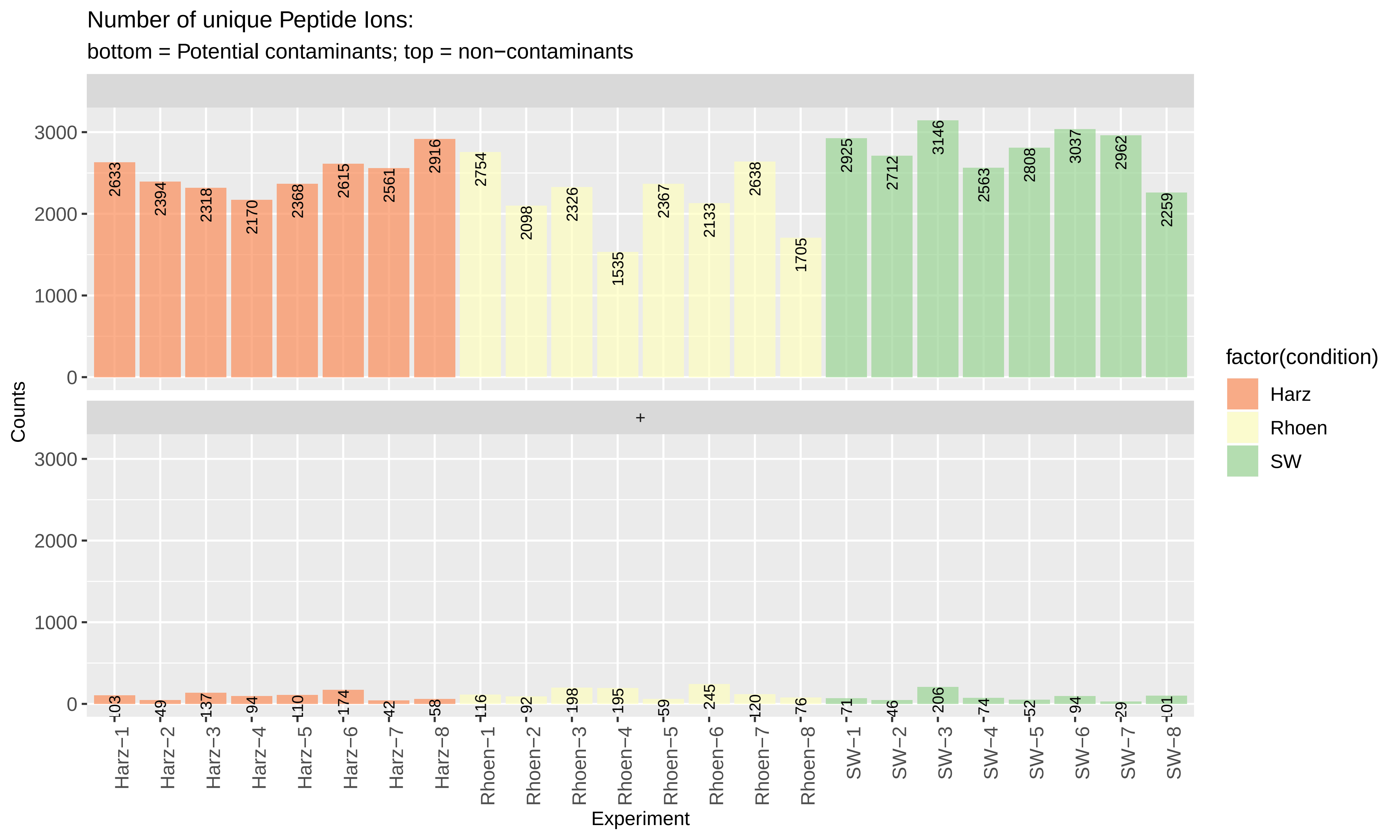

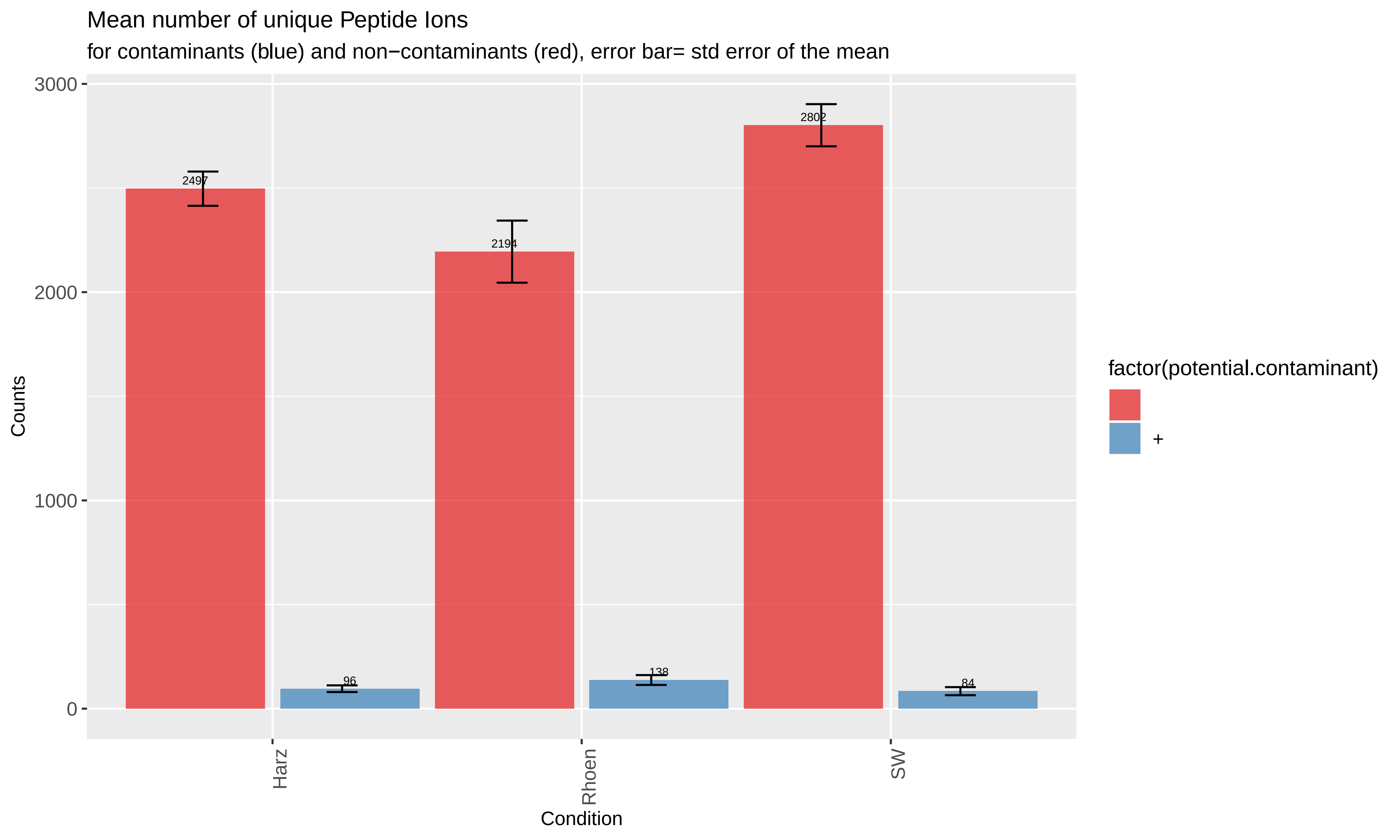

### QC_Plots_MASSERROR.pdf

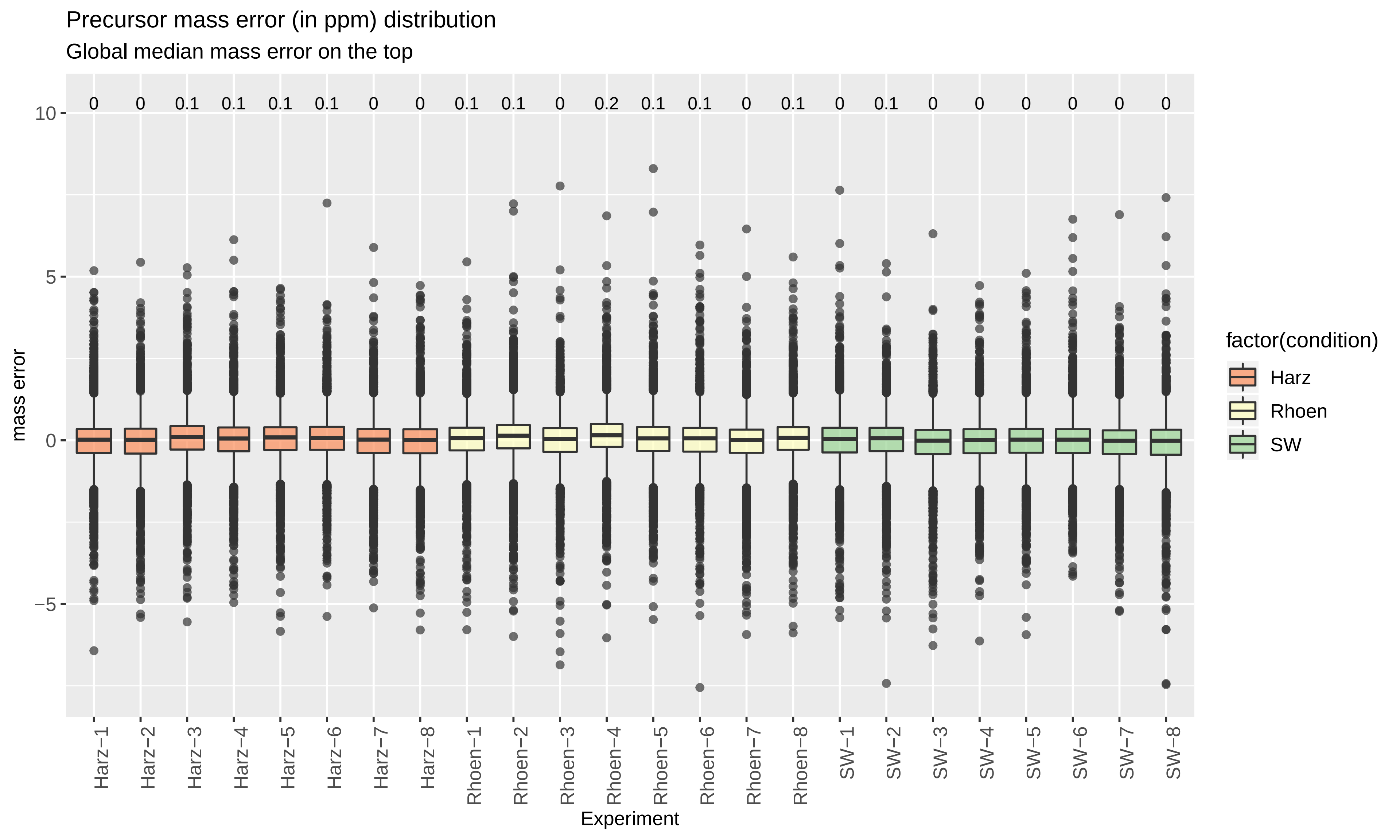

### QC_Plots_MZ.pdf

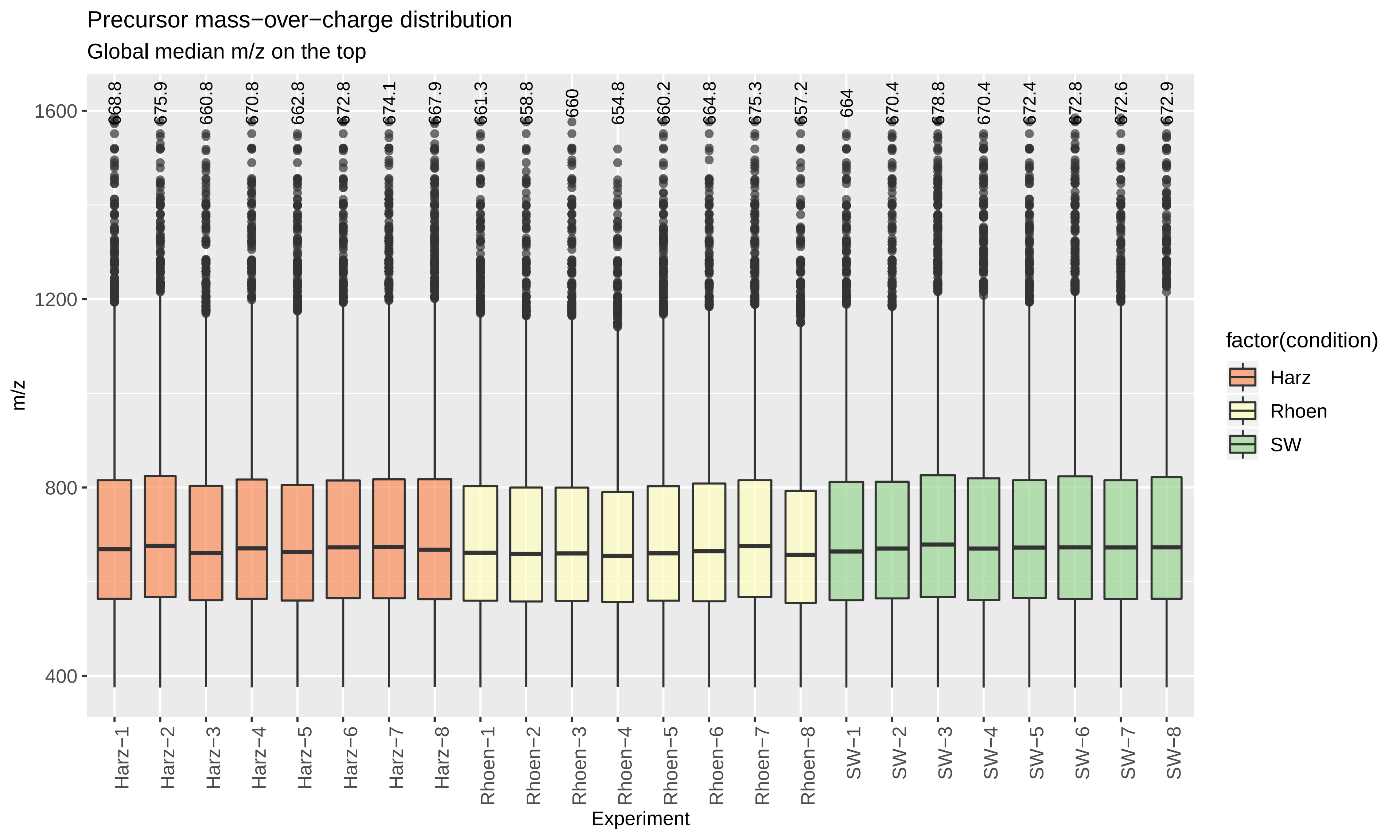

### QC_Plots_PepDetect.pdf

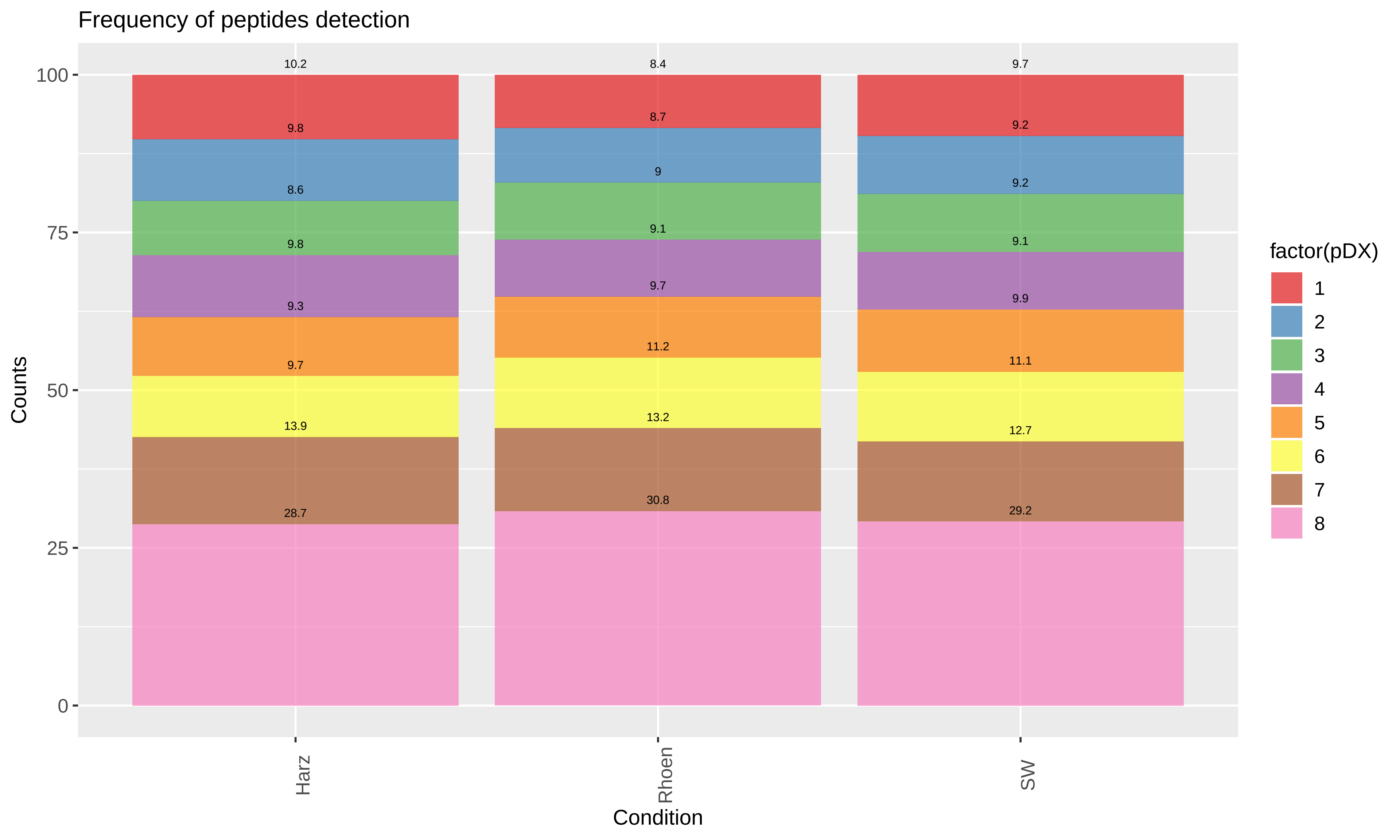

### QC_Plots_PepIonOversampling.pdf

Peptide ion oversampling

Based on all the peptides reported by MaxQuant

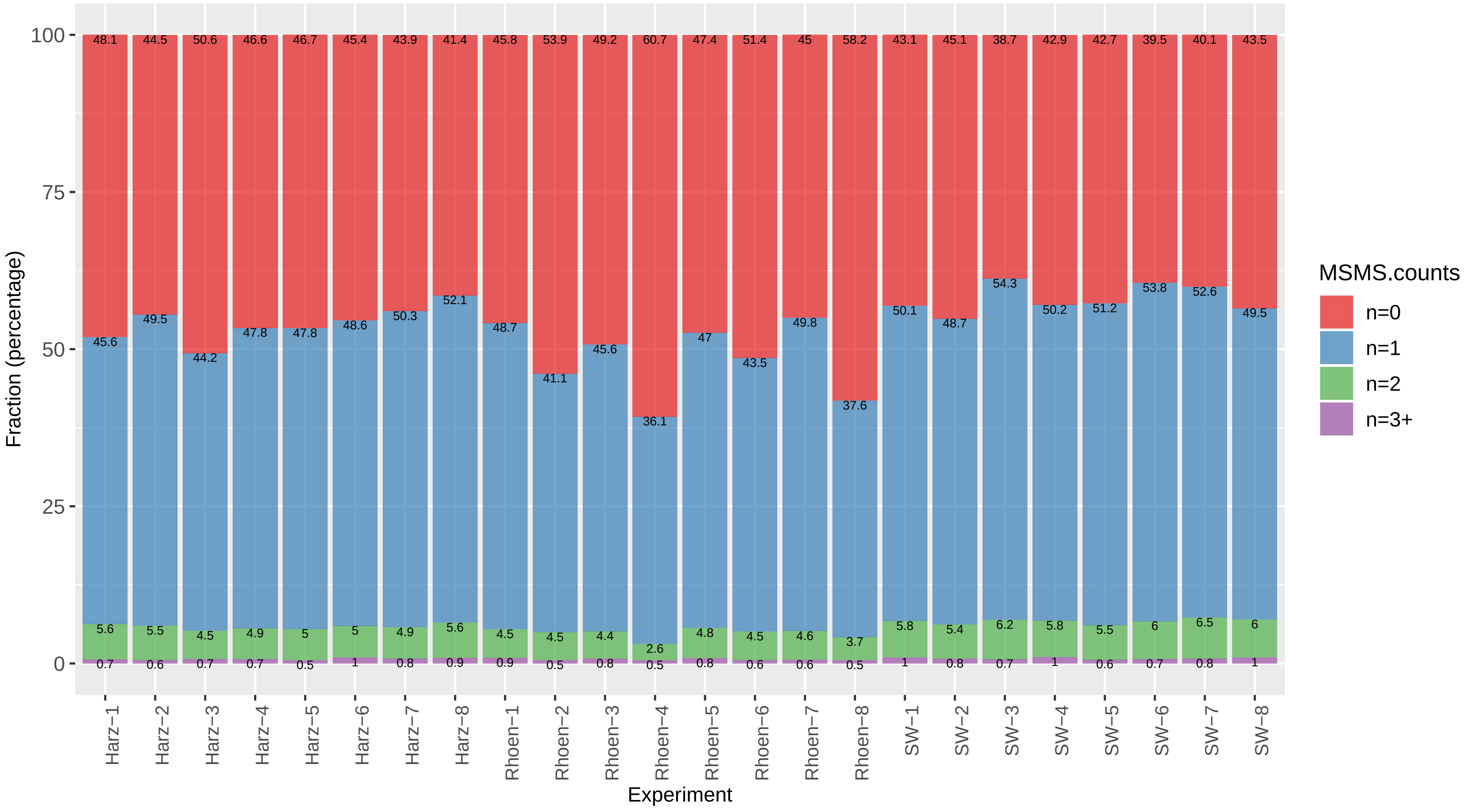

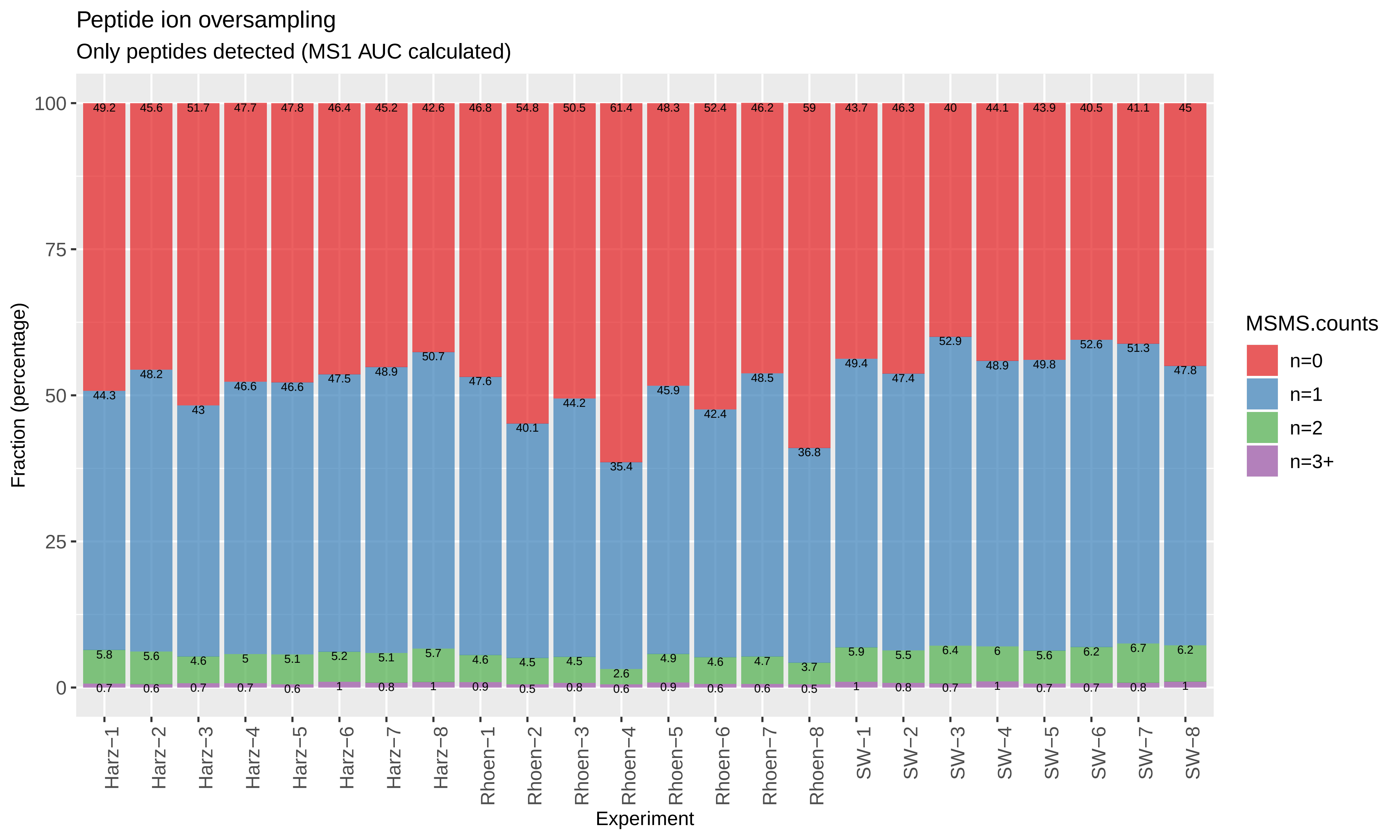

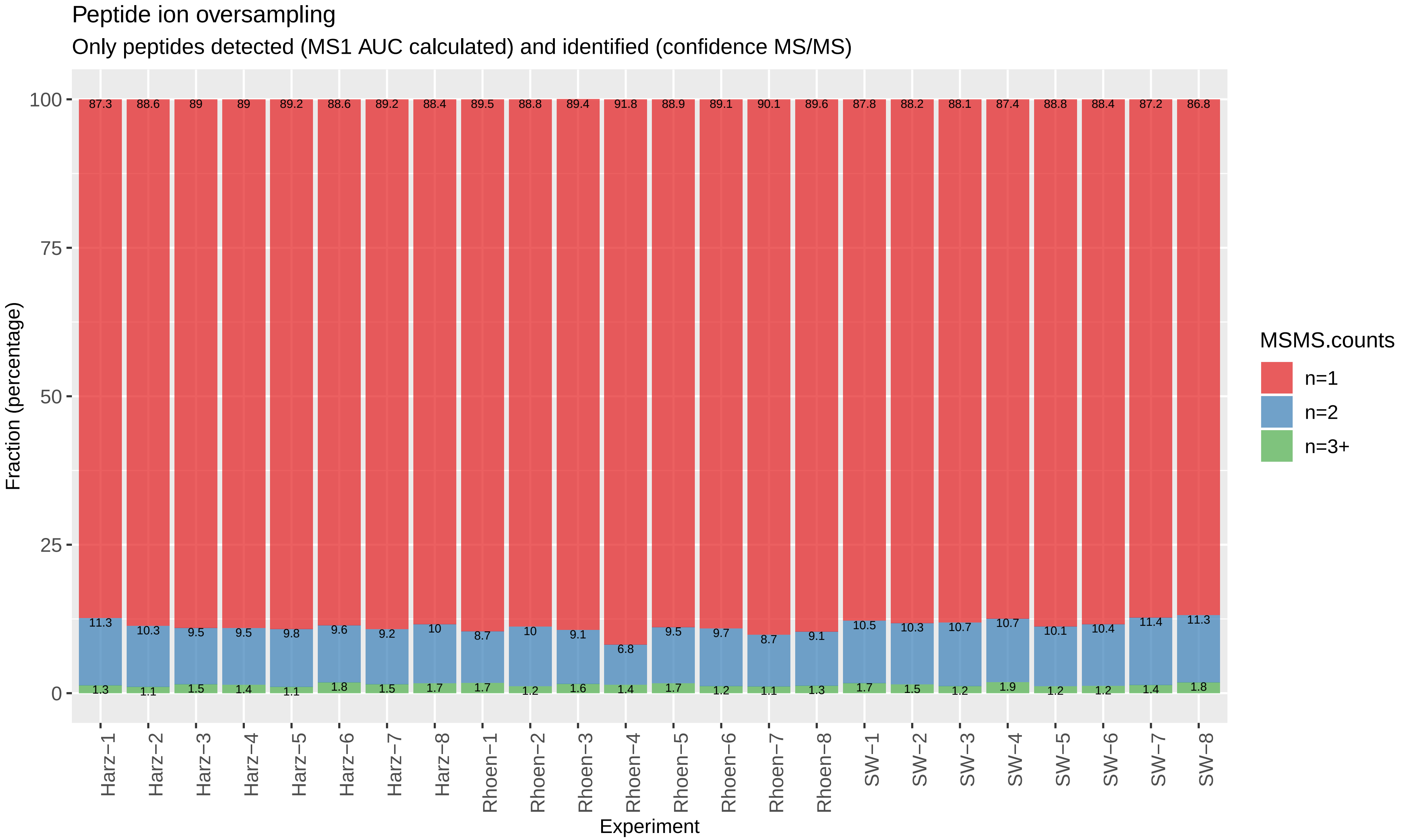

### QC_Plots_PEPTIDES.pdf

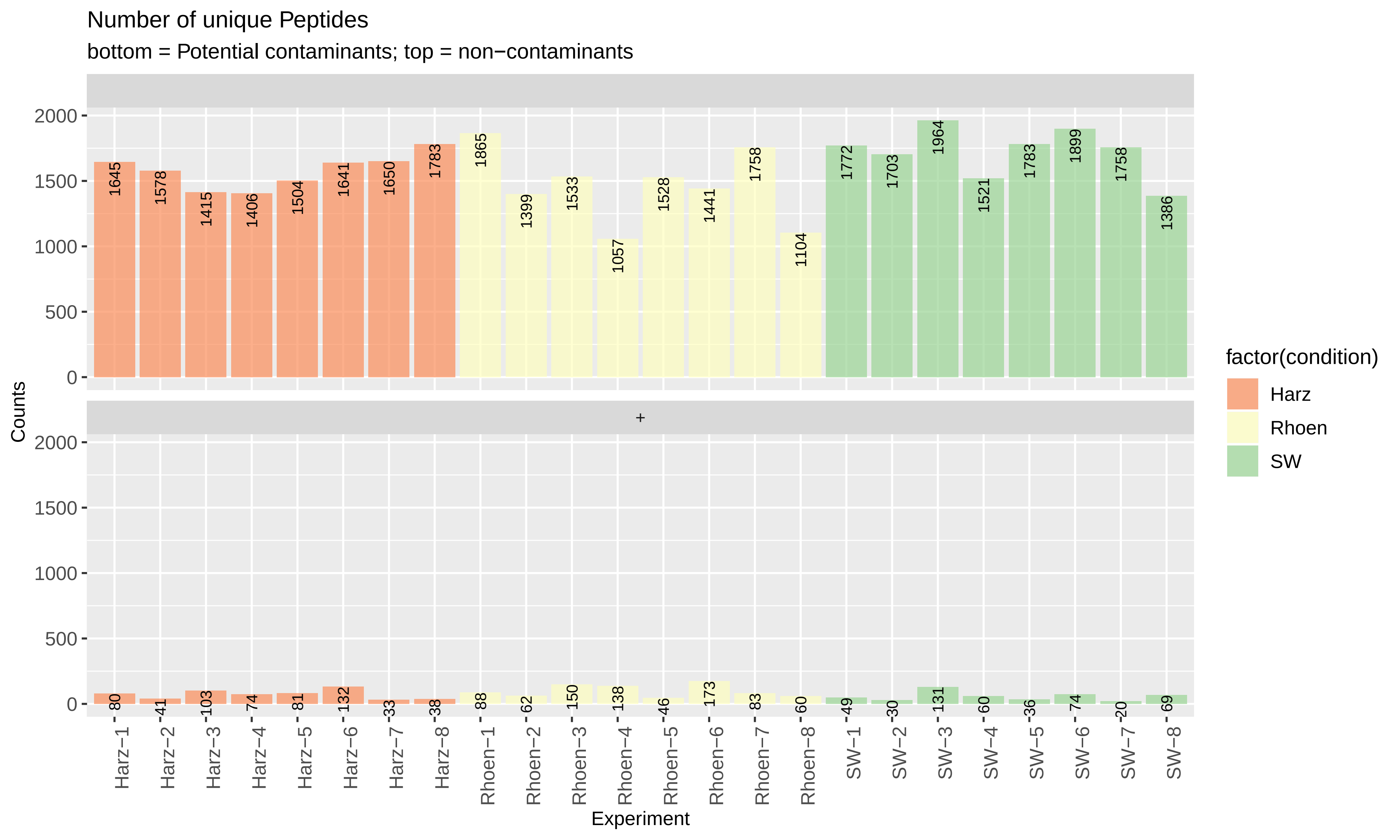

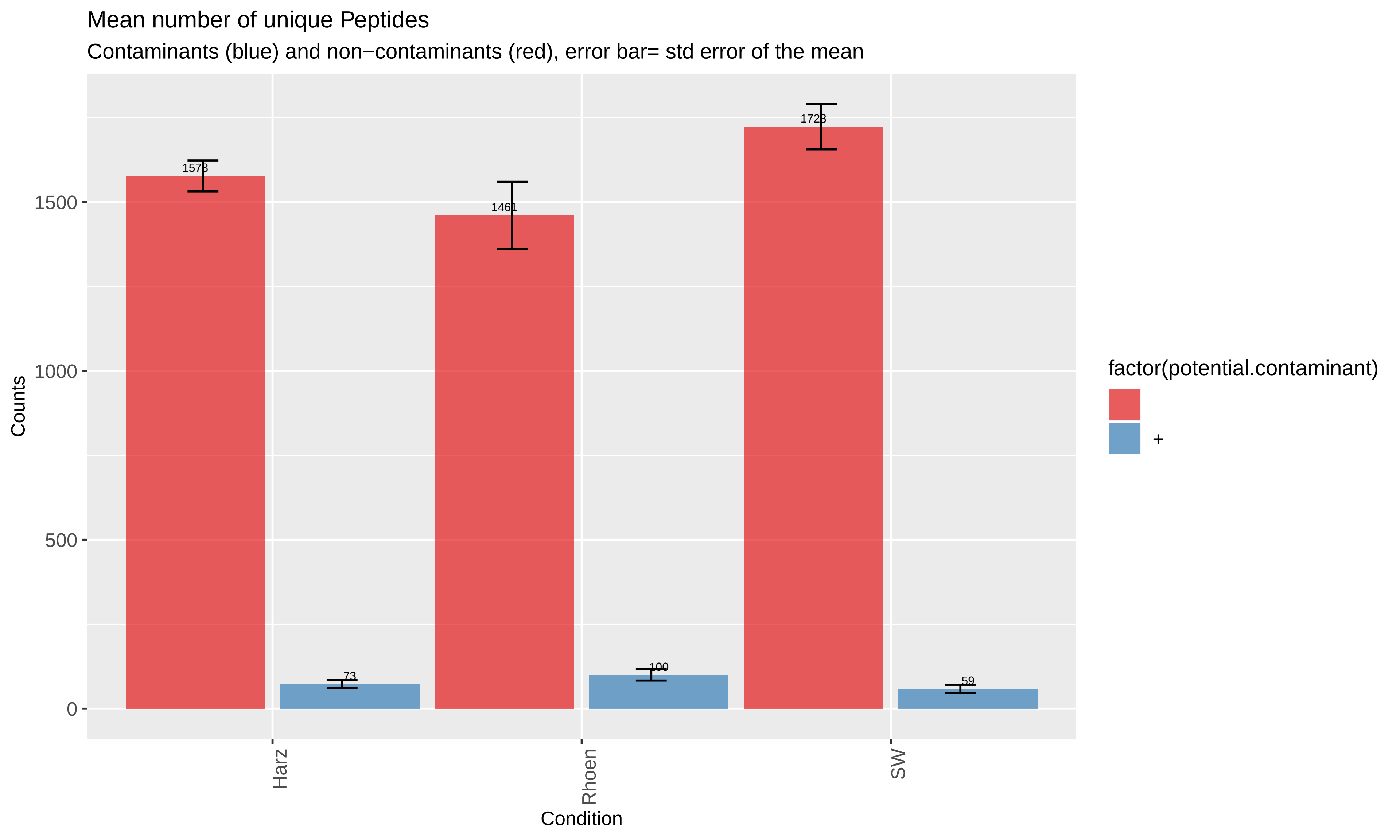

### QC_Plots_ProtDetect.pdf

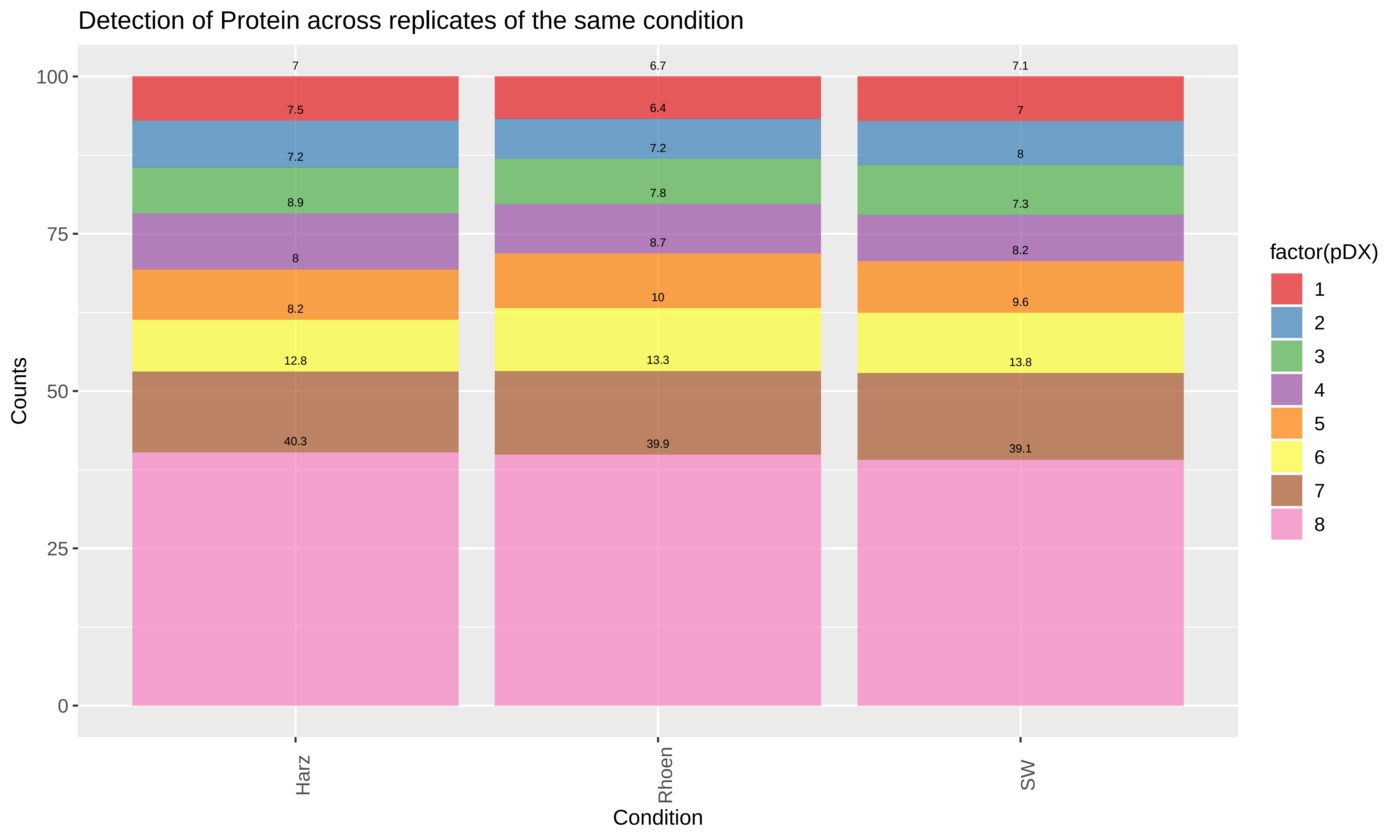

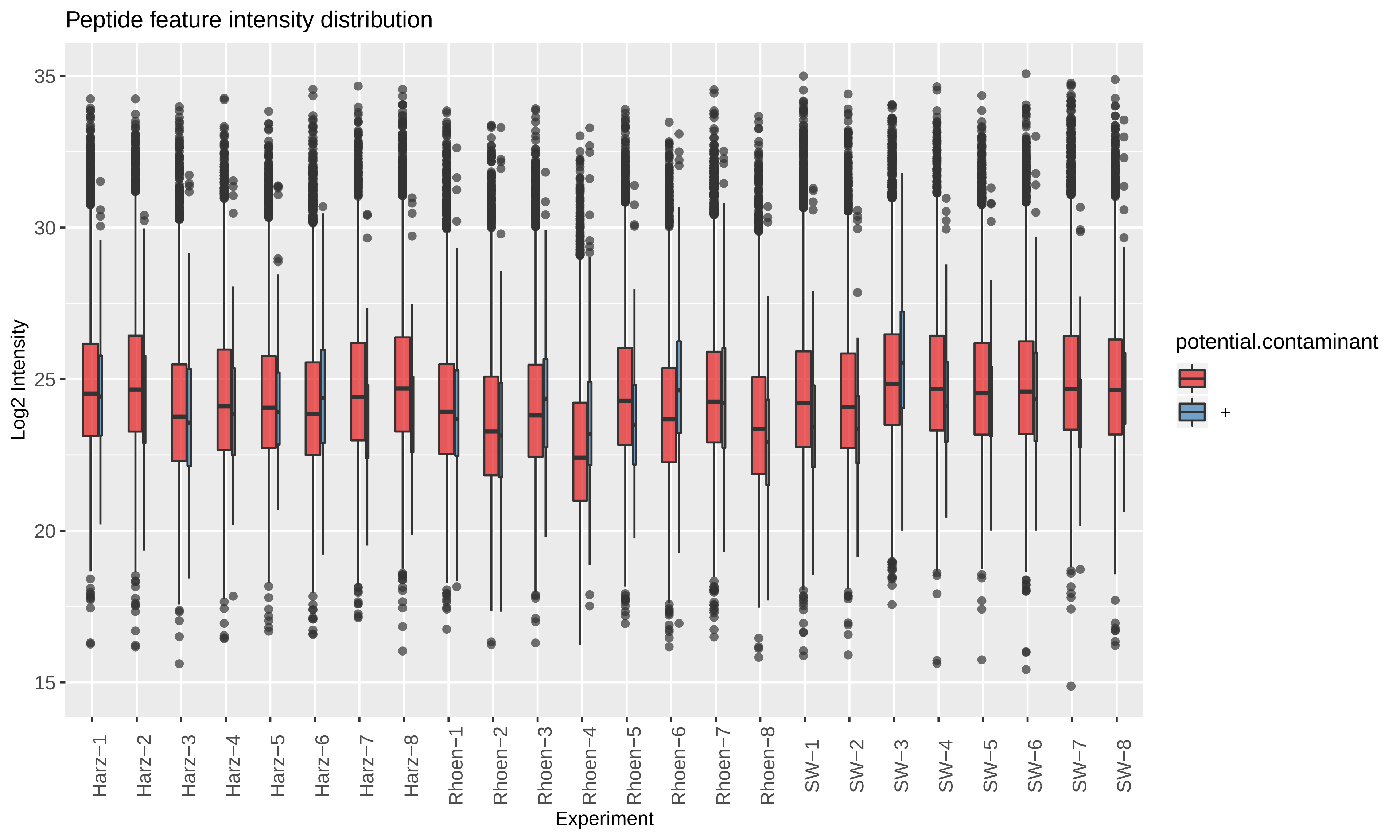

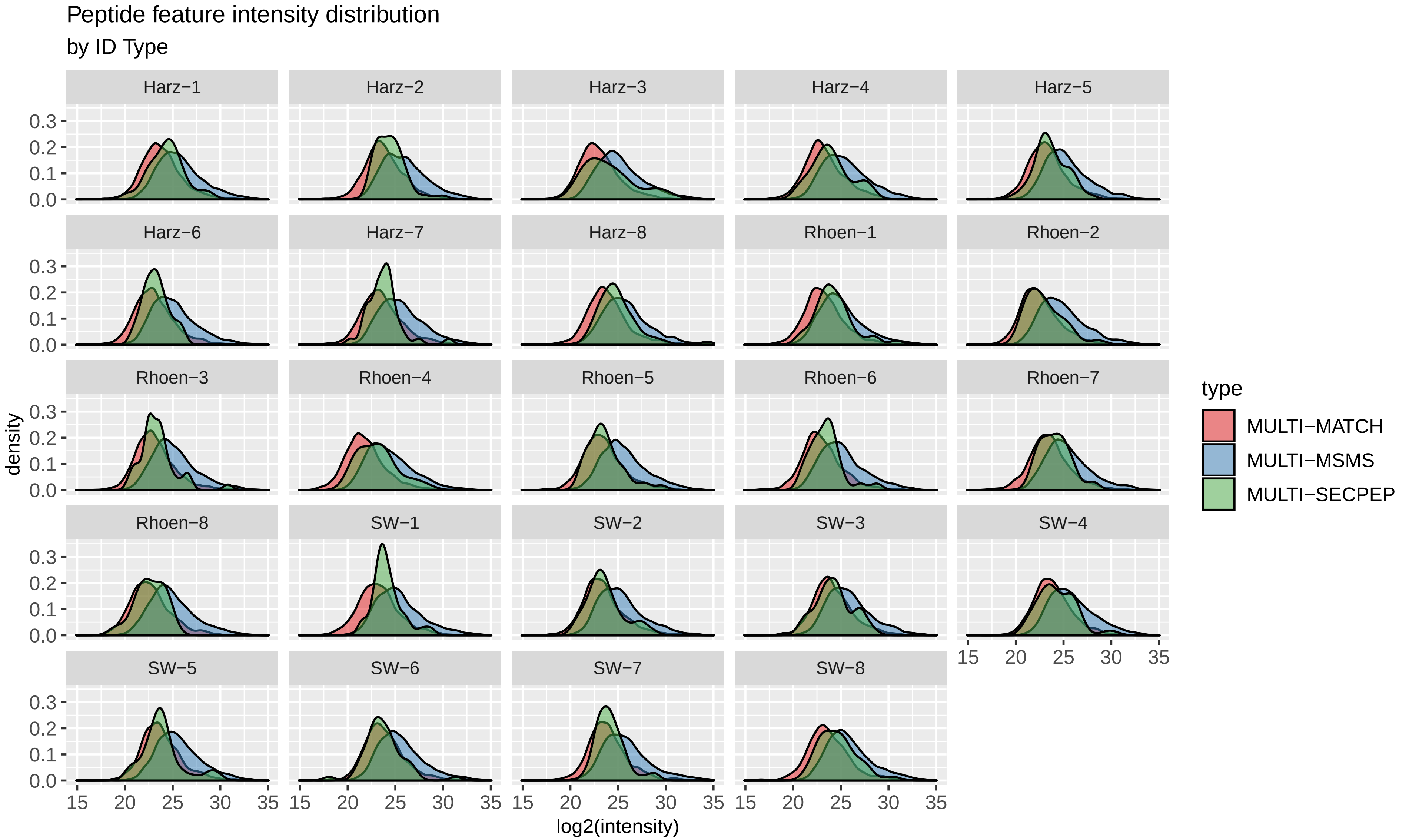

### QC_Plots_PROTEINS.pdf

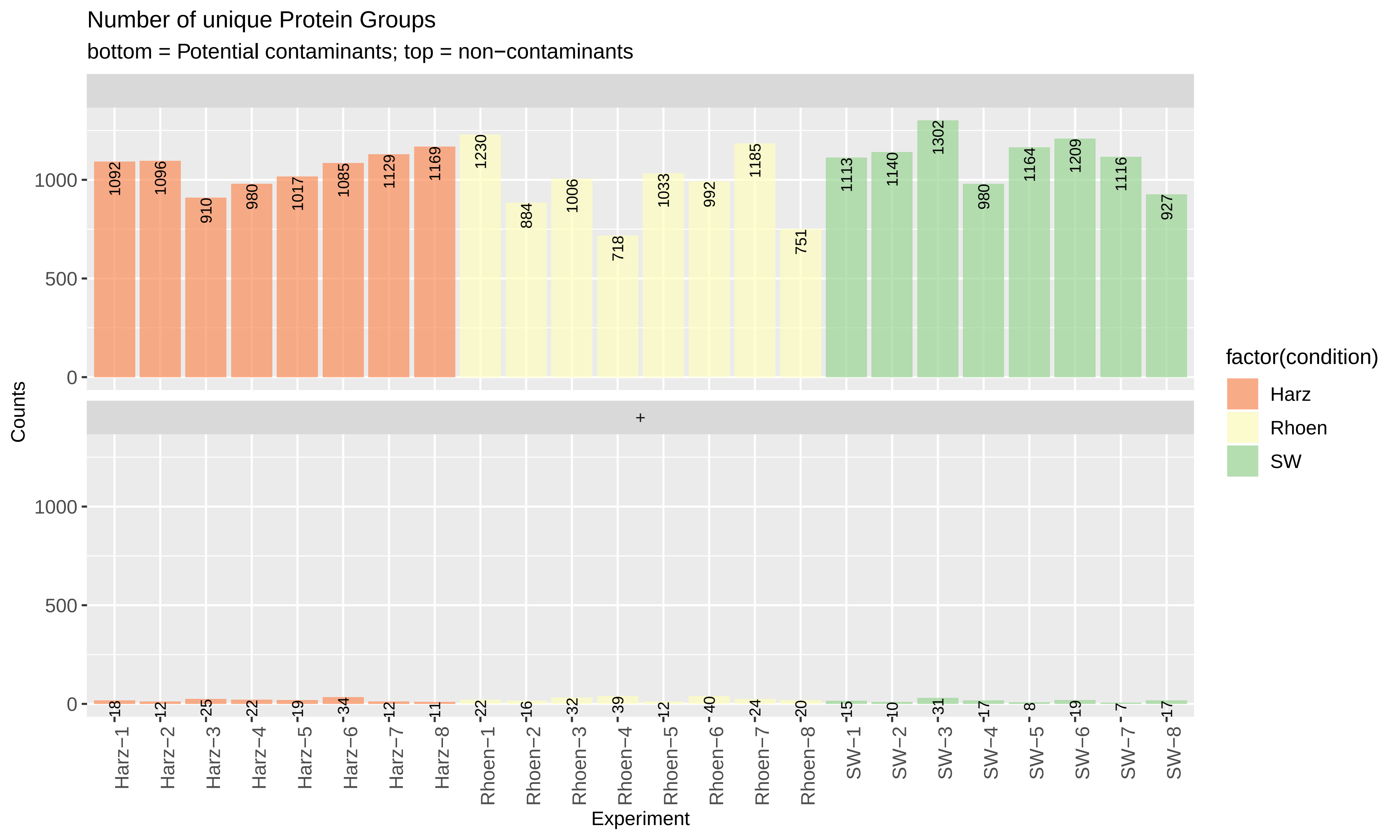

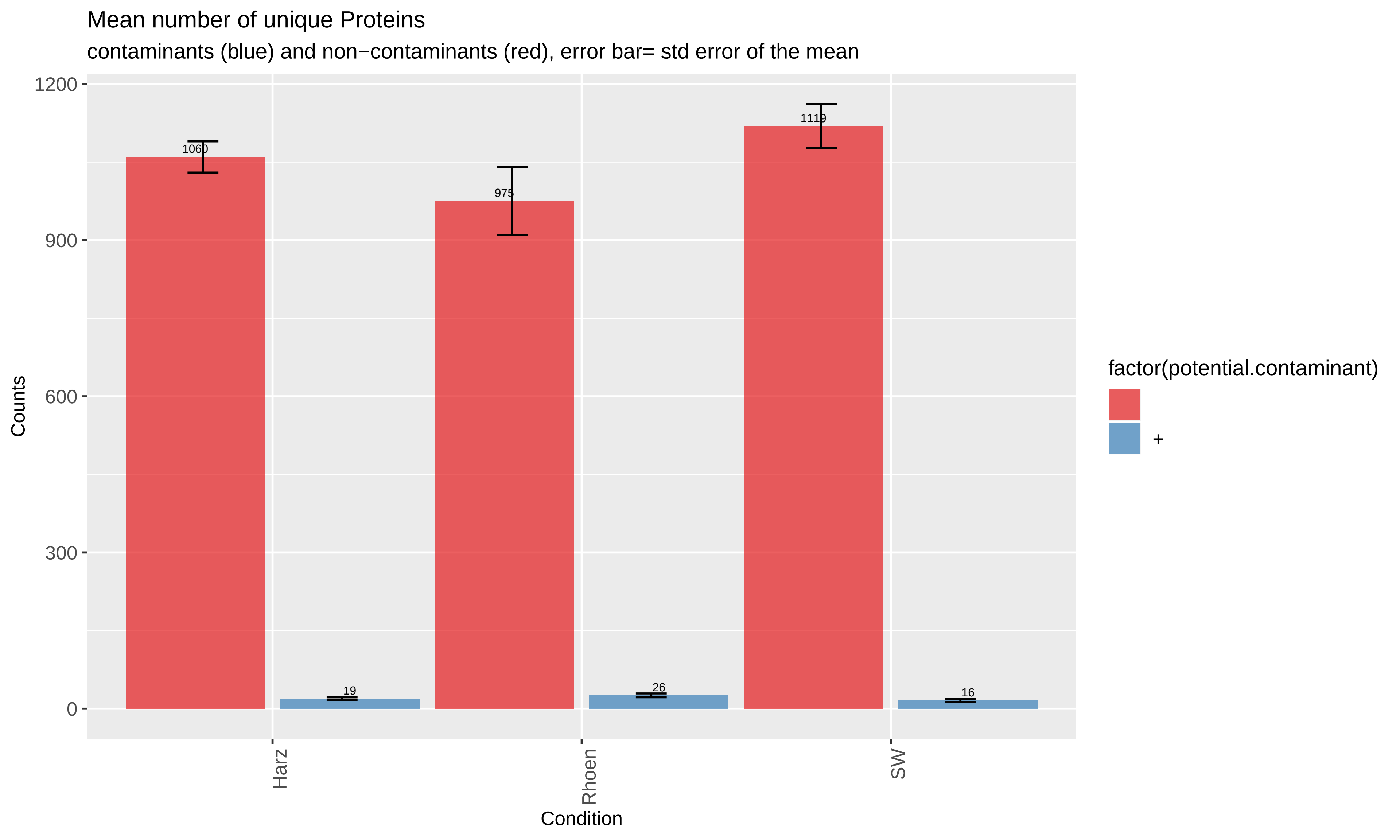

### QC_Plots_PSM.pdf

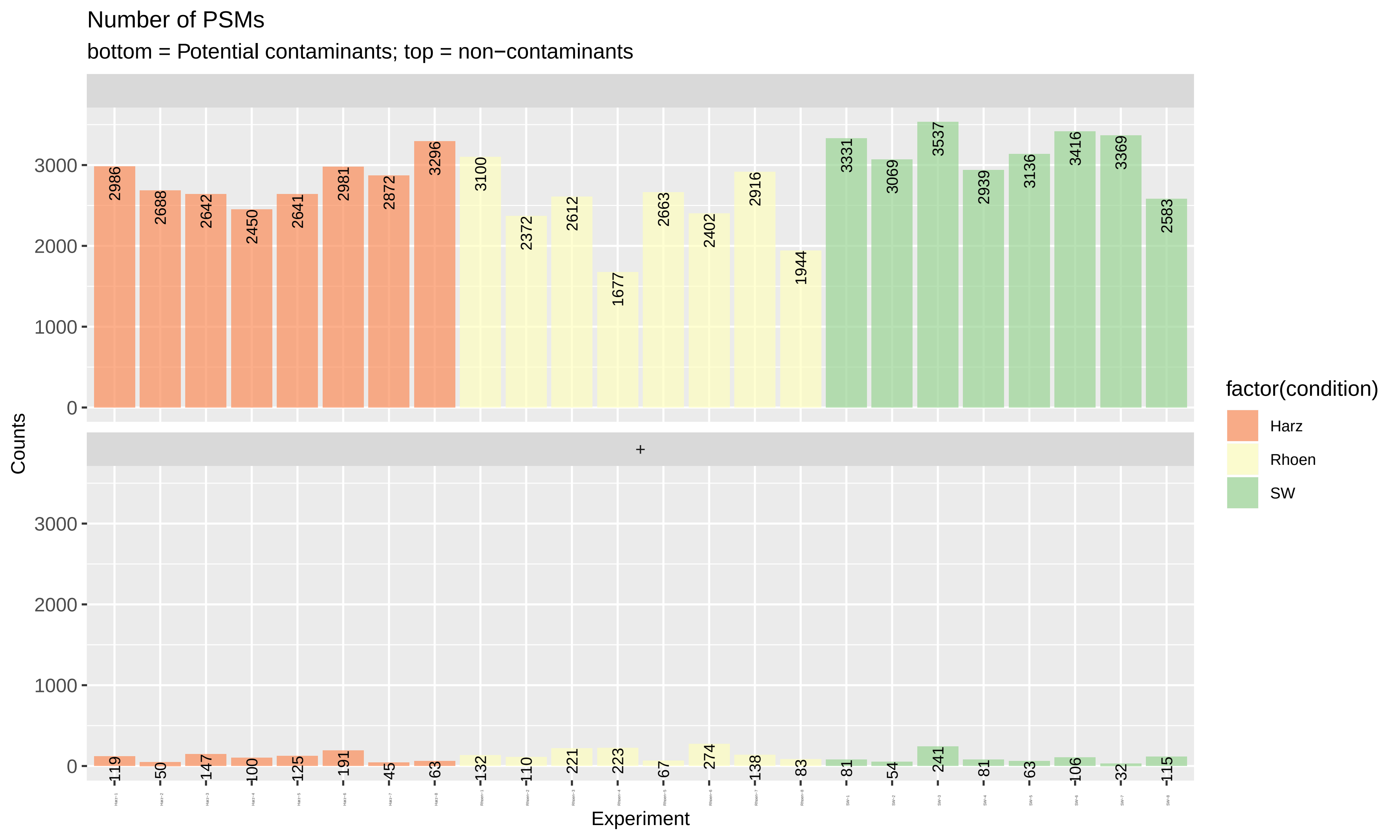

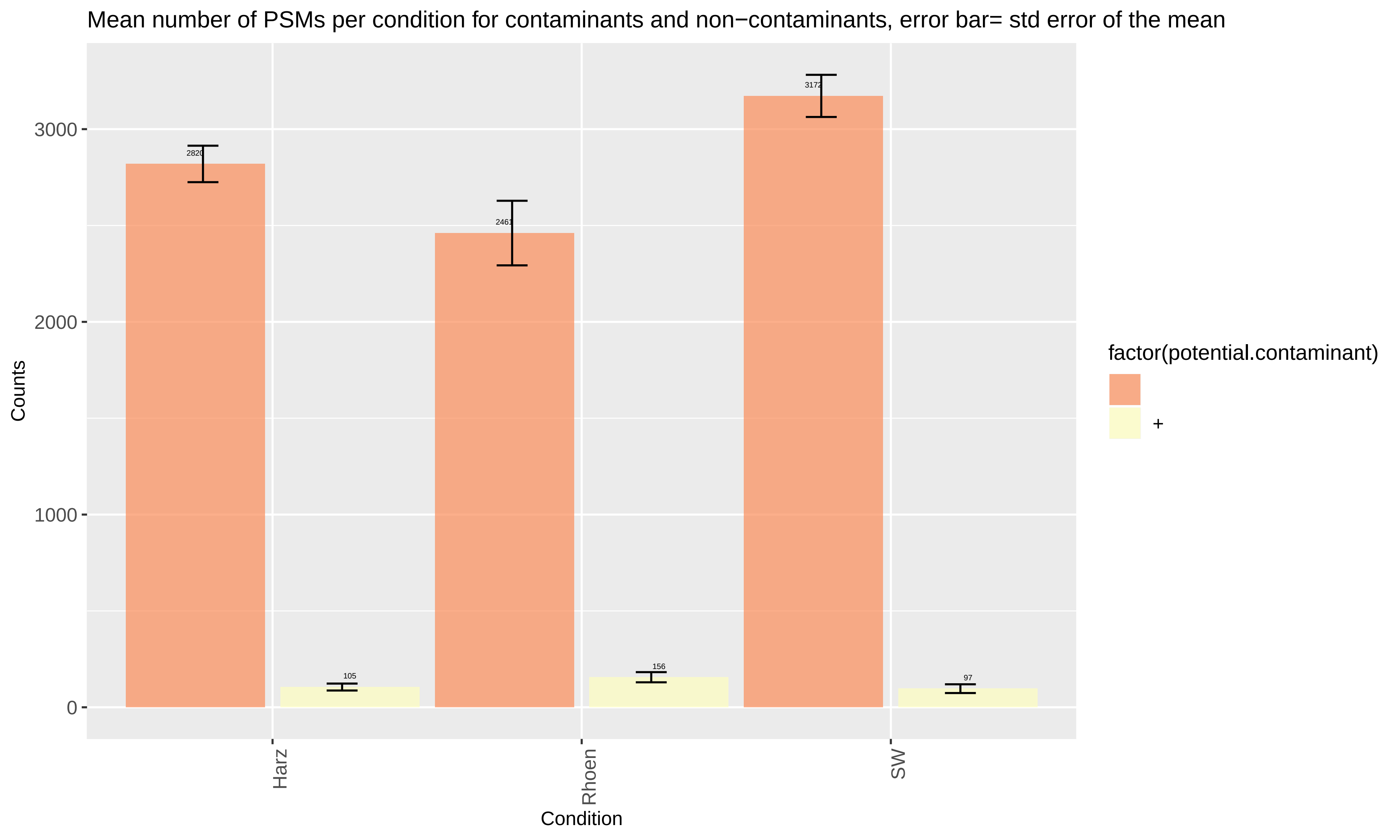

### QC_Plots_TYPE.pdf

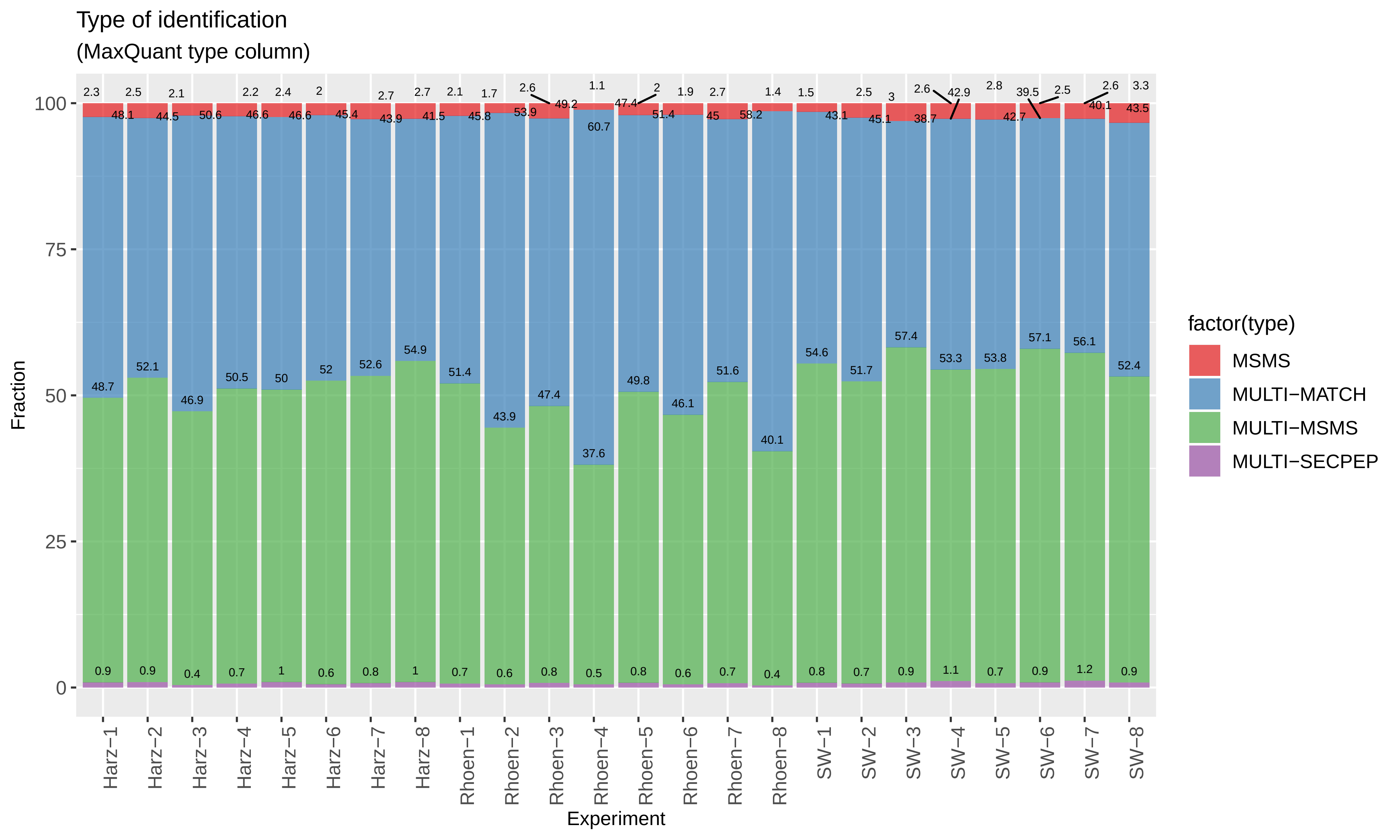

### qcPlots_evidence.qcplot.correlationMatrixBR.pdf

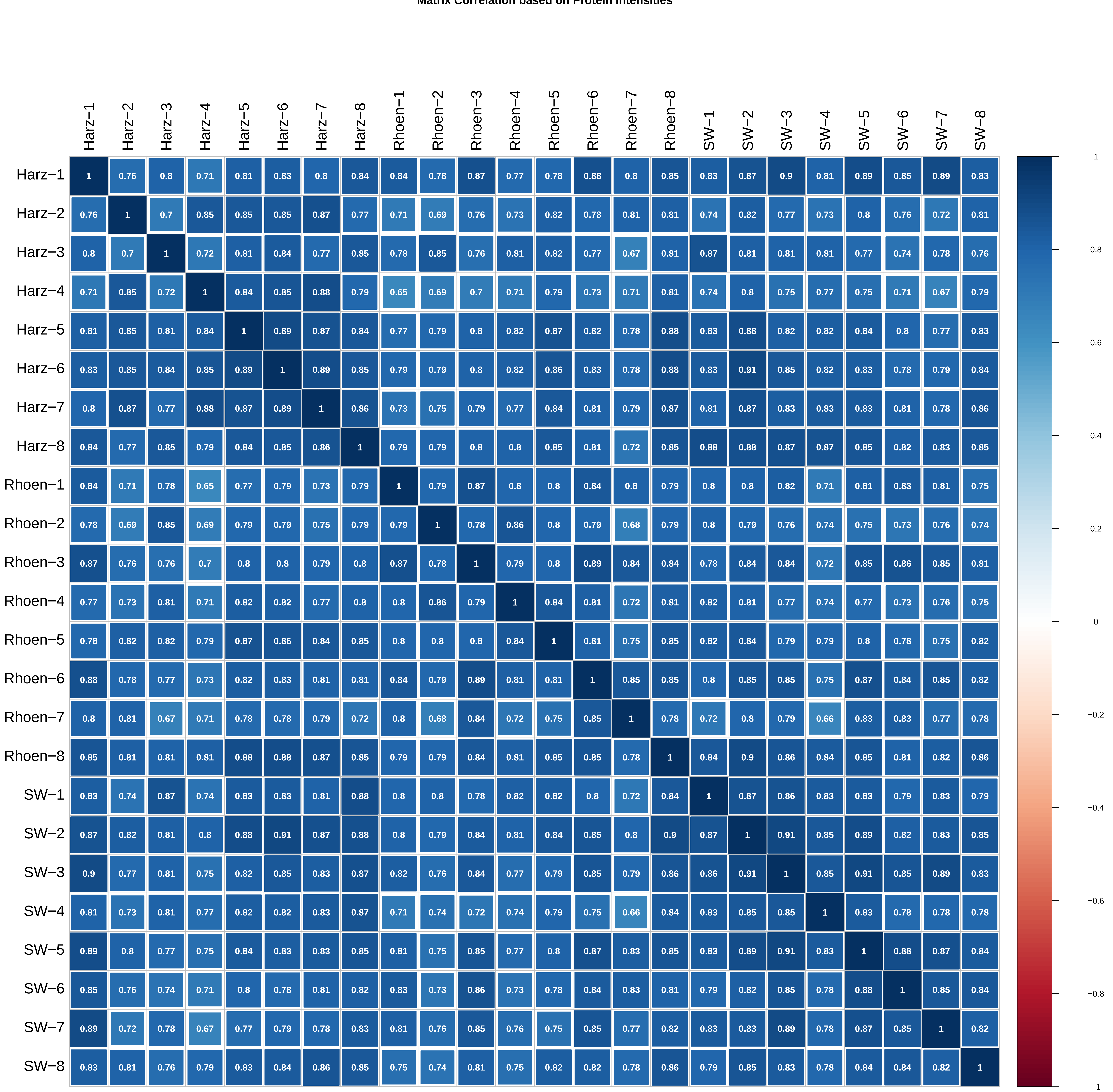

**Clustering Biological Replicates**

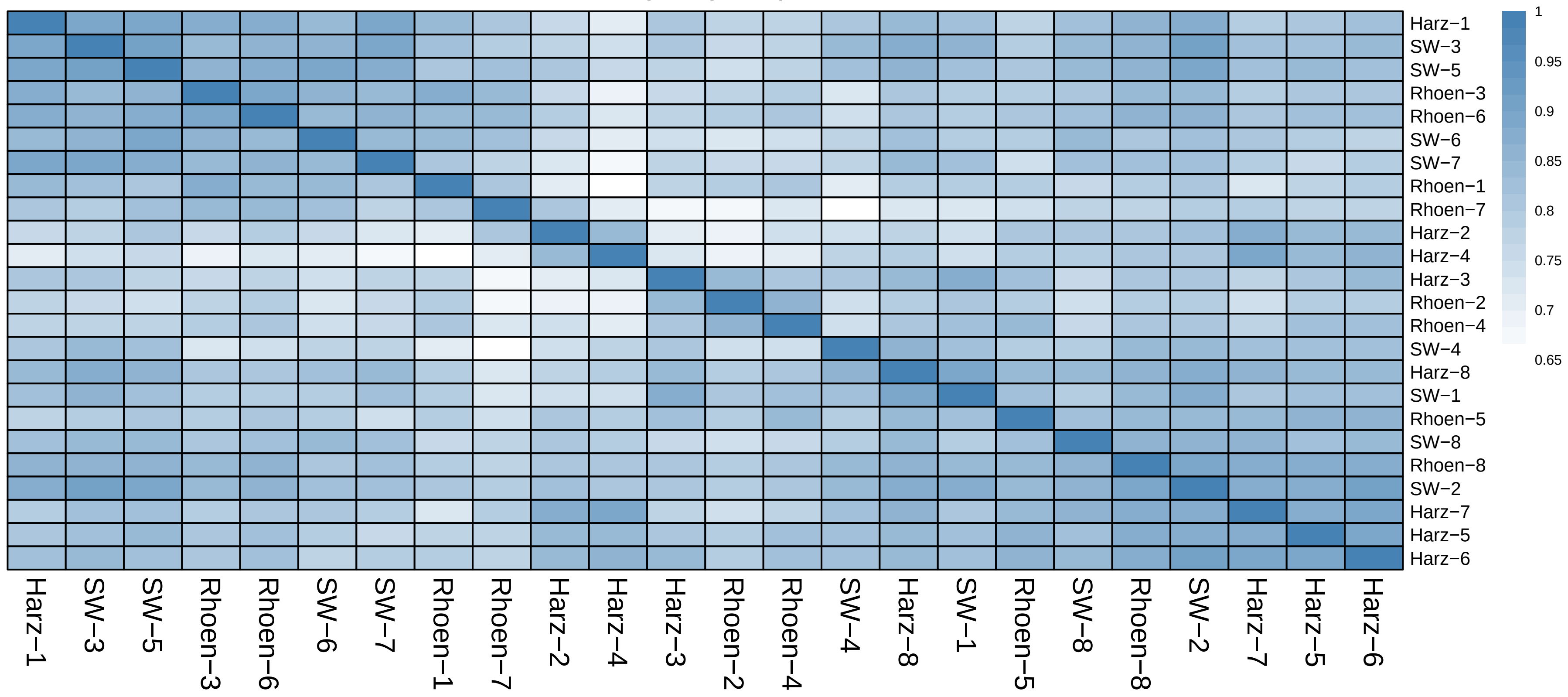

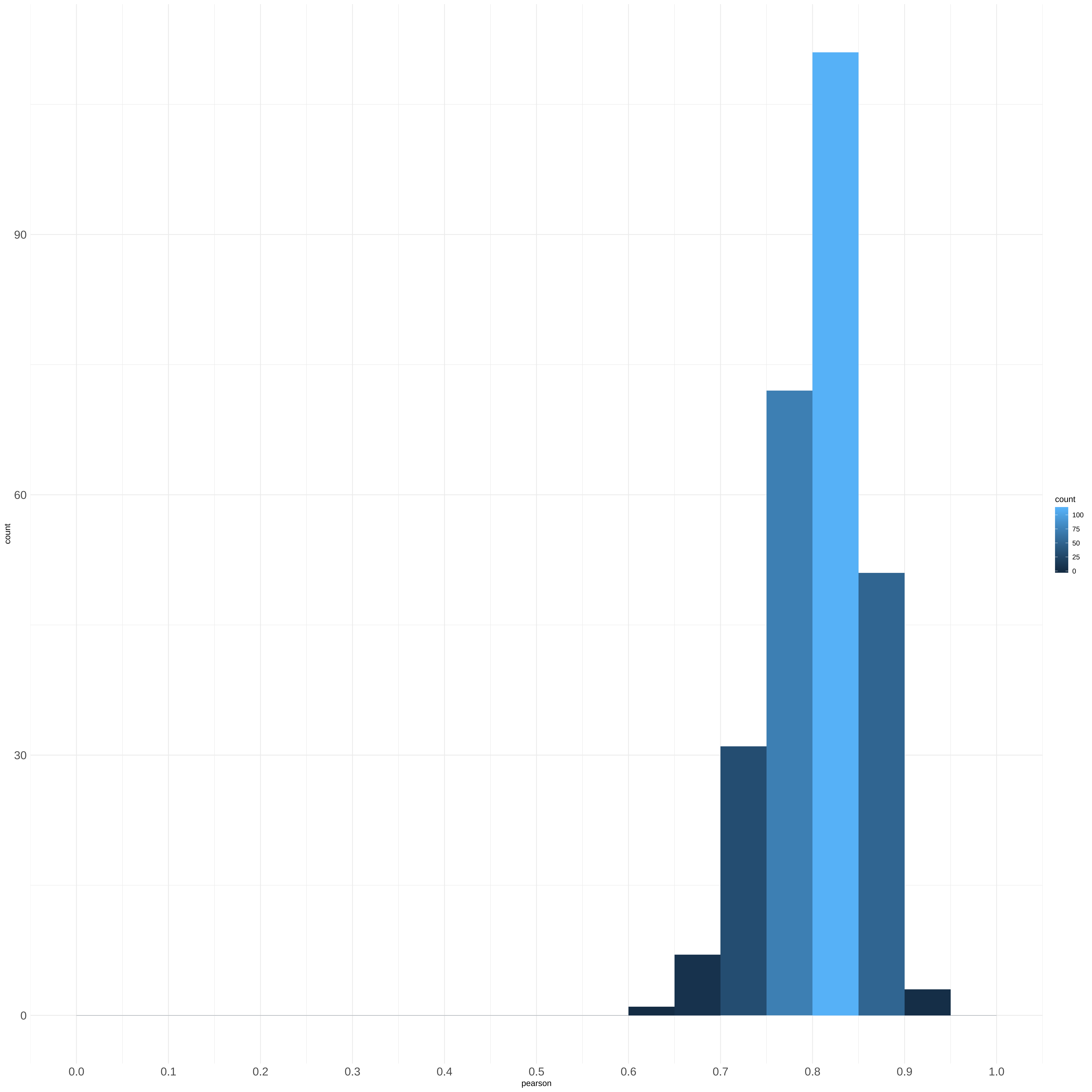

### qcPlots_evidence.qcplot.correlationMatrixConditions.pdf

**Matrix Correlation based on protein intensities**

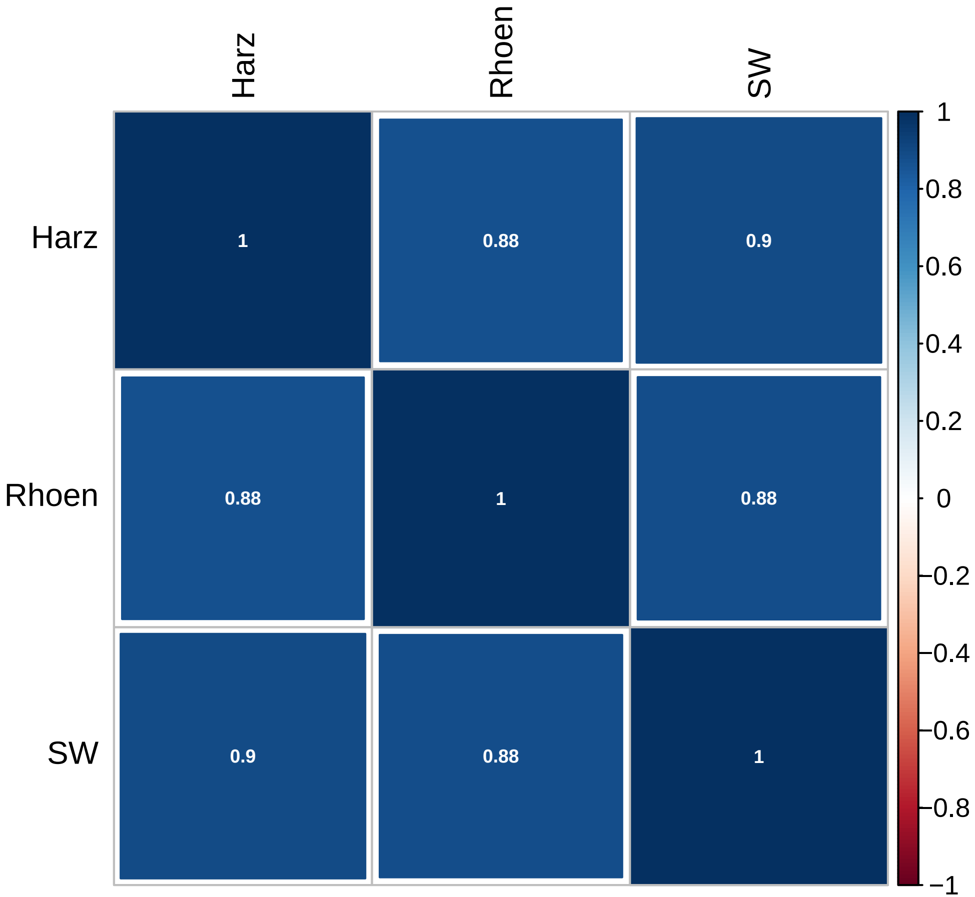

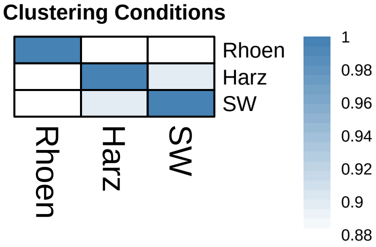

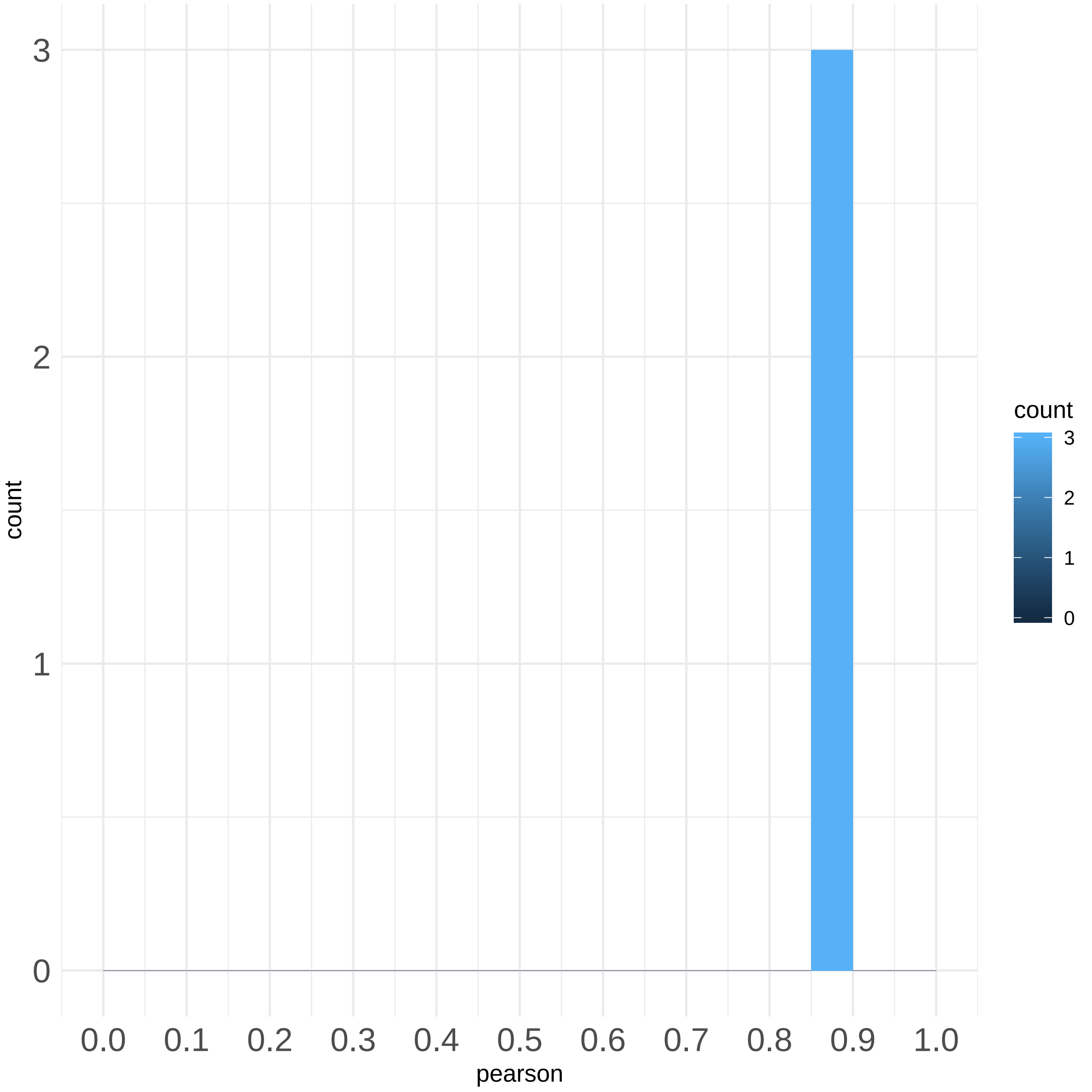

### qcPlots_evidence.qcplot.IntensityDistributions.pdf

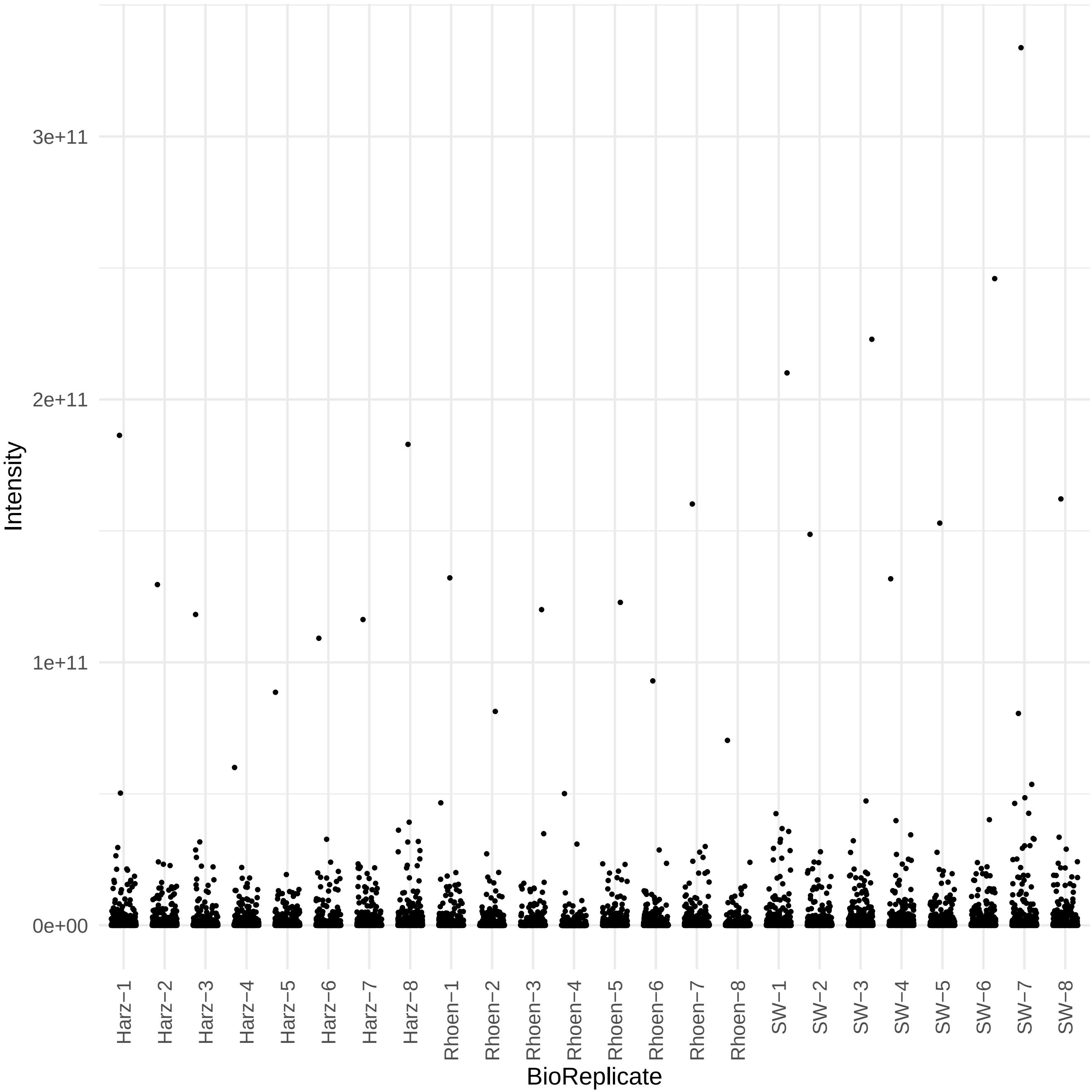

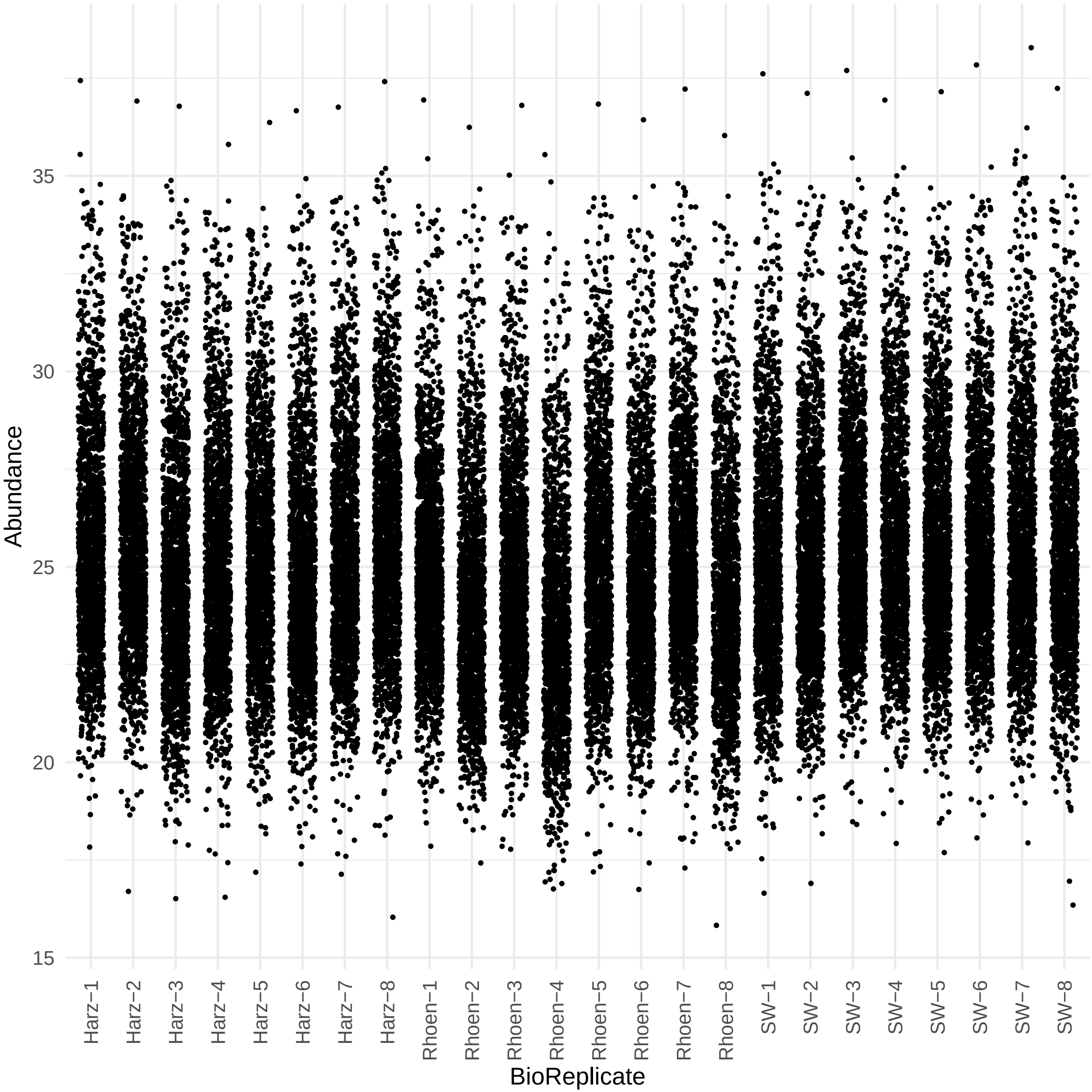

### QCSummary_ISOTOPE.pdf

Number of detected Isotope Patterns

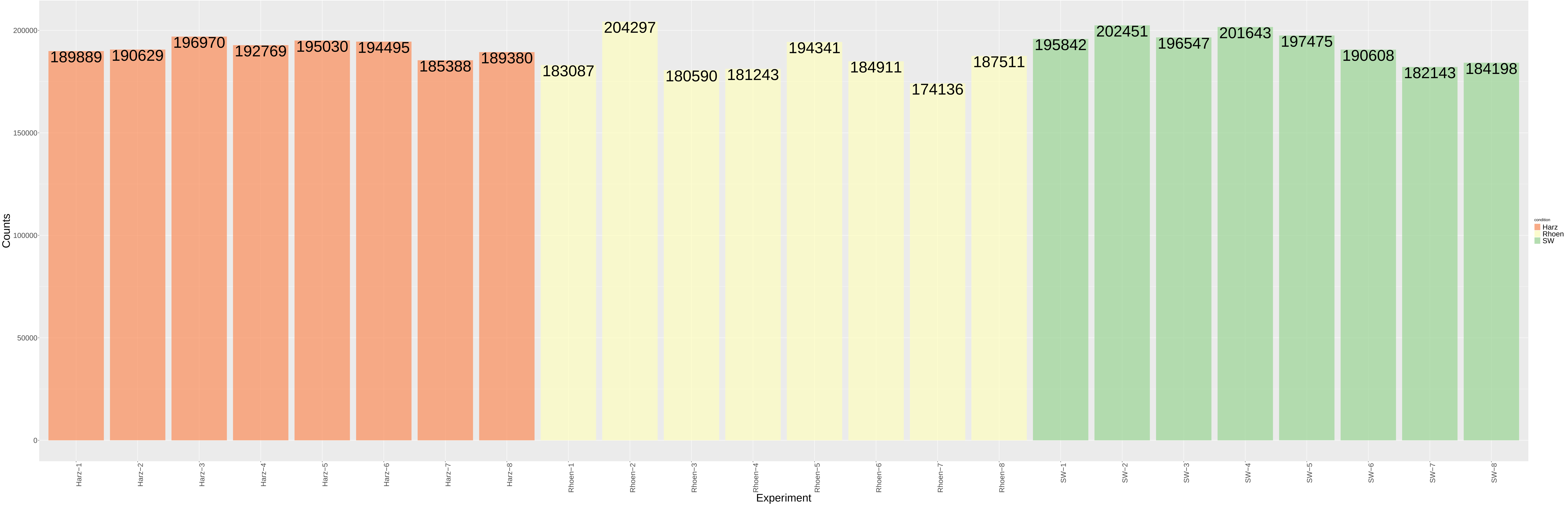

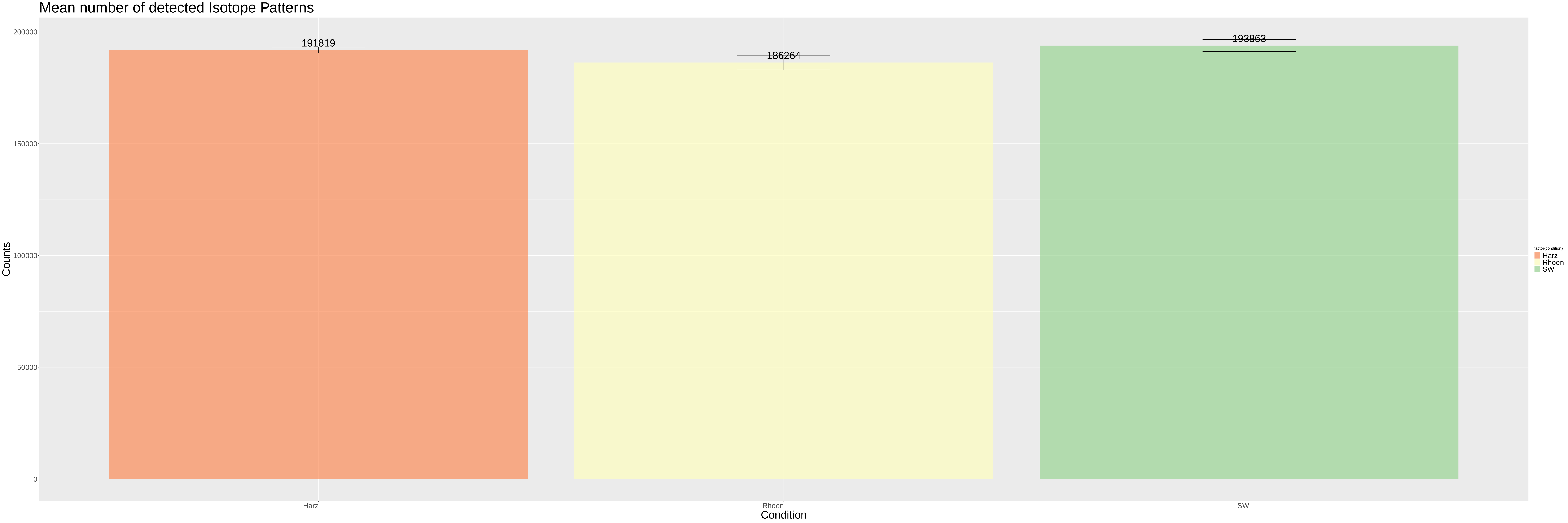

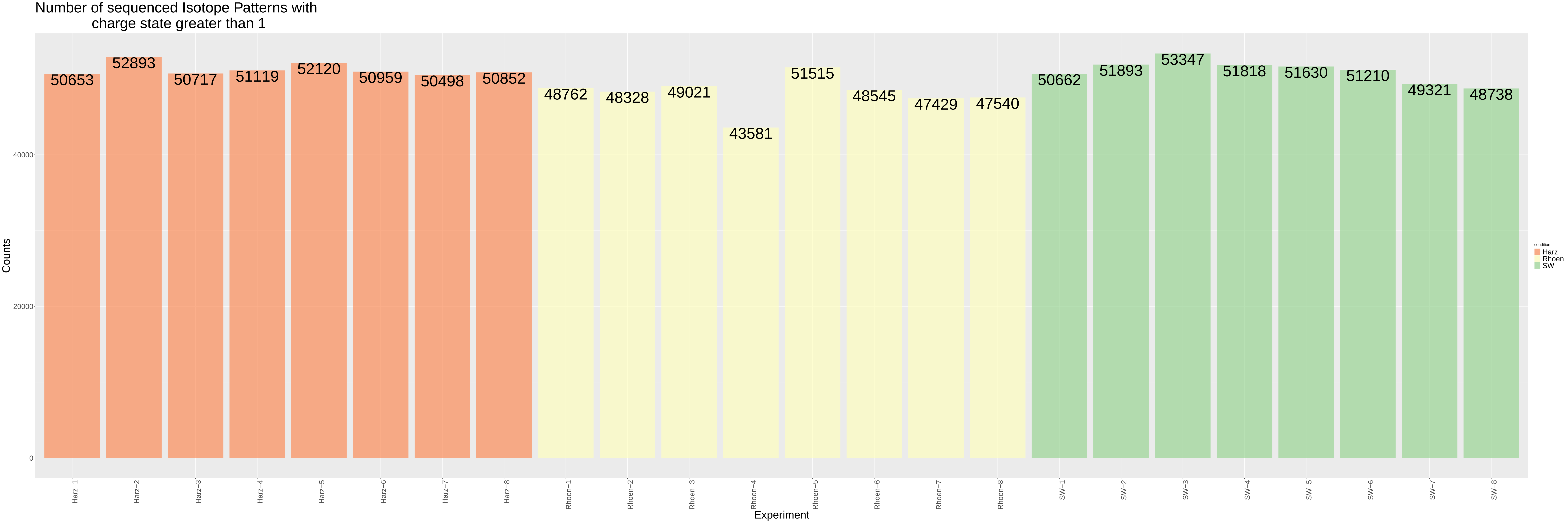

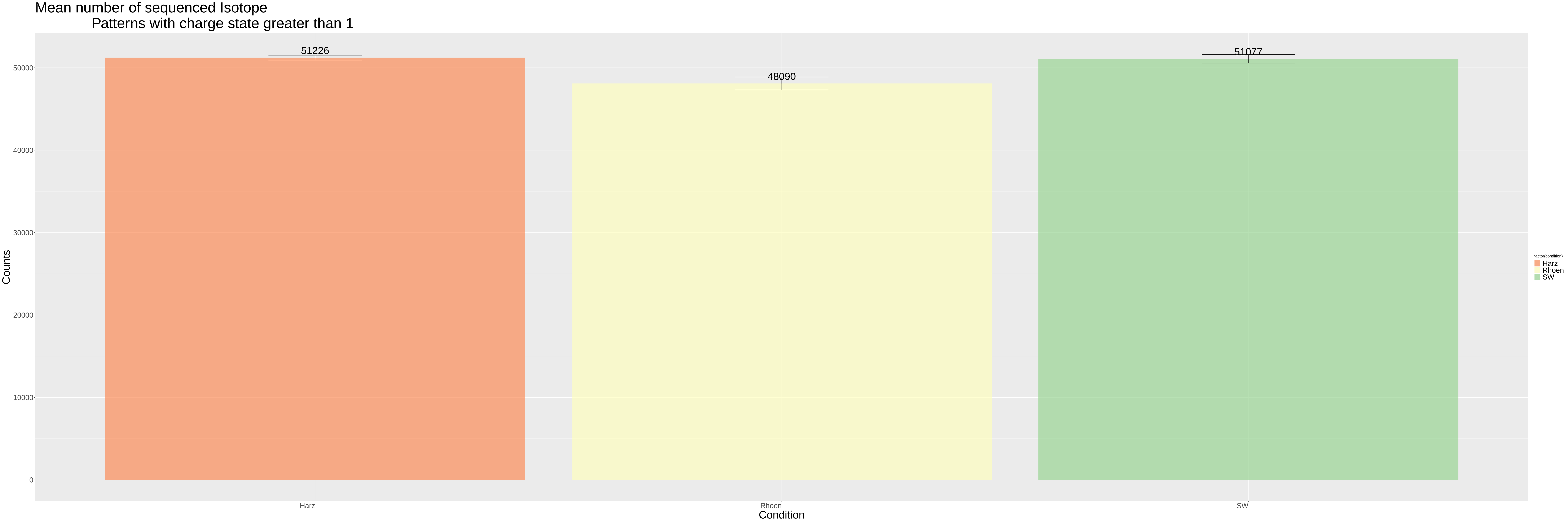

### QCSummary_MS1SCANS.pdf

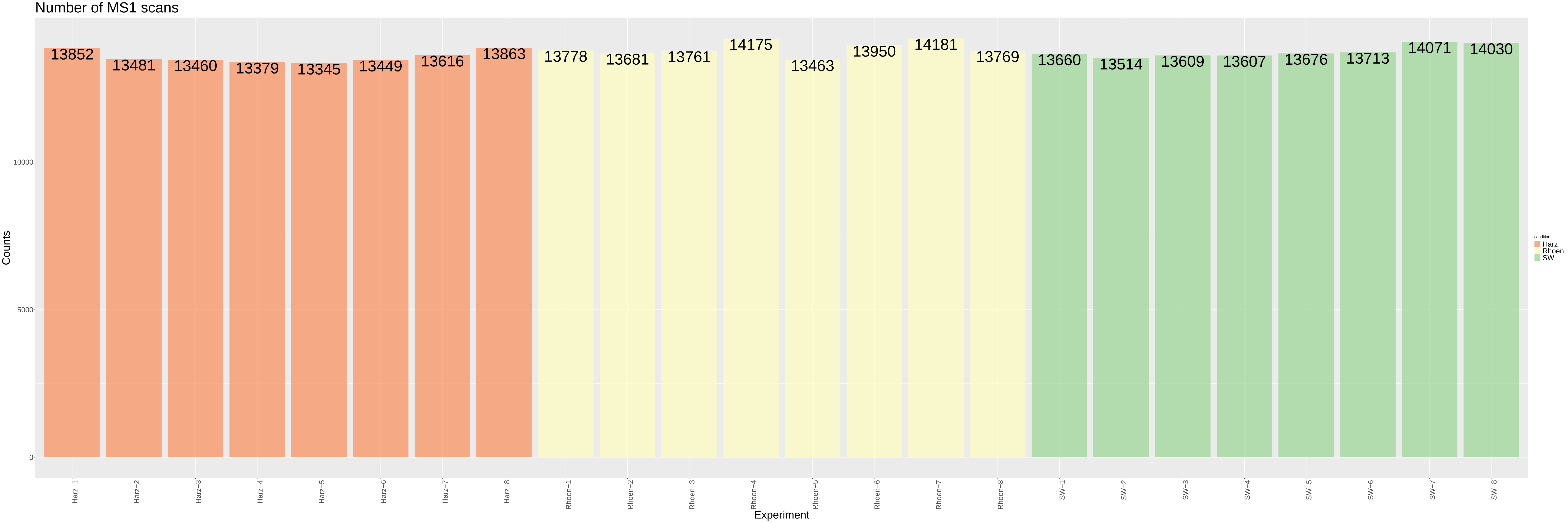

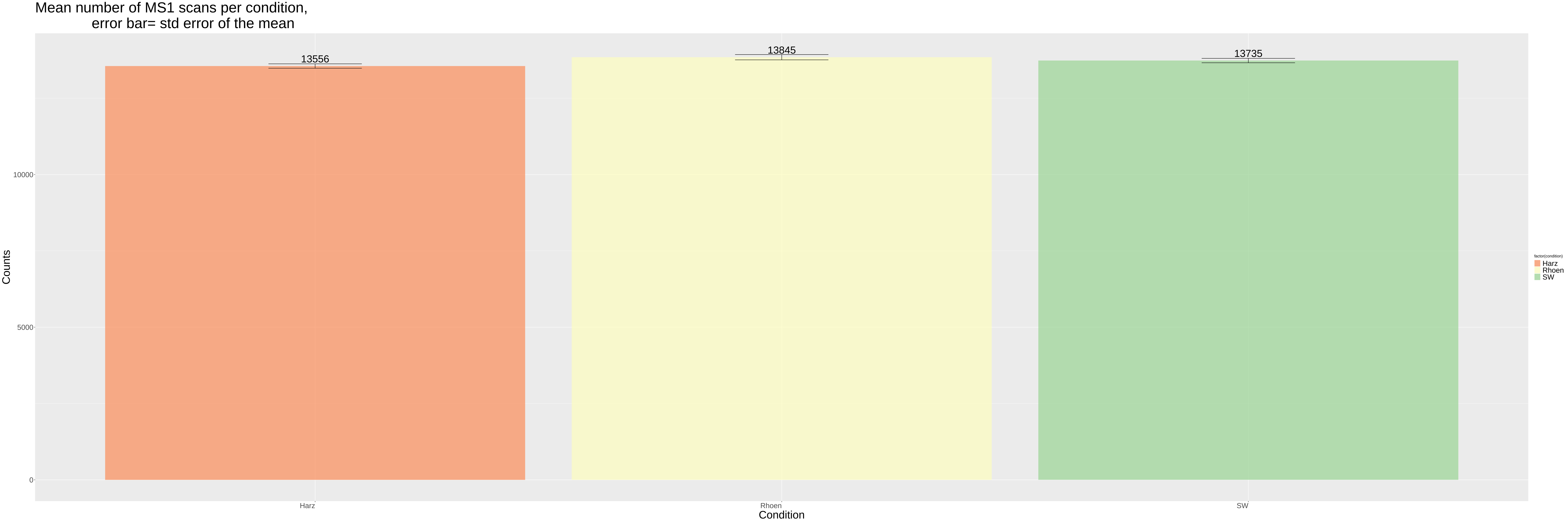

### QCSummary_MS2.pdf

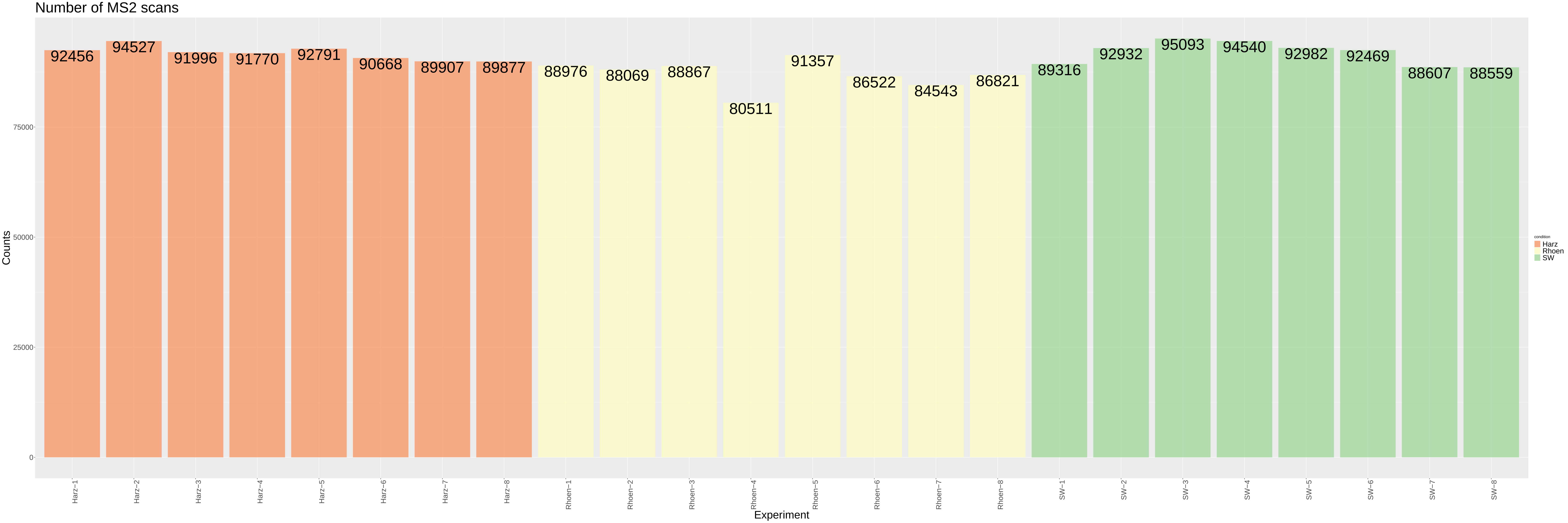

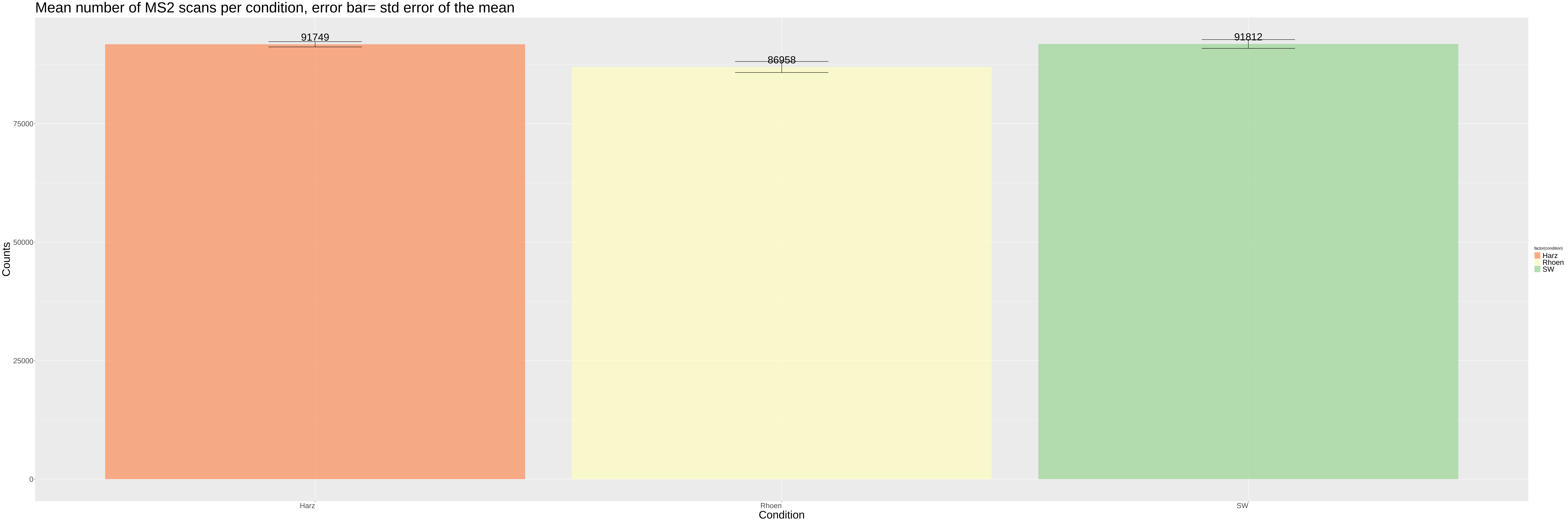

Number of MS1 and MS2 scans

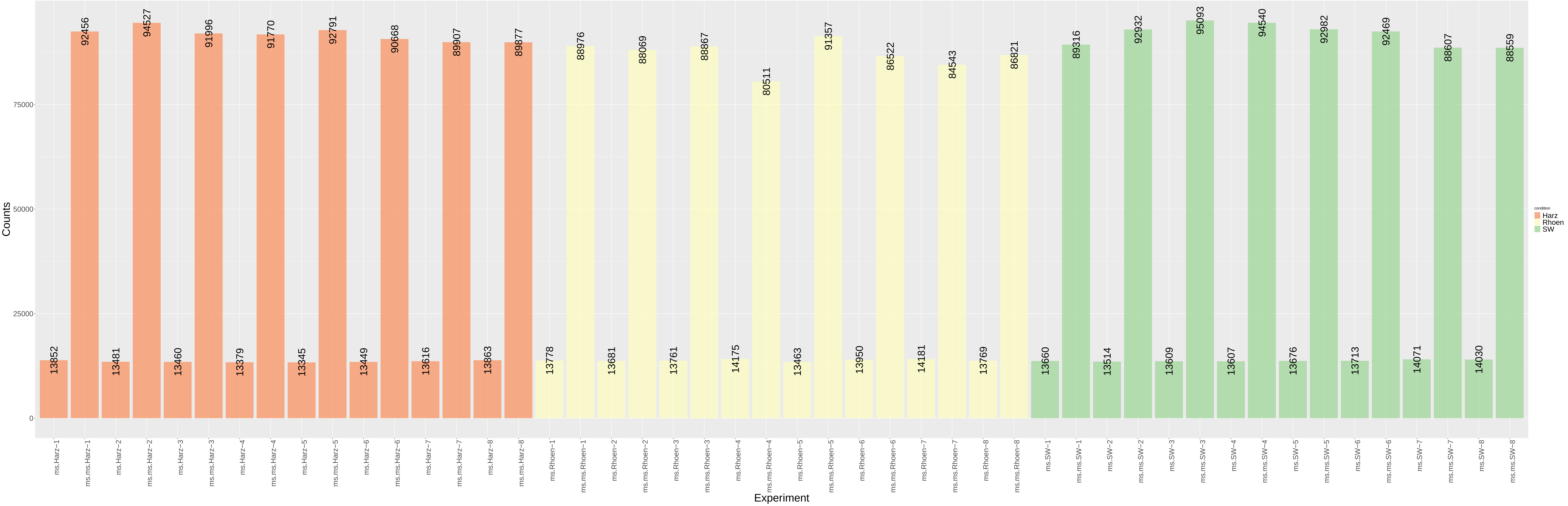

### QCSummary_MSMS.pdf

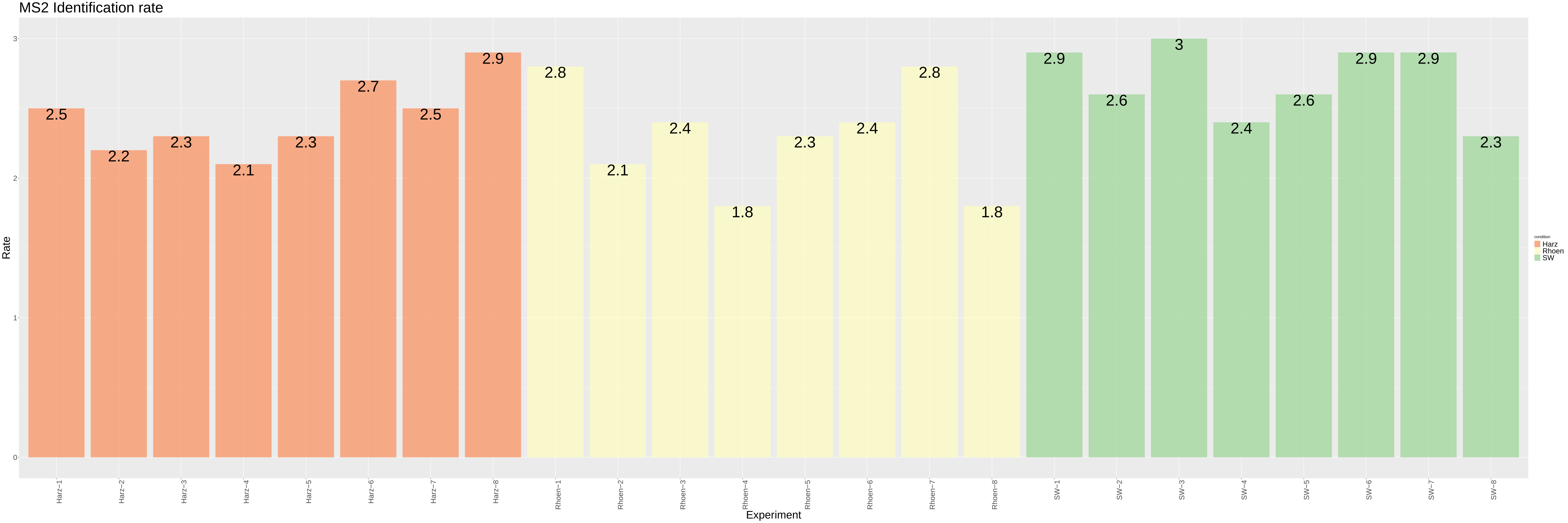

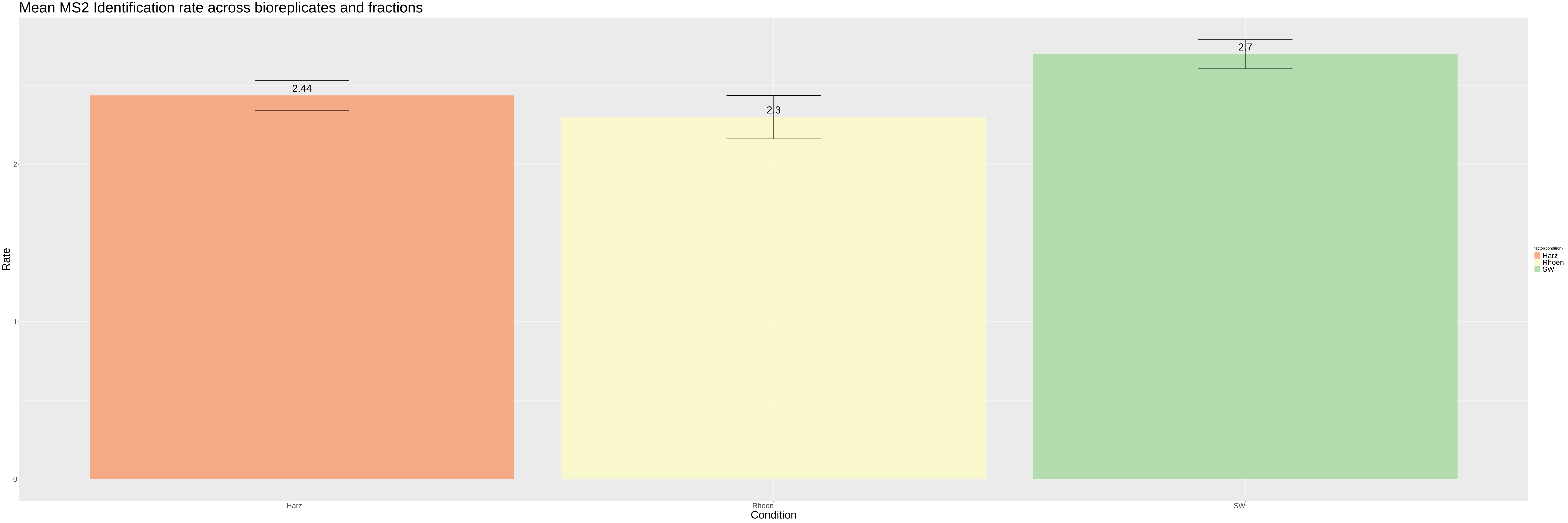
