## Supplemental Information 5 for "Abiotic drivers of protein abundance variation among natural populations": SI5.R

```
#### Code for "Abiotic drivers of protein abundances in natural populations" ####

#### Load Packages ####

library(pacman)
pacman::p_load(dplyr, preprocessCore, impute, ggeffects, rstanarm, ggplot2,
               bayesplot, vegan, WGCNA, tidyverse, flashClust, topGO,
               lintr, lattice, raster, sp, Hmisc, corrplot, caret, plyr,
               forcats, scales) 

# Read, transform, normalize and impute raw protein intensity data (SI2, spreadsheet 1 "Protein intensities raw")
expr <- read.csv("data.csv", header = TRUE, row.names = 1, sep = ";")

# Format fasta headers (first column) to only include the first Majority protein identifier
expr$V1 = sub(";.*", "", expr$V1) # Deletes everything after ";"
expr$V1 <- sub(" .*", "", expr$V1) # Deletes "[NA..]"

# Rename first column header
expr <- expr %>% 
  rename(
    Fasta = V1
  )

# Quantile normalization
expr_long <- data.frame(t(expr)) # Transpose matrix
expr_long_matrix <- as.matrix(expr_long) # Set to "matrix" object
expr_long_norm <- normalize.quantiles(expr_long_matrix) # perform quantile normalization
expr_long_norm <- as.data.frame(expr_long_norm) # set as data.frame

#### Log2 transformation ####
expr_long_norm_log <- log2(expr_long_norm)

#### Data imputation ####
sum(is.na(expr_long_norm_log)) # check how many missing values are in the data
expr_matrix2 <- as.matrix(expr_long_norm_log) # set as matrix
expr_imp <- impute.knn(expr_matrix2, k = 10, rowmax = 0.8, colmax = 0.8, maxp = 1500, rng.seed=362436069) # impute missing values using the KNN algorithm
expr2 <- expr_imp$data # get the imputed data in a new object
expr2 <- as.data.frame(expr2) # set as data.frame

#### Non-metric multidimensional scaling ####
expr_dist <- vegdist(expr2, method = "bray") # Compute Bray-Curtis distance matrix

nMDS = metaMDS(expr_dist, k=2, distance = "bray",
               maxit = 999, trymax = 500, wascores = TRUE) # Perform nMDS

stressplot(nMDS) # Check nMDS model fit

# Plot the nMDS
region=c(rep("Rhoen",8),rep("Harz",8),rep("Black Forest",8)) # Make group-vector
ordiplot(NMDS,type="n")
ordihull(NMDS,groups=region,draw="polygon",col="grey90",label=F)
points(NMDS$species, col="grey",cex=0.6, pch=21)
orditorp(NMDS,display="sites",col=c(rep("cadetblue3",8),rep("chocolate1",8),rep("chartreuse4",8)),
         air=0.1,cex=1.25)

Axis2_scores <- scores(NMDS$species, choices=c(1,2)) # get the axis scores for species (proteins)


#### WGCNA (this script was adapted from https://horvath.genetics.ucla.edu/html/CoexpressionNetwork/Rpackages/WGCNA/) ####

options(stringsAsFactors = FALSE)
dim(expr2)

gsg = goodSamplesGenes(expr2, verbose = 3) # Run this check if there are protein outliers
gsg$allOK # If the statement returns TRUE, all proteins have passed the cuts
table(rownames(env)==rownames(expr2)) # Check if datasets align; must return "TRUE"

A = adjacency(t(expr2),type="signed") # this calculates the whole network connectivity
k = as.numeric(apply(A,2,sum))-1 # standardized connectivity
Z.k = scale(k)
thresholdZ.k = -2.5
outlierColor = ifelse(Z.k<thresholdZ.k,"red","black")

sampleTree = flashClust(as.dist(1-A), method = "average")

# Convert traits to a color representation where red indicates high values
traitColors = data.frame(numbers2colors(env,signed=FALSE))
dimnames(traitColors)[[2]] = paste(names(env))
datColors = data.frame(outlier = outlierColor,traitColors)

plotDendroAndColors(sampleTree,groupLabels=names(datColors),
                    colors=datColors,main="Sample Dendrogram and Trait Heatmap")

powers = c(c(1:10), seq(from =10, to=30, by=1))
sft = pickSoftThreshold(Expr, powerVector=powers, verbose =5, networkType="signed") #call network topology analysis function

plot(sft$fitIndices[,1], -sign(sft$fitIndices[,3])*sft$fitIndices[,2], xlab= "Soft Threshold (power)", ylab="Scale Free Topology Model Fit, signed R^2", type= "n", main= paste("Scale independence"))
text(sft$fitIndices[,1], -sign(sft$fitIndices[,3])*sft$fitIndices[,2], labels=powers, col="red")
abline(h=0.90, col="red")

plot(sft$fitIndices[,1], sft$fitIndices[,5], xlab= "Soft Threshold (power)", ylab="Mean Connectivity", type="n", main = paste("Mean connectivity"))
text(sft$fitIndices[,1], sft$fitIndices[,5], labels=powers, col="red")

softPower = 18
adjacency = adjacency(expr2, power = softPower, type = "signed") #specify network type

TOM = TOMsimilarity(adjacency, TOMType="signed") # specify network type
dissTOM = 1-TOM

geneTree = flashClust(as.dist(dissTOM), method="average")

plot(geneTree, xlab="", sub="", main= "Gene Clustering on TOM-based dissimilarity", labels= FALSE, hang=0.04)

minModuleSize = 30
dynamicMods = cutreeDynamic(dendro= geneTree, distM= dissTOM, deepSplit=2, pamRespectsDendro= FALSE, minClusterSize = minModuleSize)

dynamicColors= labels2colors(dynamicMods)
MEList= moduleEigengenes(expr2, colors= dynamicColors,softPower = 18)
MEs= MEList$eigengenes
MEDiss= 1-cor(MEs)
METree= flashClust(as.dist(MEDiss), method= "average")

save(dynamicMods, MEList, MEs, MEDiss, METree, file= "Network_allSamples_signed_RLDfiltered.RData")

plot(METree, main= "Clustering of module eigengenes", xlab= "", sub= "")

MEDissThres = 0.2
merge = mergeCloseModules(expr2, dynamicColors, cutHeight= MEDissThres, verbose =3)
mergedColors = merge$colors
mergedMEs = merge$newMEs

plotDendroAndColors(geneTree, cbind(dynamicColors, mergedColors), c("Dynamic Tree Cut", "Merged dynamic"), dendroLabels= FALSE, hang=0.03, addGuide= TRUE, guideHang=0.05) # plot dendrogram with module colors below it

moduleColors = mergedColors
colorOrder = c("grey", standardColors(50))
moduleLabels = match(moduleColors, colorOrder)-1
MEs = mergedMEs

nGenes = ncol(expr2)
nSamples = nrow(expr2)

#Recalculate MEs with color labels
MEs0 = moduleEigengenes(expr2, moduleColors)$eigengenes
MEs = orderMEs(MEs0)
moduleTraitCor = cor(MEs, env, use= "p")
moduleTraitPvalue = corPvalueStudent(moduleTraitCor, nSamples)

#Print correlation heatmap between modules and traits
textMatrix= paste(signif(moduleTraitCor, 2), "\n(", 
                  signif(moduleTraitPvalue, 1), ")", sep= "")
dim(textMatrix)= dim(moduleTraitCor)
par(mar= c(6, 8.5, 3, 3))

labeledHeatmap(Matrix= moduleTraitCor, 
               xLabels= names(env), 
               yLabels= names(MEs), 
               ySymbols= names(MEs), 
               colorLabels= FALSE, 
               colors= blueWhiteRed(50), 
               textMatrix= textMatrix, 
               setStdMargins= FALSE, 
               cex.text= 0.5, 
               zlim= c(-1,1), 
               main= paste("Module-environment relationships"))

# Save the network data for SI

modNames = substring(names(MEs), 3)
geneModuleMembership = as.data.frame(cor(expr2, MEs, use = "p"))
MMPvalue = as.data.frame(corPvalueStudent(as.matrix(geneModuleMembership), nSamples))
names(geneModuleMembership) = paste("MM", modNames, sep="")
names(MMPvalue) = paste("p.MM", modNames, sep = "")
geneTraitSignificance = as.data.frame(cor(expr2, env, use = "p"))
GSPvalue = as.data.frame(corPvalueStudent(as.matrix(geneTraitSignificance), nSamples))
names(geneTraitSignificance) = paste("GS.", names(env), sep="")
names(GSPvalue) = paste("p.GS.", names(env), sep = "")
ProteinInfo_Final = data.frame(moduleColor = moduleColors,
                               geneTraitSignificance,
                               GSPvalue,
                               geneModuleMembership)
write.csv(ProteinInfo_Final, file = "Global_WGCNA_Results_Final.csv")


#### GLM example for individual proteins (example: Cytochrome_P4506 (GAXX02001139.1.p1) and elevation) ####

protein_of_interest <- expr2$GAXX02001139.1.p1 # Exract abundance values using the fasta header
elevation <- env$elevation # Get the elevation data for springs
poie <- data.frame(protein_of_interest, elevation) # put them together into a data.frame

poie$Region <- c("Rhoen", "Rhoen", "Rhoen", "Rhoen", "Rhoen", "Rhoen", "Rhoen", "Rhoen",
                    "Harz", "Harz", "Harz", "Harz", "Harz", "Harz", "Harz", "Harz",
                    "Black Forest", "Black Forest", "Black Forest", "Black Forest",
                    "Black Forest", "Black Forest", "Black Forest", "Black Forest") # Add the sampling region (origin of samples)

fit1 <- stan_glm(protein_of_interest ~ elevation + Region, data = poie) # Model fit
dat1 <- ggpredict(fit1, terms = c("elevation", "Region")) # data of model fit

p1 <- plot(dat1, rawdata = TRUE, ci.style = "ribbon", alpha = 0.1, jitter = 0.2,
           colors = c("cadetblue3", "chocolate1", "chartreuse4")) +
  labs(
    x = "Elevation (m.a.s.l)",
    y = expression(log[2]~protein~intensity),
    title = "Cytochrome P450 6a2 (3A4)"
  ) # Plot the regression


#### GO enrichment ####

# Read proteins of a certain module etc. (fasta headers in tab-delimited file (.txt or .csv))
blackModule <- read.csv("blackModule.csv", sep = ",", header = TRUE)

black <- blackModule %>%
  dplyr::select(Protein, p.GS.Oxygen) # Sort by gene significance for the abiotic variable that is significantly associated with the module

## Sort by Gene Significance p-value:

black_sorted <- black[order(black$p.GS.Oxygen), ]

# The background "GO_database.txt" is a .txt file giving all GO terms (separated by ";") for all proteins identified in the study
gene_to_go_mapping <- readMappings(file = "GO_database.txt", sep = "\t", IDsep = ",")

## Convert into topgo's genelist format:
topgo_genelist <- black_sorted$p.GS.Oxygen
names(topgo_genelist) <- black_sorted$Protein

# Define a cut-off for running fisher's exact test:  

cutoff_for_top_fivepercent <- quantile(x = topgo_genelist, probs = 0.05)

### "CC", "MF", "BP" = "Cellular Compartment", "Molecular Function", "Biological Process"

for (go_category in c("CC")) {
  # STEP TWO
  ## Build the GOdata object in topGO
  my_go_data <- new("topGOdata",
                    description = paste("GOtest", go_category, sep = "_"),
                    ontology    = go_category,
                    geneSel     = function(x) {
                      # fails to run without thi
                      return(x <= cutoff_for_top_fivepercent)
                    },
                    allGenes    = topgo_genelist,
                    gene2GO     = gene_to_go_mapping,
                    annot       = annFUN.gene2GO,
                    nodeSize    = 100) # Modify to reduce/increase stringency.
  # STEP THREE
  ## Calculate ks test using 'weight01' algorithm:
  result_weight_ks <- runTest(object    = my_go_data,
                              algorithm = "weight01",
                              statistic = "ks")
  ## Calculate fisher exact test using 'weight01' algorithm:
  result_weight_fisher <- runTest(object    = my_go_data,
                                  algorithm = "weight01",
                                  statistic = "fisher")
  ## Combine results from statistical tests:
  result_weight_output <- GenTable(object    = my_go_data,
                                   weight_ks = result_weight_ks,
                                   weight_fisher = result_weight_fisher,
                                   orderBy   = "weight_ks",
                                   topNodes  = length(score(result_weight_ks)))
  ## Correct ks test for multiple testing:
  result_weight_output$weight_ks <- as.numeric(result_weight_output$weight_ks)
  result_weight_output$weight_fisher <- as.numeric(result_weight_output$weight_fisher)
  result_weight_output$weight_ks_adjusted <- p.adjust(p = result_weight_output$weight_ks,
                                                      method = c("fdr"))
  result_weight_output$weight_fisher_adjusted <- p.adjust(p = result_weight_output$weight_fisher,
                                                          method = c("fdr"))
  ## Subset calls with significance higher than expected:
  result_weight_output_sig <- subset(x      = result_weight_output,
                                     subset = (Significant > Expected) &
                                       (weight_ks < 0.1))
  ## Update column names:
  colnames(result_weight_output_sig)[6] <- gsub(" ", "_",
                                                colnames(result_weight_output_sig)[6])
  ## For significant terms, add an additional field called 'category' which
  ## will be used for plotting of individual go category:
  result_weight_output_sig$category <- go_category
  ## Remove gaps between Terms, which can cause downstream problems:
  result_weight_output_sig$Term <- gsub(" ", "_", result_weight_output_sig$Term)
}


#### Get BioClim variables ####

# Longitude and latitude values of springs
lat <- c(50.613586, 50.599372, 50.588919, 50.465871,
         50.503505, 50.517887, 50.609672, 50.599378,
         51.637179, 51.702130, 51.767917, 51.767861,
         51.742708, 51.742801, 51.726299, 51.647887,
         47.652422, 47.652422, 47.652778, 47.597678,
         47.662372, 47.656950, 47.655839, 47.658853)

long <- c(9.764174, 9.999089, 10.004149, 10.00545,
          9.929572, 9.932069, 9.6454665, 9.669365,
          10.40629, 10.34304, 10.610061, 10.61202,
          10.67571, 10.67580, 10.564328, 10.74177,
          7.975453, 7.975456, 7.9736110, 7.840197,
          7.901392, 7.907233, 7.7676080, 7.770014)

site <- c("R1", "R2", "R3", "R5", "R6", "R7", "R8", "R9",
          "H1", "H2", "H3", "H4", "H5", "H6", "H7", "H9",
          "BF1", "BF2", "BF3", "BF4", "BF5", "BF6", "BF7", "BF8")

data <- data.frame(long, lat, row.names = site)

# Get the BioClim data with resulution 0.5 (minutes of a degree)
r <- getData("worldclim",var="bio",res=0.5)

values <- extract(r,data)
df <- cbind.data.frame(coordinates(data),values)

## Correlations between Clim variables
Clim_rcorr <- rcorr(as.matrix(Clim))
corrplot(Clim_cor)

## Remove variables highly correlation (>0.8)
Clim_cor <- cor(Clim)
hc = findCorrelation(Clim_cor, cutoff = 0.8)
hc = sort(hc)
reduced_Clim = Clim_cor[,-c(hc)]


#### Protein Families (Pfam) frequency plot (the same code holds true for the COG family frequencies) ####

# Read the InterProScan file (this is the output file of the IPS for ALL proteins identified in the study)
IPS <- read.csv("IPS.csv", sep = ";", header = TRUE)
IPS <- as.data.frame(IPS)

# Parse from DAP.txts all IDs from IPS df
HBF <- read.table("H_BF_List.txt", header = FALSE) # This would be a tab-delimited .txt file containing the fasta headers of all DAPs identified between H and BF

# Extract information in IPS on DAPs in HBF
Hbf_Ips <- as.data.frame(IPS[IPS$FASTA.header %in% HBF$V1,])

# Plot the frequencies of IPS information for the differentially abundant proteins
Pfam <- "Pfam" # For choosing Pfam from IPS annotations
pfam_hbf <- hbf_down_ips[hbf_ips$Analysis %in% Pfam, ] # Choose all pfam terms from data.frame
pfam_hbf2 <- pfam_down_hbf[!(is.na(pfam_hbf$InterPro.description) | pfam_hbf$InterPro.description==""), ] # Delete all NAs
pfam_hbf_unique <- distinct(pfam_hbf2, FASTA.header, .keep_all = TRUE) # Select only one observation of each protein

# Plot the frequency plot
ggplot(data=pfam_hbf_unique, aes(x=forcats::fct_infreq(InterPro.description))) + 
  geom_bar(aes(y = (..count..)/sum(..count..)), fill = "grey") +
  theme_classic() +
  coord_flip() +
  scale_y_continuous(labels = percent) +
  ylab("Relative frequency of Pfam families in DAPs between Harz and Black Forest") +
  xlab("")
```
